# GLIS1 regulates trabecular meshwork function and intraocular pressure and is associated with glaucoma in humans

**DOI:** 10.1101/2021.07.02.450719

**Authors:** K. Saidas Nair, Chitrangda Srivastava, Robert V. Brown, Swanand Koli, Hélène Choquet, Hong Soon Kang, Yien-Ming Kuo, Sara A. Grimm, Caleb Sutherland, Alexandra Badea, G. Allan Johnson, Yin Zhao, Jie Yin, Kyoko Okamoto, Graham Clark, Teresa Borras, Gulab Zode, Krishnakumar Kizhatil, Subhabrata Chakrabarti, Simon W.M. John, Eric Jorgenson, Anton M. Jetten

## Abstract

Chronically elevated intraocular pressure (IOP) is the major risk factor of primary open-angle glaucoma, a leading cause of blindness. Dysfunction of the trabecular meshwork (TM), which controls the outflow of aqueous humor (AqH) from the anterior chamber, is the major cause of elevated IOP. Here, we demonstrate that mice deficient in the Krüppel-like zinc finger transcriptional factor GLI-similar-1 (GLIS1) develop chronically elevated IOP. Magnetic resonance imaging and histopathological analysis reveal that deficiency in GLIS1 expression induces progressive degeneration of the TM, leading to inefficient AqH drainage from the anterior chamber and elevated IOP. Transcriptome and cistrome analyses identified several glaucoma- and extracellular matrix-associated genes as direct transcriptional targets of GLIS1. We also identified a significant association between *GLIS1* variant rs941125 and glaucoma in humans (*P*=4.73x10^-6^), further supporting a role for *GLIS1* into glaucoma etiology. Our study identifies GLIS1 as a critical regulator of TM function and maintenance, AqH dynamics, and IOP.

## Introduction

Glaucoma is a heterogeneous group of progressive optic neuropathies characterized by the degeneration of the optic nerve that results in irreversible blindness^1^. Primary open-angle glaucoma (POAG) and primary angle closure glaucoma (PACG) are the two most common forms of glaucoma in adults^2–5^, while primary congenital glaucoma (PCG) accounts for up to 18% of childhood blindness^6^. Age, ethnicity, gender, environmental and genetic factors all contribute to glaucoma susceptibility^7–9^. However, elevated intraocular pressure (IOP) is the major causal risk factor for glaucoma.

Normal IOP is required to maintain proper physiological function of the eye and also to maintain the structure of the globe of the eye. The maintenance of homeostatic IOP is critically dependent on the balance between the inflow and outflow of aqueous humor (AqH)^10^. AqH is secreted by the ciliary body into the ocular anterior chamber (AC) where it nourishes avascular tissues. The AqH subsequently exits through specialized drainage structures located at the junction where the iris meets the cornea (iridocorneal angle). The ocular drainage structures are primarily composed of the trabecular meshwork (TM) and Schlemm’s canal (SC). AqH first flows through the TM into the SC and subsequently enters the episcleral veins before returning back to the systemic circulation^10–12^. TM dysfunction has been causally linked to impaired AqH drainage (increased outflow resistance) and elevated IOP^10, 13–15^.

An increasing number of rare mutations and common genetic variants in a variety of genes, including *MYOC*, *CYP1B1*, *GLIS3*, *LOXL4*, *LTBP2*, *PITX2*, and *OPTN,* have been associated with elevated IOP and different types of glaucoma ^4, 5, 9, 16–24^.

GLI-Similar 1 (GLIS1), together with GLIS2 and -3, comprise a subfamily of Krüppel-like zinc finger (ZF) transcriptional factors^25–27^. In contrast to GLIS2 and GLIS3, relatively little is known about the physiological functions of GLIS1. To obtain greater insights into the biological roles of GLIS1, we analyzed *Glis1*-KO mice for phenotypic alterations and found that these mice develop an enlarged eye phenotype.

In this study, we examine the function of GLIS1 in ocular tissues in more detail and demonstrate that GLIS1 plays a critical regulatory role in maintaining normal TM structure and IOP. We show that GLIS1 is expressed in the TM, a tissue critical in the regulation of outflow resistance. Deficiency in *GLIS1* induces progressive degeneration of the TM, leading to inefficient AqH drainage and elevated IOP. To obtain insights into potential mechanisms that may underlie this phenotype, changes in the expression of target genes were examined. Combined RNA-Seq and ChIP-Seq analyses identified a number of genes that are directly regulated by GLIS1, including *MYOC, CYP1B1, LOXL4*, and *LTBP2*, genes previously implicated in glaucoma^6, 28^. Importantly, we have detected significant associations between common genetic variants in the *GLIS1* region and POAG in humans, thereby supporting the role of *GLIS1* as a glaucoma risk gene. These variants may impact TM functions and compromise AqH drainage thereby contributing to elevated IOP and glaucoma.

## Results

### Identification of GLIS1 physiological functions

To obtain insights into the physiological functions of GLIS1^29, 30^, *Glis1*-KO mice were examined for any potential phenotypic alterations. In these mice most of exon 4 (840 bp) was replaced by lacZ containing three Stop codons (lacZ-Stop3) generating a fusion protein (GLIS1N-βGal) consisting of the N-terminus of GLIS1 and β-galactosidase (β- Gal). This protein lacks the entire DNA-binding domain (DBD) and C-terminus of GLIS1, including its transactivation domain (TAD)(Supplementary Figure 1a). The fusion protein was undetectable by immunohistochemical staining for β-Gal (1:1000, PR-Z3781, Promega) in several tissues suggesting that it may be proteolytically degraded. Reporter transactivation analysis demonstrated that mutations in the ZF motifs that abolish their tetrahedral configuration, and deletion of the C-terminal TAD greatly decreased or fully abolished GLIS1 transcriptional activity (Supplementary Figure 1b). These data indicate that loss of the ZFs and TAD in *Glis1*-KO mice abolish the ability of GLIS1 to recognize the GLIS binding site (GLISBS) and to regulate the transcription of target genes. Supporting the specificity of the *Glis1*-KO, deletion of exon 4 had no significant effect on the expression of *Dmrtb1, Slc1a7, Dio1,* and *Cpt2*, genes neighboring *Glis1,* nor the expression of *Glis2* and *Glis3* in *Glis1*-KO kidneys and testes (Supplementary Figure 1c and d).

Evaluation of 1-6 months C57BL/6NCrl *Glis1*-KO mice revealed that these mice developed enlarged eyes (Figure 1a), while no other obvious abnormalities were observed. Similarly, no eye enlargement was observed in 129S6/SvEvTac *Glis1*-KO mice. Male and female KO mice in both backgrounds noticeably developed this abnormal eye phenotype between 2-3 months of age, which became more pronounced with age. Because protruding eyes are a well-established comorbidity commonly associated with Graves’ disease, an autoimmune disease leading to hyperthyroidism^31^, and since GLIS1 and GLIS3 family members have been implicated in several thyroid gland-associated diseases^25, 27, 32, 33^, we examined whether this *Glis1*-KO phenotype was related to the development of Graves’ disease that is characterized by high circulating levels of T3/T4 and low TSH. However, our analysis of serum T3, T4 and TSH showed that their levels were not significantly different between WT and *Glis1*-KO C57BL/6NCrl mice indicating that this phenotype was not related to the development of Graves’ disease (Supplementary Figure 2).

**Figure 1.**
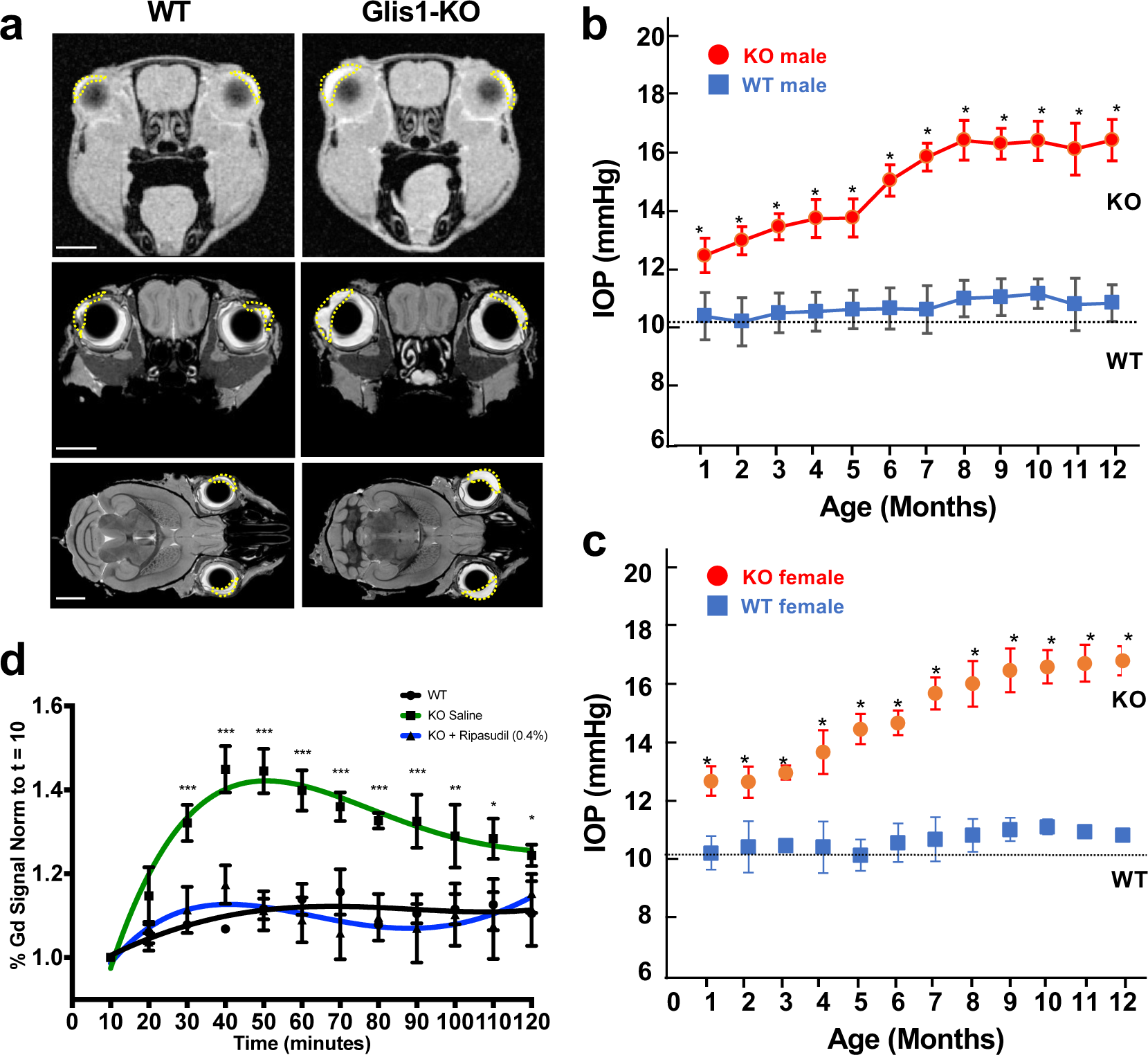
Anterior chamber is enlarged, IOP elevated, and AqH dynamics altered in *Glis1*-KO mice. **a**. Representative MRI images from 2.5 months old WT and *Glis1*-KO mice showing increased size of the anterior chamber in *Glis1*-KO mice compared to WT. The upper two images are *in vivo* images acquired by dynamic contrast enhancement in the eye. The lower four images are coronal and sagittal sections of eyes from fixed specimen stained with gadolinium using active staining. The anterior chamber is outlined by the dotted line. Scale bar = 2 mm. **b and c.** Comparison of IOP levels in male (**b**) and female (**c**) WT (squares) and *Glis1*-KO (circles) mice as function of age. Male mice examined: at 1 and 2 months (n=3); 3 (n=5); 4 and 7 (n=8); 5 (n=6); 6, 11, and 12 months (n=7). For female mice: 1 and 2 months (n=10); 3-7, 9 and 10 months (n=4); 8 (n=5); 11 and 12 months (n=3). IOP data from left and right eyes were combined and 4 IOP measurements/eye/timepoint were performed; thus, total of 24-80 measurements at each timepoint. Data are represented as means ± SD. Statistical analyses were performed with two-tailed Student’s t-test. *p<10^-5^. Dotted line indicates basal IOP level in 1-3 months old mice. **d.** AqH dynamics was examined by Gd-enhanced MRI over a 2 h period. Percent of Gd signal enhancement was determined and plotted. Left eye of 2 months-old WT mice (n=3; black line) and *Glis1*-KO mice (n=4; red line) was treated topically with saline and the right eye of *Glis1*-KO mice with Ripasudil (0.04%)(n=4; blue line). Data are represented as means ± SD). Statistical analyses were performed with two-tailed Student’s t-test. *p<10^-2^; **p<10^-3^; ***p<10^-4^.

### Intraocular Pressure is elevated in *Glis1*-KO mice

To examine whether the enlarged eye phenotype was associated with anatomic changes in intra- and periocular tissues, we performed Gadolinium magnetic resonance imaging (Gd-MRI) on formalin fixed specimen from 2.5-months-old WT and *Glis1*-KO C57BL/6NCrl mice. The contrast agent Gd has been used to assess ocular anatomy by MRI^34, 35^. Analysis of multiple MRI images through the head revealed that there was little difference between the size of the periocular tissues in *Glis1*-KO relative to WT littermates in all 3 orientations. Importantly, we consistently observed an enlargement of the AC in both the right and left eyes of the *Glis1*-KO mice (Figure 1a). The enlargement of AC observed in the *Glis1*-KO mice might be due to defective AqH drainage causing increased AqH accumulation that leads to the observed elevated IOP.

To obtain further support for this hypothesis, we measured IOP in WT and *Glis1*-KO mice over a 12-month time period. Our data demonstrated that IOP is significantly elevated in male as well as female *Glis1*-KO mice relative to age and gender matched WT littermates (Figure 1b, c). An increase in IOP was observed as early as in 1-month-old mice and then steadily increased before plateauing at 8 months. Further analysis revealed that the progressive, age-dependent increase in IOP was similar between left and right eyes (Supplementary Figure 3). Together, these observations suggested a role for GLIS1 in the regulation of IOP.

### Decreased AqH outflow in eyes from GLIS1-deficient mice

Elevated IOP is most commonly caused by outflow resistance. To determine whether the elevated IOP in *Glis1*-KO C57BL/6NCrl mice was due to changes in AqH drainage, we employed dynamic contrast enhanced MRI to evaluate AqH dynamics *in vivo*. The contrast agent Gd present in the AC enhances the T1-weighted MRI signal brightness and serves as a tracer, thereby providing a readout for AqH accumulation and outflow^34, 35^. Following administration of Gd, 2 months-old mice were scanned for 2 h at 10 min intervals and Gd accumulation in the anterior chamber was measured relative to the initial baseline (See image source files at: https://civmvoxport.vm.duke.edu/voxbase/studyhome.php?studyid=733). Our data indicated that in WT mice Gd is readily cleared from the eye (Figure 1d). A significant (29%; P<0.0001) increase in Gd accumulation in the anterior chamber was detected in *Glis1*-KO eyes as compared to the WT eyes suggesting impaired AqH exit. It is well-established that a major route of AqH exit from the anterior chamber is via the conventional drainage pathway comprising the TM and SC^10^. To obtain further support for this hypothesis, we evaluated the AqH dynamics in *Glis1*-KO mice following topical administration (5 μl, 0.4%) of Ripasudil. This drug functions as an IOP lowering Rho-kinase inhibitor that enhances AqH outflow via the TM and SC^36^. As shown in Figure 1d, Ripasudil treatment significantly reduced Gd accumulation in the ocular anterior chamber of *Glis1*-KO mice as compared to treatment with isotonic saline in the contralateral eye consistent with its IOP lowering effects. Our data suggests that the increased IOP observed in the *Glis1*-KO mice correlates with reduced AqH outflow and might involve dysfunction of the TM, a major cause of elevated IOP and glaucoma.

### Progressive disruption of ocular drainage structures in Glis1-KO mice

To determine whether structural and morphological changes of ocular drainage tissues in *Glis1*-KO mice might underlie TM dysfunction and elevated IOP, we performed a detailed ocular histological examination of WT, *Glis1*-heterozygous and *Glis1*-KO mice maintained in a C57BL/6NCrl strain. Ocular angle structures of the *Glis1*-heterozygous mice showed an intact ocular drainage tissue, unlike the *Glis1*-KO that exhibited TM degeneration (Supplementary Figure 4a and b). This lack of phenotype in heterozygous mice is consistent with that these mice did not develop elevated IOP (Supplementary Figure 4c). Histological analyses demonstrated that the angle structures of the *Glis1*-KO eyes initially appear normal. At 3 weeks, no significant difference in TM morphology was observed suggesting that GLIS1 has no major effect on TM development (Figure 2a, b; Supplementary Figure 5). Major phenotypic changes are observed by 6-8 weeks of age (Figure 2c-f). Focal regions of the angles in 6-weeks old *Glis1*-KO mice exhibited thinning of the TM (hypoplasia) in a substantial proportion of mutant eyes (Figure 2d; Supplementary Figure 6), while some local regions lacked discernible TM. Based on histology, the damage to the ocular drainage structure within an eye is quite variable at earlier time points (6 weeks) with some regions appearing much more normal. Such local variability is well documented for other glaucoma genes and may partially explain the relatively modest increase in IOP^37, 38^.

**Figure 2.**
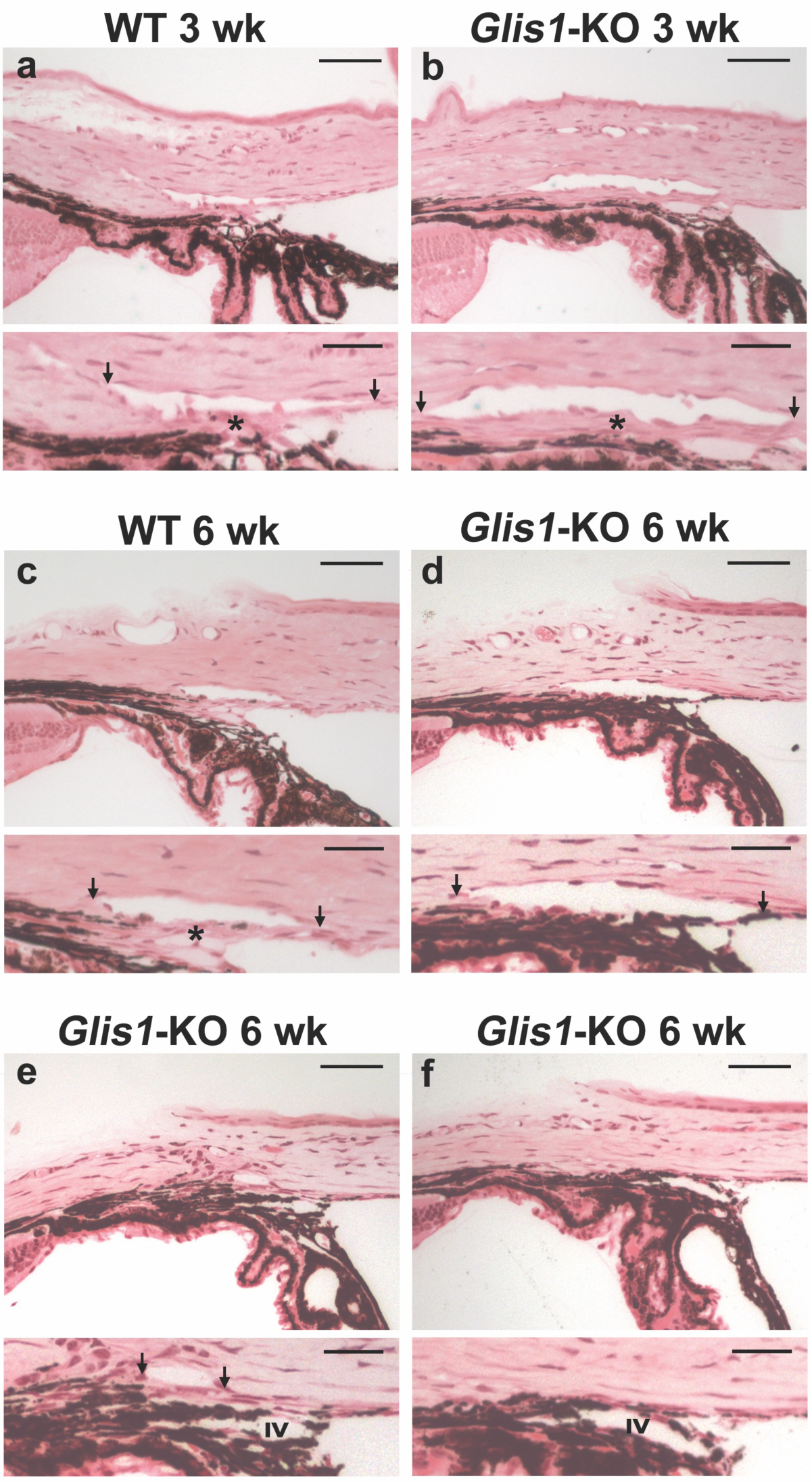
Disruption of the ocular angle drainage structures in *Glis1*-KO mice. **a, c.** WT mice maintained in C57BL/6NCrl background showed a well-developed SC and TM (*) at both 3 and 6 weeks of age. **b.** *Glis1*-KO-C57BL/6NCrl eyes exhibit a morphologically mature ocular drainage tissue at 3 weeks of age. The histological assessment of the ocular angle at 3 weeks of age was performed on 6 WT and 6 *Glis1*- KO eyes with similar results within each group. In contrast, by 6 weeks of age *Glis1*-KO eyes exhibit a variable degree of focal TM degeneration, ranging from hypoplastic TM characterized by substantial thinning of the TM **(d),** reduction in size of the Schlemm’s canal and associated TM causing partial collapse of the ocular drainage structures **(e). f.** At 6 weeks of age a small proportion of mice (<10%) showed focal regions exhibiting partial or complete collapse of the ocular drainage structures lacking TM and Schlemm’s canal. **a-f.** A magnified version of the image in the upper panel is shown in the lower panel. Arrows show edges of the SC. IV, Iris vessel. The histological assessment of the ocular angle at 6 weeks of age was performed on 10 WT and 15 *Glis1*-KO eyes with similar results within each group. Scale bar = 50 μm. Detailed measurement of the TM area was performed on 5 eyes per experimental group, shown in Supplemental figures 5 and 6.

The loss of GLIS1 function did not affect gross morphology of the SC at an early time point (4 weeks) (Supplementary Figure 7a, b) when the TM is still largely intact. However, at later ages (6 weeks and older), but more common at 3 months and older ages, there are regions where the SC becomes partially or completely collapsed (Figure 2e-f). This might be due to a regional or complete degeneration of the TM that may protect the SC from collapse. At older ages (over 6 months), in addition to the degeneration of the TM and collapse of the ocular drainage structures, *Glis1*-KO eyes exhibited anterior synechiae characterized by fusion of the iris and cornea causing angle closure (Supplementary Figure 8a). Besides the observed defects in the ocular drainage structures, no gross abnormalities were observed in other ocular tissues (Supplementary Figure 7c and 8b). We also characterized the ocular angle of *Glis1*-KO mice maintained in a 129S6/SvEvTac background. These mice exhibited thinning of the TM layer like that observed in the C57BL/6NCrl background (Supplementary Figure 9). Our data suggest that GLIS1 deficiency leads to progressive TM dysfunction and TM degeneration.

### GLIS1 is highly expressed in TM cells

Since the TM plays a major role in the regulation of AqH drainage and IOP, we decided to focus our study on the analysis of the TM. To examine whether the observed changes in the TM might be intrinsic to the loss of *GLIS1* expression in this tissue, we analyzed *GLIS1* expression in human TM (HTM) tissue and primary HTM cells. The human *GLIS1* gene and its mouse orthologue can generate two transcripts, long and short (referred to as GLIS1L and GLIS1S) that generate a 795 or a 620 amino acids protein, respectively (a 789 and 620 amino acid protein in mice) (https://useast.ensembl.org/Homo_sapiens/Gene/Summary?db=core;g=ENSG00000174332;r=1:53506237-53738106). QPCR analysis demonstrated that GLIS1L was the primary transcript in primary HTM cells, and all human tissues tested (Supplementary Figure 10), whereas GLIS1S was expressed at very low levels. In isolated HTM, characterized by their high myocilin (MYOC) expression, GLIS1 mRNA was expressed at levels comparable to that of kidney, a tissue in which GLIS1 is highly expressed^29^(Figure 3a). QPCR analysis further showed that GLIS1 mRNA was highly expressed in mouse ocular tissue enriched in the TM, moderately in the ciliary body, and at very low levels in the cornea and retina (Figure 3b). *In situ* RNA localization by RNAscope supported the expression of Glis1 transcripts in the TM and ciliary body isolated from 3 months old WT mice (Figure 3c), whereas Glis1 transcripts were not detectable in the iris and cornea. These data indicated that *GLIS1* expression is intrinsic to TM cells and suggests that the TM dysfunction observed in *Glis1*-KO mice is likely causally related to the loss of GLIS1 transcription activation function in these cells. In contrast to TM, the ciliary body, which also expressed Glis1, exhibited a properly organized epithelial layer (Supplementary Figure 7c).

**Figure 3.**
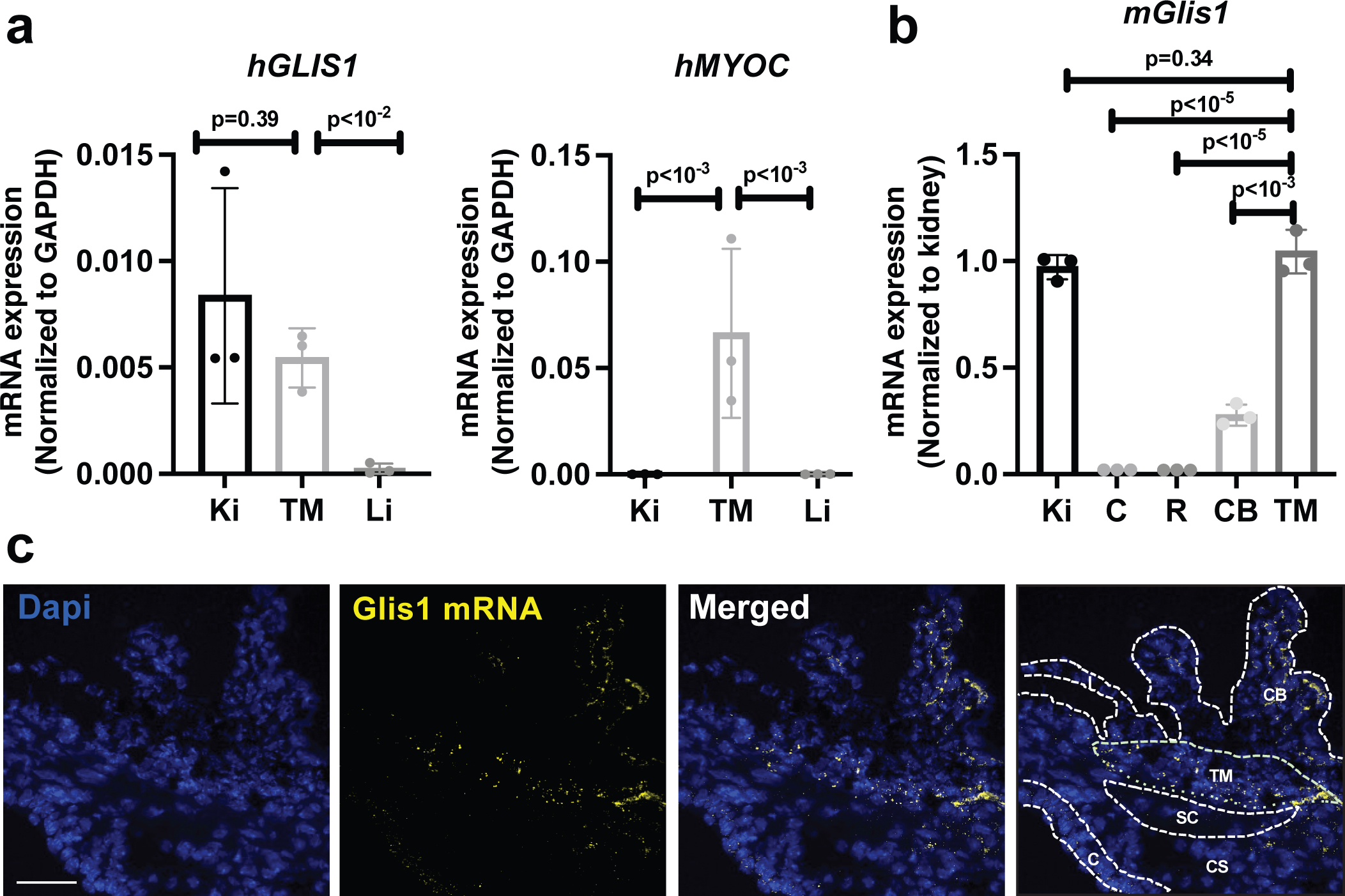
GLIS1 is highly expressed in TM. **a.** QPCR analysis of GLIS1 and MYOC mRNA expression in human TM tissue, kidney (Ki), and liver (Li) (n=3; technical replicates). Statistical analyses were performed with two-tailed Student’s t-test. Data are represented as means ± SD. P-values are indicated above the bars. **b.** Comparison of mouse GLIS1 RNA expression in several ocular tissues with that of kidney, a tissue in which GLIS1 is highly expressed (n=3; distinct samples). Statistical analyses were done with two-tailed Student’s t-test. Data are represented as means ± SD. P-values are indicated above the bars. **c.** RNAscope *in-situ* hybridization with eye sections from 3- month-old WT mouse showed that GLIS1 mRNA (yellow speckles) expression was restricted to TM and CB. Dashed lines outline different cell compartments. Kidney (Ki), Cornea (C), Retina (R), Ciliary Body (CB), Trabecular Meshwork (TM), Corneal StromaI (CS), Iris (I), SC (SC). Scale bar = 50 μm.

### Regulation of gene expression by GLIS1 in primary human TM cells

GLIS1 regulates gene transcription by binding to GLIS binding sites (GLISBS) in the promoter regulatory region of target genes^25, 26^. To investigate alterations in gene expression that might underlie the phenotypic changes observed in TM cells, we performed RNA-Seq and ChIP-Seq analyses. Transcriptome analysis was performed with HTM(shGLIS1) cells, in which GLIS1 expression was knocked down by GLIS1 shRNA lentivirus, and with control cells (HTM(Scr)) infected with scrambled shRNA lentivirus. The volcano plot in Figure 4a shows the distribution of down- and up-regulated genes in HTM(shGLIS1) cells in comparison to HTM(Scr) cells. In addition to down- regulation of GLIS1 mRNA, the expression of several genes associated with TM functions was decreased in HTM(shGLIS1) cells, including *MYOC, CHI3L1, SPARC, CYP1B1,* and *APOD*^39–41^(Figure 4a, b; Supplementary Table 2). In addition, the expression of a variety of genes encoding extracellular matrix components was reduced in HTM(shGLIS1) cells, including a number of collagen genes (e.g., *COL1A2*, *COL6A2*, *COL4A1/2*), fibulins (*FBLN1*, *FBLN5*), microfibril-related genes (*FBN2*, *LOXL1-4*, *LTBP2*), matrix metalloproteinases (*ADAMTS10*, *MMP2*), and genes involved in cell-cell and cell-ECM adhesion (e.g., *ITGA3*) (Figure 4a, b; Supplementary Table 2)^42^. In addition, the expression of a number of genes was up-regulated, including *EFEMP1* and *RTN4*. QPCR-analysis confirmed the decrease in MYOC, BMP2, LOXL4, APOD, LTBP2, and CYP1B1 mRNA expression in HTM(shGLIS1) cells (Figure 4e and Supplementary Figure 11). Several of the differentially expressed genes have previously been reported to be associated with elevated IOP and/or glaucoma, including *MYOC*, *ADAMTS10*, *LTBP2*, *LOXL1*, *TGFBR3*, *CYP1B1*, and *EFEMP1*^5, 28, 43–45^.

**Figure 4.**
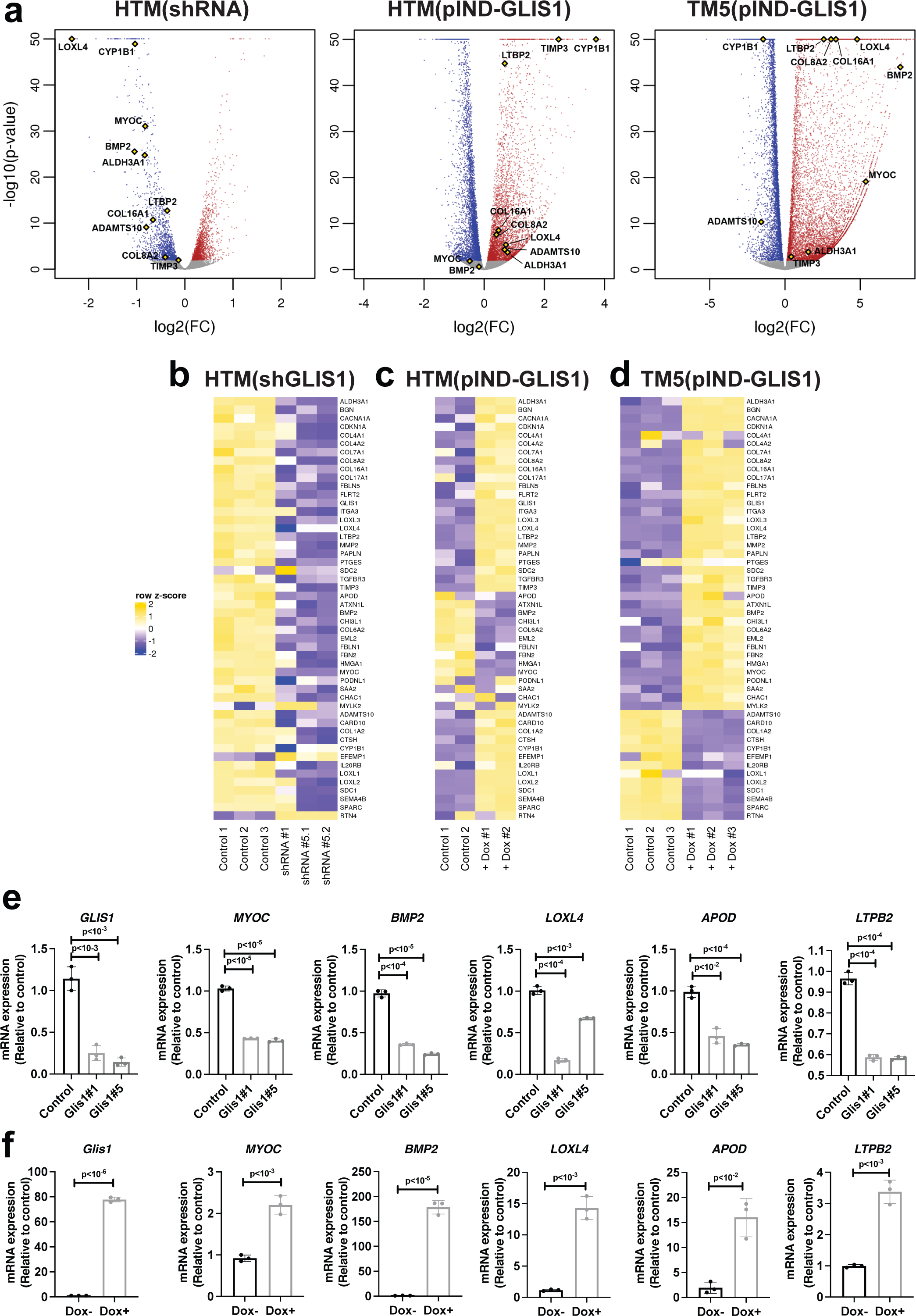
Regulation of TM/glaucoma-related gene expression by GLIS1 in TM cells. **a.** Volcano plots of genes down-regulated (blue) and up-regulated (red) in HTM(shGLIS1) and Dox-treated HTM(pIND-GLIS1) and TM5(pIND-GLIS1) cells (as determined by DESeq2 at FDR 0.01). All other genes are in gray. Several genes associated with IOP, glaucoma or ECM are indicated (yellow diamonds). The X-axis represents gene expression log2-fold change (FC) and the Y-axis represents –log10 (p-value). **b.** Heatmap of the differential expression of several TM-, glaucoma-, ECM-related mRNAs in HTM(shGLIS1) and HTM(Scr) (Control) cells; underlying data are rlog-transformed quantification scores as reported by DESeq2 followed by row-scaling at FDR 0.01. Data shown are for HTM (Scr), shGLIS1#1, and shGLIS1#5 replicates 1 and 2. **c, d.** Heatmap of the differential expression of several TM-, glaucoma-, ECM-related mRNAs in HTM(pIND-GLIS1) cells (**c**) expressing Dox-inducible Flag-GLIS1-HA treated for 18 h with or without Dox (Control) (n=2) and TM5(pIND-GLIS1) cells (n-3)(**d**). **e.** QPCR analysis of several genes down-regulated in HTM(shGLIS1) compared to HTM(Scr) cells (n=3, independent replicates). **f.** QPCR analysis (n=3, independent replicates) of several genes induced by murine GLIS1 in TM5 cells expressing Dox-inducible Flag-GLIS1-HA. Cells were treated for 18 h with or without Dox (+/- Dox). Data in **e** and f are represented as means ± SD. Statistical analyses were performed with two-tailed Student’s t-test. P-values are shown above the bars.

The regulation of many of these genes by GLIS1 was further supported by gene expression analysis in TM cells overexpressing GLIS1. For this analysis we used HTM(pIND-GLIS1) transiently expressing Dox-inducible Flag-GLIS1-HA and TM5(pIND- GLIS1), stably expressing a Dox-inducible Flag-GLIS1-HA. Dox treatment greatly induced GLIS1 mRNA expression in TM5(pIND-GLIS1) cells and accumulation of Flag- GLIS1-HA protein in the nucleus (Figure 4f; Supplementary Figure 12). Transcriptome analysis showed that induction of GLIS1 expression in Dox-treated HTM(pIND-GLIS1) and TM5(pIND-GLIS1) cells enhanced the expression of many, but not all, of the same genes that were down-regulated by shGLIS1 RNAs in HTM cells (Figure 4a, c, d; Supplementary Table 2). QRT-PCR analysis showed that the decreased expression of *MYOC, BMP2, LOXL4, APOD*, and *LTBP2* in HTM(shGLIS1) correlated with increased expression in Dox-treated TM5(pIND-GLIS1) cells (Figure 4e and f). Similarly, the induction of CYP1B1 mRNA in HTM(pIND-GLIS1) cells correlated with decreased expression in HTM(shGLIS1) (Supplementary Figure 11). As indicated above, decreased expression in HTM(shGLIS1) did not always perfectly correlate with increased expression in HTM(pIND-GLIS1) and/or TM5(pIND-GLIS1) cells (Supplementary Table 2). Such differences might, among other things, be due to variations in the epigenome and the transcription regulatory machinery between primary and immortalized TM cells or different efficiencies of the shGLIS1 used. It might further relate to differences in the expression levels of endogenous GLIS1 or that of GLIS1 target genes in HTM versus TM5 cells or variations in the binding affinity of GLIS1 to GLIS binding sites of target genes.

To determine which of the differentially expressed genes were direct transcriptional targets (cistrome) of GLIS1, we performed ChIP-Seq analysis. Since no suitable GLIS1 antibody was available for ChIP-Seq and it was not feasible to establish primary HTM cells stably expressing GLIS1, hence we utilized the TM5(pIND-GLIS1) cells. ChIP-Seq analysis to identify direct transcriptional targets of GLIS1 showed an enrichment for GLIS1 binding (Figure 5a). ChIP-Seq analysis identified a total of 46,947 distinct GLIS1 binding peaks in Dox-treated TM5(pIND-GLIS1) cells. About 10% of GLIS1 binding peaks were within proximal promoter regions 1 kb upstream of transcription start sites (TSSs), whereas 16% were further upstream (Figure 5b). GLIS1 binding was most highly enriched at introns within the gene body as we reported for GLIS3^32, 46^. Homer *de novo* and known motif analyses identified a G/C-rich GLISBS-like consensus sequences as the top motifs (Figure 5c). This sequence was very similar to the consensus GLISBS reported previously^32, 46, 47^, indicating that our ChIP-Seq was successful in detecting specific GLIS1 binding sequences. Our GLIS1 ChIP-Seq analysis identified a number of additional motifs, including motifs for bZIP transcription factors (e.g., ATF3, FRA1, BATF, and JUNB, members of the AP-1 complex), forkhead box (FOX) proteins ^48^, and TEA domain transcription factors (TEAD) that play a key role in the Hippo pathway ^49^. These data suggested co-localization of the GLIS1 binding consensus with motifs for other transcription factors that have been previously implicated in the regulation of TM and glaucoma ^45, 50, 51^. These findings are consistent with the hypothesis that GLIS1 regulates TM gene transcription in coordination with other transcription factors.

**Figure 5.**
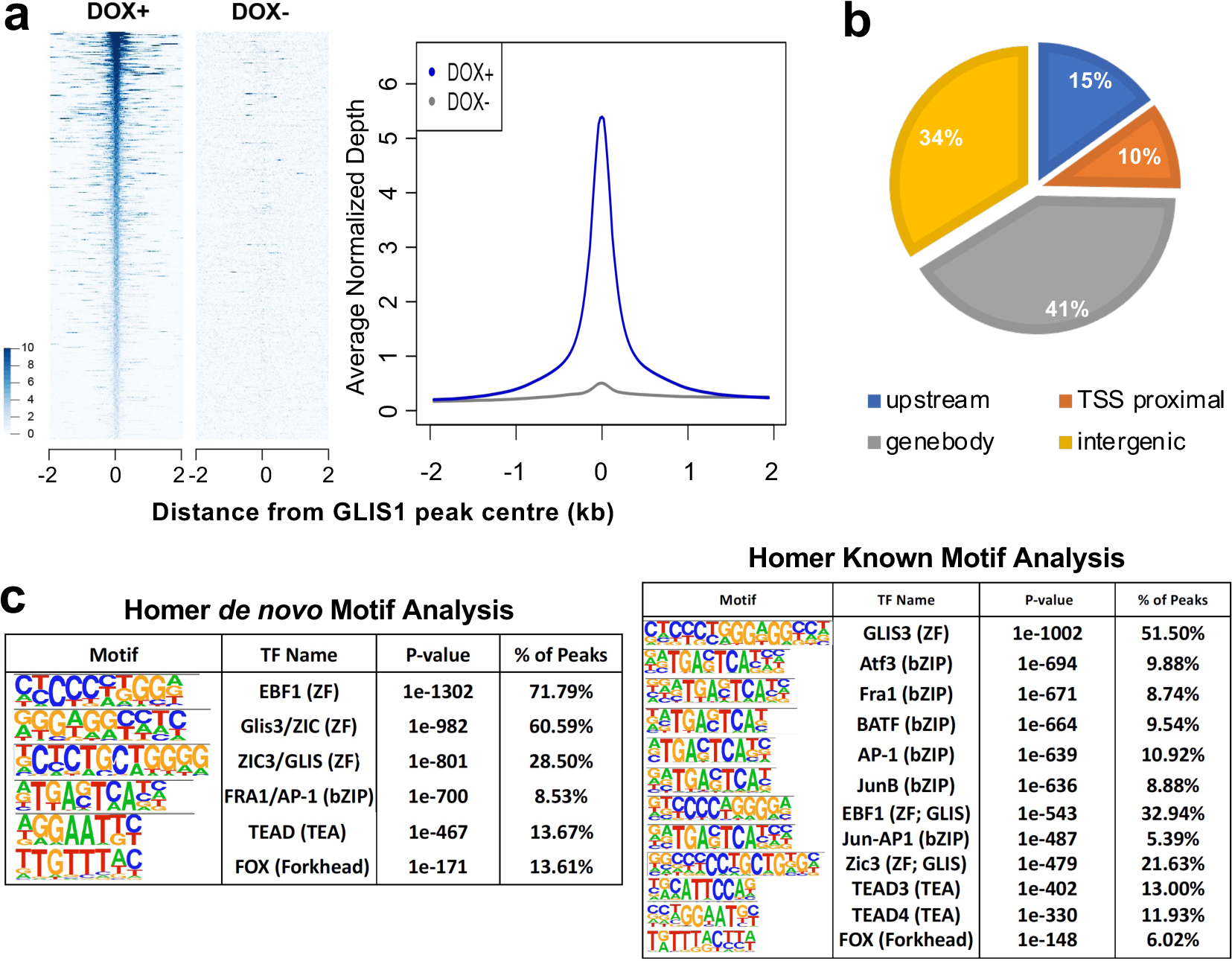
GLIS1 regulates the transcription of a subset of TM/glaucoma-related genes in HTM cells through its interaction with GLISBS. **a.** Heatmap and ChIP-Seq read density plot showing GLIS1 occupancy in TM5(pIND-GLIS1) cells treated for 18 h with Dox (+Dox) compared to untreated cells (-Dox). Each line in the heatmap represents an individual GLIS1 binding site. **b.** Pie chart showing the location of GLIS3-binding peaks within specific regions of the genome. TSS proximal: −1 kb to transcription start site (TSS); Upstream: −1 to −10 kb. **c.** Homer *de novo* and known motif analysis identified GLIS-like binding sites (GLISBS) as the top consensus sequence motif. Binding sites for transcription factors of the AP-1, FOX, and TEAD families were identified alongside GLIS consensus motifs.

Analysis of the combined RNA-Seq and ChIP-Seq data showed that the transcription of several of the differentially expressed genes with roles in TM, IOP, and glaucoma, were directly regulated by GLIS1 and included *MYOC*, *LTBP2*, *CHI3L1*, *HMGA1*, *CYP1B1*, and *LOXL1-4* (Supplementary Table 2). Genome browser tracks showing the location of GLIS1 peaks associated with several glaucoma-related genes, including *MYOC*, *CYP1B1*, and *ADAMTS10*, are shown in Figure 6. Interestingly, the GLIS1 binding peaks in the proximal promoter regions of *MYOC* and *CYP1B1* are located in a region near with a functional AP-1 responsive element (Figs. 5c and 6)^52–54^. In addition, the proximal promoter of CYP1B1 contains a G/C-rich SP1 binding sequence, which because of its similarity to the consensus GLISBS might function as a GLIS1 binding site. This suggests that these promoter regions may function as a regulatory hub for several transcription factors. Moreover, this supports our hypothesis that the transcription of *MYOC*, *CYP1B1*, and other TM genes by GLIS1 is regulated in coordination with other transcription factors, including members of the AP-1 family.

**Figure 6.**
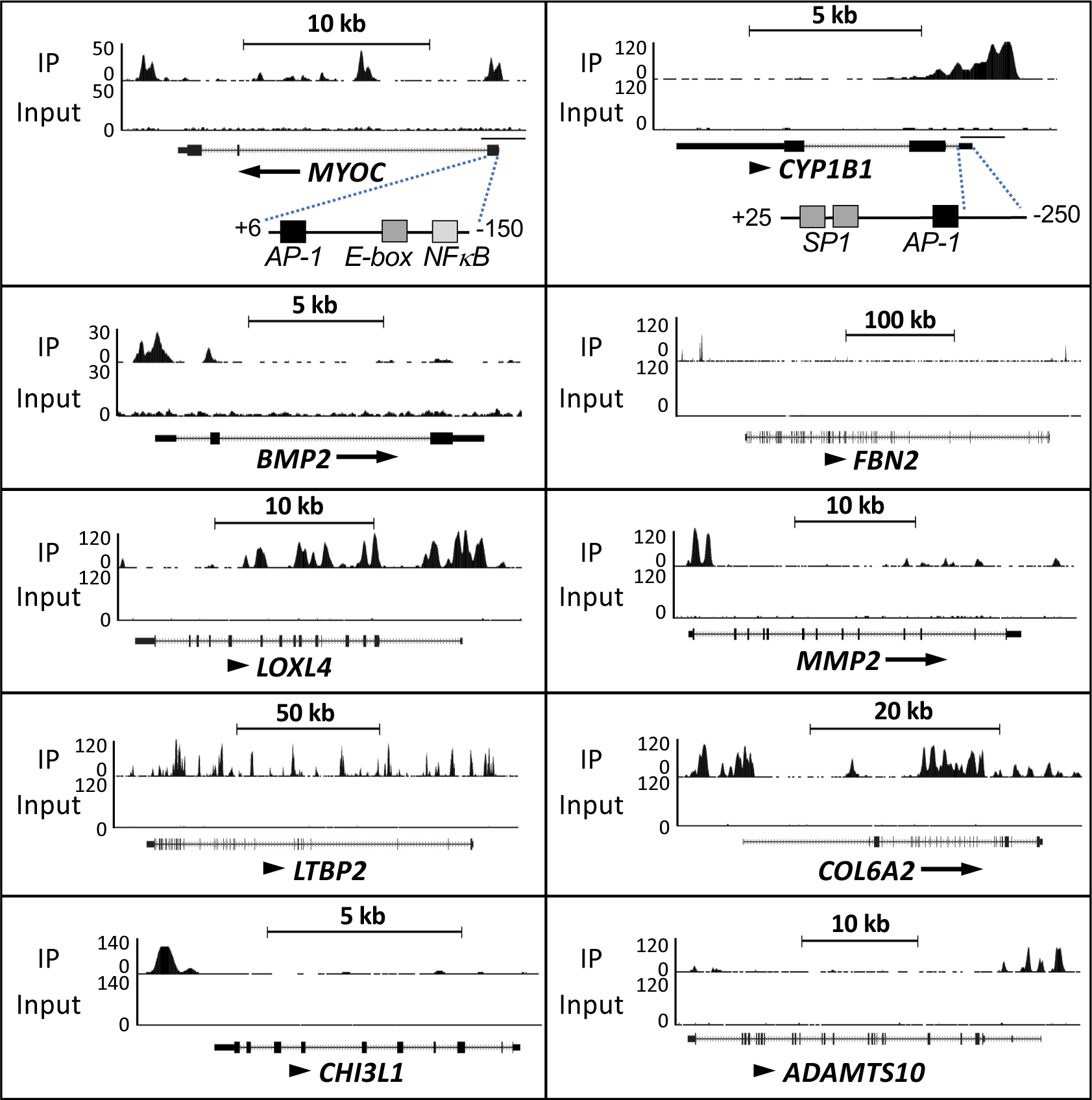
Genome browser tracks of the *MYOC*, *CHI3L1, BMP2*, *FBN2, LOXL4, MMP2, LTBP2*, *COL6A2,* CYP1B1 and *ADAMTS10* genes (https://genome.ucsc.edu/) showing GLIS1 occupancy (ChIP-Seq) in TM5(pIND-GLIS1) cells expressing Flag- GLIS1-HA. The AP-1, E-box, and NFκB binding sites in *MYOC* and the AP-1 and G/C-rich SP1 binding sites in the *CYP1B1* proximal promoter region are indicated.

KEGG pathways analysis of GLIS1 target genes down-regulated in HTM(shGLIS1), activated in HTM(pIND-GLIS1) or TM5(pIND-GLIS1) cells identified pathways associated with extracellular matrix (ECM), proteoglycans, and cellular adhesion among the top pathways in all three data sets (Supplementary Figure 13). This is consistent with recent bioinformatics analyses of TM gene expression data that identified cell matrix and cell-cell interaction related pathways among the top pathways involved in the pathogenesis of POAG ^55^.

### Association of *GLIS1* rs941125 with glaucoma

Our study of Glis1-KO mice identified a critical regulatory function for GLIS1 in the maintenance and function of the TM, a tissue that plays a major role in controlling AqH outflow and the development of glaucoma^1, 15, 56, 57^. These findings raised the question whether *GLIS1* might be involved in the pathogenesis of human glaucoma as well. To assess this, we examined the association of SNPs in the *GLIS1* region and the risk of glaucoma, combining information from the GERA and UKB cohorts^9^. rs941125, which localizes to intron 1 of *GLIS1*, was the most strongly associated SNP in the region, reaching a Bonferroni corrected level of significance (Odds Ratio = 0.94, *P*=4.73X10^-6^) (Figure 7 and Supplementary Table 3). The association of rs941125 with POAG has recently been confirmed and replicated in additional cohorts at a genome-wide level of significance (*P*=2.01X10^-11^, meta-analysis)^58^. Furthermore, rs941125 is significantly associated with variation in *GLIS1* gene expression in several tissues in GTEx (https://gtexportal.org/home/snp/rs941125). Together, these findings support a role for *GLIS1* in glaucoma pathogenesis in humans.

**Figure 7.**
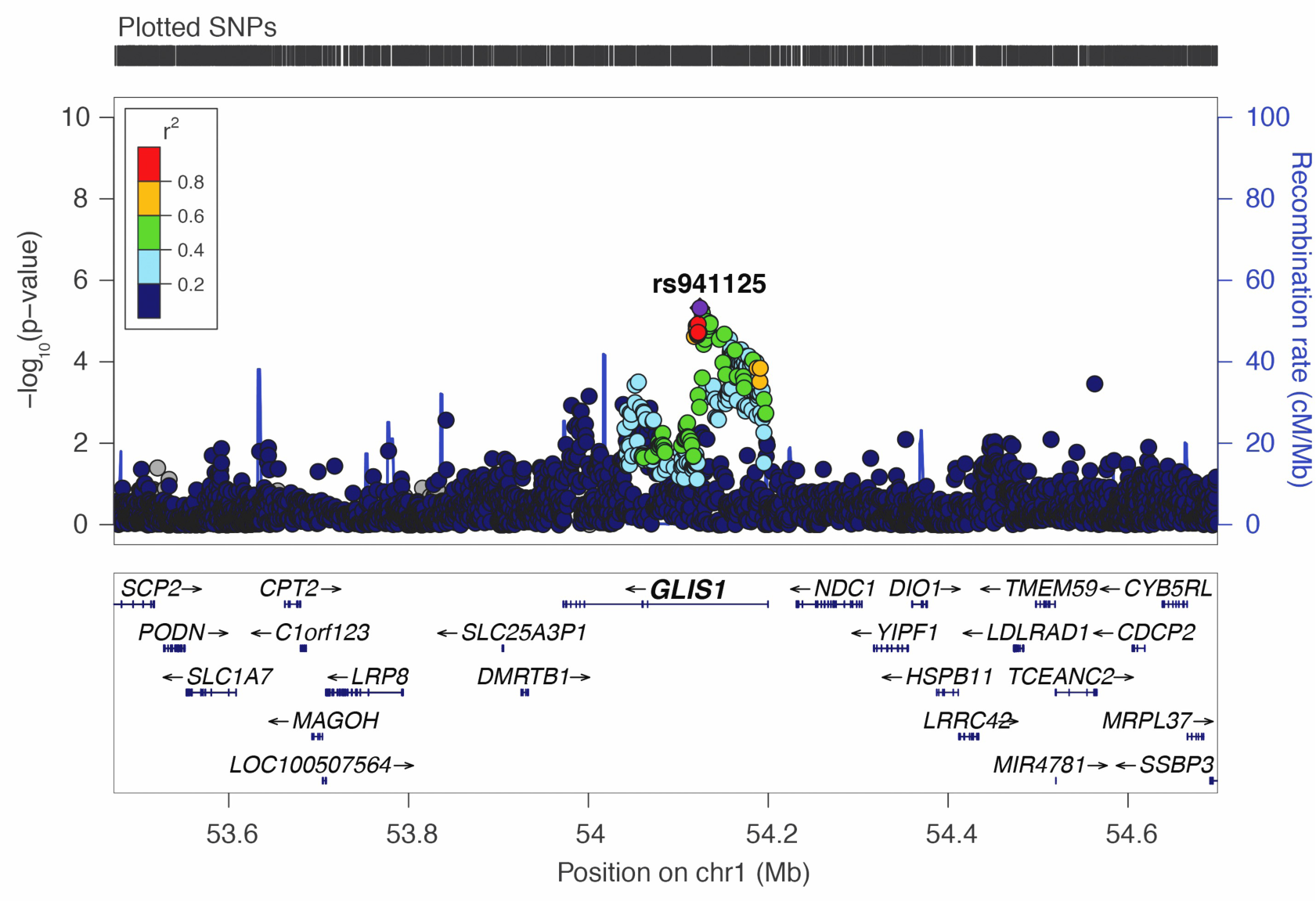
Regional plot at the *GLIS1* genomic region showing association with glaucoma in the combined (GERA + UK Biobank) multiethnic meta-analysis. Top SNP rs941125 is significantly associated with glaucoma after Bonferroni correction (p=4.73X10^-6^).

## Discussion

In this study, we identify a critical role for the transcription factor GLIS1 in the maintenance of TM/ocular drainage tissue, AqH dynamics, and IOP. The TM is an essential component of the ocular drainage structure and TM abnormalities play a major role in the development of elevated IOP and glaucoma^1, 5, 15, 57, 59, 60^. Utilizing *Glis1*-KO mice, we demonstrate that the loss of GLIS1 function causes a progressive degeneration of the TM leading to a disruption of ocular drainage structures. As a consequence of these changes, the AqH drainage is significantly compromised in *Glis1*-KO eyes causing elevated IOP. In addition, we identified several transcriptional targets of GLIS1 in TM cells that previously have been implicated in TM-related functions, IOP homeostasis, and ocular hypertension/glaucoma, including *MYOC*, *ADAMTS10*, *LTBP2*, *LOXL1*, *TGFBR3*, *CYP1B1*, and *EFEMP1*^5, 28, 43, 44^. The reduced expression of a set of TM genes together is likely responsible for the TM dysfunction and elevated IOP in *Glis1*-KO mice. Importantly, we have detected significant associations between common genetic variants in the *GLIS1* region and POAG in humans, thereby supporting the role of *GLIS1* as a glaucoma risk gene. These variants may impact TM functions and compromise AqH drainage by altering GLIS1 expression and/or function and leading to elevated IOP and glaucoma.

The expression of *Glis1* in the TM and the degeneration of the TM in *Glis1*-KO mice suggested that GLIS1 plays a critical regulatory role in maintaining TM cell function and survival. It is possible that GLIS1 deficiency leads to the disruption of a cell-intrinsic biological process important in maintaining normal structure and TM function. Excessive loss of TM cells has been proposed to be a critical pathophysiological feature resulting in defective AqH drainage and high IOP^14^. Although pronounced loss of TM cells in POAG patients was observed decades ago^13, 61^, much remains yet to be understood with regards to cellular processes regulating the maintenance of the TM and survival of TM cells and its relevance to IOP and the different forms of glaucoma^14, 15, 56^. Various changes in the TM, including alterations in ECM and mitochondria, increased oxidative stress and apoptosis, have been implicated in outflow resistance, elevated IOP, and increased glaucoma risk^15, 41, 42, 55, 62–65^. Specifically, the ECM has been identified as a major player in maintaining the structural integrity and functionality of the TM^55, 59, 61, 64–66^. Consistent with this, our gene profiling analyses identified ECM-related processes, cytoskeleton, and cellular adhesion as key pathways that are impacted as a consequence of *GLIS1* knockdown or ectopic expression of *GLIS1* (Figure 4e and f; Supplementary Figure 13). Moreover, several ECM-related genes, which levels changed in TM cells upon *GLIS1* knockdown or overexpression, have previously been implicated in elevated IOP and glaucoma, including members of the collagen I, IV, and VI families, *LTBP2*, a regulator of TGFβ signaling and ECM deposition^28, 67^. Furthermore, the expression of a number of microfibril-associated genes are also impacted, including *FBN2*, encoding a microfibril-associated glycoprotein that contributes to elastin assembly in the ECM of the TM^60, 68^. It further includes *LOXL1-4*, encoding lysyl oxidases that mediate the cross- linking of several extracellular matrix proteins, such as collagens and elastin^21, 69^, and *ADAMTS10*, encoding a metalloproteinase involved in ECM assembly^44, 70^. Alterations in the expression of these genes are likely to impact ECM assembly as well as its biomechanical properties, and physiological processes, such as differentiation, survival, and tissue organization that are regulated by the ECM-dependent signaling pathways^71^. Thus, altered cell-ECM interaction or abnormal ECM organization in *Glis1*-KO mice might adversely affect the maintenance and biomechanical properties of the TM, thereby leading to progressive degeneration of the TM and disruption of the AqH drainage and subsequently to the development of elevated IOP and glaucoma^14, 42, 62, 63^. Given a role for ECM-dependent pathways, it is possible that change in biomechanical properties of the TM (TM stiffness) may contribute to IOP changes, especially during an early time window prior to any obvious structural changes not captured by histological assessment.

In addition to these ECM and adhesion-related genes, transcriptome analysis showed that GLIS1 impacts the expression of several other TM-, IOP- and glaucoma-related genes, including *MYOC* and *CYP1B1* (Figure 4a, e, f; Supplementary Figure 12). It is interesting to note that the phenotypic changes in the TM observed in *Glis1*-KO mice exhibit some resemblance with those seen in *Cyp1b1*-deficient mice, including the collapse and degeneration of the TM^37, 61^. Thus, the reduced *CYP1B1* expression might contribute to the structural changes in the TM and elevated IOP observed in *Glis1*-KO mice. In the case of MYOC, mutations in *MYOC* are thought to act by a gain of function mechanism resulting in the misfolding and accumulation of mutant MYOC leading to ER stress and apoptosis of TM cells^72, 73^. Moreover, studies with *Myoc* knockout mice suggest that loss of *Myoc* function by itself does not cause ocular drainage tissue abnormalities or IOP elevation^74^. These studies indicate that reduced expression of *Myoc* observed in *Glis1*-KO mice is unlikely by itself inducing TM abnormalities and elevated IOP. We hypothesize that reduced expression of a set of genes rather than one particular single gene is causing the TM abnormalities and high IOP in GLIS1 deficiency, as we reported for the development of congenital hypothyroidism and neonatal diabetes in *Glis3*-KO mice^32, 47^. Thus, the altered expression of a set of TM genes, including CYP1B1, ADAMTS10, and LTBP2, may underlie TM dysfunction and elevated IOP. Therefore, the effect of Glis1 deficiency on mouse TM is likely a cumulative effect of disruption of multiple pathways.

To establish which of the differentially expressed genes were directly regulated by GLIS1, we performed ChIP-Seq analysis. This analysis revealed that GLIS1 binding was associated with *MYOC, ADAMTS10, CYP1B1*, *MMP2,* and many other genes, suggesting that these genes are direct transcriptional targets of GLIS1 (Supplementary Table 2). Interestingly, many of the direct targets of GLIS1 have a role in the ECM or TM function and have been implicated in the pathogenesis of elevated IOP and glaucoma^6, 28, 44, 56^. Homer motif analysis suggested co-localization of GLIS1 binding peaks with motifs of other transcription factors, including binding sites for members of the AP-1, TEAD, and forkhead box (FOX) families (Figure 5). Interestingly, the proximal promoters of *MYOC* and *CYP1B1* have been reported to contain functional binding sites for AP1-related transcription factors near the location of the GLIS1 binding peaks (Figure 6)^52–54, 75, 76^. The forkhead box member, FOXC1, has been implicated in the regulation of TM functions and glaucoma^77, 78^, while the Hippo pathway through activation of TEAD transcription factors regulates ECM in TM cells and appears to have a role in glaucoma^50, 51, 79^. Together, these observations support a model in which a selective set of GLIS1 target genes are co-regulated with other transcription factors through their interaction within the same regulatory regions in TM-specific genes.

Our genetic studies provide further validation for a role of GLIS1 in glaucoma pathogenesis and extend our findings in mice to humans. The observed ocular drainage defects exhibited by *Glis1*-KO mice is more severe than those observed in POAG and likely due to a complete deficiency of GLIS1 and the suppression of many target genes, as compared to TM specific changes originating from potential gene dosage effects of POAG-associated variants. The lead SNP rs941125 is not only associated with glaucoma in multiple independent cohorts^58^, it is also detected as an eQTL associated with change in *GLIS1* expression (https://gtexportal.org/home/snp/rs941125). We note, however, that GTEx does not contain information on eye tissues and that the eQTL analysis for rs941125 is based on expression data from brain tissue. Future studies need to determine whether rs941125 or other SNPs in linkage disequilibrium with rs941125 are located within gene regulatory elements (such as enhancers) that affect *GLIS1* expression and how alteration of these regulatory elements predispose the eyes to TM dysfunction leading to IOP elevation. Since POAG is multifactorial, it is likely that variants in other genes together with *GLIS1* SNPs may cooperate to induce high IOP in patients. The absence of such other modifier alleles in *Glis1* heterozygous mice might explain why the presence of a single knockout allele of *Glis1* is not sufficient to induce ocular drainage defects leading to elevated IOP.

In this study, we identify a critical role for GLIS1 in the maintenance and regulation of TM function, AqH dynamics, and IOP. GLIS1 together with other transcription factors might be part of a regulatory network required for proper maintenance and functioning of ocular drainage tissue and IOP homeostasis ^45^. Thus, *Glis1-*KO mice provide us with a valuable model to uncover cellular and molecular mechanisms that underlie the regulation of TM maintenance and ocular drainage tissue homeostasis and potentially lead to new insights into the pathogenesis of glaucoma. In addition, our data further suggest that altered expression of *GLIS1* in individuals carrying the risk allele may confer increased susceptibility towards developing POAG possibly by impacting the TM and thereby contributing to elevated IOP. Finally, as has been shown for the hedgehog/GLI signaling pathway, regulation of GLIS proteins^25^ by primary cilium-associated G protein-coupled receptors might be useful for the development of new therapeutic strategies in the management of various pathologies, including glaucoma.

## METHODS

### Glis1-deficient mice

Glis1-deficient mice (*Glis1*-KO) were described previously^30^. Mice were bred into the C57BL/6NCrl Charles River, Wilmington, MA) and 129S6/SvEvTac (Taconic, Rensselaer, NY) backgrounds for at least 7 generations. We ensured that mice maintained in C57BL/6NCrl neither carried homozygous RD8 mutation nor exhibited retinal degeneration (based on ocular histological assessment). *Glis1*-KO mice on both backgrounds developed enlarged eyes and appeared to exhibit a similar phenotype. Most experiments were carried out with *Glis1*-KO C57BL/6NCrl mice. Mice were supplied *ad libitum* with autoclaved NIH-31 rodent diet (Harlan Laboratories, Madison, WI) and provided with distilled drinking water and were group-housed in individually ventilated cages (Techniplast, Exton, PA). Experiments took place in an AAALAC accredited facility maintained at 70-73°F, relative humidity 40-60%, and 12h:12h light- dark cycle. All mice were negative for rodent murine pathogens. Littermate wild-type (WT) mice were used as controls. All animal protocols followed the guidelines outlined by the NIH Guide for the Care and Use of Laboratory Animals and were approved by the Institutional Animal Care and Use Committee at the NIEHS. Routine genotyping was carried out with the following primers: Glis1-F, 5’-AGCTAGTGGCTTTCGCCAACA; Glis1-R, 5’- GAACAAGATAGAATCATGG-TATATCC and Neo-pro, 5’- ACGCGTCACCTTAATATGCG.

### T3, T4, and thyroid stimulating hormone (TSH) assays

Blood levels of T3 and T4 were measured by radioimmunoassay (MP Biomedicals, Orangeburg, NY) as described previously^32^. Serum TSH was analyzed with a mouse pituitary magnetic bead panel kit (EMD Millipore Corp., Billerica, MA.

### RNAscope *in situ* hybridization

*In situ* hybridization was carried out on formalin-fixed paraffin embedded tissue sections using the RNAScope system (Advanced Cell Diagnostic, Hayward, CA) per manufacturer’s instructions and as previously described^80^.

### Intraocular Pressure Measurement

IOP of both eyes in age- and gender-matched WT and *Glis1*-KO littermates was measured using the Icare TonoLab rebound tonometer (Icare, Helsinki, Finland). Immediately prior to measurement animals were briefly sedated using isoflurane. IOP was measured at least 4 times/eye within a 30 min time frame with each individual IOP recorded representing the average of six measurements, giving a total of 24 rebounds from the same eye.

### Ocular angle assessment

Hematoxylin and eosin-stained ocular sections cut from plastic embedded eyes were assessed to examine the morphology of the ocular angle structures. Briefly, mice were euthanized and eyes enucleated and immediately immersed in cold fixative (1% PFA, 2% glutaraldehyde, and 0.1 M cacodylate buffer) for at least 48 h, after which they were transferred to cold 0.1 M cacodylate buffer solution and stored at 4°C. We have found that using this fixative, greatly improves capturing the SC in its intact/non-collapsed conformation. For example, at around 3 months of age 100% of WT eyes showed intact SC. Samples were embedded in glycol methacrylate, and serial sagittal sections (2 μm) passing through the optic nerve were cut and stained with hematoxylin and eosin (H&E). Ten similarly spaced sections corresponding to the peripheral, mid- peripheral and central regions (passing through the optic nerve) of the eye were evaluated^81^. Both angles of a section were considered for evaluation. The prominent angle relevant structures, including TM, SC, Iris, ciliary processes were histologically examined. A representative image showing angle relevant tissues are indicated in Supplementary Figure 14. Eyes assessed for each of genotype included both sexes.

The sections corresponding to the central region of the eye were used for measuring the TM area, which in our experience provides the most reliable assessment. We imaged on average five consecutive serial sections per eye from both wild type and *Glis1-* KO mice. Cross-section images were taken using a 20X objective on the Zeiss Axiophot microscope with a 12 Mp Insight camera. TM images were captured by SPOT5.6 imaging software, assigned an accurate 20X calibration (to account for the magnification of the acquired image), and the TM area (marked in Supplementary Figs. 5, 6) was measured using the “region” tool under the EDIT menu. The mean TM area was calculated from a minimum of 5 central ocular sections/eye.

### Assessment of the SC

Briefly, the anterior segment is excised and the iris removed. The anterior segment cup is relaxed by making four centripetal cuts. These cuts generated four fan shaped quadrants attached at the center. The SC runs along the rim (limbus) of each fan shaped quadrant. Whole mounts of the anterior segments from control and *Glis1*-KO mice were stained with an endomucin antibody^82^. Briefly, the anterior segment is excised and the iris removed. The anterior segment cup is relaxed by making four centripetal cuts. These cuts generated four fan shaped quadrants attached at the center. The Schlemm’s canal runs along the rim (limbus) of each fan shaped quadrant. The anterior segment was stained with endomucin (5µg/ml; Thermo Fisher Scientific), whole mounted, and the entire limbus encompassing each of the four quadrants imaged using a Leica LSM SP8 confocal system and DM6000 vertical microscope. The limbal region was imaged with a 20x/0.75 IMM CORR CS2–multi-immersion objective using glycerol immersion media. Overlapping regions (10%) were collected as Z-stacks at a resolution of 541x541x1 µm. The overlapping Z-stack of a quadrant was stitched using XuvStitch freeware^83^. The confocal Z-stack of the stitched quadrants of the limbus were rendered in three dimensions using the Surpass mode of Imaris 9.2 (Bitplane). Imaris Surface tool was used to render a surface on to the endomucin labeled Schlemm’s canal. The volume was obtained from the “Statistic” tab under the surface algorithms in the software. The data was downloaded as a .csv file. The volumes of Schlemm’s canal in quadrats were graphed using PlotsOfData-A web app for visualizing data together with their summaries^84^.

### AqH dynamics by Gadolinium magnetic resonance imaging (Gd MRI)

AqH dynamics was analyzed by 3D Gd MRI as described^34, 35^. The MR imaging data are accessible via the following url: https://civmvoxport.vm.duke.edu/voxbase/studyhome.php?studyid=733. All MRI measurements were performed utilizing a 7.1-Tesla/22-cm horizontal bore Magnex magnet with an Agilent Direct Drive Console (Santa Clara, CA, USA), providing up to 770 mT/m gradient strength, and a 35 mm transmit–receive birdcage coil. Mice (n=5) were anesthetized with isoflurane (2% for induction and 1.5% for maintenance) and kept warm with warm circulating air during the MRI experiment. Respiration rate was monitored using a small pneumatic pillow (SA Instruments, Inc., Stony Brook, NY, USA). Gadolinium-DTPA (Magnevist, Schering, Germany) was intraperitoneally (i.p.) injected at a dose of 0.3 mmol/kg after one T1-weighted MR image was acquired at baseline. The MR contrast was thus administered at a dose calculated to normalize to the body mass for each animal.

Some mice were treated with the IOP lowering compound Ripasudil (0.4% normal isotonic saline; Sigma), which was administered in 5 μl drops to the right eye, while normal isotonic saline was added to the left eye. T1-weighted MR images were acquired using a gradient echo sequence repeated over 2 h, sampling at 10 minutes intervals, for a total of 12 scans. The imaging parameters were repetition time/echo time = 200/1.92 ms, field of view = 14.4 × 14.4 mm^2^, matrix 192x192, flip angle 20 degrees, BW 62.5 kHz, and the in-plane resolution was 75 × 75 μm^2^. 11 coronal slices, 0.38 mm thick were acquired. 16 averages were used for each scan, resulting in total scan time for each temporal sampling interval of 10 min 12 seconds. An additional scan was acquired at the end of the dynamic contrast enhanced study, using identical parameters but increasing the number of averages to 64, to help delineate the anatomy. Select specimens were imaged *ex vivo* using a multi gradient echo sequence. Eye specimens were prepared after trans cardiac perfusion fixed with a mixture of saline and ProHance (10%), followed by a mixture of formalin and ProHance (10%). The imaging parameters were repetition time/echo time = 50/3 ms, field of view = 25.6 × 12.8 x 12.8 mm^2^, matrix 512x256x256, flip angle 60 degrees, BW 62.5 kHz, and the 3D isotropic resolution was 50 μm x 50 μm × 50 μm.

Regions of interest (ROIs) were manually drawn in the T1-weighted imaging slice bisecting the center of the globes, using ImageJ v1.47 (Wayne Rasband, National Institutes of Health, Bethesda, MD, USA). We measured the enhancement of the signal intensity (brightness of voxels) due to Gd accumulation in the anterior chamber, and our measurements were averaged over the entire ROIs. To compensate for inter animal variability and to reflect the relative enhancement in each animal, these measurements were normalized to the 10 min baseline. Several studies have demonstrated a linear relationship between the concentration of Gd and the spin lattice relaxation rate (R1=1/T1) over limited ranges of concentration^85–87^. MØrkenborg et. al.^88^ have performed experiments at 7T, the field used in these studies and demonstrated a linear correlation (r^2^ >0.92) between signal intensity in a gradient echo for concentrations ranging between 0-3.0 mmol/lGd-DTPA. Thus, while concentration was not measured directly by measurement of R1, it can be inferred from the signal intensity. In addition, we have a curve normalization to the same time (10 min) acquired in each animal to adjust for different gains. The averaged time course from each ROI measurement before and after Gd injection was fitted into a sixth-degree polynomial using MatLab2016a (The MathWorks, Inc., Natick, MA, USA). The peak percentage (%) Gd signal enhancement, time to peak, initial rate of Gd signal increase within the first 10 minutes after Gd injection, and the area under the curve were extracted from the time courses and compared between both eyes of the same groups using two-tailed paired *t*-tests, and across groups using one-way ANOVA and post hoc Tukey’s tests. Data are presented as mean ± standard deviation unless otherwise specified. Results were considered significant when *P* < 0.05.

### Cell lines

Human kidney HEK-293T cells were obtained from ATCC and grown in DMEM plus 10% FBS. Primary HTM cells were provided by Dr. T. Borras. Cells were grown in Modified IMEM (Cat. No. A1048901) supplemented with 10% fetal bovine serum (FBS) (Gibco, Grand Island, NY) and 50 μg/ml gentamycin (ThermoFisher) and used at <3 passages. Immortalized HTM-like HTM5 cells (TM5)^90^ were cultured in DMEM/F12 supplemented with 10% FBS, L-glutamine, penicillin (100 units/ml), streptomycin (0.1 mg/ml), and amphotericin B (4 mg/ml). Cells were tested negative for mycoplasma at NIEHS or UCSF.

### Reporter assay

HEK-293T cells were transfected in Opti-MEM with pTAL-Luc-(GLISBS)6 reporter, in which the luciferase reporter is under the control of six copies of GLISBS, pCMV-β-Gal, and a pCMV10 expression plasmid containing wild type Flag-GLIS1 or a Flag-GLIS1 mutant using Lipofectamine 2000. 24 hours later cells were harvested into 125 μl reporter lysis buffer and luciferase activity and β-galactosidase levels measured using a luciferase assay kit (Promega, Madison, WI) and a luminometric β-galactosidase detection kit (Takara Bio, Palo Alto, CA) following the manufacturer’s protocol. Experiments were carried out in independent triplicates^89^.

### GLIS1 shRNA knockdown

GLIS1 knockdown in HTM cells was performed by infecting cells with GLIS1 shRNA lentivirus (Dharmacon; GLIS1#1-TRCN0000107705 and GLIS1#5-TRCN0000107709 or scrambled shRNA (control)(MOI 1:10). These cells are referred to as HTM(shGLIS1) and HTM(Scr). 48 h later cells were collected and RNA isolated with a Purelink RNA mini kit (ThermoFisher Sci., Rockford, IL) for RNA-Seq analysis as described^32, 46^.

### Quantitative-PCR

Kidney, ciliary body, TM, cornea, and retina were dissected from eyes of 3-month-old WT mice (n=3), RNA was isolated using a RNeasy mini kit (Qiagen) and reverse-transcribed using the High-Capacity cDNA Archive Kit (Applied Biosystems, Foster City, CA). Glis1 expression was then measured by digital droplet PCR (ddPCR) using the QX200™ Droplet Digital™ PCR System (BioRad) and normalized to Hsp90a01 expression. Primers for Glis1 (Mouse; PrimePCR ddPCR Expression Probe Assay (BioRad); FAM; dMmuCPE5121630) and Hsp90ab1 (Mouse; PrimePCR ddPCR Expression Probe Assay (BioRad); HEX; dMmuCPE5097465). Accepted ddPCR reads had a minimum of 12,000 events and cDNA concentrations were adjusted to be within 10 to 10,000 positive events. To analyze gene expression from cultured cells, QPCR analysis was performed using SYBR Green I (Applied Biosystems, Foster City, CA). RNA from cultured HTM and TM5 cells was isolated with a Purelink RNA mini Kit (ThermoFisher) and QPCR analysis performed as described previously^32, 46^. RNAs from human tissues were from a Clontech Human Total RNA Master Panel II (#636643). Primer sequences are listed in Supplementary Table 1.

### ChIP-Seq analysis with Flag-GLIS1-HA HTM and TM5 cells

Since no suitable GLIS1 antibody is available for ChIP-Seq analysis, we used primary HTM cells transiently expressing doxycycline (Dox)-inducible Flag-GLIS1-HA and a TM-like cell line, TM5, that stably expressed Dox-inducible Flag-GLIS1-HA. First, a pIND20(Flag-GLIS1-HA) plasmid was generated by inserting Flag-GLIS1-HA into the (Dox)-inducible lentiviral expression vector pIND20^91^ and are referred to as HTM(pIND- GLIS1) and TM5(pIND-GLIS1), respectively. Lentivirus was generated by transient transfection of pIND20(Flag-GLIS3-HA) in HEK293T cells together with psPAX2 and pMD2.G plasmids. TM5 cells were infected with the pIND20(Flag-GLIS1-HA) lentivirus for 48 h and then selected in medium supplemented with 750 μg/ml G418 (Invitrogen, Carlsbad, CA). Flag-GLIS1-HA expression was induced by the addition of 300 μg/ml Dox (Sigma–Aldrich, St. Louis, MO). The expression of Flag-GLIS1-HA protein was examined by immunofluorescence. The relative fluorescent signal in nuclei was determined using ImageJ software (Fuji) as described^92^. To identify genes directly regulated by GLIS1, ChIP-Seq analysis was performed using TM5 cells stably expressing doxycycline (Dox)- inducible GLIS1-HA. ChIP analysis was performed as described previously^32, 46^. Cells were treated with and without Dox for 18 h and crosslinked with 1% formaldehyde in PBS for 10 min at RT and then quenched by glycine (final 125 mM) for 10 min at RT. Cells were washed two times with PBS and then sonicated for 40 min (S220 focused-ultrasonicator, Covaris, Woburn, MA). After removal of cell debris, chromatin was incubated overnight with HA antibody (Cell Signaling, #3724) and subsequently, incubated with Dynabeads Protein G (ThermoFisher Scientific, 10004D) for 3 h at 4°C to pulldown GLIS1-HA-chromatin complexes. The chromatin-bound beads were then washed and reverse crosslinked. Libraries were made with the ChIPed-DNA using Nextflex ChIP-Seq Library Prep kit (PerkinElmer). Sequencing reading was performed with a NovaSeq 6000 system (Illumina). TM5(-Dox) cells served as negative control to determine specificity^93^. UCSC Genome Browser Human Feb. 2009 (GRCh37/hg19) Assembly was used to generate the genome browser tracks.

### ChIP-seq analysis

ChIP-seq data was generated as single-end reads with a NovaSeq 6000 (Illumina). Raw sequence reads were filtered to remove any entries with a mean base quality score < 20. Adapters were removed via Cutadapt v1.12 with parameters “-a AGATCGGAAGAG -O 5 -q 0”, then reads were filtered to exclude those with length <30bp after trimming. Filtered and trimmed reads were mapped against the hg19 reference assembly (excluding haplotype chromosomes) via Bowtie v1.2, with only uniquely-mapped hits accepted. Duplicate mapped reads were removed by Picard tools MarkDuplicates.jar (v1.110). Initial peak calls were made with HOMER (v4.10.3) with parameters “-style factor -fdr 0.00001”, comparing each ChIP sample (Dox+ or Dox-) against its associated input sample. The Dox+ peak set was then filtered to exclude any peak that (a) overlapped a Dox- peak, (b) has fold change over input <8x (as reported by HOMER), or (c) has fold change over local signal <8x (as reported by HOMER). The Dox+ peaks were re-sized to 200bp centered on the called peak midpoints prior to downstream analysis. Enriched motifs were identified by HOMER ‘findMotifsGenome’ at “-size given” and all other parameters default. Coverage tracks for genome browser views were generated by extending each uniquely mapped non-duplicate read to the estimated average fragment size of 150bp, depth normalizing to 25M reads, then converting to bedGraph format with BEDtools v2.24.0 genomeCoverageBed and subsequently to bigwig format with UCSC utility bedGraphToBigWig v4.

### RNA-seq analysis

RNA-seq data was generated as paired-end reads with a NextSeq 5000 (Illumina). Raw sequence reads were filtered to remove any entries with a mean base quality score < 20 for either end in the pair. Filtered reads were then mapped to the hg19 reference assembly (excluding haplotype chromosomes) via STAR v2.5 with parameters “-- outSAMattrIHstart 0 --outFilterType BySJout --alignSJoverhangMin 8 --limitBAMsortRAM 55000000000 --outSAMstrandField intronMotif --outFilterIntronMotifs RemoveNoncanonical”. Counts per gene were determined via featureCounts (Subread v1.5.0-p1) with parameters “-s0 -Sfr” for Gencode V28 gene models. Differential analysis was performed with DESeq2 v1.14.1.

### Pathway analysis

Pathway analysis was performed via DAVID tools (v6.8) for KEGG pathway analysis^94, 95^.

### Immunostaining

Expression of GLIS1-βGAL fusion protein was examined by staining tissue sections with chicken anti-βGAL (1:500, ab9361, Abcam) and Alexa Fluor@ 488 donkey anti-chicken IgG (1:2000, A11039, Invitrogen) as described previously^96^. Expression of Anti-HA fusion protein was examined by staining cells with rabbit Anti-HA (1:250; #3724, Cell Signaling Technology) and Alexa Fluor@ 488 donkey anti-rabbit IgG (1:2000, A21208, Invitrogen). Fluorescence observed in a Zeiss LSM 710 confocal microscope.

### Genetic association analyses

To determine the association of genetic variants in the *GLIS1* region with glaucoma, we utilized the Genetic Epidemiology Research in Adult Health and Aging (GERA) cohort comprising of 4,986 POAG cases and 58,426 controls and a multiethnic UK Biobank (UKB; https://www.ukbiobank.ac.uk/) cohort consisting of 7,329 glaucoma (subtype unspecified) cases and 169,561 controls from five ethnic groups (European, East Asian, South Asian, African British, and mixed ancestries. The GERA cohort consists of 110,266 adult men and women, 18 years and older, who are of non-Hispanic white, Hispanic/Latino, Asian or African American ethnicity. Participants from the GERA cohort are members of the Kaiser Permanente Northern California (KPNC) integrated health care delivery system and provided self-reported information via the Research Program on Genes, Environment, and Health (RPGEH) survey. The UKB is a large prospective study following the health of approximately 500,000 participants from 5 ethnic groups (European, East Asian, South Asian, African British, and mixed ancestries) in the UK aged between the ages of 40 and 69. For UKB participants, demographic information and medical history were ascertained through touch-screen questionnaires. UKB participants also underwent a wide range of physical and cognitive assessments, including blood sampling. GERA individuals’ DNA samples were extracted using Oragene kits (DNA Genotek Inc., Ottawa, ON, Canada) at KPNC and genotyped at the Genomics Core Facility of UCSF. DNA samples were genotyped at over 665,000 genetic markers on four race/ethnicity specific Affymetrix Axiom arrays (Affymetrix, Santa Clara, CA, USA) optimized for European, Latino, East Asian, and African-American individuals. We performed genotype quality control (QC) procedures for the GERA samples on an array- wise basis. Briefly, we included genetic markers with initial genotyping call rate ≥ 97%, genotype concordance rate > 0.75 across duplicate samples, and allele frequency difference ≤ 0.15 between females and males for autosomal markers. Approximately 94% of samples and over 98% of genetic markers assayed reached QC procedures. Moreover, genetic markers with genotype call rates < 90% were excluded, as well as genetic markers with a MAF < 1%. We also performed imputation on an array-wise basis. Following the prephasing of genotypes with Shape-IT v2.r7271959, we imputed genetic markers from the cosmopolitan 1000 Genomes Project reference panel (phase I integrated release; http://1000genomes.org) using IMPUTE2 v2.3.060. We used the information r2 from IMPUTE2 as a QC parameter, which is an estimate of the correlation of the imputed genotype to the true genotype^9, 22, 97^. *GLIS1* gene locus was defined as ±500 kb upstream and downstream of the sequence using UCSC Genome Browser Assembly February 2009 (GRCh37/hg19). PLINK v1.9 was used to perform a logistic regression of the outcome and each SNP. Other statistical analyses and data management were performed in the language-and-environment R, version 3.6.0, using functions from the default libraries. All study procedures were approved by the KPNC Institutional Review Board and the protocols followed are compliant with specific Ethical Regulations. Written informed consent was obtained from all participants. The GLIS1 variant-level associations with glaucoma are fully disclosed in the manuscript (Supplementary Table 3 “Association of GLIS1 SNPs with POAG in the multiethnic meta-analysis (GERA+UKB)”). The meta-analysis GWAS summary statistics of glaucoma are available from the NHGRI-EBI GWAS Catalog study https://www.ebi.ac.uk/gwas/search?query=GCST006065.

### Statistical analysis

Data are presented as mean ± standard deviation (SD) and were analyzed using 2-tailed Student’s *t*-test using using Microsoft Excel and/or Prism 8.4 (GraphPad). To identify genetic variants in *GLIS1* associated with glaucoma, we performed logistic regression analysis adjusted for age, sex, and ancestry principal components.

### Data availability

The ChIP-seq and RNA-seq data described in this manuscript have been deposited in the NCBI Gene Expression Omnibus (GEO) with accession <u>GSE156846</u>.

The meta-analysis GWAS summary statistics of glaucoma are available from the NHGRI-EBI GWAS Catalog (https://www.ebi.ac.uk/gwas/downloads/summary-statistics), study accession number GCST006065 (https://www.ebi.ac.uk/gwas/search?query=GCST006065).

The MR imaging data will be accessible after registration at the following url (a password will be issued the following day that will provide access): https://civmvoxport.vm.duke.edu/voxbase/studyhome.php?studyid=733.

## Acknowledgements

AMJ research was supported by the Intramural Research Program of the National Institute of Environmental Health Sciences (NIEHS), the National Institutes of Health (NIH) [Z01-ES-100485]. The authors thank Laura Miller-de Graff for her outstanding assistance with breeding the knockout mice. This work was supported in part by the National Eye Institute (NEI) grants R01 EY027004 (HC, EJ) EY022891 (KSN), EY028175 (KK), EY011721 (SWMJ), EY026220 EY026177 (GZ), and EY12731291 (TB), the National Institute of Diabetes and Digestive and Kidney Diseases (NIDDK) grant R01 DK116738 (HC, EJ), NEI P30 EY002162 core grant for vision research (UCSF, Ophthalmology), Research to Prevent Blindness unrestricted grant (UCSF Ophthalmology), and grants from That Man May See Inc., Brightfocus Foundation (G2019360), UCSF Academic Senate Committee on Research (RAP grant), Marin Community Foundation-Kathlyn Masneri and Arno Masneri Fund (KSN). SC is supported by grant from Department of Biotechnology, Government of India grant (BT/PR32404/MED/30/2136/2019). SWMJ is an HHMI investigator and received funding from the Precision Medicine Initiative at Columbia University.

## Contributions

HSK, RVB, and CB performed the preliminary characterization of *Glis1*-KO mice and initiated the project; AB, RVB, CS, and GAJ analyzed AqH dynamics by MRI; RVB and CS carried out IOP analysis; CS and HSK carried out TM cell culture, gene expression, and ChIP-Seq analysis; YK, SK, YZ, and KSN performed ocular histological analysis; KSN, HC, JY, SC, and EJ, performed genetic association analyses; SG, performed the bioinformatic analyses; KO, analyzed GLIS1 transcriptional activity; GZ and TB, provided primary hTM cells and advice; KK, GC, SWMJ, analyzed the SC and provided advice on the ocular histology data; RVB, CS, KSN and AMJ, designed experiments; AMJ and KSN wrote the manuscript. RVB, CS, SG, TB, SK, HC, KK, SJ, and EJ contributed and reviewed the manuscript; AMJ oversaw the characterization of the initial mouse phenotype and molecular studies and KSN the eye structural and human genetic analyses.

## SUPPLEMENTARY INFORMATION

Supplementary Figures 1-14

Supplementary Tables 1-3

Source files:

Source file for imaging data in Figure 1a and c.

Source file for Figure 1b and b and Supplementary Figs. 3 and 4c Source file for Figure 3a and b

Source file for Figure 4e and f

Source file for Supplementary Figure 1b, c and d

Source file for Supplementary Figure 2

Source file for Supplementary Figure 5

Source file for Supplementary Figure 6

Source file for Supplementary Figure 7

Source file for Supplementary Figure 10

Source file for Supplementary Figure 11

Source file for Supplementary Figure 12

Source file for Supplementary Figure 13

Source file for Supplementary Figure 14

## REFERENCES

1. Weinreb, R. N., Aung, T., Medeiros, F. A. The pathophysiology and treatment of glaucoma: a review. Jama 311, 1901–1911 (2014).

2. Quigley, H. A., Broman, A. T. The number of people with glaucoma worldwide in 2010 and 2020. Br. J. Ophthalmol. 90, 262–267 (2006).

3. Weinreb, R. N., et al. Primary open-angle glaucoma. Nat. Rev. Dis. Primers doi: 10.1038/nrdp.2016.67 (2016).

4. Wiggs, J. L., Pasquale, L. R. Genetics of glaucoma. Hum. Mol. Genet. 26, R21–R27 (2017).

5. Choquet, H., Wiggs, J. L., Khawaja, A. P. Clinical implications of recent advances in primary open-angle glaucoma genetics. Eye (Lond) 34, 29–39 (2020).

6. Lewis, C. J., Hedberg-Buenz, A., DeLuca, A. P., Stone, E. M., Alward, W. L. M., Fingert, J. H. Primary congenital and developmental glaucomas. Hum. Mol. Genet. 26, R28–R36 (2017).

7. Libby, R. T., Gould, D. B., Anderson, M. G., John, S. W. Complex genetics of glaucoma susceptibility. Annu. Rev. Genomics Hum. Genet. 6, 15–44 (2005).

8. Bailey, J. N., et al. Genome-wide association analysis identifies TXNRD2, ATXN2 and FOXC1 as susceptibility loci for primary open-angle glaucoma. Nat. Genet. 48, 189-194 (2016).

9. Choquet, H., et al. A multiethnic genome-wide association study of primary open- angle glaucoma identifies novel risk loci. Nat. Commun. doi: 10.1038/s41467-018-04555-4 (2018).

10. Costagliola, C., et al. How many aqueous humor outflow pathways are there? Surv. Ophthalmol. 65, 144–170 (2020).

11. Abu-Hassan, D. W., Acott, T. S., Kelley, M. J. The Trabecular Meshwork: A Basic Review of Form and Function. J. Ocul. Biol. 2, doi: 10.13188/2334-2838.1000017 (2014).

12. Sacca, S. C., Pulliero, A., Izzotti, A. The dysfunction of the trabecular meshwork during glaucoma course. J. Cell. Physiol. 230, 510–525 (2015).

13. Alvarado, J., Murphy, C., Juster, R. Trabecular meshwork cellularity in primary open-angle glaucoma and nonglaucomatous normals. Ophthalmology 91, 564–579 (1984).

14. Grierson, I., Hogg, P. The proliferative and migratory activities of trabecular meshwork cells. Progr. Retin. Eye Res. 15, 33–67 (1995).

15. Stamer, W. D., Clark, A. F. The many faces of the trabecular meshwork cell. Exp. Eye Res. 158, 112–123 (2017).

16. Youngblood, H., Hauser, M. A., Liu, Y. Update on the genetics of primary open-angle glaucoma. Exp. Eye Res. 188, 107795 (2019).

17. Bonnemaijer, P. W. M., et al. Genome-wide association study of primary open-angle glaucoma in continental and admixed African populations. Hum. Genet, 137, 847–862 (2018).

18. Khawaja, A. P., et al. Genome-wide analyses identify 68 new loci associated with intraocular pressure and improve risk prediction for primary open-angle glaucoma. Nat. Genet. 50, 778–782 (2018).

19. Khor, C. C., et al. Genome-wide association study identifies five new susceptibility loci for primary angle closure glaucoma. Nat. Genet. 48, 556–562 (2016).

20. Nongpiur, M. E., et al. Evaluation of Primary Angle-Closure Glaucoma Susceptibility Loci in Patients with Early Stages of Angle-Closure Disease. Ophthalmology, 125, 664–670 (2018).

21. Shiga, Y., et al. Genome-wide association study identifies seven novel susceptibility loci for primary open-angle glaucoma. Hum. Mol. Genet. 27, 1486–1496 (2018).

22. Choquet, H., et al. A large multi-ethnic genome-wide association study identifies novel genetic loci for intraocular pressure. Nat. Commun. doi: 10.1038/s41467-017-01913-6 (2017).

23. Huang, L., et al. Genome-wide analysis identified 17 new loci influencing intraocular pressure in Chinese population. Sci. China Life Sci. 62, 153–164 (2019).

24. Zhuang, W., et al. Genotype-ocular biometry correlation analysis of eight primary angle closure glaucoma susceptibility loci in a cohort from Northern China. PLoS One 13, e0206935 (2018).

25. Jetten, A. M. GLIS1-3 transcription factors: critical roles in the regulation of multiple physiological processes and diseases. Cell Mol. Life Sci. 75, 3473–3494 (2018).

26. Scoville, D. W., Kang, H. S., Jetten, A. M. Transcription factor GLIS3: Critical roles in thyroid hormone biosynthesis, hypothyroidism, pancreatic beta cells and diabetes. Pharmacol. Ther. doi: 10.1016/j.pharmthera.2020.107632 (2020).

27. Dimitri, P. The role of GLIS3 in thyroid disease as part of a multisystem disorder. Best Pract. Res. Clin. Endocrinol. Metab. 31, 175–182 (2017).

28. Lim, S. H., et al. CYP1B1, MYOC, and LTBP2 mutations in primary congenital glaucoma patients in the United States. Am. J. Ophthalmol. 155, 508–517.e505 (2013).

29. Kim, Y. S., Lewandoski, M., Perantoni, A. O., Kurebayashi, S., Nakanishi, G., Jetten, A. M. Identification of Glis1, a novel Gli-related, Kruppel-like zinc finger protein containing transactivation and repressor functions. J. Biol. Chem. 277, 30901–30913 (2002).

30. Nakashima, M., et al. A novel gene, GliH1, with homology to the Gli zinc finger domain not required for mouse development. Mech. Dev. 119, 21–34 (2002).

31. Dong, Y. H., Fu, D. G. Autoimmune thyroid disease: mechanism, genetics and current knowledge. Eur. Rev. Med. Pharmacl. Sci. 18, 3611–3618 (2014).

32. Kang, H. S., et al. GLIS3 is indispensable for TSH/TSHR-dependent thyroid hormone biosynthesis and follicular cell proliferation. J. Clin. Invest. 127, 4326–4337 (2017).

33. Nikiforova, M. N., et al. GLIS Rearrangement is a Genomic Hallmark of Hyalinizing Trabecular Tumor of the Thyroid Gland. Thyroid 29, 161–173 (2019).

34. Ho, L. C., et al. In vivo assessment of aqueous humor dynamics upon chronic ocular hypertension and hypotensive drug treatment using gadolinium-enhanced MRI. Invest. Ophthalmol. Vis. Sci. 55, 3747–3757 (2014).

35. Crosbie, D. E., Keaney, J., Tam, L. C. S., Daniel Stamer, W., Campbell, M., Humphries, P. Age-related changes in eye morphology and aqueous humor dynamics in DBA/2J mice using contrast-enhanced ocular MRI. Magn. Reson. Imaging 59, 10–16 (2019).

36. Kaneko, Y., et al. Effects of K-115 (Ripasudil), a novel ROCK inhibitor, on trabecular meshwork and Schlemm’s canal endothelial cells. Sci. Rep. doi: 10.1038/srep19640 (2016).

37. Libby, R. T., et al. Modification of ocular defects in mouse developmental glaucoma models by tyrosinase. Science 299, 1578–1581 (2003).

38. Smith, R. S., et al. Haploinsufficiency of the transcription factors FOXC1 and FOXC2 results in aberrant ocular development. Hum. Mol. Genet. 9, 1021–1032 (2000).

39. Sethi, A., Mao, W., Wordinger, R. J., Clark, A. F. Transforming growth factor-beta induces extracellular matrix protein cross-linking lysyl oxidase (LOX) genes in human trabecular meshwork cells. Invest. Ophthalmol. Vis. Sci. 52, 5240–5250 (2011).

40. Carnes, M. U., Allingham, R. R., Ashley-Koch, A., Hauser, M. A. Transcriptome analysis of adult and fetal trabecular meshwork, cornea, and ciliary body tissues by RNA sequencing. Exp. Eye Res. 167, 91–99 (2018).

41. Borras, T. Mechanosensitive Genes in the Trabecular Meshwork at Homeostasis. In: Ophthalmology Research: Mechanisms of the Glaucomas (eds Tombran-Tink J., Barnstable C. J., Shields M. B.). Humana Press (2008).

42. Filla, M. S., Dimeo, K. D., Tong, T., Peters, D. M. Disruption of fibronectin matrix affects type IV collagen, fibrillin and laminin deposition into extracellular matrix of human trabecular meshwork (HTM) cells. Exp. Eye Res. 165, 7–19 (2017).

43. Mackay, D. S., Bennett, T. M., Shiels, A. Exome Sequencing Identifies a Missense Variant in EFEMP1 Co-Segregating in a Family with Autosomal Dominant Primary Open-Angle Glaucoma. PLoS One doi: 10.1371/journal.pone.0132529 (2015).

44. Kuchtey, J., et al. Mapping of the disease locus and identification of ADAMTS10 as a candidate gene in a canine model of primary open angle glaucoma. PLoS Genet. doi: 10.1371/journal.pgen.1001306 (2011).

45. Moazzeni, H., Mirrahimi, M., Moghadam, A., Banaei-Esfahani, A., Yazdani, S., Elahi, E. Identification of genes involved in glaucoma pathogenesis using combined network analysis and empirical studies. Hum. Mol. Genet. 28, 3637–3663 (2019).

46. Jeon, K., Kumar, D., Conway, A. E., Park, K., Jothi, R., Jetten, A. M. GLIS3 Transcriptionally Activates WNT Genes to Promote Differentiation of Human Embryonic Stem Cells into Posterior Neural Progenitors. Stem Cells 37, 202–215 (2019).

47. Scoville, D., Lichti-Kaiser, K., Grimm, S., Jetten, A. GLIS3 binds pancreatic beta cell regulatory regions alongside other islet transcription factors. J. Endocrinol. 243, 1–14 (2019).

48. Jin, Y., Liang, Z., Lou, H. The Emerging Roles of Fox Family Transcription Factors in Chromosome Replication, Organization, and Genome Stability. Cells doi: 10.3390/cells9010258 (2020).

49. Lin, K. C., Park, H. W., Guan, K. L. Regulation of the Hippo Pathway Transcription Factor TEAD. Trends Biochem. Sci. 42, 862–872 (2017).

50. Peng, J., Wang, H., Wang, X., Sun, M., Deng, S., Wang, Y. YAP and TAZ mediate steroid-induced alterations in the trabecular meshwork cytoskeleton in human trabecular meshwork cells. Int. J. Mol. Med. 41, 164–172 (2018).

51. Wang, X., et al. Mutual regulation of the Hippo/Wnt/LPA/TGFbeta signaling pathways and their roles in glaucoma (Review). Int. J. Mol. Med. 41, 1201–1212 (2018).

52. Kirstein, L., Cvekl, A., Chauhan, B. K., Tamm, E. R. Regulation of human myocilin/TIGR gene transcription in trabecular meshwork cells and astrocytes: role of upstream stimulatory factor. Genes Cells 5, 661–676 (2000).

53. Hwang, Y. P., et al. WY-14643 Regulates CYP1B1 Expression through Peroxisome Proliferator-Activated Receptor alpha-Mediated Signaling in Human Breast Cancer Cells. Int. J. Mol. Sci. doi: 10.3390/ijms20235928 (2019).

54. Zheng, W., Jefcoate, C. R. Steroidogenic factor-1 interacts with cAMP response element-binding protein to mediate cAMP stimulation of CYP1B1 via a far upstream enhancer. Mol. Pharmacol. 67, 499–512 (2005).

55. Liesenborghs, I., et al. Comprehensive bioinformatics analysis of trabecular meshwork gene expression data to unravel the molecular pathogenesis of primary open-angle glaucoma. Acta Ophthalmol. 98, 48–57 (2020).

56. Borras, T. Gene expression in the trabecular meshwork and the influence of intraocular pressure. Prog. Retin. Eye Res. 22, 435–463 (2003).

57. Liu, Y., Allingham, R. R. Major review: Molecular genetics of primary open-angle glaucoma. Exp. Eye Res. 160, 62–84 (2017).

58. Gharahkhani, P., et al. Genome-wide meta-analysis identifies 127 open-angle glaucoma loci with consistent effect across ancestries. Nat. Commun. 12, doi: 10.1038/s41467-020-20851-4 (2021).

59. Chatterjee, A., Villarreal, G., Jr., Rhee, D. J. Matricellular proteins in the trabecular meshwork: review and update. J. Ocul. Pharmacol. Ther. 30, 447–463 (2014).

60. Comes, N., Borras, T. Individual molecular response to elevated intraocular pressure in perfused postmortem human eyes. Physiol. Genomics 38, 205–225 (2009).

61. Teixeira, L. B., Zhao, Y., Dubielzig, R. R., Sorenson, C. M., Sheibani, N. Ultrastructural abnormalities of the trabecular meshwork extracellular matrix in Cyp1b1-deficient mice. Vet. Pathol. 52, 397–403 (2015).

62. Filla, M. S., Faralli, J. A., Peotter, J. L., Peters, D. M. The role of integrins in glaucoma. Exp. Eye Res. 158, 124–136 (2017).

63. WuDunn, D. Mechanobiology of trabecular meshwork cells. Exp. Eye Res. 88, 718–723 (2009).

64. Vranka, J. A., Kelley, M. J., Acott, T. S., Keller, K. E. Extracellular matrix in the trabecular meshwork: intraocular pressure regulation and dysregulation in glaucoma. Exp. Eye Res. 133, 112–125 (2015).

65. Acott, T. S., Kelley, M. J. Extracellular matrix in the trabecular meshwork. Exp. Eye Res. 86, 543–561 (2008).

66. O’Callaghan, J., Cassidy, P. S., Humphries, P. Open-angle glaucoma: therapeutically targeting the extracellular matrix of the conventional outflow pathway. Expert Opin. Ther. Targets 21, 1037–1050 (2017).

67. Saeedi, O., Yousaf, S., Tsai, J., Palmer, K., Riazuddin, S., Ahmed, Z. M. Delineation of Novel Compound Heterozygous Variants in LTBP2 Associated with Juvenile Open Angle Glaucoma. Genes (Basel) doi: 10.3390/genes9110527 (2018).

68. Thomson, J., Singh, M., Eckersley, A., Cain, S. A., Sherratt, M. J., Baldock, C. Fibrillin microfibrils and elastic fibre proteins: Functional interactions and extracellular regulation of growth factors. Semin. Cell Dev. Biol. 89, 109–117 (2019).

69. Wordinger, R. J., Clark, A. F. Lysyl oxidases in the trabecular meshwork. J Glaucoma 23, S55–58 (2014).

70. Kuchtey, J., Kuchtey, R. W. The microfibril hypothesis of glaucoma: implications for treatment of elevated intraocular pressure. J. Ocul. Pharmacol. Ther. 30, 170–180 (2014).

71. Dzamba, B. J., DeSimone, D. W. Extracellular Matrix (ECM) and the Sculpting of Embryonic Tissues. Curr. Top. Dev. Biol. 130, 245–274 (2018).

72. Zode, G. S., et al. Reduction of ER stress via a chemical chaperone prevents disease phenotypes in a mouse model of primary open angle glaucoma. J. Clin. Invest. 121, 3542–3553 (2011).

73. Wang, H., Li, M., Zhang, Z., Xue, H., Chen, X., Ji, Y. Physiological function of myocilin and its role in the pathogenesis of glaucoma in the trabecular meshwork (Review). Int. J. Mol. Med. 43, 671–681 (2019).

74. Kim, B. S., et al. Targeted disruption of the myocilin gene (Myoc) suggests that human glaucoma-causing mutations are gain of function. Mol. Cell. Biol. 21, 7707–7713 (2001).

75. Polansky, J. R. Current perspectives on the TIGR/MYOC gene (Myocilin) and glaucoma. Ophthalmol. Clin. North Am. 16, 515–527, v-vi (2003).

76. Saura, M., Cabana, M., Ayuso, C., Valverde, D. Mutations including the promoter region of myocilin/TIGR gene. Eur. J. Hum. Genet. 13, 384–387 (2005).

77. Paylakhi, S. H., et al. FOXC1 in human trabecular meshwork cells is involved in regulatory pathway that includes miR-204, MEIS2, and ITGbeta1. Exp. Eye Res. 111, 112–121 (2013).

78. Souzeau, E., et al. Glaucoma spectrum and age-related prevalence of individuals with FOXC1 and PITX2 variants. Eur. J. Hum. Genet. 25, 839–847 (2017).

79. Ho, L. T. Y., Skiba, N., Ullmer, C., Rao, P. V. Lysophosphatidic Acid Induces ECM Production via Activation of the Mechanosensitive YAP/TAZ Transcriptional Pathway in Trabecular Meshwork Cells. Invest. Ophthalmol. Vis. Sci. 59, 1969–1984 (2018).

80. Wang, F., et al. RNAscope: a novel in situ RNA analysis platform for formalin-fixed, paraffin-embedded tissues. J. Mol. Diagn. 14, 22–29 (2012).

81. Labelle-Dumais, C., et al. Loss of PRSS56 function leads to ocular angle defects and increased susceptibility to high intraocular pressure. Dis. Model Mech. doi: 10.1242/dmm.042853 (2020).

82. Kizhatil, K., Ryan, M., Marchant, J. K., Henrich, S., John, S. W. Schlemm’s canal is a unique vessel with a combination of blood vascular and lymphatic phenotypes that forms by a novel developmental process. PLoS Biol. doi: 10.1371/journal.pbio.1001912 (2014).

83. Emmenlauer, M., et al. XuvTools: free, fast and reliable stitching of large 3D datasets. J. Microsc. 233, 42–60 (2009).

84. Postma, M., Goedhart, J. PlotsOfData-A web app for visualizing data together with their summaries. PLoS Biol. doi: 10.1371/journal.pbio.3000202 (2019).

85. Tweedle, M. F., et al. Dependence of MR signal intensity on Gd tissue concentration over a broad dose range. Magn. Reson. Med. 22, 191–194 (1991).

86. Takeda, M., et al. Concentration of gadolinium-diethylene triamine pentaacetic acid in human kidney--study on proper time for dynamic magnetic resonance imaging of the human kidney on low and high magnetic fields. Tohoku J. Exp. Med. 171, 119–128 (1993).

87. Verma, S., et al. Overview of dynamic contrast-enhanced MRI in prostate cancer diagnosis and management. AJR Am. J. Roentgenol. 198, 1277–1288 (2012).

88. Morkenborg, J., Pedersen, M., Jensen, F. T., Stodkilde-Jorgensen, H., Djurhuus, J. C., Frokiaer, J. Quantitative assessment of Gd-DTPA contrast agent from signal enhancement: an in-vitro study. Magn. Reson. Imaging 21, 637–643 (2003).

89. Beak, J. Y., Kang, H. S., Kim, Y. S., Jetten, A. M. Functional analysis of the zinc finger and activation domains of Glis3 and mutant Glis3(NDH1). Nucleic Acids Res. 36, 1690–1702 (2008).

90. Pang, I. H., Shade, D. L., Clark, A. F., Steely, H. T., DeSantis, L. Preliminary characterization of a transformed cell strain derived from human trabecular meshwork. Curr. Eye Res. 13, 51–63 (1994).

91. Meerbrey, K. L., et al. The pINDUCER lentiviral toolkit for inducible RNA interference in vitro and in vivo. Proc. Natl. Acad. Sci. USA 108, 3665–3670 (2011).

92. Schindelin, J., et al. Fiji: an open-source platform for biological-image analysis. Nat. Methods 9, 676–682 (2012).

93. Narlikar, L., Jothi, R. ChIP-Seq data analysis: identification of protein-DNA binding sites with SISSRs peak-finder. Methods Mol. Biol. 802, 305–322 (2012).

94. Huang da, W., Sherman, B. T., Lempicki, R. A. Systematic and integrative analysis of large gene lists using DAVID bioinformatics resources. Nat. Protoc. 4, 44–57 (2009).

95. Huang da, W., Sherman, B. T., Lempicki, R. A. Bioinformatics enrichment tools: paths toward the comprehensive functional analysis of large gene lists. Nucleic Acids Res. 37, 1–13 (2009).

96. Zhao, F., et al. Elimination of the male reproductive tract in the female embryo is promoted by COUP-TFII in mice. Science 357, 717–720 (2017).

97. Kvale, M. N., et al. Genotyping Informatics and Quality Control for 100,000 Subjects in the Genetic Epidemiology Research on Adult Health and Aging (GERA) Cohort. Genetics 200, 1051–1060 (2015).

