## Supplemental data for "GLIS1 regulates trabecular meshwork function and intraocular pressure and is associated with glaucoma in humans"

### SUPPLEMENTARY FILES

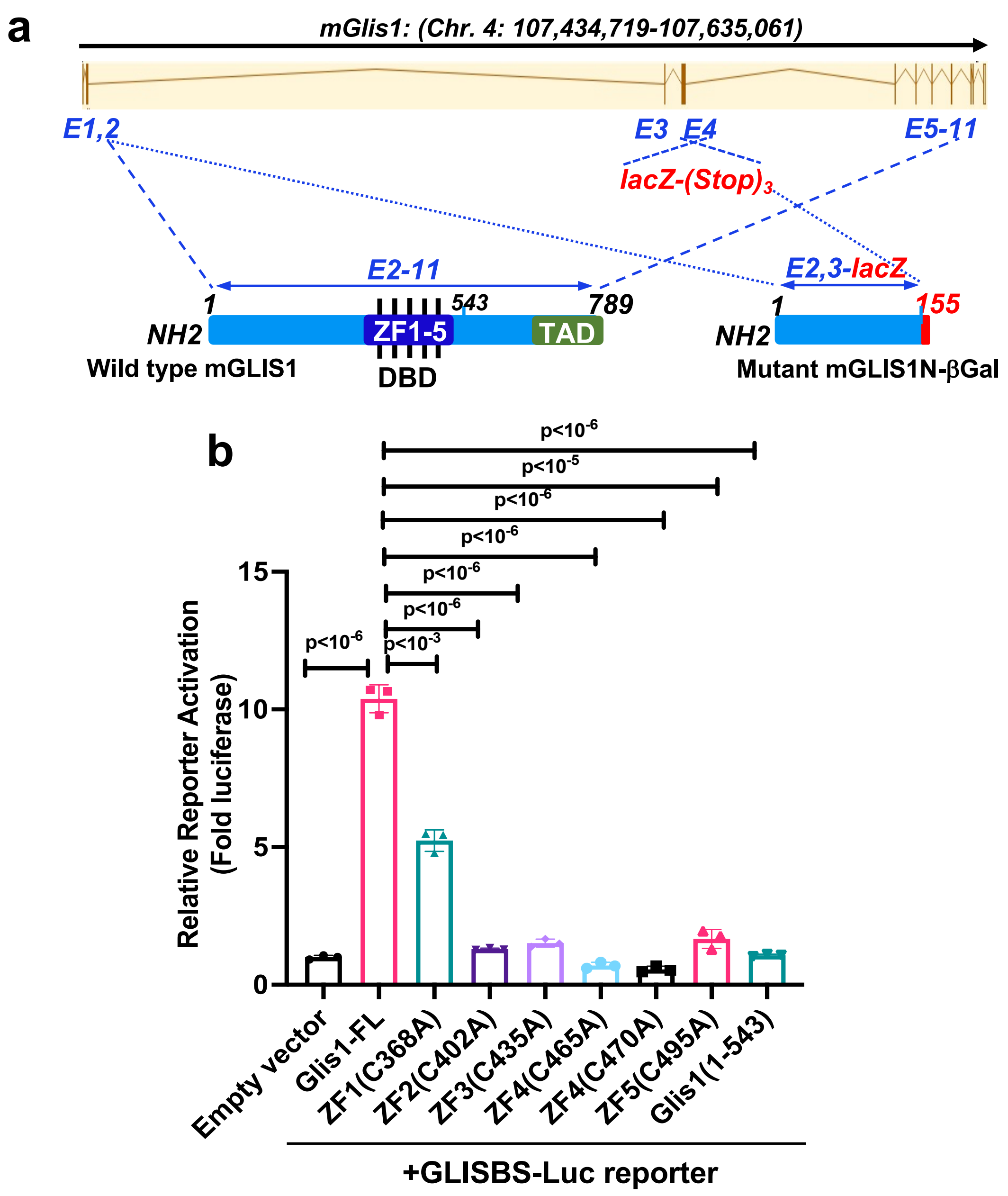

Supplementary Figure 1a, b.

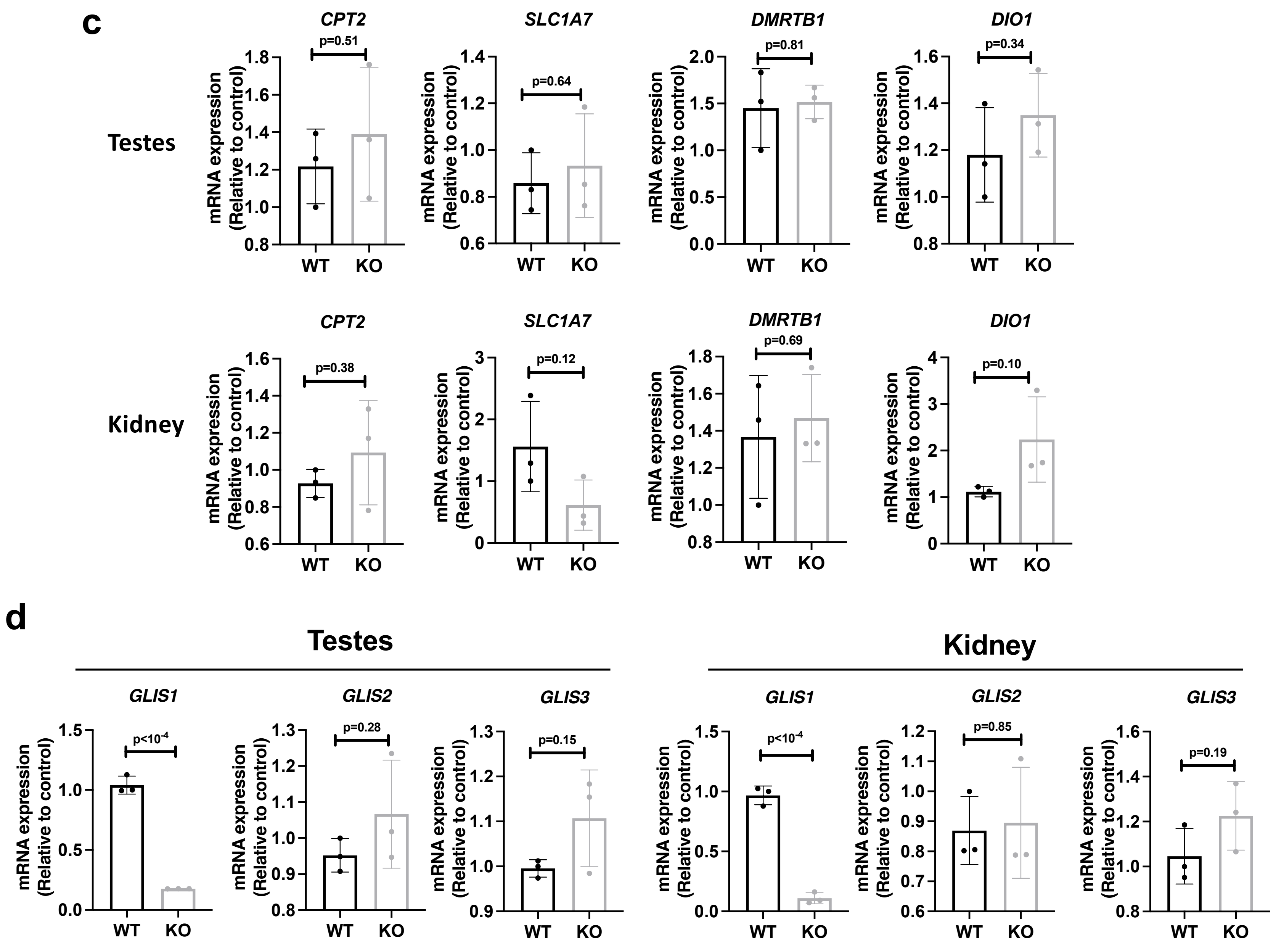

Supplementary Fig. 1c, d

**Supplementary Figure 1. a. Genomic map of the mouse *Glis1* gene.** In *Glis1*-KO mice 840bp of exon 4 is replaced by lacZ-neo cassette (lacZ-(Stop)3) resulting in the deletion of the DBD and TAD as indicated. This generates a GLIS1ΔC-βGal fusion protein. **b.** The ZF motifs (DBD) and TAD are critical for Glis1 transcriptional activity. The effect of ZF mutations and loss of the TAD on GLIS1 transcriptional activity was examined in HEK293T cells co-transfected with β-Gal and GLISBS-LUC reporters, and the expression plasmid indicated. The relative Luc reporter activation was determined and plotted (n=3, independent replicates). **c, d**. Deletion of exon 4 in *Glis1*-KO mice had no significant effect on the expression of the *Glis1* neighboring genes, *Dmrtb1, Slc1a7, Dio1,* and *Cpt2* (**c**), nor the expression of *Glis2* and *Glis3* (**d**) in *Glis1*-KO kidneys and testes. Data in **b-d** are represented as means ± SD. Statistical analyses were performed with two-tailed Student's t-test. P-values are indicated above bars.

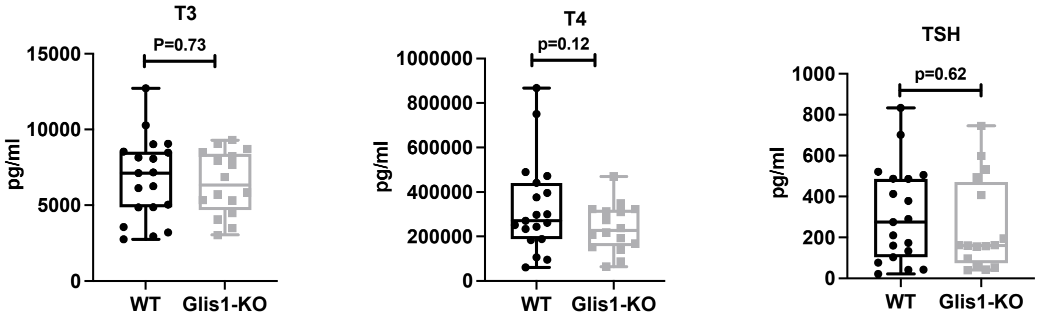

**Supplementary Figure 2. *Glis1*-KO eye phenotype is not related to Graves' disease.** Blood T3 and T4, and TSH levels in 3 months old WT (n=19 mice) and *Glis1-KO* mice (n=16 mice) were assayed. Data are presented as box plots with quartiles and ranges. Statistical analyses were performed with two-tailed Student's t-test. Median, upper, and lower quartile values: for T3 in WT (7125, 8544, 4860); KO (6339, 8405, 4688); T4 in WT (270415, 441439, 188291); KO (227577, 320409, 156793); TSH in WT (275, 487, 104); KO (161.5, 472, 74.25). Confidence intervals (CI) for T3, T4, and TSH were all 95%.

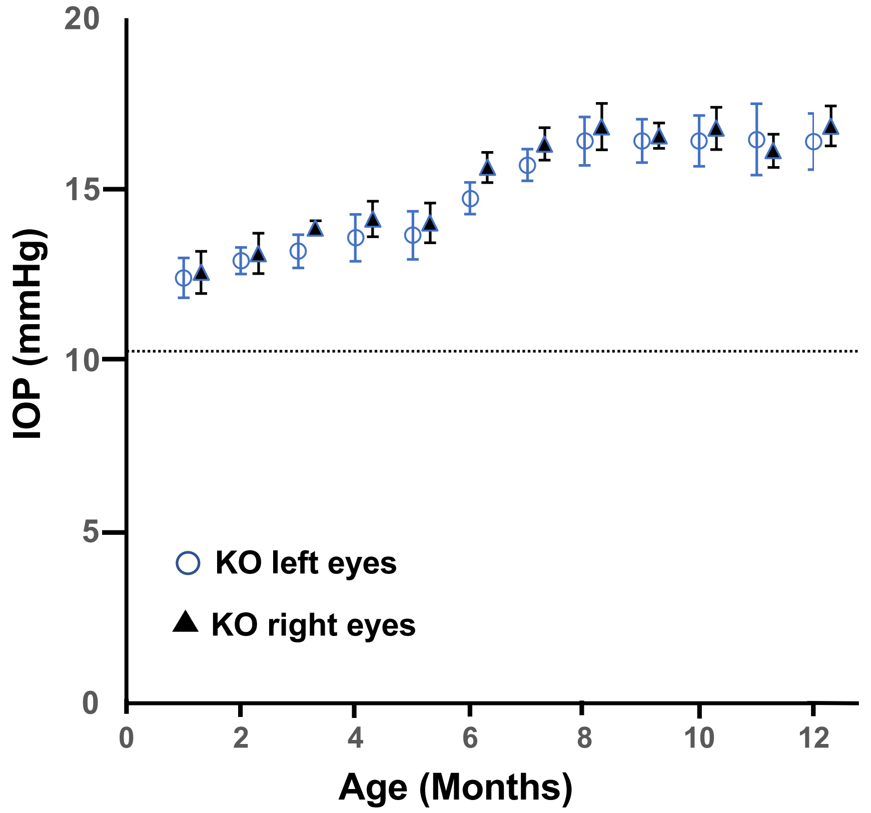

**Supplementary Figure 3. Comparison of IOP levels between left and right eyes of male *Glis1*-KO mice as function of age.** IOP levels in left and right eyes were measured. Number of mice examined: at 1 and 2 months (n=3); 3 (n=5); 4 and 7 months (n=8); 5 (n=6); 6, 11, and 12 months (n=7)(4 IOP measurements/eye/timepoint). Dotted line indicates basal IOP level in 1-3 months old male WT mice. Data are represented as means ± SD. Statistical analyses were performed with two-tailed Student's t-test. No significant statistical differences were found between right (triangles) and left (circles) eyes. p values at 1 to 12 months are 1.0, 0.91, 0.12, 0.44, 0.76, 0.044, 0.29, 0.75, 0.36, 0.94, 0.13, 0.87, respectively.

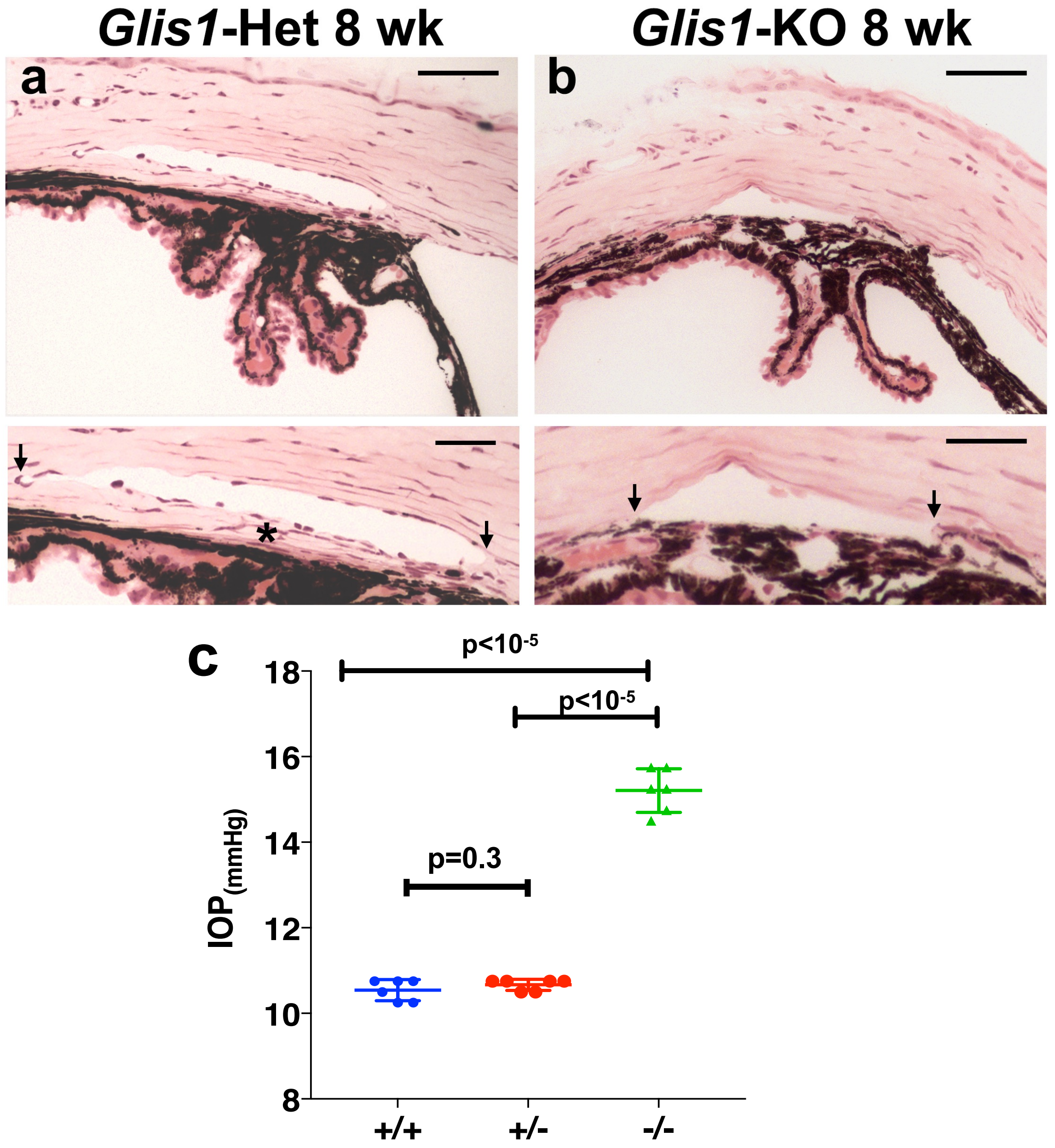

**Supplementary Figure 4. *Glis1*-heterozygous mice exhibit a well-developed ocular drainage structure.**

**a, b.** Representative histological images of the ocular angle structure from 8-weeks-old *Glis1*-heterozygous **(a)**

and *Glis1*-KO mice **(b)** maintained in C57BL/6NCrl background. *Glis1*-heterozygous eyes showed a well-developed SC and TM (*). In contrast, age-matched *Glis1*-KO eyes exhibited substantial thinning of the TM. Arrows show edges of the SC. The histological assessment was performed on 6 eyes for each genotype with similar results. Scale bar = 50 μm in upper panel and 25 μm lower panel of images in **a** and **b**. **c.** IOP comparison. No significant difference in IOP was observed between 6 months-old WT and heterozygous mice, while homozygous *Glis1*-KO mice showed significantly elevated IOP. Data are represented as means ± SD (p=0.3; 6 eyes each, IOP measured four times in each eye). Statistical analyses were performed with two-tailed Student's t-test.

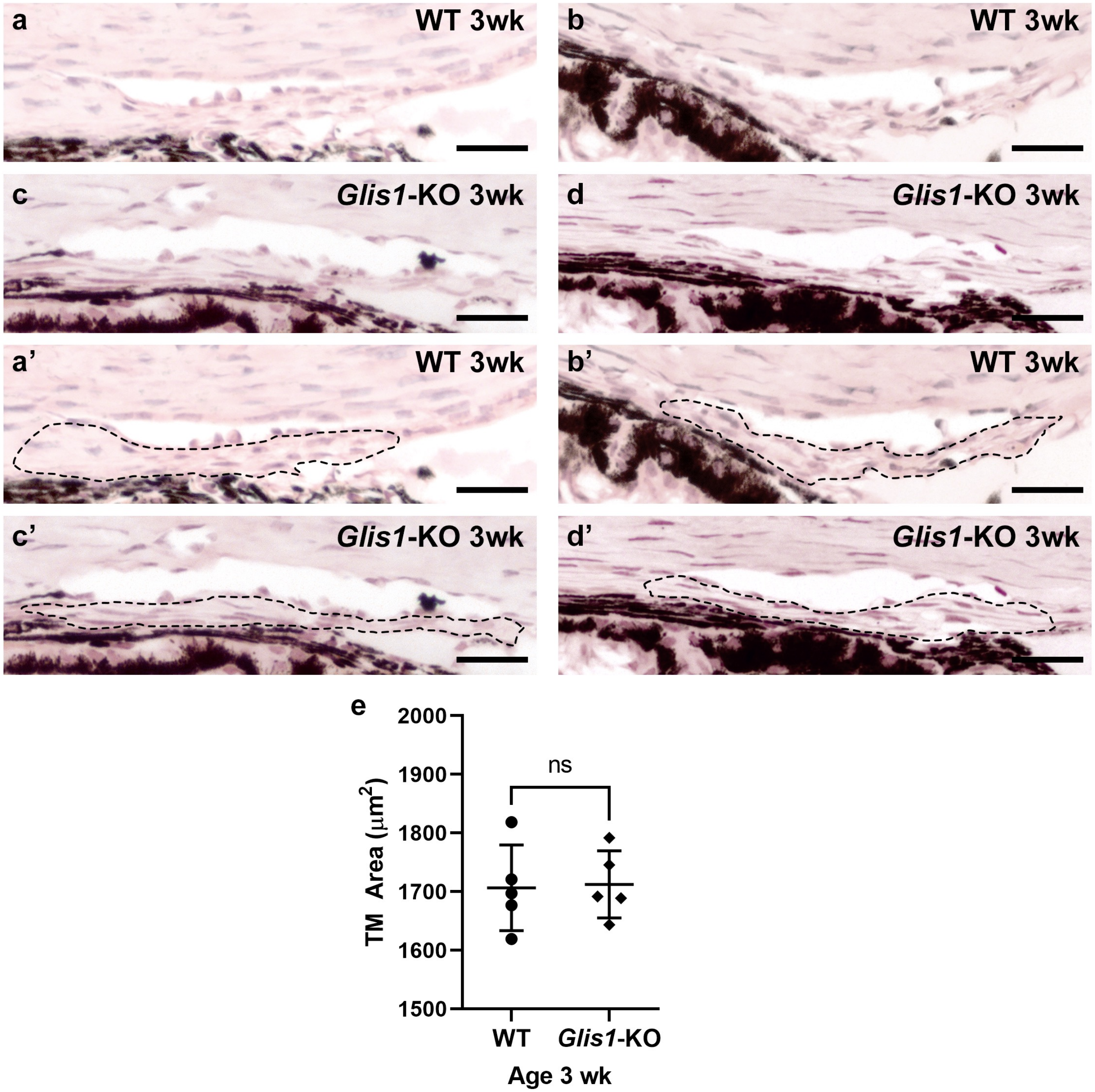

**Supplementary Figure 5. Comparison of the TM status of WT and *Glis1*-KO mice.** Representative histological images of the ocular angle structure from 3 weeks old WT (**a,b**) and *Glis1*-KO (**c,d**) mice. Lower panel shows the region selected for measuring the TM area in WT (**a',b'**) and *Glis1*-KO (**c',d'**). Images from three different mice are shown. **e**. Scatter plot showing TM area of 3 weeks-old WT and *Glis1-KO* mice (n = 5 eyes per group). Scale bar = 25 µm. Statistical analyses were performed with two-tailed Student's t-test. At 3 weeks of age, the TM area in *Glis1*-KO is not significantly (ns) different from that in WT mice (p=0.87).

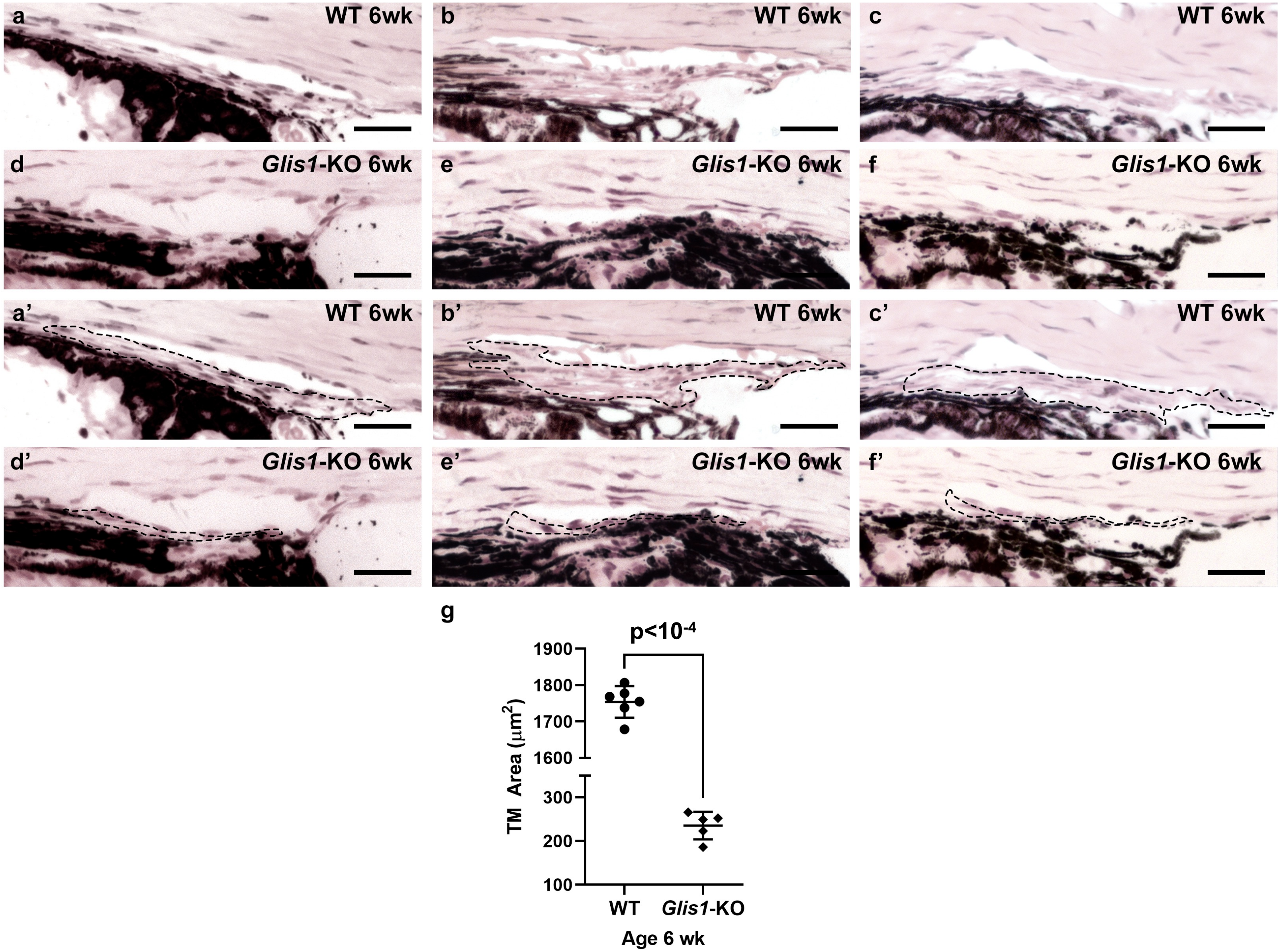

**Supplementary Figure 6. TM is substantially thinner in *Glis1*-KO mice at 6 weeks of age.** Representative histological images of the ocular angle structure from 6 weeks-old WT (**a,b,c**) and *Glis1*-KO (**d,e,f**) mice. The region selected for measuring the area of the TM is outlined in WT (**a’,b’,c’**) and *Glis1*-KO (**d’,e’,f’**) mice are shown in the lower panel. Images from three different mice are shown. **g.** Scatter plot showing TM area of 6 weeks-old WT and *Glis1-KO* mice (n = 5 eyes per group). The TM area in *Glis1*-KO is significantly smaller compared to that in WT mice. Statistical analyses were performed with two-tailed Student's t-test. Data are represented as means ± SD. $p<10^{-4}$. Scale bar = 25 μm.

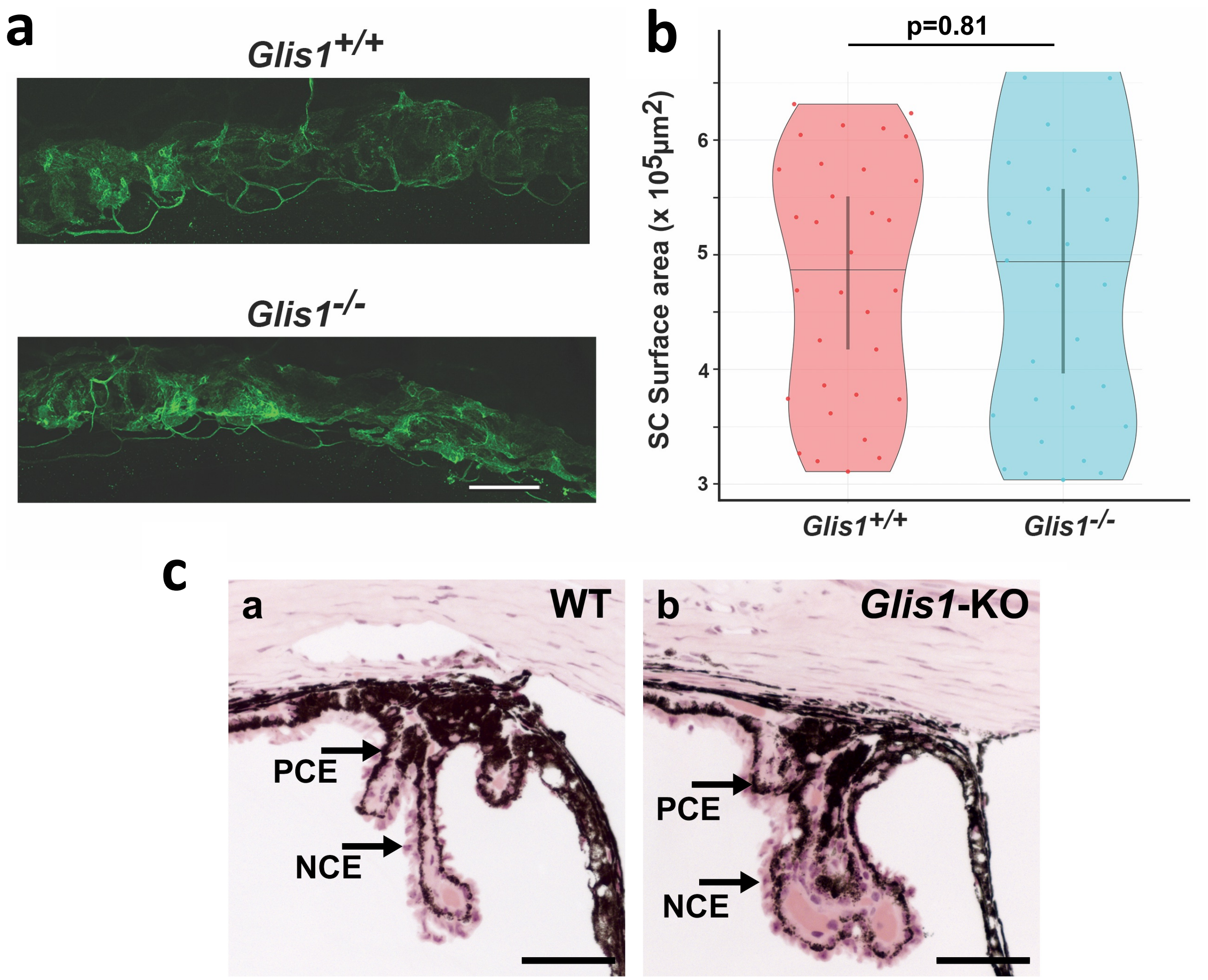

**Supplementary Figure 7. The organization of SC and ciliary body is unaffected in GLIS1 null mice.** **a**. Representative image of endomucin stained SC from a WT control mouse and *Glis1*-KO mouse at 4 weeks of age. No significant difference was observed. **b**. Quantification of volume of SC in control and *Glis1*-KO mice show no significant difference. Endomucin stained SC volume was computed using Imaris. Graph is mean and 95% confidence interval (CI) of mean. Each dot is the surface of SC in a quadrant of an eye (4 quadrants/eye). In total eight eyes per genotype were measured (a total of 32 measurements per genotype). Statistical analyses were performed with two-tailed Student's t-test. The volume of SC was not significantly different between *Glis1*-KO and wild type mice (p=0.81). Scale bar = 100µm. **c.** Representative histological image of the ciliary body of 3 months-old wild type and *Glis1*-KO mice showing normal organization of the pigmented ciliary epithelium (PCE) and nonpigmented ciliary epithelium (NCE). Scale bar = 50 µm.

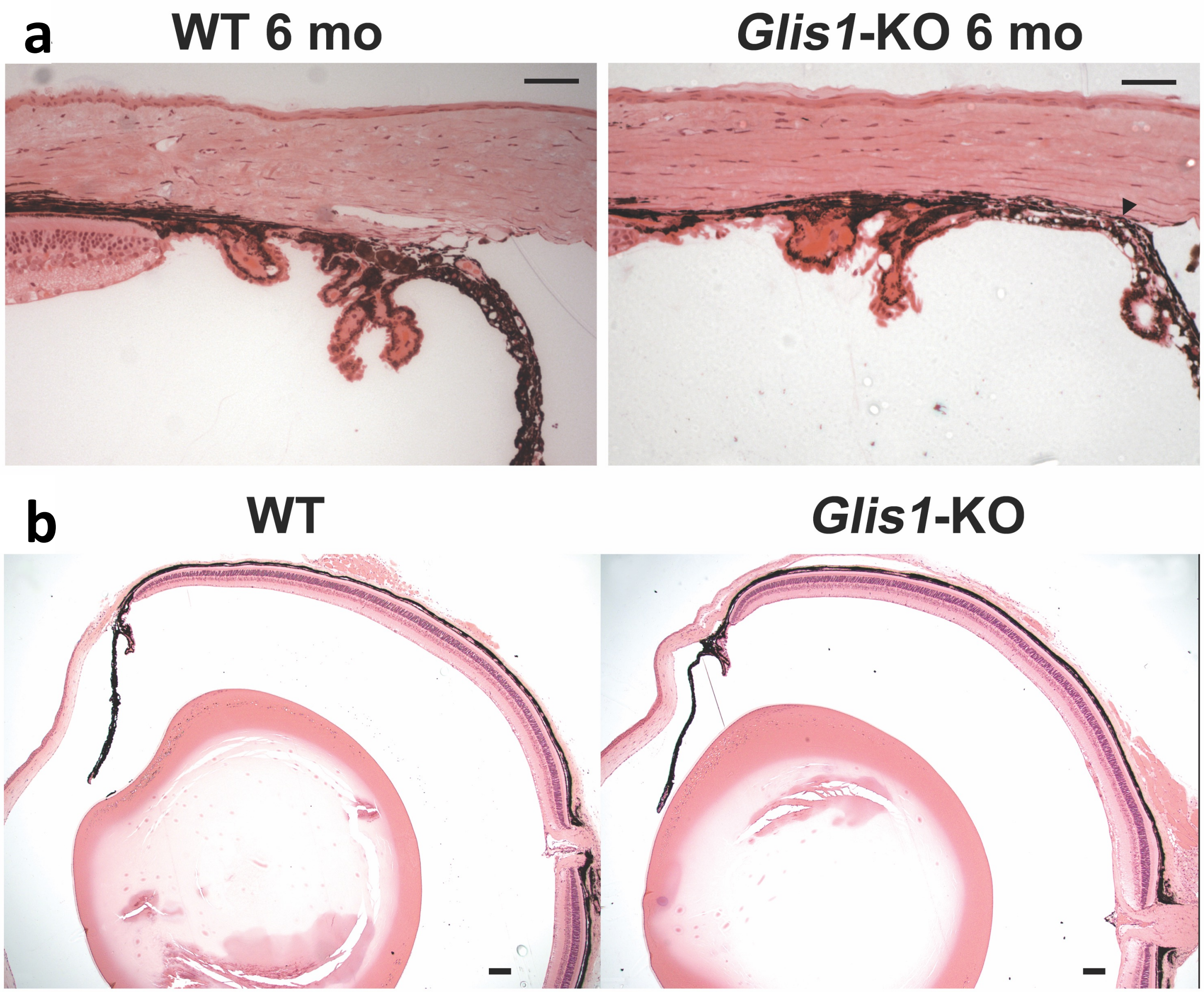

**Supplementary Figure 8. a. Ocular angle drainage structures of *Glis1*-KO mice at older ages.** The WT control mice exhibit prominent SC and TM. In contrast, *Glis1*-KO mice older than 6 months typically show no visible ocular drainage structures and exhibit synechiae characterized by fusion of the iris and cornea resulting in angle closure. Histological assessment of 6 eyes per group was performed with similar results. Scale bar = 50 µm. **b. Histological analysis of whole WT and *Glis1*-KO eyes.** Representative H&E stained ocular sections from WT and *Glis1*-KO eyes showing that *Glis1*-KO eyes do not exhibit gross morphological abnormalities besides ocular drainage tissue defect compared to the control eyes. Ten eyes per experimental group were assessed with similar results. Scale bar = 100 µm.

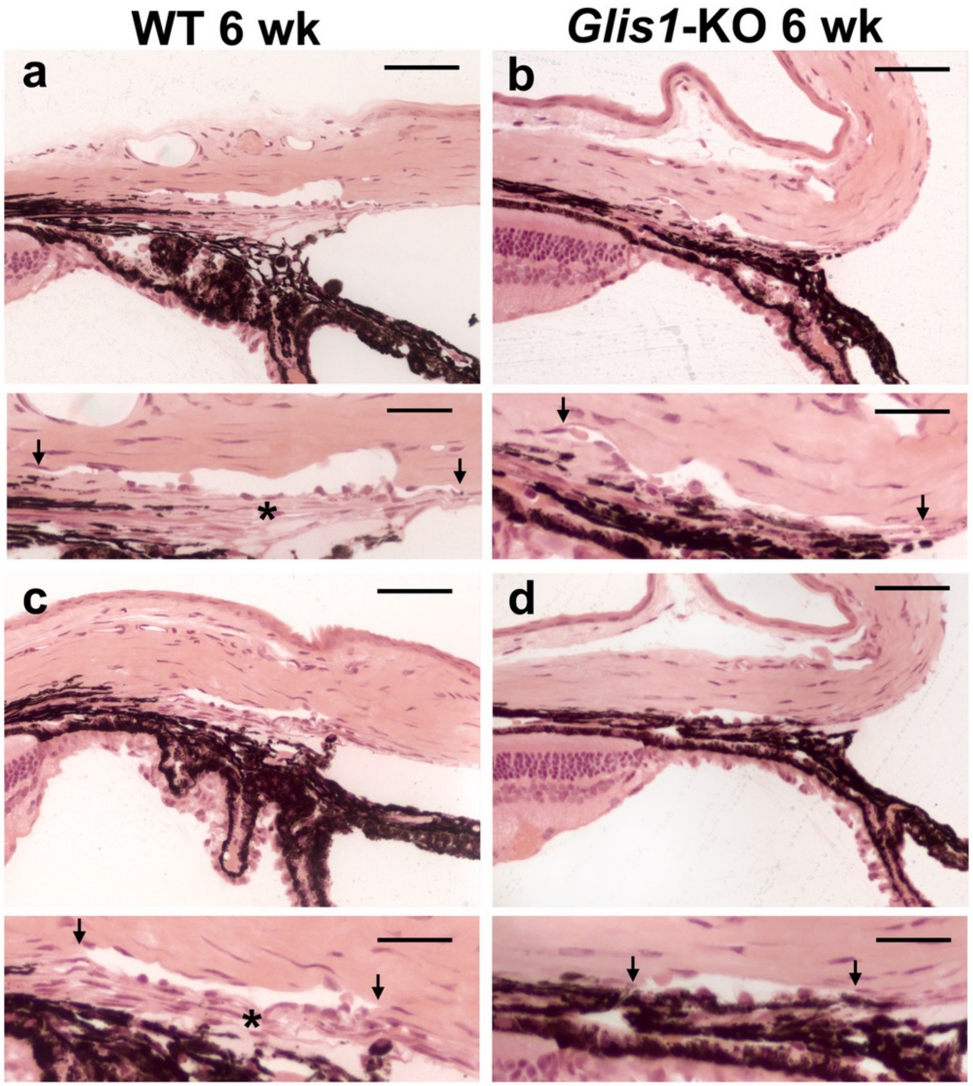

**Supplementary Figure 9. Disruption of the ocular angle drainage structures in *Glis1*-KO mice in 129S6/SvEvTac background. a-d.** Representative histological images of 2 months old WT mice *Glis1*-KO mice maintained in 129S6/SvEvTac genetic background. **a, c.** WT eyes showed a well-developed SC and TM (*). **b, d.** *Glis1*-KO eyes exhibit hypoplastic TM characterized by substantial thinning of the TM. A magnified version of the image in the upper panel is shown in the lower panel. Arrows show edges of the SC. Eight eyes in each of the experimental groups were histologically assessed with similar results. Scale bar = 50 μm for images in the upper panel and 25 μm for images in the lower panels.

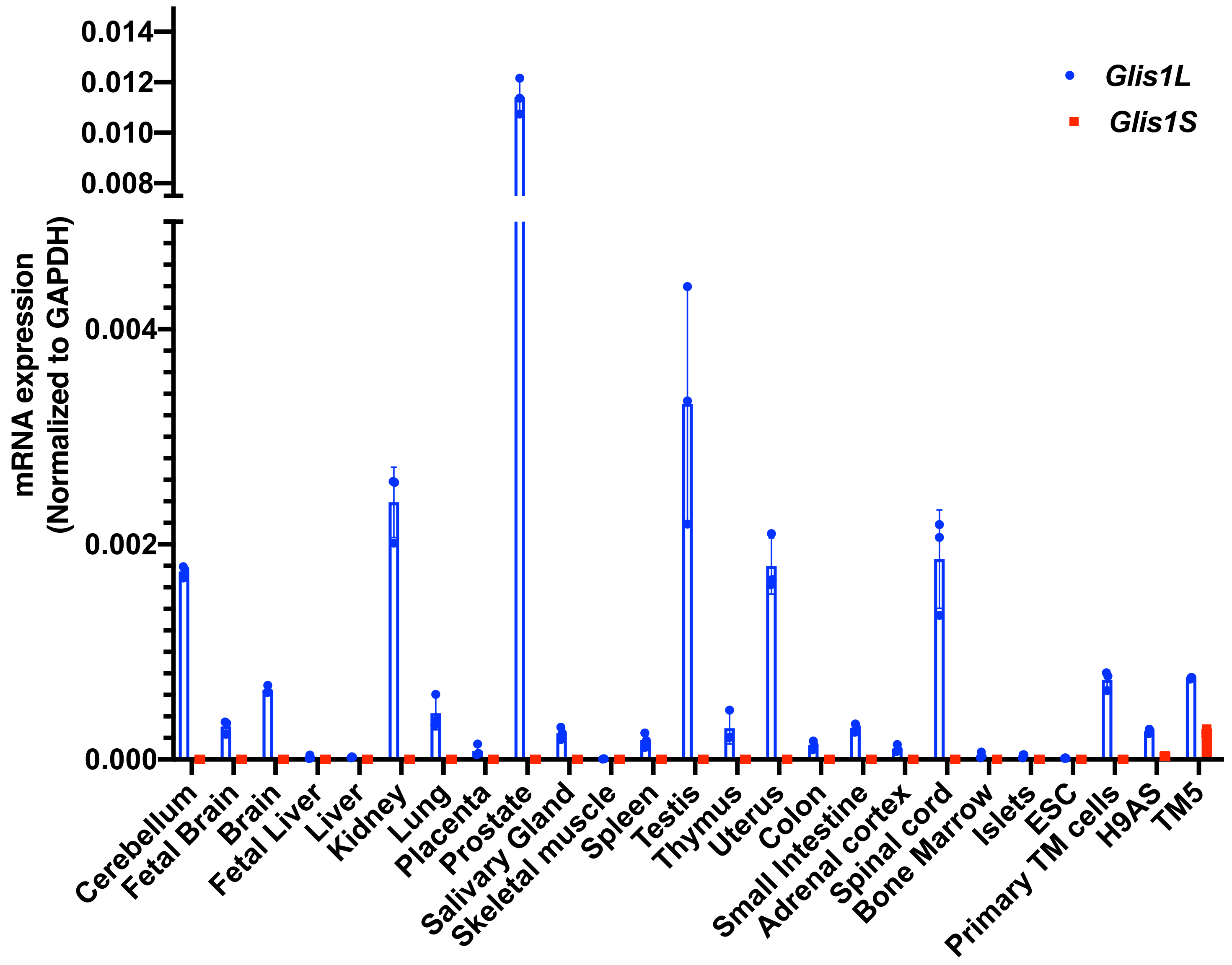

**Supplementary Figure 10. Abundancy of the long and short GLIS1 transcripts ($GLIS1_L$ and $GLIS1_S$) in several human tissues and primary human TM cells**. mRNA expression (ΔΔCt method) were analyzed by QPCR (n=3, PCR replicates), normalized to housekeeping gene (GAPDH). ESC, human embryonic stem cells. Data are represented as means ± SD.

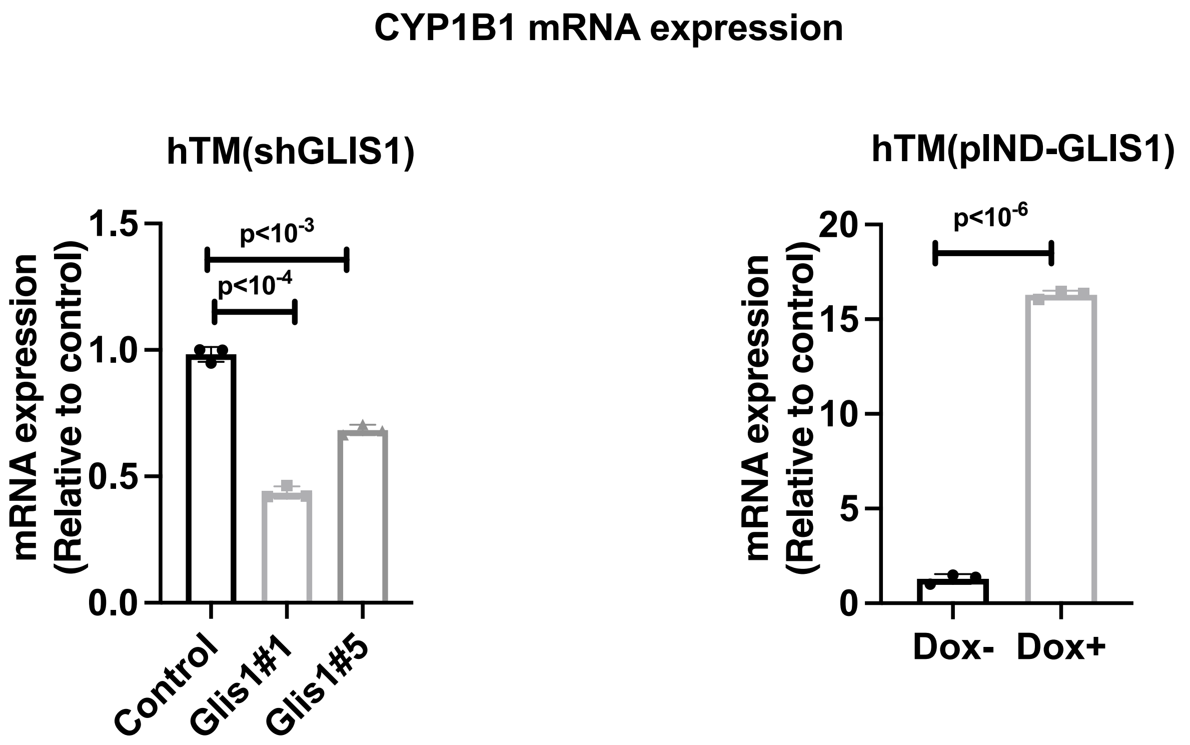

**Supplementary Figure 11. GLIS1 regulates *CYP1B1* expression in primary HTM cells.** QPCR analysis of CYP1B1 mRNA expression in HTM(shGLIS1) and HTM(Scr) (Control) cells (left panel) and in HTM(pIND-GLIS1) cells (right panel). Statistical analyses were performed with two-tailed Student's t-test. Data are represented as means ± SD (n=3 distinct replicates). P-values are indicated above bars.

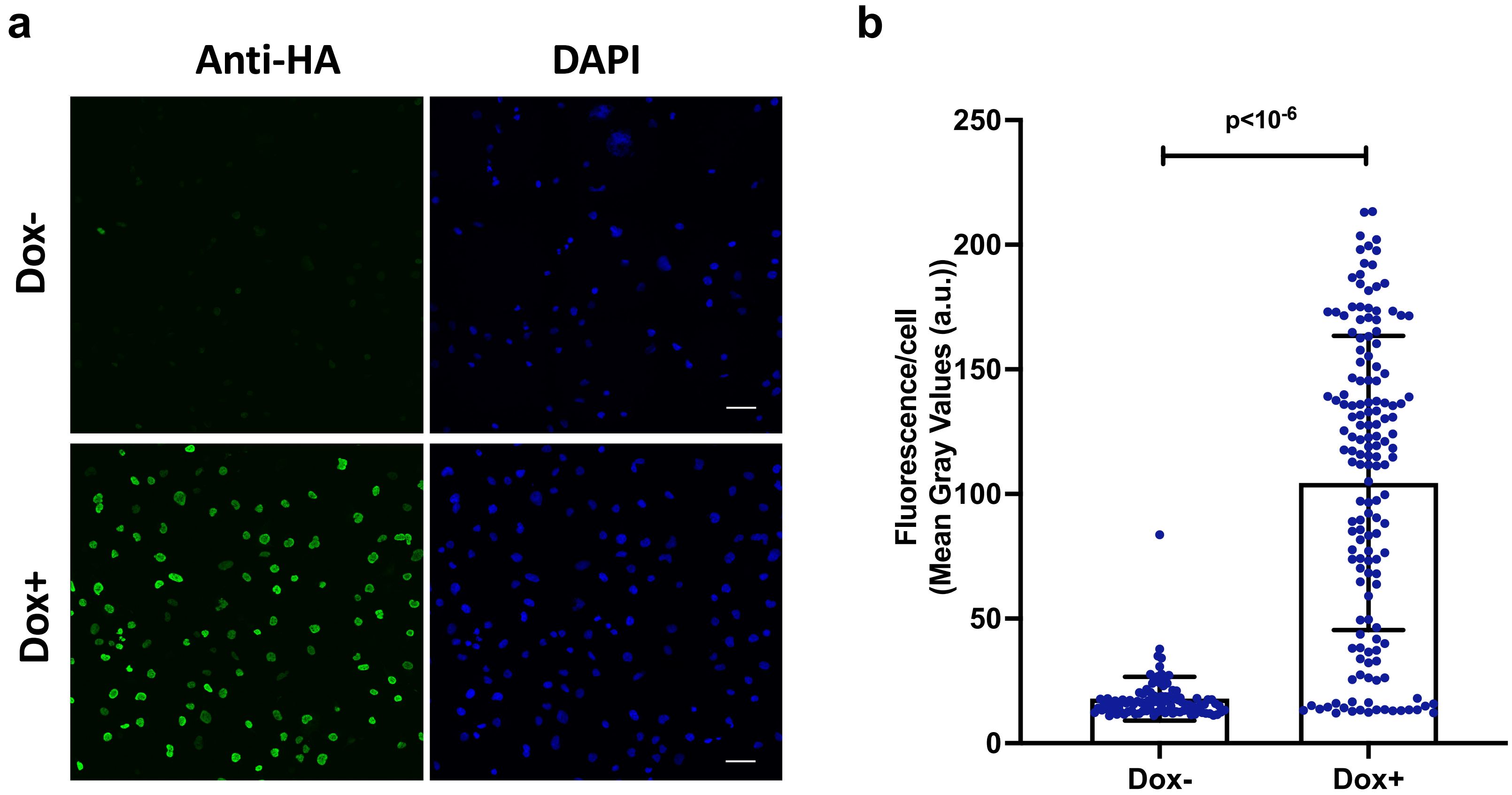

**Supplementary Figure 12. Subcellular localization of GLIS1 protein in TM5 expressing Dox-inducible Flag-GLIS1-HA. a, b.** Cells were treated for 18 h with and without Dox and then GLIS1-HA protein was visualized by immunofluorescence staining in a confocal microscope (Scale bar = 50 μm). Nuclei were stained with DAPI. The level GLIS1-HA fluorescent signal/cell (-Dox, n=97 cells; +Dox, n=152 cells; each one dish) relative to background between cells treated with or without Dox was determined as described[92]. The significantly higher GLIS1-HA fluorescent signal in Dox-treated cells is consistent with the induction of Flag-GLIS1-HA protein. Statistical analyses were performed two-tailed Student's t-test. Data are represented as means ± SD. P-value is indicated above scatter plots.

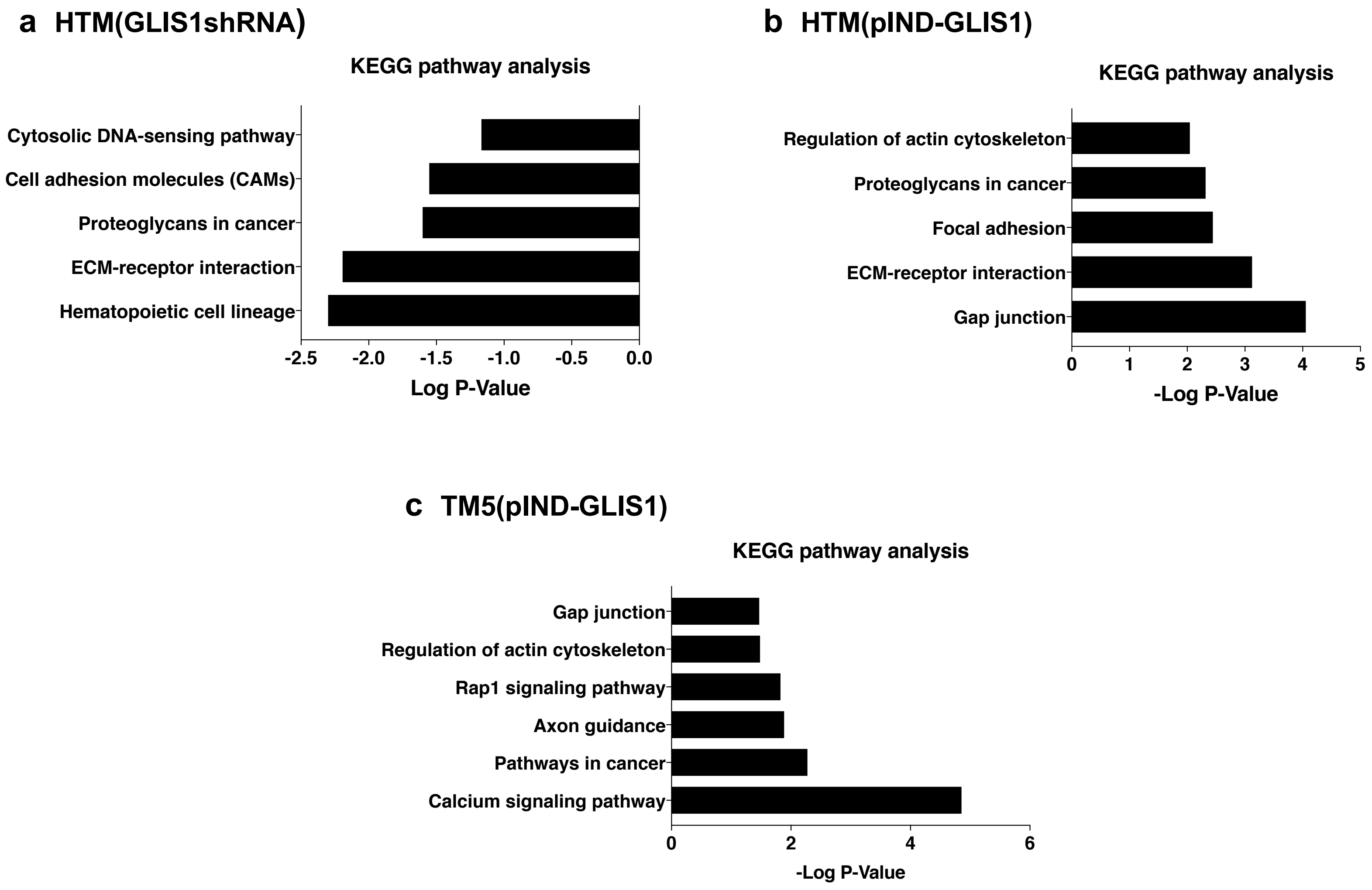

**Supplementary Figure 13. KEGG pathway analysis of GLIS1 target genes down-regulated in HTM(GLIS1shRNA) cells (a) and genes up-regulated in HTM(pIND-GLIS1) (b) and TM5(pIND-GLIS1) (c) cells.** Top pathways are indicated.

.

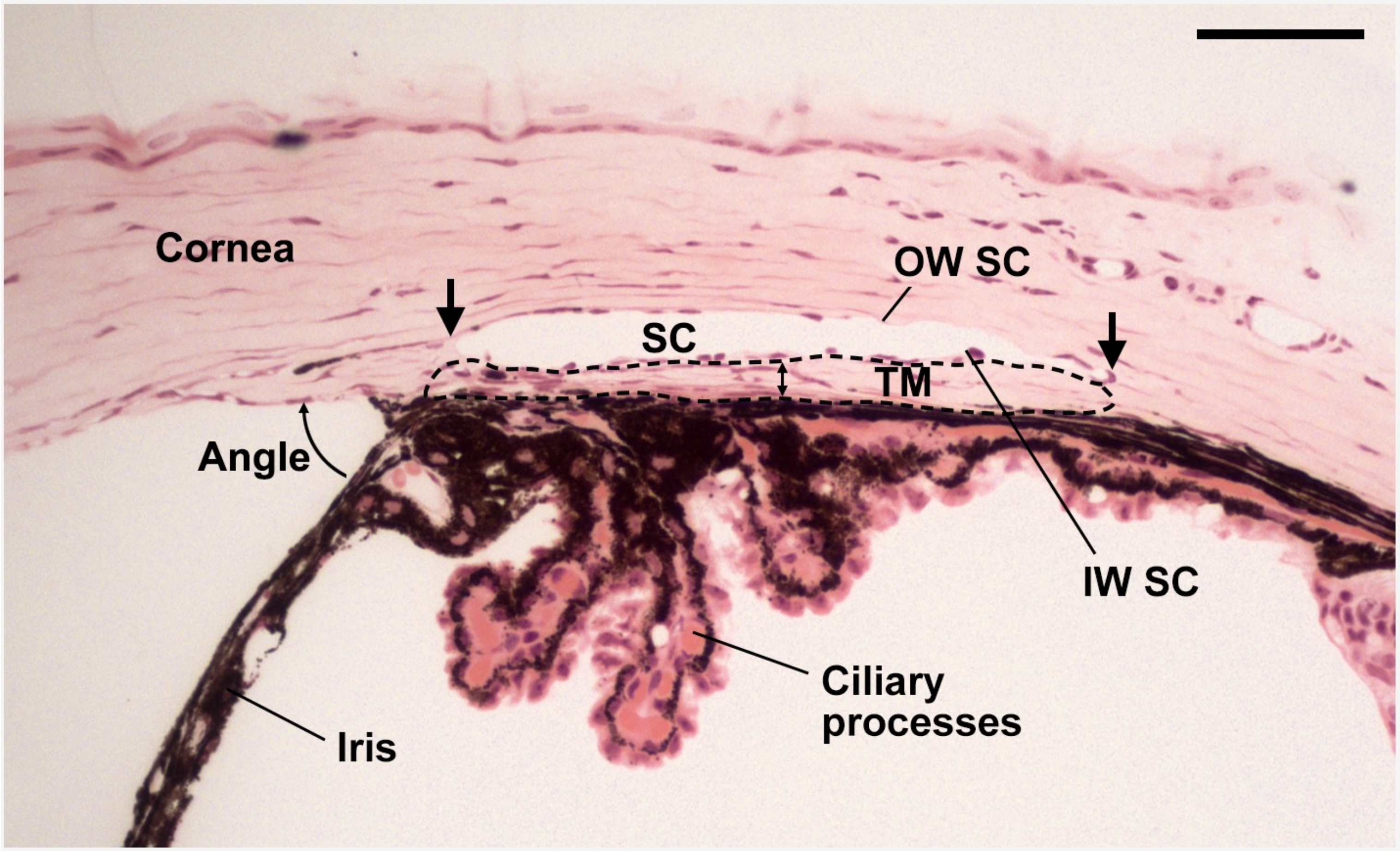

**Supplementary Figure 14. Prominent angle relevant tissues are shown in a histological section of the eye.** The ocular angle relevant structures, TM, SC, Iris, ciliary processes, inner and outer wall of SC (IW SC, OW SC) are shown to aid the reader. Region selected for measuring the TM area is marked by dotted outline. Scale bar = 50 µm.

| Gene | Primer Sequence (5'=>3') |
|---|---|
| Human | |
| MYOC | F: CAAGTCAGTTCTGGAGGAAGAG |
| MYOC | R: CTTCTCAGCCTTGCTACCTC |
| GLIS1 | F: GCCTCTGCCCAGCCCACAAG |
| GLIS1 | R: CCCACCGGCTGCTGGTTCAG |
| GLIS1 S | F: GTACCACCTGGTTGAATTTCTTCA |
| GLIS1 S | R: GTGTCCGTAGCTCCCATTCA |
| GLIS1 L | F: CAGTCCCCGAGACTACGGTC |
| GLIS1 L | R: GTCCAGCAGGCTCTTCTCTGA |
| Col8A1 | F: AAGTACATCCAGCCCATGC |
| Col8A1 | R: CCATGGAGTCCTGGCTTTC |
| LOXL4 | F: CGTGGCTACCTTTCTGAAACT |
| LOXL4 | R: TATGCTGCTTGGCACTGG |
| LTBP2 | F: GACAACAGCAAACAGCACCAA |
| LTBP2 | R: AGCGGCAGACACAGAGC |
| APOD | F: AGAGGGACAAGCATTTCATCTT |
| APOD | R: TCTCAAAGGTTGTTGGGATCTT |
| BMP2 | F: CTGTATCGCAGGCACTCAG |
| BMP2 | R: CCACTCGTTTCTGGTAGTTCTT |
| CYP1B1 | F: GGCCACTATCACTGACATCTT |
| CYP1B1 | R: CCACGACCTGATCCAATTCT |
| GAPDH | F: CCCATCACCATCTTCCAGGAG |
| GAPDH | R: CTTCTCCATGGTGGTGAAGACG |
| Mouse | |
| Glis1 | F: TTCAGCAACTCCAGCGACC |
| Glis1 | R: TTGGAGCAGCCAGGGATCTGA |
| Gapdh | F: CACATCCCAAAGCCCTCG |
| Gapdh | R:CTCAGTCCCCTCCTCAGC |
| Dio1 | F: ATCCAGTACTTCTGGTTTGTCC |
| Dio1 | R: CCTGCTGCCTTGAATGAAATC |
| Dmrtb1 | F: GCTTTGCTACCCGGATCA |
| Dmrtb1 | R: TCAGGATTCCCAGCTCCTA |
| Slc1a7 | F: CTGCCTCCTTGGCTTCTTC |
| Slc1a7 | R: CATCTTCAGCATCCGCATCA |
| Cpt2 | F: AGTATCTGCAGCACAGCATC |
| Cpt2 | R: CTGTGCACTGAGGTATCTCTTC |
| Glis2 | F: GCCTGAACAGGATGCTCGAT |
| Glis2 | : CTTCTCACCTGTGTGGGAGC |
| Glis3 | F: AAGTGGAACAGCTGCTGGG |
| Glis3 | R: GAGTGGAGGTAACTGGGAGGA |

**Supplementary Table 1. List of human and mouse primers used in QPCR analysis.**

| Gene | GLIS1 down-regulation in HTM shGLIS1#1 | GLIS1 down-regulation in HTM shGLIS1#5 | GLIS1 Overexpression in HTM | GLIS1 Overexpression in TM5 | GLIS1 Binding Peak |
|---|---|---|---|---|---|
| ADAMTS10 | down | down | up | down | + |
| ALDH3A1 | down | down | up | up | + |
| APOD | down | down | NC | NC | + |
| ATXN1L | NC | down | down | up | No |
| BGN | down | down | up | up | No |
| BMP2 | down | down | NC | up | + |
| CACNA1A | NC | down | NA | up | + |
| CARD10 | down | down | up | down | + |
| CDKN1A | down | down | NC | up | + |
| CHAC1 | down | down | NC | up | + |
| CHI3L1 | down | down | down | NC | + |
| COL16A1 | down | down | up | up | + |
| COL17A1 | down | down | NC | up | + |
| COL1A2 | NC | down | up | down | No |
| COL4A1 | NC | down | up | NC | + |
| COL4A2 | down | down | up | NC | + |
| COL6A2 | down | down | down | up | + |
| COL7A1 | down | down | NC | up | No |
| COL8A2 | down | down | up | up | No |
| CTSH | NC | down | NC | down | + |
| CYP1B1 | down | down | up | down | + |
| EFEMP1 | up | up | up | NC | No |
| EML2 | down | down | NC | up | + |
| FBLN1 | down | down | NC | NC | + |
| FBLN5 | down | down | up | up | + |
| FBN2 | down | down | NC | up | + |
| FLRT2 | down | down | NC | up | + |
| GLIS1 | down | down | up | up | + |
| HMGA1 | down | down | down | up | + |
| IL20RB | NC | NC | NC | NC | + |
| ITGA3 | NC | down | up | up | + |
| LOXL1 | down | down | up | NC | + |
| LOXL2 | NC | down | up | down | + |
| LOXL3 | down | down | up | up | + |
| LOXL4 | down | down | up | up | + |
| LTBP2 | down | down | up | up | + |
| MMP2 | down | down | up | up | + |
| MYLK | down | down | up | up | + |
| MYOC | down | down | NC | up | + |
| PAPLN | NC | down | up | up | + |
| PODN | NC | down | NC | up | + |
| PTGES | down | down | up | NC | + |
| RTN4 | up | up | NC | down | + |
| SAA2 | NA | NA | NC | up | + |
| SDC1 | down | down | up | down | + |
| SDC2 | NC | NC | up | up | + |
| SEMA4B | NC | down | NC | down | + |
| SPARC | NC | down | up | down | + |
| TFAP2B | NC | down | NC | NA | No |
| TGFBR3 | down | NC | up | up | + |
| TIMP3 | NC | down | up | up | + |

**Supplementary Table 2. Association of GLIS1 peaks (ChIP-Seq) with several GLIS1-regulated genes in GLIS1 down-regulated HTM(shGLIS1) and GLIS1-overexpressing HTM and TM5 cells (HTM(pIND-GLIS1) and TM5(pIND-GLIS1).**

NC, no statically significant change.

**Supplementary Table 3. Association of *GLIS1* SNPs with POAG in the multiethnic meta-analysis GERA+UKB.**

| CHR | BP | SNP | A1 | A2 | N | P | P.R. | OR | OR.R. | Q | I |
|---|---|---|---|---|---|---|---|---|---|---|---|
| 1 | 54123873 | rs941125 | G | T | 2 | 4.73E-06 | 4.73E-06 | 0.939 | 0.939 | 0.6693 | 0 |
| 1 | 54126754 | rs1202629 | T | A | 2 | 6.24E-06 | 6.24E-06 | 0.9305 | 0.9305 | 0.5387 | 0 |
| 1 | 54125615 | rs2874300 | G | A | 2 | 9.68E-06 | 9.68E-06 | 0.9317 | 0.9317 | 0.5574 | 0 |
| 1 | 54131753 | rs4927025 | C | T | 2 | 1.02E-05 | 1.02E-05 | 0.9326 | 0.9326 | 0.4723 | 0 |
| 1 | 54135078 | rs1465037 | C | A | 2 | 1.11E-05 | 1.11E-05 | 0.933 | 0.933 | 0.4746 | 0 |
| 1 | 54135334 | rs1554750 | T | C | 2 | 1.14E-05 | 1.14E-05 | 0.9331 | 0.9331 | 0.4745 | 0 |
| 1 | 54121976 | rs1256852 | G | A | 2 | 1.16E-05 | 1.16E-05 | 0.9414 | 0.9414 | 0.801 | 0 |
| 1 | 54119913 | rs1465035 | C | T | 2 | 1.30E-05 | 1.30E-05 | 1.0599 | 1.0599 | 0.9838 | 0 |
| 1 | 54133723 | rs6731917 | T | A | 2 | 1.32E-05 | 1.32E-05 | 0.9337 | 0.9337 | 0.5021 | 0 |
| 1 | 54120058 | rs1474462 | G | A | 2 | 1.55E-05 | 1.55E-05 | 1.0598 | 1.0598 | 0.9669 | 0 |
| 1 | 54120307 | rs1273375 | G | A | 2 | 1.58E-05 | 1.58E-05 | 1.0597 | 1.0597 | 0.9692 | 0 |
| 1 | 54133345 | rs6658469 | C | A | 2 | 1.65E-05 | 1.65E-05 | 0.9347 | 0.9347 | 0.6151 | 0 |
| 1 | 54121981 | rs1256192 | T | G | 2 | 1.85E-05 | 1.85E-05 | 0.9427 | 0.9427 | 0.786 | 0 |
| 1 | 54122770 | rs6688040 | C | T | 2 | 1.88E-05 | 1.88E-05 | 1.0592 | 1.0592 | 0.9666 | 0 |
| 1 | 54120873 | rs2141079 | C | T | 2 | 2.04E-05 | 2.04E-05 | 1.0589 | 1.0589 | 0.9742 | 0 |
| 1 | 54122030 | rs1274425 | G | A | 2 | 2.04E-05 | 2.04E-05 | 1.0589 | 1.0589 | 0.9605 | 0 |
| 1 | 54150890 | rs7518284 | A | T | 2 | 2.08E-05 | 2.08E-05 | 0.935 | 0.935 | 0.375 | 0 |
| 1 | 54121458 | rs4927023 | C | T | 2 | 2.21E-05 | 2.21E-05 | 1.0586 | 1.0586 | 0.9884 | 0 |
| 1 | 54117985 | rs1120618 | G | T | 2 | 2.39E-05 | 2.39E-05 | 1.0584 | 1.0584 | 0.8803 | 0 |
| 1 | 54145288 | rs7549345 | T | C | 2 | 2.73E-05 | 2.73E-05 | 0.9359 | 0.9359 | 0.4132 | 0 |
| 1 | 54129732 | rs1240583 | C | G | 2 | 2.78E-05 | 2.78E-05 | 0.9356 | 0.9356 | 0.5857 | 0 |
| 1 | 54156624 | rs1208235 | A | C | 2 | 2.79E-05 | 2.79E-05 | 0.9361 | 0.9361 | 0.3663 | 0 |
| 1 | 54127969 | rs1120618 | G | A | 2 | 3.69E-05 | 3.69E-05 | 0.9361 | 0.9361 | 0.5529 | 0 |
| 1 | 54158475 | rs7544346 | C | T | 2 | 4.52E-05 | 4.52E-05 | 0.9377 | 0.9377 | 0.3449 | 0 |
| 1 | 54169455 | rs4927029 | C | G | 2 | 4.97E-05 | 0.000215 | 0.9382 | 0.9387 | 0.2809 | 14 |
| 1 | 54163071 | rs1120619 | G | A | 2 | 5.11E-05 | 5.11E-05 | 1.0573 | 1.0573 | 0.521 | 0 |
| 1 | 54158155 | rs7555617 | A | C | 2 | 5.31E-05 | 5.31E-05 | 0.9383 | 0.9383 | 0.3254 | 0 |
| 1 | 54157640 | rs1208645 | C | T | 2 | 5.45E-05 | 0.000804 | 0.9386 | 0.9397 | 0.2437 | 26.43 |
| 1 | 54165624 | rs1120619 | A | G | 2 | 5.62E-05 | 5.62E-05 | 0.939 | 0.939 | 0.389 | 0 |
| 1 | 54163251 | rs7519965 | G | A | 2 | 6.03E-05 | 6.86E-05 | 0.9388 | 0.9389 | 0.3143 | 1.23 |
| 1 | 54160570 | rs1088880 | C | T | 2 | 6.21E-05 | 6.21E-05 | 0.937 | 0.937 | 0.5276 | 0 |
| 1 | 54157742 | rs1207619 | A | G | 2 | 6.25E-05 | 0.000191 | 0.9392 | 0.9396 | 0.2895 | 10.88 |
| 1 | 54160558 | rs1204433 | C | T | 2 | 6.32E-05 | 6.32E-05 | 0.9372 | 0.9372 | 0.5245 | 0 |
| 1 | 54158762 | rs1120619 | C | G | 2 | 7.16E-05 | 7.16E-05 | 0.9397 | 0.9397 | 0.3905 | 0 |
| 1 | 54161087 | rs6665257 | T | C | 2 | 7.26E-05 | 7.26E-05 | 0.9398 | 0.9398 | 0.3891 | 0 |
| 1 | 54178501 | rs1208099 | G | T | 2 | 7.29E-05 | 7.29E-05 | 0.94 | 0.94 | 0.4288 | 0 |
| 1 | 54175548 | rs1710920 | A | G | 2 | 7.30E-05 | 7.30E-05 | 0.9404 | 0.9404 | 0.5666 | 0 |
| 1 | 54159198 | rs6694399 | C | T | 2 | 7.53E-05 | 8.11E-05 | 0.9397 | 0.9398 | 0.3156 | 0.72 |
| 1 | 54178270 | rs1208090 | G | A | 2 | 7.56E-05 | 7.56E-05 | 0.9401 | 0.9401 | 0.4235 | 0 |
| 1 | 54160571 | rs1088880 | A | G | 2 | 7.95E-05 | 7.95E-05 | 0.9379 | 0.9379 | 0.4695 | 0 |
| 1 | 54158462 | rs7555908 | A | G | 2 | 8.51E-05 | 0.000177 | 0.9402 | 0.9404 | 0.299 | 7.28 |
| 1 | 54159005 | rs6658245 | T | C | 2 | 8.53E-05 | 0.000271 | 0.9402 | 0.9406 | 0.2876 | 11.59 |
| 1 | 54158690 | rs1120619 | G | A | 2 | 8.62E-05 | 0.00017 | 0.9402 | 0.9405 | 0.3003 | 6.79 |
| 1 | 54160235 | rs1088880 | C | T | 2 | 8.64E-05 | 0.000311 | 0.9402 | 0.9407 | 0.2839 | 12.92 |
| 1 | 54159079 | rs6691833 | C | G | 2 | 8.82E-05 | 0.000251 | 0.9403 | 0.9407 | 0.2904 | 10.52 |

| | | | | | | | | | | | |
|---|---|---|---|---|---|---|---|---|---|---|---|
| 1 | 54182886 | rs4927030 | A | C | 2 | 8.83E-05 | 8.83E-05 | 0.9472 | 0.9472 | 0.6947 | 0 |
| 1 | 54164799 | rs6657830 | A | G | 2 | 9.09E-05 | 0.000605 | 0.9405 | 0.9412 | 0.2651 | 19.46 |
| 1 | 54160812 | rs6673378 | G | T | 2 | 9.27E-05 | 0.000267 | 0.9405 | 0.9409 | 0.2899 | 10.71 |
| 1 | 54160709 | rs6118481 | G | T | 2 | 9.31E-05 | 0.000264 | 0.9405 | 0.9409 | 0.2903 | 10.57 |
| 1 | 54161545 | rs4927028 | T | C | 2 | 9.36E-05 | 0.000285 | 0.9405 | 0.941 | 0.2883 | 11.32 |
| 1 | 54161112 | rs6695509 | A | G | 2 | 9.63E-05 | 0.000244 | 0.9406 | 0.941 | 0.2934 | 9.41 |
| 1 | 54162471 | rs6661728 | G | A | 2 | 0.0001 | 0.000269 | 0.9408 | 0.9412 | 0.2915 | 10.13 |
| 1 | 54186489 | rs4927032 | C | T | 2 | 0.000101 | 0.000101 | 0.941 | 0.941 | 0.3521 | 0 |
| 1 | 54181942 | rs1526911 | C | G | 2 | 0.000102 | 0.000102 | 0.9411 | 0.9411 | 0.4005 | 0 |
| 1 | 54166790 | rs6588487 | G | A | 2 | 0.000102 | 0.000489 | 0.9409 | 0.9415 | 0.2747 | 16.2 |
| 1 | 54165910 | rs1120619 | C | A | 2 | 0.000103 | 0.0008 | 0.9409 | 0.9418 | 0.2591 | 21.48 |
| 1 | 54158102 | rs1120619 | G | A | 2 | 0.000103 | 0.00045 | 0.9409 | 0.9415 | 0.2774 | 15.24 |
| 1 | 54149577 | rs2948044 | G | C | 2 | 0.000103 | 0.000103 | 1.0549 | 1.0549 | 0.8749 | 0 |
| 1 | 54158220 | rs7555711 | A | G | 2 | 0.000105 | 0.000434 | 0.9409 | 0.9415 | 0.279 | 14.68 |
| 1 | 54173551 | rs1710917 | C | G | 2 | 0.000105 | 0.000105 | 0.9412 | 0.9412 | 0.4198 | 0 |
| 1 | 54168609 | rs1204008 | A | G | 2 | 0.000108 | 0.000905 | 0.9411 | 0.942 | 0.2561 | 22.45 |
| 1 | 54170801 | rs1209759 | C | G | 2 | 0.000108 | 0.001033 | 0.9412 | 0.9421 | 0.2515 | 23.94 |
| 1 | 54182502 | rs5608700 | A | C | 2 | 0.000108 | 0.000108 | 0.9413 | 0.9413 | 0.3957 | 0 |
| 1 | 54168545 | rs1120620 | T | C | 2 | 0.00011 | 0.000406 | 0.9412 | 0.9417 | 0.2818 | 13.67 |
| 1 | 54165831 | rs1120619 | A | G | 2 | 0.000112 | 0.000584 | 0.9412 | 0.9419 | 0.2714 | 17.33 |
| 1 | 54164998 | rs6658053 | A | T | 2 | 0.000118 | 0.001298 | 0.9414 | 0.9425 | 0.246 | 25.7 |
| 1 | 54166777 | rs6588486 | G | A | 2 | 0.000124 | 0.001812 | 0.9416 | 0.9428 | 0.2351 | 29.07 |
| 1 | 54170963 | rs1120620 | C | A | 2 | 0.000124 | 0.002145 | 0.9417 | 0.943 | 0.2286 | 31.02 |
| 1 | 54180287 | rs2186041 | G | A | 2 | 0.000126 | 0.000126 | 0.9421 | 0.9421 | 0.3866 | 0 |
| 1 | 54175971 | rs1879733 | T | C | 2 | 0.000128 | 0.000128 | 0.9422 | 0.9422 | 0.4155 | 0 |
| 1 | 54175861 | rs1879731 | G | A | 2 | 0.000128 | 0.000128 | 0.9422 | 0.9422 | 0.4158 | 0 |
| 1 | 54164165 | rs1204122 | A | G | 2 | 0.00013 | 0.002867 | 0.9418 | 0.9433 | 0.2186 | 33.94 |
| 1 | 54175629 | rs1204369 | T | C | 2 | 0.000139 | 0.000139 | 0.9425 | 0.9425 | 0.401 | 0 |
| 1 | 54175909 | rs1879732 | C | T | 2 | 0.000139 | 0.000139 | 0.9425 | 0.9425 | 0.401 | 0 |
| 1 | 54175360 | rs1202585 | C | T | 2 | 0.00014 | 0.00014 | 0.9425 | 0.9425 | 0.4013 | 0 |
| 1 | 54187282 | rs1738434 | T | C | 2 | 0.000142 | 0.000142 | 0.9489 | 0.9489 | 0.9162 | 0 |
| 1 | 54191089 | rs797907 | A | T | 2 | 0.000144 | 0.000144 | 0.9489 | 0.9489 | 0.865 | 0 |
| 1 | 54174607 | rs1120620 | A | G | 2 | 0.000166 | 0.000166 | 0.9431 | 0.9431 | 0.4422 | 0 |
| 1 | 54162635 | rs5902094 | A | G | 2 | 0.000171 | 0.001487 | 0.9428 | 0.9438 | 0.2511 | 24.09 |
| 1 | 54163128 | rs7554633 | A | G | 2 | 0.000172 | 0.001748 | 0.9428 | 0.9439 | 0.2454 | 25.88 |
| 1 | 54163207 | rs7512093 | T | G | 2 | 0.000173 | 0.00185 | 0.9429 | 0.944 | 0.2432 | 26.56 |
| 1 | 54171981 | rs5578613 | T | C | 2 | 0.000174 | 0.006703 | 0.9427 | 0.9447 | 0.1906 | 41.63 |
| 1 | 54152621 | rs3013777 | T | C | 2 | 0.000206 | 0.000206 | 1.0522 | 1.0522 | 0.8548 | 0 |
| 1 | 54182174 | rs1526910 | G | A | 2 | 0.00021 | 0.00021 | 0.944 | 0.944 | 0.4054 | 0 |
| 1 | 54165699 | rs1780395 | G | T | 2 | 0.000226 | 0.000226 | 1.0517 | 1.0517 | 0.7224 | 0 |
| 1 | 54172250 | rs1173588 | C | G | 2 | 0.000228 | 0.000228 | 1.0517 | 1.0517 | 0.5889 | 0 |
| 1 | 54164156 | rs1120619 | T | C | 2 | 0.000239 | 0.008502 | 0.944 | 0.946 | 0.1876 | 42.42 |
| 1 | 54183191 | rs4927031 | C | T | 2 | 0.000242 | 0.000242 | 0.9445 | 0.9445 | 0.3428 | 0 |
| 1 | 54126552 | rs1738332 | C | G | 2 | 0.000247 | 0.00272 | 0.9403 | 0.9418 | 0.2438 | 26.38 |
| 1 | 54184097 | rs1204352 | G | A | 2 | 0.000252 | 0.000252 | 0.9446 | 0.9446 | 0.3366 | 0 |
| 1 | 54172944 | rs1173589 | G | A | 2 | 0.000298 | 0.000298 | 1.0506 | 1.0506 | 0.8873 | 0 |
| 1 | 54190695 | rs797906 | C | A | 2 | 0.0003 | 0.0003 | 0.9514 | 0.9514 | 0.6426 | 0 |
| 1 | 54055693 | rs1879735 | A | G | 2 | 0.000308 | 0.000308 | 0.9508 | 0.9508 | 0.7569 | 0 |

| | | | | | | | | | | | |
|---|---|---|---|---|---|---|---|---|---|---|---|
| 1 | 54563039 | rs1181088 | G | A | 2 | 0.000342 | 0.000342 | 0.8167 | 0.8167 | 0.6779 | 0 |
| 1 | 54051898 | rs2948045 | T | C | 2 | 0.00037 | 0.00037 | 0.9514 | 0.9514 | 0.6306 | 0 |
| 1 | 54177279 | rs7539908 | T | C | 2 | 0.000376 | 0.008134 | 0.9429 | 0.9451 | 0.2088 | 36.71 |
| 1 | 54190333 | rs1205971 | A | C | 2 | 0.000379 | 0.000379 | 0.9464 | 0.9464 | 0.3423 | 0 |
| 1 | 54138854 | rs1879734 | T | C | 2 | 0.000396 | 0.000396 | 0.9481 | 0.9481 | 0.4724 | 0 |
| 1 | 54173101 | rs1173590 | G | T | 2 | 0.000439 | 0.000439 | 1.0492 | 1.0492 | 0.8901 | 0 |
| 1 | 54157498 | rs3013766 | G | A | 2 | 0.000443 | 0.000443 | 1.0534 | 1.0534 | 0.7378 | 0 |
| 1 | 54159441 | rs5636885 | C | T | 2 | 0.00045 | 0.04137 | 0.9432 | 0.9476 | 0.1273 | 57 |
| 1 | 54193023 | rs7554636 | T | C | 2 | 0.000492 | 0.000492 | 0.9472 | 0.9472 | 0.3333 | 0 |
| 1 | 54130119 | rs3006905 | G | C | 2 | 0.000512 | 0.000512 | 1.0535 | 1.0535 | 0.838 | 0 |
| 1 | 54183531 | rs7266067 | C | G | 2 | 0.000519 | 0.03806 | 0.9439 | 0.948 | 0.1353 | 55.16 |
| 1 | 54156434 | rs3013768 | G | A | 2 | 0.000531 | 0.000531 | 1.0528 | 1.0528 | 0.6014 | 0 |
| 1 | 54189197 | rs5632615 | G | A | 2 | 0.000573 | 0.000573 | 0.9482 | 0.9482 | 0.3499 | 0 |
| 1 | 54156443 | rs2948043 | A | C | 2 | 0.00058 | 0.00058 | 1.0525 | 1.0525 | 0.6144 | 0 |
| 1 | 54191103 | rs1209193 | G | A | 2 | 0.000594 | 0.000594 | 0.9483 | 0.9483 | 0.3344 | 0 |
| 1 | 54156757 | rs3013767 | C | T | 2 | 0.000595 | 0.000595 | 1.0523 | 1.0523 | 0.6041 | 0 |
| 1 | 54155902 | rs3013771 | C | T | 2 | 0.000597 | 0.000597 | 1.0523 | 1.0523 | 0.619 | 0 |
| 1 | 54138489 | rs2141080 | C | T | 2 | 0.000617 | 0.000617 | 0.9498 | 0.9498 | 0.5862 | 0 |
| 1 | 54154726 | rs882879 | G | T | 2 | 0.000633 | 0.000633 | 1.0521 | 1.0521 | 0.6277 | 0 |
| 1 | 54138698 | rs2141082 | C | T | 2 | 0.000638 | 0.000638 | 0.95 | 0.95 | 0.5911 | 0 |
| 1 | 54155959 | rs3013770 | C | T | 2 | 0.000648 | 0.000648 | 1.0521 | 1.0521 | 0.5948 | 0 |
| 1 | 54138552 | rs2141081 | C | T | 2 | 0.000649 | 0.000649 | 0.95 | 0.95 | 0.5933 | 0 |
| 1 | 54121818 | rs1203179 | C | T | 2 | 0.000651 | 0.000651 | 0.9495 | 0.9495 | 0.9784 | 0 |
| 1 | 54188994 | rs6658137 | T | C | 2 | 0.000674 | 0.000691 | 0.9489 | 0.9489 | 0.3165 | 0.32 |
| 1 | 54152703 | rs2948042 | G | A | 2 | 0.000692 | 0.000692 | 1.0517 | 1.0517 | 0.7098 | 0 |
| 1 | 54000726 | rs7520584 | A | G | 2 | 0.000695 | 0.000695 | 1.0659 | 1.0659 | 0.5177 | 0 |
| 1 | 54129591 | rs2950241 | G | A | 2 | 0.000724 | 0.000724 | 1.052 | 1.052 | 0.8595 | 0 |
| 1 | 54188096 | rs1120621 | T | C | 2 | 0.000748 | 0.001029 | 0.9493 | 0.9494 | 0.3069 | 4.2 |
| 1 | 54188577 | rs5571812 | G | A | 2 | 0.0008 | 0.001146 | 0.9492 | 0.9493 | 0.3058 | 4.66 |
| 1 | 54140843 | rs4927026 | A | G | 2 | 0.000813 | 0.01433 | 0.9458 | 0.9482 | 0.2062 | 37.41 |
| 1 | 54188674 | rs5587328 | T | C | 2 | 0.000824 | 0.001071 | 0.9494 | 0.9495 | 0.3089 | 3.41 |
| 1 | 54195018 | rs702490 | A | G | 2 | 0.000833 | 0.000833 | 1.0484 | 1.0484 | 0.4499 | 0 |
| 1 | 54156323 | rs3013769 | G | A | 2 | 0.00086 | 0.00086 | 1.0508 | 1.0508 | 0.5926 | 0 |
| 1 | 54165541 | rs1615037 | C | T | 2 | 0.00088 | 0.00088 | 1.0507 | 1.0507 | 0.6888 | 0 |
| 1 | 54143759 | rs2950244 | C | T | 2 | 0.00097 | 0.00097 | 1.0507 | 1.0507 | 0.9297 | 0 |
| 1 | 54051924 | rs4926604 | T | C | 2 | 0.001006 | 0.001006 | 1.047 | 1.047 | 0.9899 | 0 |
| 1 | 54152487 | rs3006882 | G | A | 2 | 0.001063 | 0.001063 | 1.0499 | 1.0499 | 0.6138 | 0 |
| 1 | 54038523 | rs1120617 | G | C | 2 | 0.001105 | 0.06205 | 1.0617 | 1.0653 | 0.0672 | 70.14 |
| 1 | 54175302 | rs1780392 | C | T | 2 | 0.001156 | 0.001156 | 1.0491 | 1.0491 | 0.8531 | 0 |
| 1 | 53981255 | rs499749 | G | T | 2 | 0.001161 | 0.01055 | 0.8181 | 0.81 | 0.1957 | 40.27 |
| 1 | 54177708 | rs1780391 | C | G | 2 | 0.001233 | 0.001233 | 1.0488 | 1.0488 | 0.8053 | 0 |
| 1 | 54173693 | rs1173591 | C | T | 2 | 0.001251 | 0.001251 | 1.0487 | 1.0487 | 0.8407 | 0 |
| 1 | 54123513 | rs1111658 | A | G | 2 | 0.001291 | 0.001291 | 0.9524 | 0.9524 | 0.9078 | 0 |
| 1 | 54058231 | rs4927011 | C | T | 2 | 0.001295 | 0.001295 | 1.046 | 1.046 | 0.8627 | 0 |
| 1 | 54068016 | rs4927012 | C | T | 2 | 0.001361 | 0.03366 | 1.0626 | 1.0653 | 0.1195 | 58.74 |
| 1 | 54181668 | rs1614395 | G | A | 2 | 0.00153 | 0.00153 | 1.0479 | 1.0479 | 0.4847 | 0 |
| 1 | 54043110 | rs4634849 | C | T | 2 | 0.001571 | 0.001571 | 1.0453 | 1.0453 | 0.8695 | 0 |
| 1 | 54174846 | rs2694594 | G | C | 2 | 0.001574 | 0.001574 | 1.0477 | 1.0477 | 0.7655 | 0 |

| | | | | | | | | | | | |
|---|---|---|---|---|---|---|---|---|---|---|---|
| 1 | 53990321 | rs541224 | C | T | 2 | 0.001611 | 0.001611 | 1.0657 | 1.0657 | 0.5082 | 0 |
| 1 | 53992270 | rs3855965 | C | T | 2 | 0.001633 | 0.001633 | 1.0596 | 1.0596 | 0.5207 | 0 |
| 1 | 54061333 | rs7551844 | T | C | 2 | 0.001663 | 0.001663 | 1.0451 | 1.0451 | 0.7888 | 0 |
| 1 | 54191811 | rs7266068 | T | C | 2 | 0.001782 | 0.04075 | 0.95 | 0.9531 | 0.1663 | 47.82 |
| 1 | 54046217 | rs1213729 | G | T | 2 | 0.0018 | 0.0018 | 1.0446 | 1.0446 | 0.8515 | 0 |
| 1 | 54062273 | rs2948053 | A | G | 2 | 0.001821 | 0.001821 | 0.9574 | 0.9574 | 0.7146 | 0 |
| 1 | 54197141 | rs797909 | T | C | 2 | 0.001832 | 0.009163 | 1.0454 | 1.0442 | 0.2525 | 23.64 |
| 1 | 54196929 | rs808860 | C | T | 2 | 0.001888 | 0.007025 | 1.0447 | 1.0438 | 0.2661 | 19.14 |
| 1 | 54047882 | rs1214036 | A | T | 2 | 0.0019 | 0.0019 | 1.0444 | 1.0444 | 0.8532 | 0 |
| 1 | 54187575 | rs1738850 | C | G | 2 | 0.002042 | 0.1073 | 0.9502 | 0.9553 | 0.0989 | 63.29 |
| 1 | 54061978 | rs3013754 | A | C | 2 | 0.002088 | 0.002088 | 0.9579 | 0.9579 | 0.7276 | 0 |
| 1 | 54141274 | rs4927027 | A | G | 2 | 0.002296 | 0.002296 | 0.9553 | 0.9553 | 0.6563 | 0 |
| 1 | 54143423 | rs7543166 | T | C | 2 | 0.002382 | 0.002382 | 0.9554 | 0.9554 | 0.6511 | 0 |
| 1 | 54193701 | rs910298 | T | C | 2 | 0.002456 | 0.09911 | 0.9512 | 0.9559 | 0.1109 | 60.66 |
| 1 | 54144297 | rs5588854 | T | C | 2 | 0.002578 | 0.002578 | 0.9558 | 0.9558 | 0.6224 | 0 |
| 1 | 54063938 | rs1120617 | G | A | 2 | 0.002617 | 0.002617 | 1.043 | 1.043 | 0.8339 | 0 |
| 1 | 53841367 | rs1112606 | G | A | 2 | 0.002654 | 0.3091 | 0.9014 | 0.923 | 0.0347 | 77.58 |
| 1 | 54072249 | rs4927014 | G | A | 2 | 0.002691 | 0.002691 | 1.0427 | 1.0427 | 0.7753 | 0 |
| 1 | 54071610 | rs1256387 | A | G | 2 | 0.002861 | 0.002861 | 1.0423 | 1.0423 | 0.8272 | 0 |
| 1 | 54047225 | rs3006884 | G | A | 2 | 0.00299 | 0.00299 | 1.0634 | 1.0634 | 0.5118 | 0 |
| 1 | 54110087 | rs1526907 | T | C | 2 | 0.003128 | 0.003128 | 1.0413 | 1.0413 | 0.6582 | 0 |
| 1 | 53996718 | rs542238 | G | T | 2 | 0.003382 | 0.003382 | 1.0556 | 1.0556 | 0.6589 | 0 |
| 1 | 53986683 | rs2316191 | C | T | 2 | 0.003472 | 0.003472 | 1.0608 | 1.0608 | 0.5465 | 0 |
| 1 | 54108777 | rs1088880 | C | T | 2 | 0.003598 | 0.003598 | 1.0406 | 1.0406 | 0.667 | 0 |
| 1 | 54069274 | rs1122277 | C | T | 2 | 0.003662 | 0.003662 | 0.8941 | 0.8941 | 0.5791 | 0 |
| 1 | 53989785 | rs1088879 | A | C | 2 | 0.003694 | 0.003694 | 1.061 | 1.061 | 0.5631 | 0 |
| 1 | 53994562 | rs494176 | C | T | 2 | 0.00374 | 0.00374 | 1.055 | 1.055 | 0.7052 | 0 |
| 1 | 53984934 | rs495906 | C | G | 2 | 0.003982 | 0.003982 | 1.0602 | 1.0602 | 0.4428 | 0 |
| 1 | 54108494 | rs1088880 | G | A | 2 | 0.004034 | 0.004034 | 1.0401 | 1.0401 | 0.6628 | 0 |
| 1 | 54040670 | rs1078895 | C | G | 2 | 0.004294 | 0.004294 | 1.0408 | 1.0408 | 0.7789 | 0 |
| 1 | 54042336 | rs2950249 | T | G | 2 | 0.005061 | 0.005061 | 0.8896 | 0.8896 | 0.9134 | 0 |
| 1 | 54054084 | rs941124 | A | G | 2 | 0.005103 | 0.005103 | 0.8891 | 0.8891 | 0.9336 | 0 |
| 1 | 53990051 | rs543218 | T | C | 2 | 0.005279 | 0.005279 | 0.8547 | 0.8547 | 0.6344 | 0 |
| 1 | 54061098 | rs2950253 | T | C | 2 | 0.005348 | 0.005348 | 0.8982 | 0.8982 | 0.772 | 0 |
| 1 | 54125929 | rs7690848 | C | T | 2 | 0.005367 | 0.005367 | 0.9272 | 0.9272 | 0.4494 | 0 |
| 1 | 54194992 | rs702491 | T | C | 2 | 0.005424 | 0.02226 | 1.0468 | 1.0453 | 0.2465 | 25.54 |
| 1 | 54065670 | rs3013750 | A | G | 2 | 0.005446 | 0.005446 | 0.8979 | 0.8979 | 0.6237 | 0 |
| 1 | 54056651 | rs3013758 | C | A | 2 | 0.005607 | 0.005607 | 0.8903 | 0.8903 | 0.9345 | 0 |
| 1 | 54059351 | rs3013755 | C | T | 2 | 0.005691 | 0.005691 | 0.8904 | 0.8904 | 0.978 | 0 |
| 1 | 54082352 | rs2141083 | C | T | 2 | 0.00575 | 0.00575 | 0.9624 | 0.9624 | 0.573 | 0 |
| 1 | 54051527 | rs1546806 | A | T | 2 | 0.005782 | 0.005782 | 0.8909 | 0.8909 | 0.9086 | 0 |
| 1 | 54075224 | rs2316195 | G | A | 2 | 0.00599 | 0.00599 | 0.9466 | 0.9466 | 0.9643 | 0 |
| 1 | 53996953 | rs540345 | T | C | 2 | 0.006391 | 0.1759 | 1.0604 | 1.0507 | 0.117 | 59.29 |
| 1 | 54079653 | rs2950262 | A | G | 2 | 0.006868 | 0.006868 | 0.9472 | 0.9472 | 0.899 | 0 |
| 1 | 54110425 | rs6657679 | G | A | 2 | 0.006878 | 0.006878 | 1.0378 | 1.0378 | 0.8258 | 0 |
| 1 | 54110911 | rs4926609 | T | C | 2 | 0.007115 | 0.007115 | 1.0376 | 1.0376 | 0.8416 | 0 |
| 1 | 54127356 | rs7681177 | G | A | 2 | 0.007168 | 0.06741 | 0.9338 | 0.9384 | 0.1901 | 41.76 |
| 1 | 54111733 | rs4926611 | T | C | 2 | 0.007213 | 0.007213 | 1.0375 | 1.0375 | 0.8365 | 0 |

| | | | | | | | | | | | |
|---|---|---|---|---|---|---|---|---|---|---|---|
| 1 | 54110300 | rs6682763 | C | A | 2 | 0.007236 | 0.007236 | 1.0375 | 1.0375 | 0.6804 | 0 |
| 1 | 54108112 | rs6699114 | G | A | 2 | 0.007614 | 0.007614 | 1.0373 | 1.0373 | 0.8474 | 0 |
| 1 | 54107026 | rs4927021 | C | G | 2 | 0.007837 | 0.007837 | 1.0372 | 1.0372 | 0.8518 | 0 |
| 1 | 54131586 | rs7529754 | G | A | 2 | 0.007854 | 0.007854 | 0.8872 | 0.8872 | 0.8935 | 0 |
| 1 | 54106382 | rs1275360 | C | T | 2 | 0.007918 | 0.007918 | 1.0372 | 1.0372 | 0.8545 | 0 |
| 1 | 54352928 | rs1482959 | T | C | 2 | 0.008037 | 0.2908 | 0.9241 | 0.9373 | 0.0496 | 74.06 |
| 1 | 54514570 | rs7743242 | G | C | 2 | 0.008055 | 0.008055 | 1.1836 | 1.1836 | 0.6909 | 0 |
| 1 | 54042387 | rs941127 | A | G | 2 | 0.008715 | 0.008715 | 0.8963 | 0.8963 | 0.8734 | 0 |
| 1 | 54042388 | rs941128 | T | C | 2 | 0.008735 | 0.008735 | 0.8963 | 0.8963 | 0.8739 | 0 |
| 1 | 54054618 | rs6588482 | T | A | 2 | 0.008748 | 0.008748 | 0.9634 | 0.9634 | 0.5623 | 0 |
| 1 | 54113854 | rs6687923 | A | C | 2 | 0.008889 | 0.008889 | 1.0366 | 1.0366 | 0.8569 | 0 |
| 1 | 54452210 | rs7290432 | C | G | 2 | 0.009093 | 0.009093 | 1.0863 | 1.0863 | 0.3538 | 0 |
| 1 | 53997747 | rs481452 | A | G | 2 | 0.009232 | 0.1693 | 1.0582 | 1.0494 | 0.1359 | 55.03 |
| 1 | 54447542 | rs7538972 | C | A | 2 | 0.00925 | 0.00925 | 1.0843 | 1.0843 | 0.4477 | 0 |
| 1 | 54198417 | rs797912 | C | T | 2 | 0.009492 | 0.009492 | 1.0531 | 1.0531 | 0.6706 | 0 |
| 1 | 54077731 | rs7527338 | C | T | 2 | 0.009661 | 0.009661 | 1.0366 | 1.0366 | 0.8437 | 0 |
| 1 | 54469084 | rs7266411 | C | T | 2 | 0.01003 | 0.01003 | 0.9331 | 0.9331 | 0.6705 | 0 |
| 1 | 54045624 | rs1738608 | T | C | 2 | 0.01031 | 0.01031 | 0.9456 | 0.9456 | 0.8082 | 0 |
| 1 | 53974987 | rs1074969 | G | T | 2 | 0.01052 | 0.01052 | 1.056 | 1.056 | 0.5606 | 0 |
| 1 | 54081640 | rs1078896 | C | T | 2 | 0.01067 | 0.01067 | 1.0361 | 1.0361 | 0.6322 | 0 |
| 1 | 54447214 | rs7408510 | T | C | 2 | 0.01074 | 0.01074 | 1.0826 | 1.0826 | 0.4151 | 0 |
| 1 | 54082487 | rs2177957 | C | T | 2 | 0.01079 | 0.01079 | 0.9652 | 0.9652 | 0.6584 | 0 |
| 1 | 54447203 | rs7290430 | T | C | 2 | 0.0109 | 0.0109 | 1.0826 | 1.0826 | 0.3976 | 0 |
| 1 | 54078923 | rs7542387 | A | G | 2 | 0.01111 | 0.01111 | 1.0359 | 1.0359 | 0.7921 | 0 |
| 1 | 54198925 | rs3013763 | C | T | 2 | 0.01112 | 0.01112 | 1.0413 | 1.0413 | 0.6009 | 0 |
| 1 | 54067701 | rs998526 | C | T | 2 | 0.01116 | 0.1916 | 1.095 | 1.0884 | 0.0721 | 69.08 |
| 1 | 54103375 | rs1204524 | A | G | 2 | 0.01116 | 0.01116 | 1.0354 | 1.0354 | 0.7879 | 0 |
| 1 | 54115023 | rs2037739 | G | T | 2 | 0.01123 | 0.01123 | 1.0353 | 1.0353 | 0.5373 | 0 |
| 1 | 54471953 | rs7266411 | T | G | 2 | 0.01137 | 0.01137 | 0.9333 | 0.9333 | 0.6973 | 0 |
| 1 | 54079964 | rs1180129 | G | A | 2 | 0.01156 | 0.01156 | 1.0357 | 1.0357 | 0.6975 | 0 |
| 1 | 54045718 | rs1738610 | T | A | 2 | 0.01181 | 0.01181 | 0.9466 | 0.9466 | 0.7772 | 0 |
| 1 | 54043786 | rs1158534 | G | A | 2 | 0.01182 | 0.01182 | 0.9466 | 0.9466 | 0.7743 | 0 |
| 1 | 54084091 | rs1214000 | C | A | 2 | 0.01184 | 0.01184 | 1.0353 | 1.0353 | 0.9491 | 0 |
| 1 | 53990715 | rs492600 | T | C | 2 | 0.01219 | 0.2053 | 1.055 | 1.046 | 0.1244 | 57.64 |
| 1 | 54622930 | rs7475167 | G | A | 2 | 0.01231 | 0.01231 | 0.8967 | 0.8967 | 0.7507 | 0 |
| 1 | 54077958 | rs2056695 | G | A | 2 | 0.01257 | 0.01257 | 1.0352 | 1.0352 | 0.6907 | 0 |
| 1 | 53643887 | rs1274529 | C | T | 2 | 0.0126 | 0.0126 | 1.0359 | 1.0359 | 0.5225 | 0 |
| 1 | 54446576 | rs1207385 | C | T | 2 | 0.0133 | 0.0133 | 1.0806 | 1.0806 | 0.448 | 0 |
| 1 | 53591846 | rs1383600 | A | G | 2 | 0.01345 | 0.01345 | 1.0455 | 1.0455 | 0.4847 | 0 |
| 1 | 54084347 | rs1526909 | G | C | 2 | 0.01473 | 0.01473 | 1.0341 | 1.0341 | 0.7601 | 0 |
| 1 | 54108212 | rs1120618 | C | G | 2 | 0.01476 | 0.01476 | 0.9519 | 0.9519 | 0.6073 | 0 |
| 1 | 53777631 | rs2788032 | A | C | 2 | 0.01522 | 0.01522 | 0.9386 | 0.9386 | 0.5732 | 0 |
| 1 | 54011869 | rs7523635 | G | A | 2 | 0.01556 | 0.01556 | 1.0336 | 1.0336 | 0.8552 | 0 |
| 1 | 54452523 | rs7290432 | T | C | 2 | 0.01561 | 0.03683 | 1.0798 | 1.0783 | 0.2582 | 21.78 |
| 1 | 53634415 | rs2876790 | A | G | 2 | 0.0158 | 0.03897 | 0.9383 | 0.9405 | 0.2789 | 14.7 |
| 1 | 54102943 | rs1738291 | C | T | 2 | 0.01605 | 0.01605 | 0.9524 | 0.9524 | 0.6414 | 0 |
| 1 | 54107269 | rs5636013 | C | A | 2 | 0.01619 | 0.01619 | 0.9526 | 0.9526 | 0.6296 | 0 |
| 1 | 54542919 | rs1023094 | G | C | 2 | 0.01658 | 0.01658 | 0.9497 | 0.9497 | 0.69 | 0 |

| | | | | | | | | | | | |
|---|---|---|---|---|---|---|---|---|---|---|---|
| 1 | 54085048 | rs4927019 | C | T | 2 | 0.01666 | 0.01666 | 1.0335 | 1.0335 | 0.7274 | 0 |
| 1 | 54069674 | rs1879738 | A | G | 2 | 0.01691 | 0.1575 | 1.0885 | 1.083 | 0.1155 | 59.64 |
| 1 | 54452731 | rs5953548 | G | T | 2 | 0.01693 | 0.04585 | 1.0788 | 1.077 | 0.2455 | 25.85 |
| 1 | 54060991 | rs1207200 | C | A | 2 | 0.01776 | 0.04638 | 0.9664 | 0.9674 | 0.2557 | 22.59 |
| 1 | 54042714 | rs2950250 | T | G | 2 | 0.01844 | 0.01844 | 1.0339 | 1.0339 | 0.8741 | 0 |
| 1 | 54041558 | rs4926603 | G | A | 2 | 0.01887 | 0.01887 | 0.9502 | 0.9502 | 0.6572 | 0 |
| 1 | 54013465 | rs2316194 | C | T | 2 | 0.01891 | 0.01891 | 1.0327 | 1.0327 | 0.7917 | 0 |
| 1 | 54119578 | rs1738733 | G | A | 2 | 0.01893 | 0.04244 | 0.9662 | 0.967 | 0.2666 | 18.98 |
| 1 | 54064469 | rs1320385 | A | G | 2 | 0.01897 | 0.1309 | 1.0867 | 1.0822 | 0.1437 | 53.22 |
| 1 | 54055318 | rs1120617 | T | C | 2 | 0.01924 | 0.01924 | 0.9669 | 0.9669 | 0.3439 | 0 |
| 1 | 54060016 | rs3522700 | G | C | 2 | 0.01958 | 0.02322 | 0.9668 | 0.967 | 0.3076 | 3.95 |
| 1 | 53581886 | rs1213881 | G | A | 2 | 0.01997 | 0.2366 | 0.9001 | 0.7724 | 0.049 | 74.2 |
| 1 | 53644258 | rs1256693 | G | A | 2 | 0.01999 | 0.01999 | 1.0331 | 1.0331 | 0.7471 | 0 |
| 1 | 54176721 | rs1158598 | A | G | 2 | 0.01999 | 0.1732 | 0.9418 | 0.9483 | 0.1477 | 52.29 |
| 1 | 54071428 | rs1738245 | G | A | 2 | 0.02026 | 0.05323 | 0.9675 | 0.9685 | 0.2513 | 24.01 |
| 1 | 54048719 | rs2950252 | C | G | 2 | 0.02029 | 0.02029 | 1.0334 | 1.0334 | 0.8538 | 0 |
| 1 | 54116788 | rs6685730 | C | T | 2 | 0.02033 | 0.02033 | 1.0324 | 1.0324 | 0.5638 | 0 |
| 1 | 54063550 | rs6177029 | G | A | 2 | 0.02045 | 0.02045 | 0.951 | 0.951 | 0.5029 | 0 |
| 1 | 54197688 | rs702489 | A | G | 2 | 0.02069 | 0.02069 | 1.0468 | 1.0468 | 0.7385 | 0 |
| 1 | 54060248 | rs4307514 | T | C | 2 | 0.0207 | 0.0247 | 0.9672 | 0.9673 | 0.307 | 4.17 |
| 1 | 54116416 | rs6697692 | A | G | 2 | 0.02076 | 0.02076 | 1.0323 | 1.0323 | 0.7 | 0 |
| 1 | 54072471 | rs4927015 | G | A | 2 | 0.02083 | 0.02083 | 1.0323 | 1.0323 | 0.6852 | 0 |
| 1 | 53979087 | rs943516 | G | T | 2 | 0.02098 | 0.02098 | 1.0493 | 1.0493 | 0.7242 | 0 |
| 1 | 54012282 | rs1221707 | A | G | 2 | 0.02098 | 0.02098 | 1.0321 | 1.0321 | 0.7483 | 0 |
| 1 | 53987508 | rs519786 | A | G | 2 | 0.02122 | 0.1619 | 1.0515 | 1.0449 | 0.1818 | 43.9 |
| 1 | 54059368 | rs6662142 | T | C | 2 | 0.02126 | 0.05105 | 0.9673 | 0.9682 | 0.259 | 21.5 |
| 1 | 53987717 | rs490654 | A | T | 2 | 0.02147 | 0.1654 | 1.0513 | 1.0446 | 0.1791 | 44.61 |
| 1 | 53982525 | rs559991 | G | A | 2 | 0.02202 | 0.183 | 1.0523 | 1.0445 | 0.1857 | 42.9 |
| 1 | 54061446 | rs1088879 | G | A | 2 | 0.02226 | 0.05093 | 0.9678 | 0.9687 | 0.2619 | 20.56 |
| 1 | 54066318 | rs1203667 | C | T | 2 | 0.02249 | 0.1722 | 1.0842 | 1.0789 | 0.1199 | 58.65 |
| 1 | 54011903 | rs540126 | A | G | 2 | 0.02261 | 0.02261 | 1.0318 | 1.0318 | 0.8974 | 0 |
| 1 | 54011192 | rs2986655 | A | G | 2 | 0.0227 | 0.0227 | 1.0318 | 1.0318 | 0.8975 | 0 |
| 1 | 54065179 | rs941126 | T | C | 2 | 0.02284 | 0.1694 | 1.084 | 1.0789 | 0.1227 | 58.03 |
| 1 | 54067154 | rs4926605 | T | C | 2 | 0.02291 | 0.1648 | 1.084 | 1.079 | 0.1263 | 57.23 |
| 1 | 54313904 | rs1143012 | C | A | 2 | 0.02294 | 0.02512 | 0.8937 | 0.893 | 0.3105 | 2.76 |
| 1 | 54066376 | rs1202256 | G | A | 2 | 0.02296 | 0.1752 | 1.084 | 1.0786 | 0.1187 | 58.93 |
| 1 | 54065648 | rs1211659 | G | A | 2 | 0.02298 | 0.1708 | 1.084 | 1.0787 | 0.122 | 58.18 |
| 1 | 54065924 | rs1272391 | G | A | 2 | 0.02301 | 0.171 | 1.0839 | 1.0787 | 0.1219 | 58.2 |
| 1 | 54066402 | rs1202958 | A | G | 2 | 0.02304 | 0.171 | 1.0839 | 1.0787 | 0.122 | 58.19 |
| 1 | 54066012 | rs1272733 | A | G | 2 | 0.02308 | 0.1714 | 1.0839 | 1.0787 | 0.1218 | 58.24 |
| 1 | 54066506 | rs1120617 | C | T | 2 | 0.02308 | 0.1717 | 1.0839 | 1.0787 | 0.1216 | 58.28 |
| 1 | 54065997 | rs1202946 | A | T | 2 | 0.02314 | 0.172 | 1.0839 | 1.0786 | 0.1215 | 58.31 |
| 1 | 54069983 | rs1879736 | T | C | 2 | 0.02321 | 0.1704 | 1.0838 | 1.0786 | 0.1229 | 57.98 |
| 1 | 54065831 | rs1272387 | G | T | 2 | 0.02336 | 0.1743 | 1.0837 | 1.0785 | 0.1203 | 58.56 |
| 1 | 54066038 | rs1203660 | C | T | 2 | 0.02336 | 0.1741 | 1.0837 | 1.0785 | 0.1205 | 58.52 |
| 1 | 54070013 | rs6663966 | C | T | 2 | 0.02337 | 0.172 | 1.0837 | 1.0785 | 0.1222 | 58.15 |
| 1 | 54066687 | rs1207940 | C | T | 2 | 0.02339 | 0.1743 | 1.0837 | 1.0784 | 0.1204 | 58.55 |
| 1 | 54066752 | rs1206439 | T | C | 2 | 0.02339 | 0.1743 | 1.0837 | 1.0784 | 0.1204 | 58.55 |

| | | | | | | | | | | | |
|---|---|---|---|---|---|---|---|---|---|---|---|
| 1 | 54065778 | rs1272343 | C | T | 2 | 0.02341 | 0.1748 | 1.0837 | 1.0784 | 0.1201 | 58.61 |
| 1 | 54067102 | rs1120617 | C | T | 2 | 0.02341 | 0.1746 | 1.0837 | 1.0784 | 0.1202 | 58.58 |
| 1 | 54066663 | rs1207940 | C | T | 2 | 0.02344 | 0.1748 | 1.0837 | 1.0784 | 0.1201 | 58.6 |
| 1 | 54065523 | rs1212038 | T | C | 2 | 0.02349 | 0.1756 | 1.0836 | 1.0783 | 0.1197 | 58.7 |
| 1 | 54037576 | rs5624322 | T | C | 2 | 0.02368 | 0.02368 | 0.9512 | 0.9512 | 0.5987 | 0 |
| 1 | 54069725 | rs1879737 | C | A | 2 | 0.02375 | 0.1755 | 1.0835 | 1.0782 | 0.1204 | 58.53 |
| 1 | 53893407 | rs1115981 | C | T | 2 | 0.02404 | 0.02404 | 0.9597 | 0.9597 | 0.5523 | 0 |
| 1 | 53987000 | rs545467 | C | A | 2 | 0.02448 | 0.1723 | 1.0499 | 1.0435 | 0.1787 | 44.71 |
| 1 | 53907185 | rs5594609 | C | A | 2 | 0.02466 | 0.1126 | 1.1048 | 1.1211 | 0.1237 | 57.81 |
| 1 | 54013075 | rs2316192 | A | T | 2 | 0.0249 | 0.0249 | 1.0313 | 1.0313 | 0.9926 | 0 |
| 1 | 54120011 | rs6177660 | A | G | 2 | 0.02502 | 0.02502 | 0.9539 | 0.9539 | 0.7701 | 0 |
| 1 | 54104301 | rs6177544 | G | A | 2 | 0.02515 | 0.02515 | 0.9556 | 0.9556 | 0.5204 | 0 |
| 1 | 54061779 | rs6588483 | G | A | 2 | 0.02533 | 0.02533 | 0.9527 | 0.9527 | 0.477 | 0 |
| 1 | 54117225 | rs7263800 | T | G | 2 | 0.02551 | 0.02551 | 0.8652 | 0.8652 | 0.9544 | 0 |
| 1 | 54037176 | rs5599974 | T | G | 2 | 0.02558 | 0.02558 | 0.9525 | 0.9525 | 0.6572 | 0 |
| 1 | 54065303 | rs1211646 | G | A | 2 | 0.02564 | 0.1871 | 1.0824 | 1.077 | 0.1164 | 59.44 |
| 1 | 54040105 | rs5620832 | C | G | 2 | 0.02568 | 0.02568 | 0.9525 | 0.9525 | 0.6153 | 0 |
| 1 | 54066599 | rs1738222 | A | T | 2 | 0.02571 | 0.02571 | 0.9531 | 0.9531 | 0.4683 | 0 |
| 1 | 54010926 | rs1120617 | G | A | 2 | 0.02572 | 0.02572 | 1.0311 | 1.0311 | 0.8901 | 0 |
| 1 | 54035689 | rs1078895 | T | C | 2 | 0.02577 | 0.02577 | 1.0514 | 1.0514 | 0.5656 | 0 |
| 1 | 54070831 | rs1206252 | T | C | 2 | 0.02579 | 0.1833 | 1.0822 | 1.0769 | 0.1195 | 58.75 |
| 1 | 54067311 | rs6177029 | G | A | 2 | 0.02603 | 0.02603 | 0.9531 | 0.9531 | 0.4669 | 0 |
| 1 | 54071011 | rs6176826 | G | A | 2 | 0.0261 | 0.0261 | 0.9532 | 0.9532 | 0.4732 | 0 |
| 1 | 54621886 | rs585136 | A | T | 2 | 0.02615 | 0.02615 | 1.0362 | 1.0362 | 0.8697 | 0 |
| 1 | 54584278 | rs1157962 | A | G | 2 | 0.02642 | 0.05102 | 0.9704 | 0.971 | 0.2705 | 17.64 |
| 1 | 54068567 | rs1078895 | G | T | 2 | 0.02656 | 0.1735 | 1.0818 | 1.0769 | 0.1285 | 56.72 |
| 1 | 54450901 | rs7515322 | A | G | 2 | 0.02673 | 0.1252 | 1.0711 | 1.0683 | 0.1671 | 47.62 |
| 1 | 54453500 | rs1123484 | A | G | 2 | 0.02687 | 0.03925 | 1.0751 | 1.0742 | 0.2914 | 10.15 |
| 1 | 54450110 | rs7516944 | T | G | 2 | 0.02697 | 0.06134 | 1.0706 | 1.0693 | 0.2477 | 25.17 |
| 1 | 54468086 | rs7290435 | G | T | 2 | 0.02737 | 0.02737 | 1.0788 | 1.0788 | 0.5769 | 0 |
| 1 | 53910159 | rs7267515 | C | T | 2 | 0.0275 | 0.1218 | 1.1029 | 1.1197 | 0.1192 | 58.8 |
| 1 | 54013462 | rs2316193 | G | C | 2 | 0.02764 | 0.02764 | 1.0308 | 1.0308 | 0.9881 | 0 |
| 1 | 54445348 | rs1120626 | A | G | 2 | 0.02769 | 0.3604 | 1.0574 | 1.0501 | 0.0379 | 76.8 |
| 1 | 54045226 | rs1208471 | G | A | 2 | 0.02789 | 0.02789 | 0.9566 | 0.9566 | 0.628 | 0 |
| 1 | 54451563 | rs1202217 | T | A | 2 | 0.02803 | 0.1344 | 1.0707 | 1.0676 | 0.1625 | 48.75 |
| 1 | 54450646 | rs7548544 | C | G | 2 | 0.02811 | 0.129 | 1.0706 | 1.0677 | 0.1678 | 47.44 |
| 1 | 53981725 | rs495172 | T | G | 2 | 0.02812 | 0.1859 | 1.048 | 1.0416 | 0.1804 | 44.28 |
| 1 | 54011124 | rs2986656 | T | C | 2 | 0.02842 | 0.02842 | 1.0306 | 1.0306 | 0.9202 | 0 |
| 1 | 53643199 | rs6704459 | G | C | 2 | 0.02893 | 0.02893 | 1.031 | 1.031 | 0.6946 | 0 |
| 1 | 54451080 | rs7525809 | G | A | 2 | 0.02926 | 0.1353 | 1.0701 | 1.0671 | 0.1647 | 48.2 |
| 1 | 54109522 | rs1738721 | C | T | 2 | 0.02944 | 0.02944 | 0.9552 | 0.9552 | 0.7286 | 0 |
| 1 | 54195034 | rs1240612 | C | T | 2 | 0.02976 | 0.5291 | 0.9622 | 0.9733 | 0.0218 | 81 |
| 1 | 54121391 | rs1738737 | T | C | 2 | 0.02983 | 0.02983 | 0.9552 | 0.9552 | 0.8697 | 0 |
| 1 | 53591918 | rs1679966 | C | A | 2 | 0.02988 | 0.02988 | 0.9587 | 0.9587 | 0.7461 | 0 |
| 1 | 54456041 | rs7543936 | A | G | 2 | 0.02999 | 0.07044 | 1.0724 | 1.0706 | 0.2445 | 26.16 |
| 1 | 54024598 | rs7686568 | G | A | 2 | 0.03047 | 0.03047 | 0.913 | 0.913 | 0.7939 | 0 |
| 1 | 54112990 | rs1158955 | T | C | 2 | 0.03062 | 0.03062 | 0.9554 | 0.9554 | 0.7445 | 0 |
| 1 | 54455252 | rs7290433 | C | T | 2 | 0.03078 | 0.06153 | 1.0714 | 1.07 | 0.2606 | 20.98 |

| | | | | | | | | | | | |
|---|---|---|---|---|---|---|---|---|---|---|---|
| 1 | 54454769 | rs7290433 | G | A | 2 | 0.03081 | 0.05995 | 1.0715 | 1.07 | 0.2631 | 20.14 |
| 1 | 54119284 | rs7548123 | C | G | 2 | 0.03102 | 0.03102 | 0.9555 | 0.9555 | 0.8539 | 0 |
| 1 | 54455555 | rs7290433 | A | C | 2 | 0.03124 | 0.06259 | 1.0712 | 1.0697 | 0.26 | 21.18 |
| 1 | 54120689 | rs1158863 | G | C | 2 | 0.03129 | 0.03129 | 0.9556 | 0.9556 | 0.8508 | 0 |
| 1 | 54120733 | rs1157723 | A | G | 2 | 0.03129 | 0.03129 | 0.9556 | 0.9556 | 0.8508 | 0 |
| 1 | 54455992 | rs7554087 | G | A | 2 | 0.03132 | 0.06324 | 1.0712 | 1.0697 | 0.2592 | 21.44 |
| 1 | 54013042 | rs6693508 | T | G | 2 | 0.03138 | 0.03138 | 1.03 | 1.03 | 0.8488 | 0 |
| 1 | 54439637 | rs7530434 | C | T | 2 | 0.03142 | 0.4561 | 1.0564 | 1.0477 | 0.0154 | 82.97 |
| 1 | 54419015 | rs7611634 | T | C | 2 | 0.03149 | 0.2624 | 1.1051 | 1.0932 | 0.0935 | 64.44 |
| 1 | 54578401 | rs954878 | G | A | 2 | 0.03157 | 0.08092 | 0.9712 | 0.9722 | 0.2403 | 27.47 |
| 1 | 53981615 | rs496126 | A | G | 2 | 0.03161 | 0.2648 | 1.0478 | 1.0392 | 0.1458 | 52.74 |
| 1 | 53999716 | rs498473 | G | A | 2 | 0.03167 | 0.03167 | 1.063 | 1.063 | 0.3226 | 0 |
| 1 | 53960488 | rs1288610 | A | G | 2 | 0.03174 | 0.1179 | 1.0404 | 1.0434 | 0.1469 | 52.48 |
| 1 | 54146952 | rs7289679 | A | G | 2 | 0.03184 | 0.03184 | 1.0762 | 1.0762 | 0.5684 | 0 |
| 1 | 54450025 | rs1204915 | T | C | 2 | 0.03202 | 0.09897 | 1.0683 | 1.0663 | 0.2093 | 36.56 |
| 1 | 54625682 | rs1088883 | T | C | 2 | 0.03207 | 0.03207 | 1.0353 | 1.0353 | 0.9377 | 0 |
| 1 | 54043386 | rs1158527 | G | A | 2 | 0.03225 | 0.03225 | 0.9577 | 0.9577 | 0.6221 | 0 |
| 1 | 54045180 | rs1208997 | T | C | 2 | 0.03253 | 0.03253 | 0.9578 | 0.9578 | 0.6229 | 0 |
| 1 | 54599439 | rs7367806 | T | G | 2 | 0.03257 | 0.03257 | 1.041 | 1.041 | 0.6228 | 0 |
| 1 | 54448873 | rs5731705 | C | G | 2 | 0.03276 | 0.09746 | 1.0694 | 1.0673 | 0.2138 | 35.3 |
| 1 | 53978119 | rs6588480 | A | G | 2 | 0.03312 | 0.04265 | 1.041 | 1.0405 | 0.3027 | 5.85 |
| 1 | 54045856 | rs7408474 | T | G | 2 | 0.03312 | 0.03312 | 0.958 | 0.958 | 0.6154 | 0 |
| 1 | 54106705 | rs1738715 | T | G | 2 | 0.03324 | 0.03324 | 0.956 | 0.956 | 0.6642 | 0 |
| 1 | 54118490 | rs4626818 | G | T | 2 | 0.03348 | 0.03348 | 0.9561 | 0.9561 | 0.8191 | 0 |
| 1 | 54449103 | rs7290431 | C | T | 2 | 0.03362 | 0.03362 | 1.0679 | 1.0679 | 0.3356 | 0 |
| 1 | 54087629 | rs7939775 | C | T | 2 | 0.03363 | 0.03363 | 0.9544 | 0.9544 | 0.5802 | 0 |
| 1 | 54146291 | rs5763434 | G | A | 2 | 0.0343 | 0.0343 | 1.0746 | 1.0746 | 0.5349 | 0 |
| 1 | 54453489 | rs7290432 | C | T | 2 | 0.03457 | 0.09108 | 1.0698 | 1.0675 | 0.2297 | 30.68 |
| 1 | 54457478 | rs1983585 | C | T | 2 | 0.03483 | 0.07614 | 1.0704 | 1.0686 | 0.2498 | 24.49 |
| 1 | 54461121 | rs7924597 | A | T | 2 | 0.03507 | 0.09738 | 1.0701 | 1.0675 | 0.2244 | 32.24 |
| 1 | 54454069 | rs5968363 | G | A | 2 | 0.0351 | 0.09196 | 1.0696 | 1.0674 | 0.2299 | 30.62 |
| 1 | 54042889 | rs3108391 | C | T | 2 | 0.03542 | 0.03542 | 1.043 | 1.043 | 0.4509 | 0 |
| 1 | 54116576 | rs6177660 | T | C | 2 | 0.03569 | 0.03569 | 0.9569 | 0.9569 | 0.8752 | 0 |
| 1 | 54644197 | rs621359 | G | A | 2 | 0.03578 | 0.03578 | 1.034 | 1.034 | 0.6685 | 0 |
| 1 | 54117388 | rs7610164 | A | G | 2 | 0.03613 | 0.03613 | 0.957 | 0.957 | 0.9051 | 0 |
| 1 | 53717598 | rs1905238 | C | T | 2 | 0.03628 | 0.03628 | 0.9076 | 0.9076 | 0.8562 | 0 |
| 1 | 54599341 | rs1134563 | G | A | 2 | 0.0363 | 0.0363 | 0.9572 | 0.9572 | 0.9656 | 0 |
| 1 | 54104906 | rs1738296 | T | C | 2 | 0.03632 | 0.03632 | 0.9567 | 0.9567 | 0.6822 | 0 |
| 1 | 54437173 | rs1120626 | G | A | 2 | 0.03646 | 0.1322 | 1.0738 | 1.0706 | 0.187 | 42.56 |
| 1 | 54118424 | rs1157607 | A | G | 2 | 0.03661 | 0.03661 | 0.9569 | 0.9569 | 0.8194 | 0 |
| 1 | 54133932 | rs7514580 | C | T | 2 | 0.03665 | 0.03665 | 1.0811 | 1.0811 | 0.9341 | 0 |
| 1 | 54066538 | rs1202629 | T | C | 2 | 0.03752 | 0.1463 | 1.0758 | 1.0721 | 0.1762 | 45.35 |
| 1 | 54625292 | rs660514 | A | G | 2 | 0.03776 | 0.03776 | 1.0339 | 1.0339 | 0.9662 | 0 |
| 1 | 54457482 | rs1983586 | A | G | 2 | 0.03783 | 0.1115 | 1.069 | 1.0662 | 0.2148 | 35.03 |
| 1 | 54118849 | rs7522391 | G | A | 2 | 0.03789 | 0.03789 | 0.9571 | 0.9571 | 0.8079 | 0 |
| 1 | 54103346 | rs5593205 | G | A | 2 | 0.03799 | 0.03799 | 0.9571 | 0.9571 | 0.6709 | 0 |
| 1 | 54458569 | rs5725525 | G | A | 2 | 0.03842 | 0.1171 | 1.0687 | 1.0658 | 0.2097 | 36.45 |
| 1 | 54115483 | rs7512380 | G | A | 2 | 0.03843 | 0.03843 | 0.9572 | 0.9572 | 0.7197 | 0 |

| | | | | | | | | | | | |
|---|---|---|---|---|---|---|---|---|---|---|---|
| 1 | 53946485 | rs1288622 | A | G | 2 | 0.03851 | 0.03851 | 0.9589 | 0.9589 | 0.7975 | 0 |
| 1 | 54095360 | rs1880861 | C | T | 2 | 0.03887 | 0.03887 | 0.9565 | 0.9565 | 0.587 | 0 |
| 1 | 54009279 | rs496933 | C | G | 2 | 0.03891 | 0.03891 | 1.0301 | 1.0301 | 0.6285 | 0 |
| 1 | 54444147 | rs1147666 | C | T | 2 | 0.03915 | 0.1337 | 1.1002 | 1.0948 | 0.198 | 39.65 |
| 1 | 53520594 | rs7612839 | A | T | 2 | 0.04026 | 0.04026 | 1.1748 | 1.1748 | 0.6496 | 0 |
| 1 | 54475188 | rs7290437 | C | T | 2 | 0.0403 | 0.0403 | 1.0722 | 1.0722 | 0.3822 | 0 |
| 1 | 54009328 | rs3855969 | C | T | 2 | 0.04083 | 0.4157 | 1.0453 | 1.0339 | 0.0789 | 67.61 |
| 1 | 54075857 | rs1710890 | G | A | 2 | 0.04086 | 0.04086 | 0.8699 | 0.8699 | 0.5985 | 0 |
| 1 | 53912739 | rs7267515 | G | C | 2 | 0.04126 | 0.04126 | 1.1305 | 1.1305 | 0.4799 | 0 |
| 1 | 54471517 | rs7878293 | G | A | 2 | 0.04131 | 0.4311 | 1.0859 | 1.074 | 0.0265 | 79.68 |
| 1 | 54408279 | rs6177485 | T | C | 2 | 0.04137 | 0.04137 | 0.9355 | 0.9355 | 0.5061 | 0 |
| 1 | 53927836 | rs7658208 | G | T | 2 | 0.04151 | 0.07462 | 0.9137 | 0.9098 | 0.2409 | 27.28 |
| 1 | 54535919 | rs7266413 | G | A | 2 | 0.04162 | 0.04162 | 0.9467 | 0.9467 | 0.7829 | 0 |
| 1 | 54224270 | rs1181191 | A | C | 2 | 0.04189 | 0.04189 | 0.9538 | 0.9538 | 0.9841 | 0 |
| 1 | 54440729 | rs7290250 | T | C | 2 | 0.04198 | 0.4572 | 1.0544 | 1.0458 | 0.0228 | 80.71 |
| 1 | 54458862 | rs6085784 | C | T | 2 | 0.0423 | 0.1521 | 1.0673 | 1.0637 | 0.1842 | 43.28 |
| 1 | 53655037 | rs7800331 | G | A | 2 | 0.04235 | 0.05067 | 1.1442 | 1.1434 | 0.3022 | 6.05 |
| 1 | 54443867 | rs1120626 | C | T | 2 | 0.04247 | 0.05083 | 1.071 | 1.0706 | 0.3021 | 6.1 |
| 1 | 53503094 | rs5982185 | A | C | 2 | 0.04291 | 0.04291 | 1.1729 | 1.1729 | 0.6492 | 0 |
| 1 | 54130885 | rs6158812 | C | A | 2 | 0.04312 | 0.04312 | 1.0792 | 1.0792 | 0.9281 | 0 |
| 1 | 54007612 | rs1275267 | T | C | 2 | 0.04313 | 0.04313 | 1.0284 | 1.0284 | 0.9818 | 0 |
| 1 | 54115984 | rs5619857 | C | T | 2 | 0.04328 | 0.04328 | 0.9583 | 0.9583 | 0.7747 | 0 |
| 1 | 54261335 | rs1159161 | C | T | 2 | 0.04338 | 0.324 | 0.9176 | 0.9326 | 0.1205 | 58.51 |
| 1 | 54442913 | rs1120626 | A | G | 2 | 0.04346 | 0.04346 | 1.0842 | 1.0842 | 0.6408 | 0 |
| 1 | 53842099 | rs1316436 | C | T | 2 | 0.0437 | 0.0437 | 1.0283 | 1.0283 | 0.5128 | 0 |
| 1 | 53590630 | rs3766793 | T | C | 2 | 0.04393 | 0.04393 | 1.0383 | 1.0383 | 0.5174 | 0 |
| 1 | 53958910 | rs1288613 | C | T | 2 | 0.04396 | 0.04396 | 1.0375 | 1.0375 | 0.3589 | 0 |
| 1 | 53644967 | rs6588471 | C | T | 2 | 0.04407 | 0.04407 | 1.0287 | 1.0287 | 0.7511 | 0 |
| 1 | 53924333 | rs7487855 | G | A | 2 | 0.04441 | 0.1287 | 0.9128 | 0.902 | 0.1536 | 50.88 |
| 1 | 53947791 | rs943524 | G | C | 2 | 0.04443 | 0.04443 | 0.9639 | 0.9639 | 0.4653 | 0 |
| 1 | 54074185 | rs5601102 | G | A | 2 | 0.04458 | 0.04458 | 0.8712 | 0.8712 | 0.5238 | 0 |
| 1 | 54576448 | rs7623047 | T | C | 2 | 0.04468 | 0.04468 | 1.0693 | 1.0693 | 0.7571 | 0 |
| 1 | 54001983 | rs545514 | A | G | 2 | 0.04475 | 0.1488 | 1.0396 | 1.0374 | 0.1958 | 40.25 |
| 1 | 53946644 | rs1288621 | A | G | 2 | 0.04477 | 0.04477 | 0.9639 | 0.9639 | 0.4523 | 0 |
| 1 | 54085864 | rs3006899 | T | C | 2 | 0.04483 | 0.04483 | 0.9579 | 0.9579 | 0.6446 | 0 |
| 1 | 53955202 | rs1288615 | T | C | 2 | 0.04487 | 0.04487 | 0.964 | 0.964 | 0.4113 | 0 |
| 1 | 53963481 | rs1074969 | G | A | 2 | 0.04537 | 0.0548 | 1.0373 | 1.0377 | 0.2944 | 9.04 |
| 1 | 54010065 | rs525145 | A | G | 2 | 0.04581 | 0.1589 | 1.0396 | 1.0373 | 0.1871 | 42.53 |
| 1 | 54456131 | rs7534706 | C | T | 2 | 0.04585 | 0.09925 | 1.0652 | 1.0634 | 0.2416 | 27.06 |
| 1 | 53926803 | rs1288629 | C | G | 2 | 0.04623 | 0.04623 | 0.964 | 0.964 | 0.6178 | 0 |
| 1 | 54231418 | rs4130279 | C | T | 2 | 0.04627 | 0.04627 | 0.9253 | 0.9253 | 0.4503 | 0 |
| 1 | 54014492 | rs563403 | A | G | 2 | 0.04649 | 0.3869 | 1.0495 | 1.0379 | 0.1013 | 62.75 |
| 1 | 54558042 | rs7571320 | G | A | 2 | 0.04677 | 0.04677 | 1.0745 | 1.0745 | 0.9059 | 0 |
| 1 | 54465945 | rs6177485 | C | T | 2 | 0.0471 | 0.0471 | 0.9359 | 0.9359 | 0.9941 | 0 |
| 1 | 54484696 | rs7706216 | C | A | 2 | 0.04734 | 0.2809 | 1.0809 | 1.071 | 0.114 | 59.96 |
| 1 | 54489267 | rs1158922 | A | G | 2 | 0.04734 | 0.2318 | 1.0666 | 1.0611 | 0.133 | 55.69 |
| 1 | 54031895 | rs1274914 | T | C | 2 | 0.04748 | 0.04748 | 0.9467 | 0.9467 | 0.5516 | 0 |
| 1 | 53980389 | rs4131336 | C | T | 2 | 0.04762 | 0.04762 | 0.9536 | 0.9536 | 0.4219 | 0 |

| | | | | | | | | | | | |
|---|---|---|---|---|---|---|---|---|---|---|---|
| 1 | 53699129 | rs1719265 | G | C | 2 | 0.04881 | 0.04881 | 1.0519 | 1.0519 | 0.4306 | 0 |
| 1 | 54036317 | rs4927009 | G | C | 2 | 0.04916 | 0.04916 | 0.9563 | 0.9563 | 0.4892 | 0 |
| 1 | 54657551 | rs5635308 | G | A | 2 | 0.04926 | 0.43 | 0.9587 | 0.9683 | 0.0738 | 68.71 |
| 1 | 54204324 | rs702488 | T | C | 2 | 0.04936 | 0.04936 | 1.0415 | 1.0415 | 0.4428 | 0 |
| 1 | 53926873 | rs1288630 | G | C | 2 | 0.04949 | 0.04949 | 0.9645 | 0.9645 | 0.5495 | 0 |
| 1 | 53926037 | rs1288628 | T | G | 2 | 0.04951 | 0.04951 | 0.9645 | 0.9645 | 0.5473 | 0 |
| 1 | 53589130 | rs1120610 | G | A | 2 | 0.0499 | 0.0499 | 1.0395 | 1.0395 | 0.7501 | 0 |
| 1 | 53589671 | rs3753596 | C | T | 2 | 0.05051 | 0.05051 | 1.0396 | 1.0396 | 0.6517 | 0 |
| 1 | 54129487 | rs1119744 | G | A | 2 | 0.05072 | 0.05072 | 0.8422 | 0.8422 | 0.5188 | 0 |
| 1 | 54129258 | rs5765175 | G | T | 2 | 0.05079 | 0.05079 | 1.0761 | 1.0761 | 0.9779 | 0 |
| 1 | 54185389 | rs1120621 | C | T | 2 | 0.05108 | 0.1783 | 0.9245 | 0.93 | 0.1916 | 41.35 |
| 1 | 53584825 | rs1209255 | G | T | 2 | 0.05122 | 0.1381 | 1.0445 | 1.0523 | 0.1525 | 51.15 |
| 1 | 54128434 | rs7289677 | T | A | 2 | 0.05215 | 0.05215 | 1.0756 | 1.0756 | 0.947 | 0 |
| 1 | 54436645 | rs1120626 | T | C | 2 | 0.05255 | 0.1223 | 1.0684 | 1.0663 | 0.2265 | 31.64 |
| 1 | 54075686 | rs6092622 | G | A | 2 | 0.05402 | 0.05402 | 0.8762 | 0.8762 | 0.4665 | 0 |
| 1 | 54079218 | rs7291067 | T | C | 2 | 0.05432 | 0.05432 | 0.8761 | 0.8761 | 0.4713 | 0 |
| 1 | 54148243 | rs1710910 | T | C | 2 | 0.05532 | 0.05532 | 1.0682 | 1.0682 | 0.7526 | 0 |
| 1 | 54079808 | rs2948055 | C | A | 2 | 0.05576 | 0.05576 | 0.9595 | 0.9595 | 0.5607 | 0 |
| 1 | 53904989 | rs6163920 | A | C | 2 | 0.05603 | 0.3326 | 0.9443 | 0.9522 | 0.1028 | 62.43 |
| 1 | 53587701 | rs3766788 | G | A | 2 | 0.05639 | 0.05639 | 1.0386 | 1.0386 | 0.5775 | 0 |
| 1 | 54409951 | rs8016830 | C | T | 2 | 0.05649 | 0.05649 | 0.9223 | 0.9223 | 0.4719 | 0 |
| 1 | 54439279 | rs1150540 | A | C | 2 | 0.05661 | 0.2595 | 1.0919 | 1.0837 | 0.1275 | 56.96 |
| 1 | 54630158 | rs613381 | C | T | 2 | 0.05735 | 0.05735 | 1.0284 | 1.0284 | 0.4538 | 0 |
| 1 | 54021529 | rs3534596 | T | C | 2 | 0.05823 | 0.05831 | 1.0527 | 1.0527 | 0.3172 | 0.04 |
| 1 | 54469633 | rs1389375 | G | T | 2 | 0.05871 | 0.5083 | 1.0791 | 1.0668 | 0.0163 | 82.67 |
| 1 | 54612692 | rs1164881 | T | C | 2 | 0.05873 | 0.05873 | 0.9423 | 0.9423 | 0.5261 | 0 |
| 1 | 53882558 | rs7514445 | T | C | 2 | 0.05891 | 0.2091 | 0.9189 | 0.9274 | 0.2047 | 37.83 |
| 1 | 53897118 | rs5866356 | C | G | 2 | 0.05894 | 0.3456 | 0.9434 | 0.9518 | 0.0995 | 63.14 |
| 1 | 54662686 | rs1256230 | G | A | 2 | 0.05992 | 0.05992 | 1.0269 | 1.0269 | 0.5422 | 0 |
| 1 | 53897370 | rs5964729 | C | G | 2 | 0.0601 | 0.3461 | 0.9436 | 0.952 | 0.1017 | 62.68 |
| 1 | 53589674 | rs3753597 | C | T | 2 | 0.06052 | 0.06052 | 1.0381 | 1.0381 | 0.7247 | 0 |
| 1 | 54642971 | rs4926621 | T | C | 2 | 0.06098 | 0.06098 | 0.9113 | 0.9113 | 0.5884 | 0 |
| 1 | 54440551 | rs7408449 | G | T | 2 | 0.06113 | 0.6021 | 1.0502 | 1.0389 | 0.0059 | 86.82 |
| 1 | 53972115 | rs2296760 | G | A | 2 | 0.06121 | 0.06121 | 0.9059 | 0.9059 | 0.5912 | 0 |
| 1 | 54342251 | rs1163252 | C | T | 2 | 0.0615 | 0.0615 | 0.9182 | 0.9182 | 0.3236 | 0 |
| 1 | 54441756 | rs1088882 | A | G | 2 | 0.06235 | 0.1039 | 1.0649 | 1.0636 | 0.2636 | 19.98 |
| 1 | 54479358 | rs7266411 | A | C | 2 | 0.06292 | 0.09198 | 0.9603 | 0.9587 | 0.2595 | 21.34 |
| 1 | 54011142 | rs6682171 | T | C | 2 | 0.06295 | 0.2348 | 1.0363 | 1.0335 | 0.1537 | 50.87 |
| 1 | 54151611 | rs7266064 | G | A | 2 | 0.06302 | 0.1244 | 1.0427 | 1.0408 | 0.2574 | 22.03 |
| 1 | 54662346 | rs7525757 | T | G | 2 | 0.06306 | 0.06306 | 1.0265 | 1.0265 | 0.5243 | 0 |
| 1 | 53899760 | rs6692519 | G | C | 2 | 0.06362 | 0.3498 | 0.9442 | 0.9526 | 0.1045 | 62.07 |
| 1 | 54016692 | rs511202 | A | T | 2 | 0.06436 | 0.33 | 1.0454 | 1.0374 | 0.1401 | 54.06 |
| 1 | 54622694 | rs1710992 | G | A | 2 | 0.0646 | 0.0646 | 0.9673 | 0.9673 | 0.5703 | 0 |
| 1 | 53760505 | rs5578981 | T | C | 2 | 0.06467 | 0.7072 | 0.9353 | 1.0846 | 0.0774 | 67.94 |
| 1 | 53862530 | rs1737952 | A | G | 2 | 0.06488 | 0.3194 | 0.921 | 0.9339 | 0.1473 | 52.39 |
| 1 | 54120275 | rs7755183 | C | T | 2 | 0.06506 | 0.06506 | 0.8762 | 0.8762 | 0.3623 | 0 |
| 1 | 54101062 | rs6679903 | A | G | 2 | 0.06522 | 0.06522 | 0.9616 | 0.9616 | 0.651 | 0 |
| 1 | 54664001 | rs3511518 | T | C | 2 | 0.06586 | 0.6123 | 0.9627 | 0.9759 | 0.0292 | 78.98 |

| | | | | | | | | | | | |
|---|---|---|---|---|---|---|---|---|---|---|---|
| 1 | 54698208 | rs610874 | A | T | 2 | 0.06633 | 0.06633 | 1.0256 | 1.0256 | 0.3539 | 0 |
| 1 | 54323705 | rs6177482 | T | C | 2 | 0.06703 | 0.1153 | 0.9469 | 0.9487 | 0.2712 | 17.39 |
| 1 | 54615173 | rs7518597 | G | A | 2 | 0.06706 | 0.06706 | 0.9675 | 0.9675 | 0.4389 | 0 |
| 1 | 54623252 | rs7475095 | T | C | 2 | 0.06736 | 0.06736 | 0.9678 | 0.9678 | 0.6312 | 0 |
| 1 | 54211525 | rs7290060 | G | C | 2 | 0.06739 | 0.06739 | 0.9562 | 0.9562 | 0.8035 | 0 |
| 1 | 53591463 | rs3766794 | G | A | 2 | 0.06781 | 0.06781 | 0.9669 | 0.9669 | 0.6544 | 0 |
| 1 | 53976188 | rs2280511 | C | T | 2 | 0.06844 | 0.06844 | 1.0405 | 1.0405 | 0.3438 | 0 |
| 1 | 54039326 | rs2950248 | T | C | 2 | 0.06846 | 0.1857 | 1.0636 | 1.0617 | 0.1821 | 43.84 |
| 1 | 54654129 | rs4244643 | G | T | 2 | 0.06851 | 0.06851 | 1.0273 | 1.0273 | 0.6361 | 0 |
| 1 | 54623183 | rs7290802 | C | T | 2 | 0.06856 | 0.06856 | 0.9682 | 0.9682 | 0.7032 | 0 |
| 1 | 54600471 | rs1088883 | T | C | 2 | 0.06875 | 0.06875 | 1.0349 | 1.0349 | 0.9623 | 0 |
| 1 | 53874341 | rs1212496 | T | G | 2 | 0.06902 | 0.06902 | 0.9716 | 0.9716 | 0.5888 | 0 |
| 1 | 54620583 | rs1710992 | C | T | 2 | 0.06905 | 0.06905 | 0.9678 | 0.9678 | 0.5629 | 0 |
| 1 | 54456325 | rs7266410 | A | C | 2 | 0.06906 | 0.1045 | 0.9588 | 0.9568 | 0.2501 | 24.39 |
| 1 | 54661150 | rs946448 | C | T | 2 | 0.06928 | 0.06928 | 1.0259 | 1.0259 | 0.5124 | 0 |
| 1 | 54632658 | rs1135357 | G | A | 2 | 0.06982 | 0.06982 | 0.96 | 0.96 | 0.8766 | 0 |
| 1 | 54117630 | rs5882096 | C | T | 2 | 0.07011 | 0.2802 | 0.9001 | 0.916 | 0.2442 | 26.26 |
| 1 | 53926426 | rs1885588 | T | C | 2 | 0.07024 | 0.08723 | 0.9152 | 0.9121 | 0.2883 | 11.31 |
| 1 | 53948733 | rs1298638 | T | C | 2 | 0.07027 | 0.07027 | 0.965 | 0.965 | 0.9769 | 0 |
| 1 | 54627760 | rs618873 | C | G | 2 | 0.07028 | 0.07028 | 1.0293 | 1.0293 | 0.445 | 0 |
| 1 | 54090216 | rs6043062 | T | C | 2 | 0.07085 | 0.1093 | 0.8836 | 0.8874 | 0.292 | 9.94 |
| 1 | 54106668 | rs1738301 | G | C | 2 | 0.07163 | 0.07163 | 0.9622 | 0.9622 | 0.4498 | 0 |
| 1 | 54617963 | rs1112592 | C | A | 2 | 0.07195 | 0.07195 | 0.9682 | 0.9682 | 0.5227 | 0 |
| 1 | 53897818 | rs7597721 | A | T | 2 | 0.07311 | 0.2827 | 0.9416 | 0.9469 | 0.137 | 54.77 |
| 1 | 53997844 | rs1203284 | C | G | 2 | 0.07347 | 0.1799 | 0.9377 | 0.9347 | 0.1634 | 48.52 |
| 1 | 53985443 | rs3887579 | A | C | 2 | 0.07389 | 0.07389 | 1.027 | 1.027 | 0.6853 | 0 |
| 1 | 54111547 | rs1180175 | C | G | 2 | 0.07412 | 0.07412 | 0.843 | 0.843 | 0.6276 | 0 |
| 1 | 54646041 | rs780503 | A | G | 2 | 0.07418 | 0.07418 | 1.0289 | 1.0289 | 0.5761 | 0 |
| 1 | 54616538 | rs6667290 | A | C | 2 | 0.07419 | 0.07419 | 0.9683 | 0.9683 | 0.4948 | 0 |
| 1 | 54221402 | rs7290060 | A | G | 2 | 0.0745 | 0.0745 | 0.958 | 0.958 | 0.4624 | 0 |
| 1 | 53875471 | rs1342969 | C | T | 2 | 0.07514 | 0.07514 | 0.9719 | 0.9719 | 0.4941 | 0 |
| 1 | 53792115 | rs5853408 | G | T | 2 | 0.07521 | 0.07521 | 0.8858 | 0.8858 | 0.54 | 0 |
| 1 | 54615558 | rs6699826 | C | T | 2 | 0.07538 | 0.07538 | 0.9684 | 0.9684 | 0.4772 | 0 |
| 1 | 54120838 | rs1159061 | T | C | 2 | 0.0757 | 0.0757 | 0.9636 | 0.9636 | 0.7805 | 0 |
| 1 | 54038001 | rs1983485 | T | C | 2 | 0.07595 | 0.2411 | 1.0619 | 1.0594 | 0.1465 | 52.57 |
| 1 | 53533727 | rs6151760 | G | C | 2 | 0.07626 | 0.07626 | 1.1595 | 1.1595 | 0.7903 | 0 |
| 1 | 54119089 | rs1180294 | C | T | 2 | 0.07708 | 0.07708 | 0.8772 | 0.8772 | 0.3585 | 0 |
| 1 | 54406760 | rs6177485 | C | T | 2 | 0.07709 | 0.3949 | 0.9477 | 0.9589 | 0.1344 | 55.36 |
| 1 | 53998415 | rs3888344 | T | G | 2 | 0.0771 | 0.1603 | 0.9386 | 0.9362 | 0.193 | 41 |
| 1 | 54698112 | rs643805 | T | A | 2 | 0.07809 | 0.07809 | 1.0246 | 1.0246 | 0.385 | 0 |
| 1 | 54637697 | rs670877 | A | G | 2 | 0.07906 | 0.1843 | 1.0273 | 1.0257 | 0.2213 | 33.15 |
| 1 | 54463737 | rs1044323 | C | A | 2 | 0.07911 | 0.07911 | 0.9716 | 0.9716 | 0.5636 | 0 |
| 1 | 53886578 | rs1147254 | A | G | 2 | 0.07978 | 0.07978 | 0.9254 | 0.9254 | 0.3527 | 0 |
| 1 | 53833205 | rs6176837 | A | G | 2 | 0.07983 | 0.07983 | 0.9247 | 0.9247 | 0.4187 | 0 |
| 1 | 54536010 | rs6674224 | A | T | 2 | 0.08002 | 0.154 | 0.9704 | 0.9677 | 0.1926 | 41.09 |
| 1 | 54658463 | rs7537833 | G | C | 2 | 0.08003 | 0.08003 | 1.0248 | 1.0248 | 0.3489 | 0 |
| 1 | 54637915 | rs6690858 | C | T | 2 | 0.0814 | 0.0814 | 1.0282 | 1.0282 | 0.4846 | 0 |
| 1 | 54211771 | rs1725982 | T | C | 2 | 0.0816 | 0.0816 | 0.9587 | 0.9587 | 0.8217 | 0 |

| | | | | | | | | | | | |
|---|---|---|---|---|---|---|---|---|---|---|---|
| 1 | 53960235 | rs1288611 | G | A | 2 | 0.08174 | 0.08174 | 1.0598 | 1.0598 | 0.3312 | 0 |
| 1 | 53614388 | rs1136528 | G | A | 2 | 0.08302 | 0.09514 | 1.051 | 1.0519 | 0.2949 | 8.84 |
| 1 | 54293297 | rs7864528 | C | T | 2 | 0.08335 | 0.3813 | 0.9311 | 0.9438 | 0.1303 | 56.3 |
| 1 | 53952777 | rs1288616 | G | A | 2 | 0.08371 | 0.08371 | 0.9648 | 0.9648 | 0.6649 | 0 |
| 1 | 53877725 | rs6701841 | G | C | 2 | 0.08385 | 0.1067 | 0.9736 | 0.974 | 0.2941 | 9.16 |
| 1 | 54426758 | rs7721578 | T | C | 2 | 0.08407 | 0.08407 | 1.0775 | 1.0775 | 0.4402 | 0 |
| 1 | 54642866 | rs5606756 | G | C | 2 | 0.08416 | 0.08416 | 0.9716 | 0.9716 | 0.4313 | 0 |
| 1 | 54524608 | rs1496833 | G | A | 2 | 0.08422 | 0.4754 | 1.0579 | 1.0484 | 0.0457 | 74.96 |
| 1 | 53990937 | rs7289333 | C | G | 2 | 0.08463 | 0.08463 | 0.9156 | 0.9156 | 0.3986 | 0 |
| 1 | 53590423 | rs3766792 | C | T | 2 | 0.08503 | 0.08503 | 1.0348 | 1.0348 | 0.7804 | 0 |
| 1 | 53895695 | rs7290300 | C | T | 2 | 0.0854 | 0.399 | 0.9487 | 0.9569 | 0.0973 | 63.62 |
| 1 | 54119083 | rs6588485 | A | G | 2 | 0.08618 | 0.08618 | 1.1193 | 1.1193 | 0.3568 | 0 |
| 1 | 53870466 | rs4926992 | C | A | 2 | 0.0864 | 0.0864 | 0.9767 | 0.9767 | 0.7068 | 0 |
| 1 | 54153321 | rs3013776 | A | G | 2 | 0.08709 | 0.1088 | 0.9459 | 0.9466 | 0.294 | 9.17 |
| 1 | 54036593 | rs3013760 | G | A | 2 | 0.08812 | 0.3227 | 0.9433 | 0.9466 | 0.1066 | 61.59 |
| 1 | 54636684 | rs1399448 | G | A | 2 | 0.08856 | 0.08856 | 1.1309 | 1.1309 | 0.9634 | 0 |
| 1 | 54673879 | rs3491185 | C | G | 2 | 0.08891 | 0.08891 | 0.8951 | 0.8951 | 0.6707 | 0 |
| 1 | 54001823 | rs4126639 | A | G | 2 | 0.08911 | 0.08911 | 1.0244 | 1.0244 | 0.4453 | 0 |
| 1 | 54613529 | rs1157051 | T | G | 2 | 0.08916 | 0.08916 | 1.0884 | 1.0884 | 0.6683 | 0 |
| 1 | 53959435 | rs7529321 | G | A | 2 | 0.0896 | 0.323 | 0.9647 | 0.9597 | 0.0525 | 73.41 |
| 1 | 53960377 | rs1484444 | G | A | 2 | 0.09066 | 0.2227 | 0.9575 | 0.9516 | 0.1238 | 57.79 |
| 1 | 54690867 | rs5680264 | G | A | 2 | 0.09078 | 0.09078 | 1.0559 | 1.0559 | 0.6466 | 0 |
| 1 | 54608251 | rs680509 | C | T | 2 | 0.09121 | 0.09121 | 0.969 | 0.969 | 0.4617 | 0 |
| 1 | 54032922 | rs5865177 | G | C | 2 | 0.09159 | 0.3294 | 0.9716 | 0.9666 | 0.0451 | 75.08 |
| 1 | 54108462 | rs1157990 | G | A | 2 | 0.09425 | 0.09425 | 0.9152 | 0.9152 | 0.4298 | 0 |
| 1 | 54602347 | rs2183302 | G | A | 2 | 0.09434 | 0.09434 | 1.0321 | 1.0321 | 0.9776 | 0 |
| 1 | 54177891 | rs1120620 | T | C | 2 | 0.09438 | 0.09438 | 0.9321 | 0.9321 | 0.3766 | 0 |
| 1 | 53911237 | rs5589533 | C | T | 2 | 0.09464 | 0.2553 | 0.9584 | 0.9519 | 0.097 | 63.69 |
| 1 | 54557290 | rs1204688 | C | T | 2 | 0.09525 | 0.1564 | 0.9766 | 0.9774 | 0.2602 | 21.13 |
| 1 | 53506847 | rs1118852 | G | A | 2 | 0.09544 | 0.09544 | 1.0941 | 1.0941 | 0.7123 | 0 |
| 1 | 53909648 | rs1461186 | G | A | 2 | 0.09559 | 0.3327 | 0.9425 | 0.9505 | 0.1568 | 50.11 |
| 1 | 54088450 | rs5978933 | C | T | 2 | 0.09566 | 0.09566 | 0.9124 | 0.9124 | 0.9713 | 0 |
| 1 | 53883374 | rs1418857 | G | T | 2 | 0.09681 | 0.1547 | 0.9636 | 0.9617 | 0.2255 | 31.92 |
| 1 | 53949203 | rs1288620 | C | T | 2 | 0.09686 | 0.2201 | 0.9453 | 0.9495 | 0.226 | 31.78 |
| 1 | 53970392 | rs2858428 | C | T | 2 | 0.09754 | 0.09754 | 1.0476 | 1.0476 | 0.6718 | 0 |
| 1 | 53965231 | rs1887403 | C | T | 2 | 0.09781 | 0.09781 | 1.0756 | 1.0756 | 0.3957 | 0 |
| 1 | 53506133 | rs7756267 | C | A | 2 | 0.09846 | 0.09846 | 1.0932 | 1.0932 | 0.7213 | 0 |
| 1 | 53878779 | rs6681936 | C | T | 2 | 0.09894 | 0.09894 | 0.9745 | 0.9745 | 0.5039 | 0 |
| 1 | 53961239 | rs1710858 | T | G | 2 | 0.09958 | 0.2432 | 0.9586 | 0.9523 | 0.1132 | 60.14 |
| 1 | 54109981 | rs1526908 | T | C | 2 | 0.09973 | 0.09973 | 1.1137 | 1.1137 | 0.4475 | 0 |
| 1 | 53948220 | rs943522 | C | T | 2 | 0.1005 | 0.1005 | 0.9302 | 0.9302 | 0.4373 | 0 |
| 1 | 54688516 | rs642990 | T | C | 2 | 0.1008 | 0.1008 | 1.0228 | 1.0228 | 0.4381 | 0 |
| 1 | 54036761 | rs2950247 | A | T | 2 | 0.1012 | 0.3308 | 0.9452 | 0.9483 | 0.1135 | 60.08 |
| 1 | 53827678 | rs7165499 | G | A | 2 | 0.1015 | 0.1015 | 0.9166 | 0.9166 | 0.5969 | 0 |
| 1 | 54698111 | rs643806 | T | A | 2 | 0.1018 | 0.1018 | 1.0228 | 1.0228 | 0.478 | 0 |
| 1 | 54209853 | rs1136427 | T | C | 2 | 0.1019 | 0.1019 | 0.9311 | 0.9311 | 0.3823 | 0 |
| 1 | 53956083 | rs6176844 | A | T | 2 | 0.1023 | 0.1023 | 1.0859 | 1.0859 | 0.4827 | 0 |
| 1 | 54181586 | rs1204077 | G | A | 2 | 0.1024 | 0.1024 | 0.9328 | 0.9328 | 0.3831 | 0 |

| | | | | | | | | | | | |
|---|---|---|---|---|---|---|---|---|---|---|---|
| 1 | 54449746 | rs7266410 | C | T | 2 | 0.1025 | 0.1025 | 0.966 | 0.966 | 0.6194 | 0 |
| 1 | 53960216 | rs1406257 | T | C | 2 | 0.1032 | 0.2389 | 0.9595 | 0.9542 | 0.126 | 57.29 |
| 1 | 54664921 | rs4926623 | A | G | 2 | 0.104 | 0.4494 | 0.9736 | 0.9781 | 0.0839 | 66.52 |
| 1 | 54614363 | rs1167485 | C | T | 2 | 0.1046 | 0.1046 | 0.9644 | 0.9644 | 0.9375 | 0 |
| 1 | 54500362 | rs1088883 | G | A | 2 | 0.1049 | 0.3116 | 0.9771 | 0.9794 | 0.1603 | 49.28 |
| 1 | 54636977 | rs1049317 | T | C | 2 | 0.1061 | 0.1061 | 0.9735 | 0.9735 | 0.34 | 0 |
| 1 | 53965248 | rs7666658 | G | C | 2 | 0.1062 | 0.2928 | 0.9585 | 0.9502 | 0.073 | 68.9 |
| 1 | 54603700 | rs913813 | A | G | 2 | 0.1065 | 0.1065 | 1.0311 | 1.0311 | 0.9772 | 0 |
| 1 | 53579402 | rs1288373 | C | T | 2 | 0.1076 | 0.1076 | 0.9553 | 0.9553 | 0.405 | 0 |
| 1 | 54620354 | rs7523092 | C | G | 2 | 0.1081 | 0.1081 | 1.0372 | 1.0372 | 0.912 | 0 |
| 1 | 54602838 | rs913812 | T | C | 2 | 0.1084 | 0.1084 | 1.0308 | 1.0308 | 0.845 | 0 |
| 1 | 53958815 | rs1120616 | A | C | 2 | 0.1091 | 0.1313 | 0.9609 | 0.9599 | 0.2805 | 14.13 |
| 1 | 53909906 | rs7408400 | G | A | 2 | 0.11 | 0.2699 | 0.9602 | 0.9538 | 0.1008 | 62.87 |
| 1 | 54607157 | rs671539 | G | A | 2 | 0.11 | 0.11 | 0.9705 | 0.9705 | 0.5012 | 0 |
| 1 | 54032759 | rs8027232 | G | T | 2 | 0.1103 | 0.3407 | 0.9719 | 0.9653 | 0.0434 | 75.48 |
| 1 | 53533333 | rs4926948 | A | G | 2 | 0.1111 | 0.1111 | 1.0378 | 1.0378 | 0.943 | 0 |
| 1 | 54513827 | rs914722 | T | A | 2 | 0.1112 | 0.3167 | 0.9777 | 0.9798 | 0.1566 | 50.16 |
| 1 | 53876149 | rs1342970 | G | T | 2 | 0.1121 | 0.1121 | 0.9751 | 0.9751 | 0.557 | 0 |
| 1 | 54613369 | rs1710991 | A | C | 2 | 0.1123 | 0.1123 | 0.9692 | 0.9692 | 0.8743 | 0 |
| 1 | 54507506 | rs7948100 | C | A | 2 | 0.1124 | 0.3938 | 1.0517 | 1.0454 | 0.1067 | 61.58 |
| 1 | 54019045 | rs502312 | A | G | 2 | 0.1125 | 0.1125 | 1.0294 | 1.0294 | 0.5057 | 0 |
| 1 | 54375701 | rs1120624 | C | T | 2 | 0.1125 | 0.1125 | 1.0227 | 1.0227 | 0.7398 | 0 |
| 1 | 53792241 | rs1144102 | C | A | 2 | 0.1127 | 0.1127 | 0.9305 | 0.9305 | 0.6786 | 0 |
| 1 | 54507000 | rs1141656 | G | A | 2 | 0.1128 | 0.3938 | 1.0517 | 1.0453 | 0.107 | 61.51 |
| 1 | 54621273 | rs628358 | G | A | 2 | 0.1128 | 0.1128 | 1.0244 | 1.0244 | 0.8078 | 0 |
| 1 | 54566310 | rs6694397 | G | A | 2 | 0.113 | 0.3567 | 0.9779 | 0.9805 | 0.1396 | 54.19 |
| 1 | 54034274 | rs6686640 | G | A | 2 | 0.1133 | 0.3398 | 0.9719 | 0.9648 | 0.0438 | 75.4 |
| 1 | 53856739 | rs7267513 | G | A | 2 | 0.1135 | 0.1135 | 1.0989 | 1.0989 | 0.8411 | 0 |
| 1 | 53887526 | rs6075435 | T | G | 2 | 0.1136 | 0.3101 | 0.9515 | 0.9566 | 0.174 | 45.88 |
| 1 | 54508231 | rs7507037 | C | A | 2 | 0.1147 | 0.394 | 1.0514 | 1.0451 | 0.1085 | 61.17 |
| 1 | 54629968 | rs1895927 | G | A | 2 | 0.1151 | 0.1151 | 1.1204 | 1.1204 | 0.8962 | 0 |
| 1 | 53878651 | rs6656566 | G | A | 2 | 0.1154 | 0.1154 | 0.9757 | 0.9757 | 0.4429 | 0 |
| 1 | 53988806 | rs2478979 | G | A | 2 | 0.1154 | 0.1154 | 0.9225 | 0.9225 | 0.5933 | 0 |
| 1 | 54094024 | rs2948056 | T | G | 2 | 0.1157 | 0.2766 | 1.1076 | 1.0942 | 0.2344 | 29.27 |
| 1 | 53587022 | rs1737731 | C | T | 2 | 0.1171 | 0.1171 | 0.923 | 0.923 | 0.9545 | 0 |
| 1 | 54507907 | rs7566527 | C | T | 2 | 0.1173 | 0.3999 | 1.0509 | 1.0447 | 0.1058 | 61.76 |
| 1 | 54608658 | rs1710990 | T | C | 2 | 0.1175 | 0.1175 | 1.0805 | 1.0805 | 0.6927 | 0 |
| 1 | 54345573 | rs7266236 | G | A | 2 | 0.1176 | 0.1306 | 1.0446 | 1.0455 | 0.2959 | 8.48 |
| 1 | 53884364 | rs7948129 | G | A | 2 | 0.1181 | 0.3305 | 0.952 | 0.9574 | 0.1673 | 47.56 |
| 1 | 53885808 | rs7450680 | A | G | 2 | 0.1182 | 0.3156 | 0.9521 | 0.9571 | 0.1753 | 45.56 |
| 1 | 53897517 | rs1203307 | G | A | 2 | 0.1184 | 0.1184 | 0.9705 | 0.9705 | 0.5432 | 0 |
| 1 | 53860103 | rs7267514 | C | A | 2 | 0.1191 | 0.1191 | 1.098 | 1.098 | 0.8736 | 0 |
| 1 | 54572175 | rs1204617 | T | C | 2 | 0.1195 | 0.3203 | 0.9781 | 0.9803 | 0.1684 | 47.29 |
| 1 | 54579848 | rs7463119 | G | A | 2 | 0.1198 | 0.5001 | 1.0525 | 1.0411 | 0.079 | 67.58 |
| 1 | 54310748 | rs7556609 | G | C | 2 | 0.1215 | 0.1215 | 0.9693 | 0.9693 | 0.4765 | 0 |
| 1 | 53639682 | rs7990163 | C | T | 2 | 0.1219 | 0.1219 | 0.9576 | 0.9576 | 0.8792 | 0 |
| 1 | 54465705 | rs7513522 | A | G | 2 | 0.122 | 0.1939 | 1.0219 | 1.0209 | 0.2612 | 20.77 |
| 1 | 53879927 | rs1710850 | A | C | 2 | 0.1227 | 0.1227 | 1.0302 | 1.0302 | 0.8807 | 0 |

| | | | | | | | | | | | |
|---|---|---|---|---|---|---|---|---|---|---|---|
| 1 | 54550595 | rs7446676 | C | A | 2 | 0.123 | 0.5113 | 1.0507 | 1.0412 | 0.0605 | 71.62 |
| 1 | 54127832 | rs5923876 | T | C | 2 | 0.1231 | 0.1231 | 1.0606 | 1.0606 | 0.6585 | 0 |
| 1 | 54598051 | rs1449454 | A | G | 2 | 0.1232 | 0.1232 | 0.962 | 0.962 | 0.6529 | 0 |
| 1 | 53907000 | rs1088879 | C | T | 2 | 0.1242 | 0.1242 | 0.9633 | 0.9633 | 0.3864 | 0 |
| 1 | 53815324 | rs5738095 | G | A | 2 | 0.1253 | 0.1253 | 0.863 | 0.863 | 0.9933 | 0 |
| 1 | 54616689 | rs9326027 | C | T | 2 | 0.1253 | 0.1581 | 1.0366 | 1.0381 | 0.2652 | 19.46 |
| 1 | 53482013 | rs1213250 | A | G | 2 | 0.1256 | 0.1256 | 0.9788 | 0.9788 | 0.8051 | 0 |
| 1 | 54444174 | rs1008406 | T | A | 2 | 0.1257 | 0.2125 | 1.0589 | 1.0569 | 0.2379 | 28.2 |
| 1 | 54470234 | rs1207515 | A | G | 2 | 0.1262 | 0.1661 | 1.0211 | 1.0207 | 0.2822 | 13.52 |
| 1 | 54528088 | rs7578915 | T | C | 2 | 0.1267 | 0.4906 | 1.0503 | 1.0417 | 0.0704 | 69.45 |
| 1 | 53889547 | rs7524189 | T | G | 2 | 0.1268 | 0.3443 | 0.9532 | 0.9586 | 0.1652 | 48.07 |
| 1 | 54482323 | rs7408512 | A | G | 2 | 0.127 | 0.127 | 1.0553 | 1.0553 | 0.5757 | 0 |
| 1 | 54686374 | rs612711 | T | C | 2 | 0.1271 | 0.1271 | 1.021 | 1.021 | 0.3789 | 0 |
| 1 | 53579182 | rs3766781 | G | A | 2 | 0.1272 | 0.1272 | 0.9765 | 0.9765 | 0.7215 | 0 |
| 1 | 54645313 | rs7528869 | G | A | 2 | 0.1275 | 0.1275 | 1.0232 | 1.0232 | 0.3747 | 0 |
| 1 | 53890718 | rs6686699 | C | T | 2 | 0.1276 | 0.3366 | 0.9532 | 0.9585 | 0.1713 | 46.56 |
| 1 | 54440345 | rs1214425 | A | C | 2 | 0.1276 | 0.1276 | 0.9751 | 0.9751 | 0.8102 | 0 |
| 1 | 54189769 | rs7524579 | G | T | 2 | 0.1282 | 0.1282 | 0.9373 | 0.9373 | 0.4048 | 0 |
| 1 | 54645262 | rs7520966 | T | C | 2 | 0.1282 | 0.1282 | 1.0231 | 1.0231 | 0.36 | 0 |
| 1 | 54442349 | rs7408880 | T | C | 2 | 0.1284 | 0.2086 | 1.0585 | 1.0566 | 0.2439 | 26.36 |
| 1 | 54442533 | rs7797718 | T | C | 2 | 0.1285 | 0.209 | 1.0584 | 1.0565 | 0.2432 | 26.56 |
| 1 | 54530630 | rs1146211 | C | A | 2 | 0.1286 | 0.4947 | 1.05 | 1.0414 | 0.0695 | 69.65 |
| 1 | 54142060 | rs7289678 | G | A | 2 | 0.1288 | 0.1288 | 1.055 | 1.055 | 0.7207 | 0 |
| 1 | 54441994 | rs7408880 | T | C | 2 | 0.1289 | 0.2093 | 1.0583 | 1.0565 | 0.2432 | 26.57 |
| 1 | 54616753 | rs1999175 | T | C | 2 | 0.1289 | 0.1476 | 1.0367 | 1.0376 | 0.2874 | 11.63 |
| 1 | 53866575 | rs1288592 | A | G | 2 | 0.1293 | 0.2251 | 0.9695 | 0.9637 | 0.1615 | 48.98 |
| 1 | 54441973 | rs7408450 | T | A | 2 | 0.1293 | 0.2111 | 1.0583 | 1.0564 | 0.2425 | 26.8 |
| 1 | 54552911 | rs1203340 | G | A | 2 | 0.1295 | 0.282 | 0.9787 | 0.9803 | 0.2011 | 38.82 |
| 1 | 54436691 | rs1120626 | A | G | 2 | 0.1296 | 0.3769 | 1.052 | 1.0472 | 0.1219 | 58.2 |
| 1 | 53510925 | rs7519017 | A | G | 2 | 0.1298 | 0.1298 | 1.0844 | 1.0844 | 0.5922 | 0 |
| 1 | 54469227 | rs635509 | A | G | 2 | 0.1298 | 0.1434 | 1.021 | 1.0209 | 0.3054 | 4.8 |
| 1 | 54024086 | rs6177027 | C | T | 2 | 0.1299 | 0.3576 | 0.9727 | 0.9646 | 0.0389 | 76.55 |
| 1 | 53529606 | rs1205820 | G | T | 2 | 0.13 | 0.144 | 0.8858 | 0.8858 | 0.3003 | 6.79 |
| 1 | 53597703 | rs1120610 | G | A | 2 | 0.1303 | 0.1926 | 1.0247 | 1.0239 | 0.2659 | 19.21 |
| 1 | 54540040 | rs1490119 | G | A | 2 | 0.1303 | 0.4981 | 1.0497 | 1.041 | 0.069 | 69.75 |
| 1 | 54661740 | rs946442 | T | C | 2 | 0.1303 | 0.1303 | 1.0812 | 1.0812 | 0.7456 | 0 |
| 1 | 54468689 | rs633315 | G | A | 2 | 0.1307 | 0.1503 | 1.021 | 1.0207 | 0.3003 | 6.78 |
| 1 | 54618435 | rs7528345 | A | T | 2 | 0.1307 | 0.1307 | 1.0229 | 1.0229 | 0.655 | 0 |
| 1 | 54601232 | rs2026110 | A | G | 2 | 0.1311 | 0.1311 | 1.0289 | 1.0289 | 0.8959 | 0 |
| 1 | 54668278 | rs1890566 | A | G | 2 | 0.1317 | 0.1317 | 0.9797 | 0.9797 | 0.3711 | 0 |
| 1 | 54275152 | rs1256284 | G | C | 2 | 0.1318 | 0.1318 | 1.0301 | 1.0301 | 0.6132 | 0 |
| 1 | 54469976 | rs585269 | C | T | 2 | 0.1325 | 0.1686 | 1.0209 | 1.0205 | 0.2873 | 11.67 |
| 1 | 54514675 | rs1412528 | T | C | 2 | 0.1325 | 0.1325 | 1.1195 | 1.1195 | 0.5194 | 0 |
| 1 | 53502810 | rs1487053 | A | G | 2 | 0.1328 | 0.1328 | 1.0861 | 1.0861 | 0.7687 | 0 |
| 1 | 54041766 | rs2177955 | T | C | 2 | 0.133 | 0.3738 | 1.0499 | 1.047 | 0.1121 | 60.38 |
| 1 | 54677182 | rs3766460 | G | A | 2 | 0.133 | 0.133 | 0.9798 | 0.9798 | 0.4788 | 0 |
| 1 | 54635805 | rs1088883 | T | G | 2 | 0.1331 | 0.1696 | 0.9228 | 0.9246 | 0.2912 | 10.23 |
| 1 | 54627830 | rs618465 | A | G | 2 | 0.134 | 0.134 | 0.9691 | 0.9691 | 0.8653 | 0 |

| | | | | | | | | | | | |
|---|---|---|---|---|---|---|---|---|---|---|---|
| 1 | 53850364 | rs7812285 | C | T | 2 | 0.1345 | 0.2835 | 1.0824 | 1.0741 | 0.2237 | 32.44 |
| 1 | 54470237 | rs666828 | T | C | 2 | 0.1345 | 0.1883 | 1.0205 | 1.02 | 0.2713 | 17.38 |
| 1 | 54471747 | rs7266411 | C | T | 2 | 0.1345 | 0.1898 | 0.9796 | 0.9802 | 0.2726 | 16.92 |
| 1 | 53884235 | rs1766636 | G | A | 2 | 0.1348 | 0.5622 | 0.9578 | 0.9684 | 0.0641 | 70.82 |
| 1 | 53831650 | rs7289946 | T | C | 2 | 0.1352 | 0.3071 | 0.8587 | 0.8509 | 0.1225 | 58.08 |
| 1 | 54608256 | rs7521143 | G | C | 2 | 0.1353 | 0.1353 | 1.0762 | 1.0762 | 0.7593 | 0 |
| 1 | 53960058 | rs1288612 | A | G | 2 | 0.1356 | 0.1629 | 1.0475 | 1.0467 | 0.2966 | 8.2 |
| 1 | 54468166 | rs620907 | T | G | 2 | 0.1359 | 0.1824 | 1.0206 | 1.0201 | 0.278 | 15.03 |
| 1 | 53908590 | rs1120615 | T | A | 2 | 0.1365 | 0.1365 | 0.9644 | 0.9644 | 0.3627 | 0 |
| 1 | 54033242 | rs1211954 | C | G | 2 | 0.1365 | 0.3706 | 0.9738 | 0.967 | 0.0408 | 76.11 |
| 1 | 54087425 | rs1120618 | T | A | 2 | 0.1366 | 0.1366 | 0.9177 | 0.9177 | 0.959 | 0 |
| 1 | 54125882 | rs7440221 | C | T | 2 | 0.1369 | 0.1369 | 0.8859 | 0.8859 | 0.7189 | 0 |
| 1 | 54470035 | rs1120627 | G | C | 2 | 0.1369 | 0.1844 | 1.0205 | 1.02 | 0.277 | 15.39 |
| 1 | 53515994 | rs1209552 | G | A | 2 | 0.137 | 0.1434 | 0.8882 | 0.8882 | 0.3096 | 3.15 |
| 1 | 54548012 | rs7965923 | G | T | 2 | 0.138 | 0.5164 | 1.0487 | 1.0398 | 0.0656 | 70.49 |
| 1 | 53611964 | rs1679934 | G | A | 2 | 0.1382 | 0.3468 | 0.9747 | 0.9776 | 0.1746 | 45.74 |
| 1 | 54088569 | rs7289672 | A | G | 2 | 0.1382 | 0.1382 | 0.9181 | 0.9181 | 0.9618 | 0 |
| 1 | 54436245 | rs1209478 | A | G | 2 | 0.1383 | 0.5214 | 1.0671 | 1.0562 | 0.0545 | 72.96 |
| 1 | 54346114 | rs1512938 | G | A | 2 | 0.1384 | 0.1926 | 1.0506 | 1.0559 | 0.2292 | 30.85 |
| 1 | 53500645 | rs1899705 | C | G | 2 | 0.1388 | 0.1388 | 1.0846 | 1.0846 | 0.7305 | 0 |
| 1 | 54369261 | rs7515148 | C | T | 2 | 0.1389 | 0.4175 | 0.9518 | 0.9597 | 0.1426 | 53.48 |
| 1 | 53500798 | rs1783802 | A | T | 2 | 0.139 | 0.139 | 1.0846 | 1.0846 | 0.7297 | 0 |
| 1 | 54098858 | rs1148313 | G | A | 2 | 0.1393 | 0.1393 | 0.8836 | 0.8836 | 0.8102 | 0 |
| 1 | 54089280 | rs6588484 | T | G | 2 | 0.1394 | 0.1394 | 0.9183 | 0.9183 | 0.9624 | 0 |
| 1 | 54560672 | rs6674672 | G | A | 2 | 0.1394 | 0.3654 | 0.9793 | 0.9816 | 0.1537 | 50.86 |
| 1 | 54483636 | rs7408512 | C | T | 2 | 0.1396 | 0.1396 | 1.0535 | 1.0535 | 0.5701 | 0 |
| 1 | 54223298 | rs1223981 | A | G | 2 | 0.1397 | 0.1397 | 1.0329 | 1.0329 | 0.4498 | 0 |
| 1 | 54470757 | rs679801 | T | C | 2 | 0.1397 | 0.1937 | 1.0206 | 1.02 | 0.2742 | 16.37 |
| 1 | 53893557 | rs1204833 | C | G | 2 | 0.1398 | 0.3503 | 0.9689 | 0.9648 | 0.0766 | 68.11 |
| 1 | 54476972 | rs5907129 | A | G | 2 | 0.1398 | 0.3182 | 1.049 | 1.0456 | 0.1719 | 46.42 |
| 1 | 54628704 | rs604206 | T | C | 2 | 0.1398 | 0.1398 | 1.0229 | 1.0229 | 0.7128 | 0 |
| 1 | 54483641 | rs7408513 | C | G | 2 | 0.1399 | 0.1399 | 1.0535 | 1.0535 | 0.5707 | 0 |
| 1 | 53890704 | rs6661567 | G | A | 2 | 0.1402 | 0.3901 | 0.9548 | 0.9609 | 0.1507 | 51.57 |
| 1 | 54466145 | rs1275821 | G | A | 2 | 0.1403 | 0.1702 | 1.0206 | 1.0203 | 0.2934 | 9.4 |
| 1 | 53564492 | rs7956080 | C | T | 2 | 0.1406 | 0.2243 | 0.9114 | 0.9169 | 0.2789 | 14.71 |
| 1 | 53764635 | rs2782500 | G | A | 2 | 0.1406 | 0.8068 | 1.0354 | 0.9787 | 0.0159 | 82.81 |
| 1 | 54468638 | rs676562 | C | G | 2 | 0.1407 | 0.1854 | 1.0206 | 1.0201 | 0.2818 | 13.66 |
| 1 | 54290619 | rs1256407 | G | C | 2 | 0.1409 | 0.1409 | 1.0294 | 1.0294 | 0.6777 | 0 |
| 1 | 53880369 | rs6685296 | T | C | 2 | 0.141 | 0.141 | 1.0288 | 1.0288 | 0.8026 | 0 |
| 1 | 54094177 | rs1120618 | C | G | 2 | 0.1415 | 0.1415 | 0.8847 | 0.8847 | 0.8457 | 0 |
| 1 | 53503202 | rs6013009 | G | C | 2 | 0.1417 | 0.1417 | 1.0839 | 1.0839 | 0.7319 | 0 |
| 1 | 53897228 | rs6165758 | A | T | 2 | 0.1417 | 0.3786 | 0.9545 | 0.9604 | 0.1594 | 49.5 |
| 1 | 54468947 | rs634532 | G | C | 2 | 0.1417 | 0.1896 | 1.0205 | 1.0199 | 0.279 | 14.65 |
| 1 | 54466956 | rs605182 | A | G | 2 | 0.1419 | 0.1842 | 1.0203 | 1.0199 | 0.2825 | 13.42 |
| 1 | 54482820 | rs1114915 | C | T | 2 | 0.1421 | 0.1421 | 1.0633 | 1.0633 | 0.5816 | 0 |
| 1 | 54467767 | rs662603 | T | C | 2 | 0.1425 | 0.1885 | 1.0206 | 1.02 | 0.2817 | 13.71 |
| 1 | 53502877 | rs1710770 | A | G | 2 | 0.1426 | 0.1426 | 1.0837 | 1.0837 | 0.734 | 0 |
| 1 | 54633967 | rs671431 | G | C | 2 | 0.1426 | 0.1426 | 1.0255 | 1.0255 | 0.691 | 0 |

| | | | | | | | | | | | |
|---|---|---|---|---|---|---|---|---|---|---|---|
| 1 | 54263944 | rs7266231 | T | G | 2 | 0.1428 | 0.1428 | 1.0292 | 1.0292 | 0.6201 | 0 |
| 1 | 54469948 | rs665817 | T | G | 2 | 0.1428 | 0.2062 | 1.0202 | 1.0195 | 0.2653 | 19.41 |
| 1 | 53598104 | rs4926955 | A | G | 2 | 0.1429 | 0.1904 | 1.0242 | 1.0236 | 0.2809 | 13.99 |
| 1 | 54027035 | rs4548383 | C | T | 2 | 0.1429 | 0.4008 | 0.9752 | 0.9695 | 0.0349 | 77.53 |
| 1 | 54284551 | rs7266232 | T | C | 2 | 0.1429 | 0.1429 | 1.0292 | 1.0292 | 0.6521 | 0 |
| 1 | 54286313 | rs1256628 | A | G | 2 | 0.1429 | 0.1429 | 1.0292 | 1.0292 | 0.6577 | 0 |
| 1 | 54020542 | rs1214126 | C | T | 2 | 0.1434 | 0.3927 | 0.9732 | 0.9633 | 0.0239 | 80.39 |
| 1 | 53783845 | rs1710832 | C | T | 2 | 0.1438 | 0.1438 | 0.9371 | 0.9371 | 0.8862 | 0 |
| 1 | 54126704 | rs4927024 | T | C | 2 | 0.1442 | 0.1442 | 1.0316 | 1.0316 | 0.3744 | 0 |
| 1 | 54468837 | rs677481 | A | C | 2 | 0.1444 | 0.1982 | 1.0202 | 1.0196 | 0.2736 | 16.57 |
| 1 | 53616806 | rs1088876 | G | A | 2 | 0.1447 | 0.3242 | 1.0258 | 1.0231 | 0.1946 | 40.56 |
| 1 | 54096471 | rs1120618 | G | A | 2 | 0.145 | 0.145 | 0.8857 | 0.8857 | 0.8386 | 0 |
| 1 | 53654444 | rs7538293 | T | C | 2 | 0.1455 | 0.1455 | 0.9074 | 0.9074 | 0.6454 | 0 |
| 1 | 54083312 | rs3006898 | T | C | 2 | 0.146 | 0.146 | 1.0226 | 1.0226 | 0.3881 | 0 |
| 1 | 54289826 | rs6068403 | C | T | 2 | 0.1461 | 0.1461 | 1.029 | 1.029 | 0.674 | 0 |
| 1 | 54283836 | rs1115780 | G | A | 2 | 0.1462 | 0.1462 | 1.029 | 1.029 | 0.6561 | 0 |
| 1 | 54114033 | rs7529819 | C | T | 2 | 0.1464 | 0.1464 | 1.0221 | 1.0221 | 0.4739 | 0 |
| 1 | 54637417 | rs682705 | G | A | 2 | 0.1465 | 0.1586 | 1.0224 | 1.0222 | 0.3081 | 3.72 |
| 1 | 53507732 | rs6682813 | G | A | 2 | 0.1468 | 0.211 | 0.8907 | 0.8909 | 0.2472 | 25.33 |
| 1 | 54097814 | rs1142066 | G | A | 2 | 0.1476 | 0.1476 | 0.8863 | 0.8863 | 0.8491 | 0 |
| 1 | 54176620 | rs1150637 | G | C | 2 | 0.1476 | 0.1476 | 0.9402 | 0.9402 | 0.8167 | 0 |
| 1 | 54462969 | rs7248062 | C | A | 2 | 0.1476 | 0.2708 | 1.02 | 1.0188 | 0.2234 | 32.55 |
| 1 | 54467342 | rs650134 | T | C | 2 | 0.1482 | 0.1978 | 1.0201 | 1.0196 | 0.2783 | 14.94 |
| 1 | 54468760 | rs633697 | A | G | 2 | 0.1487 | 0.1982 | 1.0199 | 1.0194 | 0.2771 | 15.33 |
| 1 | 54468321 | rs664848 | C | T | 2 | 0.1494 | 0.2104 | 1.0197 | 1.0191 | 0.2664 | 19.05 |
| 1 | 53608781 | rs1288363 | G | C | 2 | 0.1495 | 0.4592 | 0.9751 | 0.9794 | 0.1187 | 58.91 |
| 1 | 53589404 | rs6176814 | G | T | 2 | 0.1498 | 0.1498 | 0.9746 | 0.9746 | 0.6498 | 0 |
| 1 | 54507195 | rs3402831 | A | G | 2 | 0.1498 | 0.4159 | 0.9796 | 0.9823 | 0.133 | 55.69 |
| 1 | 54558958 | rs7840272 | G | T | 2 | 0.1501 | 0.4893 | 1.0472 | 1.0393 | 0.0878 | 65.68 |
| 1 | 53503282 | rs1274741 | C | T | 2 | 0.1502 | 0.1502 | 0.9799 | 0.9799 | 0.9917 | 0 |
| 1 | 54467026 | rs605272 | T | A | 2 | 0.1504 | 0.2139 | 1.0197 | 1.0191 | 0.2655 | 19.34 |
| 1 | 54476147 | rs1710965 | G | A | 2 | 0.1508 | 0.508 | 1.0483 | 1.0412 | 0.0668 | 70.25 |
| 1 | 53516383 | rs1088876 | C | A | 2 | 0.1514 | 0.18 | 0.8917 | 0.8918 | 0.2851 | 12.49 |
| 1 | 53950781 | rs1212554 | T | C | 2 | 0.1519 | 0.2322 | 1.0327 | 1.0379 | 0.1873 | 42.48 |
| 1 | 54674389 | rs7547103 | T | C | 2 | 0.152 | 0.152 | 0.9807 | 0.9807 | 0.4104 | 0 |
| 1 | 54076386 | rs6144910 | C | T | 2 | 0.1521 | 0.1521 | 0.9145 | 0.9145 | 0.6165 | 0 |
| 1 | 54677786 | rs1739254 | G | A | 2 | 0.1521 | 0.1521 | 0.9669 | 0.9669 | 0.4733 | 0 |
| 1 | 54248518 | rs4927036 | A | G | 2 | 0.1523 | 0.1523 | 1.0284 | 1.0284 | 0.6014 | 0 |
| 1 | 54514223 | rs4129477 | T | C | 2 | 0.1525 | 0.1525 | 0.9692 | 0.9692 | 0.3286 | 0 |
| 1 | 54258880 | rs2273139 | C | T | 2 | 0.1527 | 0.1527 | 1.0285 | 1.0285 | 0.6201 | 0 |
| 1 | 54285530 | rs4926614 | G | T | 2 | 0.1529 | 0.1529 | 1.0285 | 1.0285 | 0.6732 | 0 |
| 1 | 54575746 | rs4927055 | C | T | 2 | 0.1529 | 0.3885 | 0.9799 | 0.9823 | 0.1525 | 51.16 |
| 1 | 53602711 | rs2806274 | C | A | 2 | 0.1533 | 0.455 | 0.9753 | 0.9795 | 0.1244 | 57.65 |
| 1 | 54102148 | rs7533697 | T | C | 2 | 0.1535 | 0.1535 | 0.8996 | 0.8996 | 0.6111 | 0 |
| 1 | 54496344 | rs6652171 | C | T | 2 | 0.1535 | 0.4258 | 0.9799 | 0.9826 | 0.1283 | 56.76 |
| 1 | 54272063 | rs1181146 | A | T | 2 | 0.1536 | 0.1536 | 0.9606 | 0.9606 | 0.9535 | 0 |
| 1 | 54606342 | rs6588507 | C | T | 2 | 0.1537 | 0.1537 | 0.9743 | 0.9743 | 0.6052 | 0 |
| 1 | 53644535 | rs3125256 | A | T | 2 | 0.1538 | 0.1538 | 1.0218 | 1.0218 | 0.682 | 0 |

| | | | | | | | | | | | |
|---|---|---|---|---|---|---|---|---|---|---|---|
| 1 | 54498754 | rs3437044 | C | T | 2 | 0.1539 | 0.4337 | 0.9799 | 0.9827 | 0.1254 | 57.42 |
| 1 | 54240238 | rs7266230 | A | T | 2 | 0.1542 | 0.1542 | 1.0283 | 1.0283 | 0.5942 | 0 |
| 1 | 54461703 | rs637590 | A | G | 2 | 0.1544 | 0.2859 | 1.0194 | 1.0183 | 0.2132 | 35.48 |
| 1 | 54501826 | rs1204192 | G | C | 2 | 0.1544 | 0.4285 | 0.9799 | 0.9826 | 0.1273 | 57 |
| 1 | 54217999 | rs1159174 | C | T | 2 | 0.1545 | 0.1545 | 0.9352 | 0.9352 | 0.5788 | 0 |
| 1 | 54451130 | rs652208 | T | C | 2 | 0.1545 | 0.1545 | 1.0213 | 1.0213 | 0.7244 | 0 |
| 1 | 54363420 | rs1493943 | G | A | 2 | 0.1547 | 0.5725 | 0.9268 | 0.9484 | 0.1019 | 62.64 |
| 1 | 53772495 | rs1288490 | C | T | 2 | 0.1553 | 0.7705 | 1.091 | 1.0526 | 0.0052 | 87.17 |
| 1 | 54491225 | rs6705831 | C | T | 2 | 0.1554 | 0.4206 | 0.98 | 0.9826 | 0.1333 | 55.62 |
| 1 | 54469120 | rs635059 | G | C | 2 | 0.1558 | 0.2367 | 1.0194 | 1.0187 | 0.2519 | 23.81 |
| 1 | 54655563 | rs1120629 | C | T | 2 | 0.156 | 0.156 | 1.0195 | 1.0195 | 0.7193 | 0 |
| 1 | 54606403 | rs7638567 | T | G | 2 | 0.1563 | 0.1563 | 1.0739 | 1.0739 | 0.7623 | 0 |
| 1 | 54551831 | rs7898525 | T | A | 2 | 0.1566 | 0.5217 | 1.0465 | 1.0379 | 0.0756 | 68.32 |
| 1 | 53641358 | rs3125255 | T | C | 2 | 0.1568 | 0.1568 | 1.0216 | 1.0216 | 0.7216 | 0 |
| 1 | 54628328 | rs1128292 | G | A | 2 | 0.1577 | 0.1577 | 1.0319 | 1.0319 | 0.3933 | 0 |
| 1 | 54606343 | rs662524 | G | A | 2 | 0.1592 | 0.1592 | 1.0718 | 1.0718 | 0.8538 | 0 |
| 1 | 53501456 | rs1710769 | A | G | 2 | 0.1594 | 0.1594 | 1.0795 | 1.0795 | 0.6534 | 0 |
| 1 | 54340580 | rs7441600 | T | C | 2 | 0.1596 | 0.2393 | 0.9309 | 0.9347 | 0.2725 | 16.95 |
| 1 | 54682489 | rs747753 | C | G | 2 | 0.1597 | 0.1597 | 1.0267 | 1.0267 | 0.5083 | 0 |
| 1 | 54566190 | rs6694292 | G | A | 2 | 0.1599 | 0.4413 | 0.9804 | 0.9832 | 0.1255 | 57.41 |
| 1 | 53582306 | rs3766785 | A | T | 2 | 0.1602 | 0.1602 | 0.9769 | 0.9769 | 0.3567 | 0 |
| 1 | 54082647 | rs7289470 | C | G | 2 | 0.1605 | 0.1605 | 0.9216 | 0.9216 | 0.9905 | 0 |
| 1 | 54131414 | rs1158416 | C | G | 2 | 0.1611 | 0.1611 | 0.9695 | 0.9695 | 0.4329 | 0 |
| 1 | 53642449 | rs3102047 | A | G | 2 | 0.1613 | 0.1613 | 1.0214 | 1.0214 | 0.6509 | 0 |
| 1 | 53910662 | rs1208534 | G | A | 2 | 0.1614 | 0.3918 | 0.9566 | 0.9621 | 0.1656 | 47.99 |
| 1 | 54574645 | rs1145406 | A | C | 2 | 0.1618 | 0.5501 | 1.0457 | 1.0365 | 0.066 | 70.41 |
| 1 | 53890748 | rs1497373 | C | T | 2 | 0.1632 | 0.3633 | 0.9705 | 0.9663 | 0.0828 | 66.77 |
| 1 | 54608549 | rs4625227 | C | G | 2 | 0.1635 | 0.1635 | 0.9751 | 0.9751 | 0.6283 | 0 |
| 1 | 54104730 | rs7679788 | T | C | 2 | 0.1638 | 0.1638 | 0.9077 | 0.9077 | 0.9321 | 0 |
| 1 | 54111132 | rs4926610 | G | C | 2 | 0.1639 | 0.1639 | 1.0211 | 1.0211 | 0.4172 | 0 |
| 1 | 54473368 | rs928441 | C | A | 2 | 0.1639 | 0.2687 | 0.9808 | 0.9818 | 0.2388 | 27.93 |
| 1 | 54576469 | rs7408520 | A | T | 2 | 0.1641 | 0.1641 | 1.1073 | 1.1073 | 0.7739 | 0 |
| 1 | 53668954 | rs5932720 | G | A | 2 | 0.1643 | 0.4108 | 0.8896 | 0.897 | 0.1181 | 59.05 |
| 1 | 54302323 | rs1130163 | C | T | 2 | 0.1643 | 0.1643 | 1.0278 | 1.0278 | 0.5829 | 0 |
| 1 | 54251868 | rs5763635 | G | C | 2 | 0.1645 | 0.1645 | 1.0278 | 1.0278 | 0.641 | 0 |
| 1 | 54256713 | rs4927038 | C | T | 2 | 0.1646 | 0.1646 | 1.0277 | 1.0277 | 0.6374 | 0 |
| 1 | 53738400 | rs1380325 | T | C | 2 | 0.1647 | 0.1647 | 0.9537 | 0.9537 | 0.6458 | 0 |
| 1 | 54471157 | rs681617 | A | G | 2 | 0.1647 | 0.2878 | 1.0195 | 1.0181 | 0.2309 | 30.32 |
| 1 | 54615139 | rs682159 | G | A | 2 | 0.1648 | 0.2193 | 0.9787 | 0.9794 | 0.2785 | 14.85 |
| 1 | 54484513 | rs7528535 | C | T | 2 | 0.165 | 0.207 | 0.9777 | 0.9765 | 0.2539 | 23.18 |
| 1 | 54679756 | rs1203372 | T | A | 2 | 0.165 | 0.165 | 1.0265 | 1.0265 | 0.5272 | 0 |
| 1 | 53915383 | rs1121705 | T | A | 2 | 0.1652 | 0.1652 | 0.9598 | 0.9598 | 0.6069 | 0 |
| 1 | 53961283 | rs1129413 | G | A | 2 | 0.1653 | 0.1653 | 0.9456 | 0.9456 | 0.6108 | 0 |
| 1 | 54256324 | rs5786159 | G | A | 2 | 0.1653 | 0.1653 | 1.0276 | 1.0276 | 0.6388 | 0 |
| 1 | 54261043 | rs5619355 | C | T | 2 | 0.1657 | 0.1657 | 1.0276 | 1.0276 | 0.6544 | 0 |
| 1 | 54099804 | rs3006903 | G | A | 2 | 0.1658 | 0.1658 | 1.021 | 1.021 | 0.5122 | 0 |
| 1 | 54302624 | rs1123505 | T | C | 2 | 0.166 | 0.166 | 1.0277 | 1.0277 | 0.5897 | 0 |
| 1 | 53965176 | rs943520 | A | C | 2 | 0.1664 | 0.2931 | 1.044 | 1.0407 | 0.2326 | 29.82 |

| | | | | | | | | | | | |
|---|---|---|---|---|---|---|---|---|---|---|---|
| 1 | 54608910 | rs1768473 | G | A | 2 | 0.1673 | 0.1673 | 0.9754 | 0.9754 | 0.708 | 0 |
| 1 | 54598169 | rs630076 | C | G | 2 | 0.1677 | 0.1677 | 1.0533 | 1.0533 | 0.6636 | 0 |
| 1 | 54617813 | rs2066092 | G | A | 2 | 0.1677 | 0.1677 | 1.0209 | 1.0209 | 0.648 | 0 |
| 1 | 54473488 | rs928442 | C | G | 2 | 0.168 | 0.2536 | 0.9811 | 0.9819 | 0.2512 | 24.06 |
| 1 | 54617856 | rs2066091 | G | C | 2 | 0.1688 | 0.1688 | 1.0208 | 1.0208 | 0.7081 | 0 |
| 1 | 54251031 | rs4927037 | C | T | 2 | 0.1689 | 0.1689 | 1.0274 | 1.0274 | 0.631 | 0 |
| 1 | 54607617 | rs677732 | C | T | 2 | 0.1694 | 0.1694 | 0.9752 | 0.9752 | 0.7091 | 0 |
| 1 | 54655895 | rs1088883 | G | A | 2 | 0.1695 | 0.1695 | 1.0188 | 1.0188 | 0.711 | 0 |
| 1 | 53635975 | rs1427636 | C | T | 2 | 0.1697 | 0.1697 | 0.908 | 0.908 | 0.635 | 0 |
| 1 | 53889450 | rs1204424 | C | G | 2 | 0.1699 | 0.356 | 0.9712 | 0.9677 | 0.0989 | 63.27 |
| 1 | 54230086 | rs7266070 | G | A | 2 | 0.1708 | 0.1708 | 1.0272 | 1.0272 | 0.5525 | 0 |
| 1 | 53917887 | rs7557188 | G | C | 2 | 0.1709 | 0.4105 | 0.9575 | 0.9632 | 0.1619 | 48.89 |
| 1 | 54227790 | rs1256841 | C | G | 2 | 0.171 | 0.171 | 1.0273 | 1.0273 | 0.5305 | 0 |
| 1 | 53889978 | rs1203486 | T | A | 2 | 0.1713 | 0.3631 | 0.9713 | 0.9677 | 0.0941 | 64.33 |
| 1 | 54234580 | rs1753622 | C | A | 2 | 0.1718 | 0.1718 | 1.0272 | 1.0272 | 0.5598 | 0 |
| 1 | 54679319 | rs1120630 | C | T | 2 | 0.1718 | 0.1718 | 1.0261 | 1.0261 | 0.5158 | 0 |
| 1 | 54609280 | rs7593238 | G | A | 2 | 0.1721 | 0.7198 | 1.0699 | 1.044 | 0.0186 | 81.95 |
| 1 | 54302824 | rs5567315 | C | T | 2 | 0.1723 | 0.1723 | 1.0272 | 1.0272 | 0.5976 | 0 |
| 1 | 54594347 | rs673403 | T | C | 2 | 0.1723 | 0.1723 | 1.0188 | 1.0188 | 0.6631 | 0 |
| 1 | 54617256 | rs9436480 | A | G | 2 | 0.1723 | 0.2401 | 1.0331 | 1.0363 | 0.2147 | 35.03 |
| 1 | 54477125 | rs5698039 | G | A | 2 | 0.1726 | 0.3505 | 1.0456 | 1.042 | 0.1821 | 43.83 |
| 1 | 54029856 | rs2873096 | C | A | 2 | 0.1728 | 0.4309 | 0.9769 | 0.9711 | 0.0337 | 77.82 |
| 1 | 54108906 | rs1088880 | T | C | 2 | 0.1734 | 0.1734 | 1.0206 | 1.0206 | 0.4446 | 0 |
| 1 | 54473516 | rs928443 | A | G | 2 | 0.1735 | 0.257 | 0.9819 | 0.9824 | 0.2463 | 25.61 |
| 1 | 53961148 | rs4926599 | C | T | 2 | 0.1736 | 0.2519 | 1.0433 | 1.0413 | 0.264 | 19.85 |
| 1 | 53905267 | rs1167685 | C | A | 2 | 0.1739 | 0.3143 | 0.9651 | 0.9592 | 0.1233 | 57.9 |
| 1 | 54249533 | rs1256252 | C | T | 2 | 0.1739 | 0.1739 | 1.027 | 1.027 | 0.6166 | 0 |
| 1 | 54223184 | rs1223949 | G | A | 2 | 0.1742 | 0.1742 | 1.0315 | 1.0315 | 0.9415 | 0 |
| 1 | 54526545 | rs3519120 | C | T | 2 | 0.1747 | 0.4468 | 0.9809 | 0.9836 | 0.1331 | 55.67 |
| 1 | 53914058 | rs7884902 | G | A | 2 | 0.1749 | 0.412 | 0.9579 | 0.9635 | 0.1632 | 48.57 |
| 1 | 54571778 | rs7518719 | T | A | 2 | 0.1755 | 0.5558 | 1.0442 | 1.0352 | 0.0713 | 69.25 |
| 1 | 53646557 | rs3102044 | C | T | 2 | 0.1766 | 0.1766 | 1.0206 | 1.0206 | 0.6632 | 0 |
| 1 | 54648518 | rs1213907 | G | A | 2 | 0.1766 | 0.1766 | 0.9626 | 0.9626 | 0.954 | 0 |
| 1 | 53756149 | rs6176963 | T | G | 2 | 0.1767 | 0.1767 | 1.1173 | 1.1173 | 0.5547 | 0 |
| 1 | 53579109 | rs1710802 | C | T | 2 | 0.1771 | 0.4074 | 1.0464 | 1.0417 | 0.1468 | 52.5 |
| 1 | 54125949 | rs2948047 | C | T | 2 | 0.1771 | 0.1771 | 0.9711 | 0.9711 | 0.4457 | 0 |
| 1 | 54049955 | rs7755923 | T | C | 2 | 0.1778 | 0.3249 | 0.9609 | 0.9646 | 0.2309 | 30.34 |
| 1 | 54473548 | rs928444 | C | G | 2 | 0.1791 | 0.2791 | 0.9821 | 0.9827 | 0.2345 | 29.23 |
| 1 | 53945016 | rs1288623 | C | T | 2 | 0.1795 | 0.3592 | 0.9521 | 0.9586 | 0.2742 | 16.34 |
| 1 | 53620340 | rs2388 | C | T | 2 | 0.1806 | 0.3381 | 1.0238 | 1.0216 | 0.2122 | 35.74 |
| 1 | 53619912 | rs1288343 | A | G | 2 | 0.1807 | 0.3504 | 1.0238 | 1.0215 | 0.2064 | 37.35 |
| 1 | 54472778 | rs1023213 | T | G | 2 | 0.1816 | 0.3224 | 0.9817 | 0.9829 | 0.2146 | 35.07 |
| 1 | 54466383 | rs1274023 | T | C | 2 | 0.1817 | 0.3217 | 1.0184 | 1.0171 | 0.2159 | 34.7 |
| 1 | 54694750 | rs6177646 | A | G | 2 | 0.1821 | 0.2398 | 1.028 | 1.0314 | 0.2227 | 32.74 |
| 1 | 53960614 | rs6660494 | G | T | 2 | 0.1822 | 0.1822 | 1.0438 | 1.0438 | 0.3574 | 0 |
| 1 | 53792908 | rs1207509 | C | A | 2 | 0.1823 | 0.1823 | 0.8819 | 0.8819 | 0.7817 | 0 |
| 1 | 54469883 | rs665421 | T | C | 2 | 0.1832 | 0.3384 | 1.0184 | 1.0169 | 0.2078 | 36.98 |
| 1 | 54614384 | rs6671168 | G | C | 2 | 0.1837 | 0.1837 | 0.9599 | 0.9599 | 0.9298 | 0 |

| | | | | | | | | | | | |
|---|---|---|---|---|---|---|---|---|---|---|---|
| 1 | 54312060 | rs1503822 | C | T | 2 | 0.1846 | 0.4805 | 1.0285 | 1.0244 | 0.1126 | 60.28 |
| 1 | 54189668 | rs1204338 | G | A | 2 | 0.1848 | 0.1848 | 0.9733 | 0.9733 | 0.5506 | 0 |
| 1 | 53628061 | rs1288340 | G | A | 2 | 0.1855 | 0.3434 | 1.0232 | 1.0211 | 0.2137 | 35.32 |
| 1 | 54487588 | rs1114472 | C | G | 2 | 0.1866 | 0.1866 | 1.0488 | 1.0488 | 0.508 | 0 |
| 1 | 53887569 | rs1288605 | G | A | 2 | 0.1867 | 0.541 | 0.9618 | 0.9702 | 0.1054 | 61.86 |
| 1 | 54310719 | rs7556600 | G | A | 2 | 0.1867 | 0.1867 | 0.9587 | 0.9587 | 0.5187 | 0 |
| 1 | 53868794 | rs5579425 | T | C | 2 | 0.1873 | 0.1873 | 1.0761 | 1.0761 | 0.7306 | 0 |
| 1 | 54308972 | rs1738853 | A | T | 2 | 0.188 | 0.188 | 1.031 | 1.031 | 0.9752 | 0 |
| 1 | 53895802 | rs7833705 | T | C | 2 | 0.1884 | 0.334 | 0.9665 | 0.9605 | 0.1172 | 59.26 |
| 1 | 54366599 | rs4927043 | A | G | 2 | 0.1887 | 0.3584 | 0.9687 | 0.9716 | 0.2021 | 38.53 |
| 1 | 54463775 | rs605480 | G | T | 2 | 0.1892 | 0.4124 | 1.0182 | 1.016 | 0.1669 | 47.67 |
| 1 | 53628815 | rs1288338 | C | T | 2 | 0.1898 | 0.3429 | 1.023 | 1.0209 | 0.2164 | 34.57 |
| 1 | 54581869 | rs1337378 | T | C | 2 | 0.1904 | 0.1904 | 0.9824 | 0.9824 | 0.8361 | 0 |
| 1 | 54606804 | rs3766465 | C | T | 2 | 0.1905 | 0.1905 | 0.9765 | 0.9765 | 0.6443 | 0 |
| 1 | 53656293 | rs3502723 | C | T | 2 | 0.1907 | 0.3383 | 1.0189 | 1.0174 | 0.2129 | 35.54 |
| 1 | 53777739 | rs2788031 | C | T | 2 | 0.191 | 0.191 | 0.9639 | 0.9639 | 0.5334 | 0 |
| 1 | 53876381 | rs953625 | A | G | 2 | 0.191 | 0.191 | 0.9822 | 0.9822 | 0.7008 | 0 |
| 1 | 54369674 | rs1203357 | G | C | 2 | 0.1911 | 0.567 | 0.9564 | 0.9681 | 0.1168 | 59.35 |
| 1 | 53644093 | rs1088877 | T | A | 2 | 0.1912 | 0.1912 | 1.0196 | 1.0196 | 0.8293 | 0 |
| 1 | 53644036 | rs1088877 | T | C | 2 | 0.1913 | 0.1913 | 0.9798 | 0.9798 | 0.7378 | 0 |
| 1 | 54037328 | rs1438538 | C | A | 2 | 0.1916 | 0.1916 | 0.932 | 0.932 | 0.7151 | 0 |
| 1 | 54240489 | rs7266230 | A | C | 2 | 0.1919 | 0.1919 | 1.0259 | 1.0259 | 0.6632 | 0 |
| 1 | 54588534 | rs9436934 | C | T | 2 | 0.1922 | 0.1922 | 1.0181 | 1.0181 | 0.601 | 0 |
| 1 | 54191958 | rs797908 | C | T | 2 | 0.1923 | 0.1923 | 0.9737 | 0.9737 | 0.5719 | 0 |
| 1 | 54487384 | rs7290438 | C | T | 2 | 0.1925 | 0.1925 | 1.048 | 1.048 | 0.4821 | 0 |
| 1 | 54246860 | rs3460526 | T | C | 2 | 0.1927 | 0.1927 | 0.957 | 0.957 | 0.4263 | 0 |
| 1 | 53902553 | rs7744217 | A | T | 2 | 0.1929 | 0.3141 | 0.9666 | 0.9612 | 0.1439 | 53.19 |
| 1 | 54090242 | rs2948058 | C | T | 2 | 0.1932 | 0.1932 | 1.0198 | 1.0198 | 0.4791 | 0 |
| 1 | 54615337 | rs585715 | C | T | 2 | 0.194 | 0.194 | 1.0488 | 1.0488 | 0.6471 | 0 |
| 1 | 53519895 | rs1120608 | A | T | 2 | 0.1944 | 0.1944 | 0.903 | 0.903 | 0.4055 | 0 |
| 1 | 53885577 | rs7642135 | A | G | 2 | 0.1944 | 0.3237 | 0.9668 | 0.9613 | 0.1344 | 55.38 |
| 1 | 53792975 | rs1206097 | T | C | 2 | 0.1945 | 0.1945 | 0.8846 | 0.8846 | 0.8051 | 0 |
| 1 | 54466568 | rs1507200 | C | T | 2 | 0.1951 | 0.1951 | 0.9279 | 0.9279 | 0.6862 | 0 |
| 1 | 53922026 | rs7810310 | A | G | 2 | 0.1954 | 0.344 | 0.9666 | 0.9604 | 0.1133 | 60.13 |
| 1 | 53828529 | rs5817314 | A | G | 2 | 0.1961 | 0.1961 | 1.1881 | 1.1881 | 0.6452 | 0 |
| 1 | 53899001 | rs1448412 | C | T | 2 | 0.1961 | 0.3122 | 0.9668 | 0.9617 | 0.1502 | 51.69 |
| 1 | 54005913 | rs7554233 | T | G | 2 | 0.1964 | 0.2475 | 1.0183 | 1.0197 | 0.2391 | 27.85 |
| 1 | 54320337 | rs1120622 | A | G | 2 | 0.1967 | 0.1967 | 0.9599 | 0.9599 | 0.5807 | 0 |
| 1 | 53824304 | rs1130370 | G | T | 2 | 0.1976 | 0.3444 | 0.8673 | 0.863 | 0.1591 | 49.57 |
| 1 | 54596219 | rs654403 | T | C | 2 | 0.1977 | 0.1977 | 1.0178 | 1.0178 | 0.643 | 0 |
| 1 | 53965546 | rs943519 | C | T | 2 | 0.1978 | 0.2911 | 1.041 | 1.0387 | 0.2567 | 22.28 |
| 1 | 54656765 | rs1207972 | C | T | 2 | 0.1978 | 0.1978 | 1.0763 | 1.0763 | 0.3654 | 0 |
| 1 | 53678227 | rs7725939 | A | G | 2 | 0.1982 | 0.2039 | 1.0563 | 1.0567 | 0.3091 | 3.35 |
| 1 | 54561290 | rs1152303 | C | T | 2 | 0.1989 | 0.2859 | 1.0605 | 1.0706 | 0.179 | 44.62 |
| 1 | 53621394 | rs1288345 | A | C | 2 | 0.2 | 0.4337 | 1.0222 | 1.0191 | 0.1685 | 47.28 |
| 1 | 54198205 | rs7938098 | C | T | 2 | 0.2001 | 0.2001 | 0.9678 | 0.9678 | 0.9224 | 0 |
| 1 | 54605107 | rs636426 | T | C | 2 | 0.2004 | 0.2004 | 1.0642 | 1.0642 | 0.8145 | 0 |
| 1 | 54015832 | rs1208185 | A | C | 2 | 0.2006 | 0.2006 | 1.018 | 1.018 | 0.3454 | 0 |

| | | | | | | | | | | | |
|---|---|---|---|---|---|---|---|---|---|---|---|
| 1 | 53596099 | rs7408278 | C | T | 2 | 0.2009 | 0.3405 | 1.0367 | 1.0472 | 0.1024 | 62.52 |
| 1 | 53868230 | rs7267514 | G | A | 2 | 0.2016 | 0.2016 | 1.0735 | 1.0735 | 0.7132 | 0 |
| 1 | 54688787 | rs1078897 | C | G | 2 | 0.2018 | 0.2018 | 1.0189 | 1.0189 | 0.4373 | 0 |
| 1 | 54689461 | rs1088884 | T | C | 2 | 0.202 | 0.202 | 1.0191 | 1.0191 | 0.511 | 0 |
| 1 | 53918739 | rs1151053 | T | C | 2 | 0.2022 | 0.3302 | 0.9672 | 0.9618 | 0.1368 | 54.83 |
| 1 | 54656651 | rs4927066 | C | G | 2 | 0.2029 | 0.2029 | 1.0175 | 1.0175 | 0.6029 | 0 |
| 1 | 54310214 | rs1120621 | A | G | 2 | 0.2044 | 0.2044 | 0.9605 | 0.9605 | 0.4204 | 0 |
| 1 | 53947018 | rs1140881 | G | A | 2 | 0.2046 | 0.3508 | 0.9671 | 0.9604 | 0.1113 | 60.57 |
| 1 | 53638446 | rs1157721 | C | T | 2 | 0.2049 | 0.2718 | 0.9359 | 0.9312 | 0.2215 | 33.08 |
| 1 | 53589801 | rs1679971 | A | G | 2 | 0.2055 | 0.2055 | 0.9776 | 0.9776 | 0.9978 | 0 |
| 1 | 53829540 | rs7408397 | T | G | 2 | 0.2057 | 0.2057 | 1.1936 | 1.1936 | 0.5979 | 0 |
| 1 | 53654975 | rs7752222 | G | A | 2 | 0.2059 | 0.2059 | 0.9185 | 0.9185 | 0.9665 | 0 |
| 1 | 53796206 | rs1415142 | A | G | 2 | 0.2059 | 0.2059 | 0.886 | 0.886 | 0.8882 | 0 |
| 1 | 54469109 | rs635045 | A | G | 2 | 0.2061 | 0.3893 | 1.0176 | 1.0159 | 0.1919 | 41.28 |
| 1 | 54492546 | rs6000408 | C | T | 2 | 0.2066 | 0.2066 | 1.0799 | 1.0799 | 0.7098 | 0 |
| 1 | 54289020 | rs1048946 | C | T | 2 | 0.2072 | 0.2072 | 1.0289 | 1.0289 | 0.8649 | 0 |
| 1 | 53948905 | rs1737195 | T | C | 2 | 0.2073 | 0.3537 | 0.9673 | 0.9606 | 0.1105 | 60.74 |
| 1 | 53954869 | rs7897924 | G | C | 2 | 0.2074 | 0.3549 | 0.9673 | 0.9605 | 0.1093 | 61 |
| 1 | 54551875 | rs7516791 | G | T | 2 | 0.2084 | 0.3097 | 0.9828 | 0.9837 | 0.2475 | 25.23 |
| 1 | 53577701 | rs922682 | C | T | 2 | 0.2087 | 0.2087 | 0.9339 | 0.9339 | 0.427 | 0 |
| 1 | 54609652 | rs599597 | C | T | 2 | 0.2094 | 0.2094 | 0.975 | 0.975 | 0.7895 | 0 |
| 1 | 54342517 | rs1154486 | T | C | 2 | 0.2098 | 0.3871 | 1.0312 | 1.0282 | 0.1963 | 40.12 |
| 1 | 53868228 | rs7267514 | A | G | 2 | 0.2102 | 0.2102 | 1.0721 | 1.0721 | 0.7295 | 0 |
| 1 | 54081664 | rs1120617 | A | G | 2 | 0.2104 | 0.2104 | 0.9805 | 0.9805 | 0.5976 | 0 |
| 1 | 54264780 | rs6177480 | C | A | 2 | 0.2105 | 0.2105 | 0.9588 | 0.9588 | 0.4038 | 0 |
| 1 | 53881254 | rs6683597 | A | G | 2 | 0.2106 | 0.2106 | 0.9832 | 0.9832 | 0.9619 | 0 |
| 1 | 53949869 | rs1154546 | T | C | 2 | 0.2112 | 0.3438 | 0.9676 | 0.9616 | 0.1273 | 56.98 |
| 1 | 54167478 | rs1126767 | C | T | 2 | 0.2112 | 0.2112 | 1.0432 | 1.0432 | 0.9847 | 0 |
| 1 | 54480410 | rs1180550 | T | A | 2 | 0.2113 | 0.2113 | 1.0443 | 1.0443 | 0.3777 | 0 |
| 1 | 54605177 | rs636533 | G | A | 2 | 0.212 | 0.212 | 1.0625 | 1.0625 | 0.8367 | 0 |
| 1 | 53951565 | rs1288618 | C | T | 2 | 0.2121 | 0.4371 | 0.9615 | 0.9666 | 0.1756 | 45.48 |
| 1 | 54563239 | rs6672905 | A | T | 2 | 0.2127 | 0.4485 | 0.9827 | 0.9849 | 0.1576 | 49.93 |
| 1 | 53586010 | rs7803088 | C | T | 2 | 0.213 | 0.3196 | 1.0424 | 1.0559 | 0.1288 | 56.66 |
| 1 | 53899019 | rs1766641 | C | G | 2 | 0.2134 | 0.2134 | 1.0255 | 1.0255 | 0.9468 | 0 |
| 1 | 54629940 | rs1125337 | T | C | 2 | 0.2134 | 0.2134 | 1.028 | 1.028 | 0.455 | 0 |
| 1 | 53930360 | rs6173884 | G | A | 2 | 0.2141 | 0.3535 | 0.9679 | 0.9619 | 0.121 | 58.41 |
| 1 | 54327983 | rs1274410 | A | G | 2 | 0.2146 | 0.2146 | 0.9614 | 0.9614 | 0.5735 | 0 |
| 1 | 53893551 | rs1159052 | C | T | 2 | 0.215 | 0.215 | 1.0264 | 1.0264 | 0.8628 | 0 |
| 1 | 54003748 | rs1273993 | G | A | 2 | 0.2154 | 0.2154 | 1.0175 | 1.0175 | 0.3843 | 0 |
| 1 | 54466789 | rs1120627 | C | G | 2 | 0.2154 | 0.4315 | 1.0171 | 1.0151 | 0.1724 | 46.3 |
| 1 | 53657684 | rs6176959 | T | G | 2 | 0.2156 | 0.2156 | 1.0936 | 1.0936 | 0.8266 | 0 |
| 1 | 54362193 | rs2268185 | T | G | 2 | 0.216 | 0.216 | 0.9613 | 0.9613 | 0.4495 | 0 |
| 1 | 53926750 | rs7821931 | C | G | 2 | 0.2163 | 0.3539 | 0.9681 | 0.9622 | 0.1231 | 57.94 |
| 1 | 54323368 | rs1120622 | T | C | 2 | 0.2172 | 0.2172 | 0.9617 | 0.9617 | 0.5962 | 0 |
| 1 | 54308934 | rs7408443 | G | T | 2 | 0.2173 | 0.4964 | 1.0985 | 1.0955 | 0.0781 | 67.79 |
| 1 | 54697515 | rs1399413 | G | A | 2 | 0.2173 | 0.2173 | 0.91 | 0.91 | 0.3329 | 0 |
| 1 | 54588882 | rs7528864 | A | G | 2 | 0.2175 | 0.2175 | 1.0171 | 1.0171 | 0.6049 | 0 |
| 1 | 54689643 | rs1078897 | A | G | 2 | 0.2175 | 0.2175 | 1.0185 | 1.0185 | 0.5323 | 0 |

| | | | | | | | | | | | |
|---|---|---|---|---|---|---|---|---|---|---|---|
| 1 | 54323891 | rs1202270 | T | C | 2 | 0.2176 | 0.2176 | 0.9617 | 0.9617 | 0.5955 | 0 |
| 1 | 53776929 | rs7289537 | C | T | 2 | 0.2177 | 0.2177 | 0.9671 | 0.9671 | 0.4841 | 0 |
| 1 | 54324061 | rs1120622 | A | G | 2 | 0.2181 | 0.2181 | 0.9617 | 0.9617 | 0.5945 | 0 |
| 1 | 53972049 | rs2296761 | A | C | 2 | 0.2182 | 0.2182 | 0.9474 | 0.9474 | 0.7727 | 0 |
| 1 | 53937115 | rs7855232 | A | T | 2 | 0.2184 | 0.3588 | 0.9682 | 0.9621 | 0.1188 | 58.89 |
| 1 | 54167387 | rs7588551 | T | C | 2 | 0.2185 | 0.2185 | 0.9742 | 0.9742 | 0.9826 | 0 |
| 1 | 54363076 | rs1158216 | G | A | 2 | 0.2186 | 0.2186 | 0.9616 | 0.9616 | 0.4467 | 0 |
| 1 | 54614409 | rs581554 | C | T | 2 | 0.2191 | 0.2191 | 0.9808 | 0.9808 | 0.3941 | 0 |
| 1 | 54363750 | rs3468408 | C | T | 2 | 0.2198 | 0.2198 | 0.9618 | 0.9618 | 0.4442 | 0 |
| 1 | 53621345 | rs1288344 | G | T | 2 | 0.2204 | 0.4418 | 1.0213 | 1.0184 | 0.1791 | 44.61 |
| 1 | 54364755 | rs7521654 | G | A | 2 | 0.2206 | 0.2206 | 0.9617 | 0.9617 | 0.4461 | 0 |
| 1 | 54163836 | rs1158263 | G | C | 2 | 0.2207 | 0.2207 | 0.9743 | 0.9743 | 0.9864 | 0 |
| 1 | 54325688 | rs1120623 | T | C | 2 | 0.2211 | 0.2211 | 0.9619 | 0.9619 | 0.5938 | 0 |
| 1 | 54324598 | rs6177482 | C | T | 2 | 0.2213 | 0.2213 | 0.9601 | 0.9601 | 0.6016 | 0 |
| 1 | 53952714 | rs7586836 | G | A | 2 | 0.2214 | 0.3571 | 0.9683 | 0.9621 | 0.1217 | 58.24 |
| 1 | 54614145 | rs6696001 | C | T | 2 | 0.222 | 0.222 | 0.9635 | 0.9635 | 0.9044 | 0 |
| 1 | 53939697 | rs7757662 | T | A | 2 | 0.2227 | 0.3604 | 0.9685 | 0.9624 | 0.1209 | 58.44 |
| 1 | 54591265 | rs612297 | A | G | 2 | 0.2231 | 0.2231 | 1.0169 | 1.0169 | 0.6086 | 0 |
| 1 | 54190045 | rs7547962 | C | T | 2 | 0.2234 | 0.2234 | 0.9759 | 0.9759 | 0.3199 | 0 |
| 1 | 53977292 | rs527014 | C | T | 2 | 0.2235 | 0.2235 | 0.9537 | 0.9537 | 0.8332 | 0 |
| 1 | 53866131 | rs7267514 | T | A | 2 | 0.2239 | 0.2239 | 1.0696 | 1.0696 | 0.6823 | 0 |
| 1 | 53640580 | rs3102046 | T | C | 2 | 0.225 | 0.225 | 1.0188 | 1.0188 | 0.9305 | 0 |
| 1 | 54144459 | rs7538181 | G | A | 2 | 0.225 | 0.225 | 1.0461 | 1.0461 | 0.7701 | 0 |
| 1 | 54311234 | rs1738857 | C | T | 2 | 0.2251 | 0.5598 | 1.0258 | 1.0215 | 0.0874 | 65.77 |
| 1 | 54365194 | rs1449994 | G | A | 2 | 0.2253 | 0.2253 | 1.0792 | 1.0792 | 0.6001 | 0 |
| 1 | 54445959 | rs1209406 | G | A | 2 | 0.2257 | 0.64 | 1.0438 | 1.0349 | 0.0406 | 76.16 |
| 1 | 54093593 | rs3013744 | A | G | 2 | 0.2261 | 0.2261 | 1.0183 | 1.0183 | 0.4778 | 0 |
| 1 | 54555405 | rs1981039 | T | A | 2 | 0.2261 | 0.3626 | 0.9835 | 0.9847 | 0.2233 | 32.56 |
| 1 | 54591269 | rs688404 | C | T | 2 | 0.2264 | 0.2264 | 1.0168 | 1.0168 | 0.6135 | 0 |
| 1 | 53596130 | rs7267087 | G | A | 2 | 0.2267 | 0.2267 | 0.9756 | 0.9756 | 0.9058 | 0 |
| 1 | 54317485 | rs1127931 | G | A | 2 | 0.2267 | 0.2267 | 0.962 | 0.962 | 0.5154 | 0 |
| 1 | 54649636 | rs1469147 | G | A | 2 | 0.2285 | 0.2285 | 1.0701 | 1.0701 | 0.6815 | 0 |
| 1 | 54532687 | rs6177759 | A | G | 2 | 0.2286 | 0.2286 | 0.9657 | 0.9657 | 0.9615 | 0 |
| 1 | 53880664 | rs6588479 | A | G | 2 | 0.2287 | 0.2287 | 0.9838 | 0.9838 | 0.7855 | 0 |
| 1 | 53574123 | rs1769289 | C | T | 2 | 0.229 | 0.229 | 0.9283 | 0.9283 | 0.936 | 0 |
| 1 | 54443432 | rs1112361 | A | G | 2 | 0.2291 | 0.6 | 1.044 | 1.0366 | 0.0585 | 72.07 |
| 1 | 54693425 | rs1158900 | A | G | 2 | 0.2291 | 0.2508 | 1.0244 | 1.0256 | 0.2816 | 13.75 |
| 1 | 54187449 | rs1157925 | C | T | 2 | 0.2292 | 0.242 | 0.9762 | 0.976 | 0.3005 | 6.73 |
| 1 | 53641911 | rs7531337 | C | A | 2 | 0.2295 | 0.2295 | 0.982 | 0.982 | 0.9455 | 0 |
| 1 | 54591322 | rs612368 | G | A | 2 | 0.2296 | 0.2296 | 1.0165 | 1.0165 | 0.786 | 0 |
| 1 | 54608746 | rs1768474 | C | T | 2 | 0.2301 | 0.2301 | 0.9787 | 0.9787 | 0.8828 | 0 |
| 1 | 54133184 | rs7965589 | G | C | 2 | 0.2305 | 0.2305 | 1.049 | 1.049 | 0.9744 | 0 |
| 1 | 54539339 | rs6674720 | C | G | 2 | 0.2307 | 0.3666 | 0.9836 | 0.9848 | 0.2249 | 32.12 |
| 1 | 53646540 | rs3102045 | T | A | 2 | 0.2309 | 0.2309 | 1.0185 | 1.0185 | 0.8957 | 0 |
| 1 | 53585254 | rs6176811 | G | A | 2 | 0.2313 | 0.3476 | 1.0402 | 1.0572 | 0.0998 | 63.09 |
| 1 | 54094104 | rs2950266 | G | C | 2 | 0.2314 | 0.2314 | 1.0181 | 1.0181 | 0.4708 | 0 |
| 1 | 54276225 | rs1181156 | A | T | 2 | 0.2314 | 0.2314 | 0.9667 | 0.9667 | 0.6945 | 0 |
| 1 | 54243532 | rs7266231 | G | A | 2 | 0.2316 | 0.2316 | 1.0238 | 1.0238 | 0.4552 | 0 |

| | | | | | | | | | | | |
|---|---|---|---|---|---|---|---|---|---|---|---|
| 1 | 54462254 | rs6698321 | T | C | 2 | 0.2317 | 0.2317 | 1.03 | 1.03 | 0.6675 | 0 |
| 1 | 54312428 | rs1275397 | C | T | 2 | 0.2318 | 0.2318 | 0.9627 | 0.9627 | 0.5956 | 0 |
| 1 | 53883775 | rs7761176 | C | A | 2 | 0.2321 | 0.2428 | 1.037 | 1.0377 | 0.301 | 6.53 |
| 1 | 54005523 | rs1275023 | G | A | 2 | 0.2323 | 0.2913 | 1.0169 | 1.0185 | 0.2248 | 32.12 |
| 1 | 54553207 | rs9651202 | T | C | 2 | 0.233 | 0.367 | 0.9837 | 0.9849 | 0.2261 | 31.75 |
| 1 | 53931213 | rs1342971 | C | A | 2 | 0.2336 | 0.3782 | 0.9692 | 0.9628 | 0.1115 | 60.53 |
| 1 | 54319776 | rs1158971 | T | C | 2 | 0.2346 | 0.2346 | 0.9634 | 0.9634 | 0.5548 | 0 |
| 1 | 54639004 | rs1180092 | C | T | 2 | 0.2348 | 0.3693 | 1.0693 | 1.0615 | 0.2759 | 15.75 |
| 1 | 54331152 | rs1202151 | G | A | 2 | 0.2352 | 0.2352 | 0.963 | 0.963 | 0.5859 | 0 |
| 1 | 53881081 | rs6686055 | T | C | 2 | 0.2354 | 0.2354 | 0.984 | 0.984 | 0.7471 | 0 |
| 1 | 54390466 | rs1272873 | C | T | 2 | 0.2359 | 0.2359 | 0.9795 | 0.9795 | 0.8152 | 0 |
| 1 | 53941008 | rs1288643 | A | G | 2 | 0.2365 | 0.4516 | 0.9636 | 0.9683 | 0.182 | 43.87 |
| 1 | 54292026 | rs1181186 | G | A | 2 | 0.2365 | 0.2365 | 0.9671 | 0.9671 | 0.6822 | 0 |
| 1 | 54279889 | rs1434913 | G | A | 2 | 0.2366 | 0.2366 | 0.961 | 0.961 | 0.4502 | 0 |
| 1 | 54527647 | rs1212632 | A | G | 2 | 0.2369 | 0.2369 | 0.9796 | 0.9796 | 0.5583 | 0 |
| 1 | 54686464 | rs613158 | C | T | 2 | 0.2375 | 0.4832 | 1.0164 | 1.0246 | 0.0155 | 82.93 |
| 1 | 53672688 | rs1072706 | T | C | 2 | 0.2377 | 0.2377 | 1.0172 | 1.0172 | 0.5512 | 0 |
| 1 | 53792335 | rs3737980 | G | A | 2 | 0.2381 | 0.2381 | 0.8921 | 0.8921 | 0.7275 | 0 |
| 1 | 54444816 | rs1206061 | A | G | 2 | 0.2381 | 0.6213 | 1.0433 | 1.0351 | 0.0548 | 72.88 |
| 1 | 54410017 | rs7290248 | T | A | 2 | 0.2382 | 0.2382 | 1.0441 | 1.0441 | 0.468 | 0 |
| 1 | 54561916 | rs7555099 | C | T | 2 | 0.2384 | 0.4073 | 0.9839 | 0.9854 | 0.2041 | 38 |
| 1 | 53971807 | rs1710864 | C | T | 2 | 0.2387 | 0.2387 | 0.9453 | 0.9453 | 0.5245 | 0 |
| 1 | 54445241 | rs1120626 | T | C | 2 | 0.2388 | 0.6428 | 1.0425 | 1.0343 | 0.0418 | 75.86 |
| 1 | 54630794 | rs1126690 | G | T | 2 | 0.2389 | 0.2389 | 0.98 | 0.98 | 0.7697 | 0 |
| 1 | 53946661 | rs1203146 | T | A | 2 | 0.2392 | 0.4262 | 0.9611 | 0.962 | 0.1486 | 52.07 |
| 1 | 53949573 | rs7551111 | A | C | 2 | 0.2393 | 0.2958 | 1.0829 | 1.1079 | 0.2043 | 37.93 |
| 1 | 53511229 | rs1208848 | G | A | 2 | 0.2396 | 0.2396 | 1.0397 | 1.0397 | 0.8113 | 0 |
| 1 | 54643655 | rs7730008 | C | T | 2 | 0.2399 | 0.2399 | 0.967 | 0.967 | 0.8948 | 0 |
| 1 | 53502765 | rs1871750 | T | C | 2 | 0.24 | 0.24 | 1.0251 | 1.0251 | 0.456 | 0 |
| 1 | 53794611 | rs1897099 | G | A | 2 | 0.2403 | 0.2403 | 0.9331 | 0.9331 | 0.5546 | 0 |
| 1 | 54595215 | rs1158648 | C | T | 2 | 0.2407 | 0.2407 | 0.9839 | 0.9839 | 0.5046 | 0 |
| 1 | 54228278 | rs1274641 | C | T | 2 | 0.2417 | 0.2417 | 0.9615 | 0.9615 | 0.5398 | 0 |
| 1 | 53964385 | rs1088879 | C | T | 2 | 0.2418 | 0.2963 | 1.0178 | 1.0192 | 0.2362 | 28.74 |
| 1 | 54673097 | rs1088883 | C | T | 2 | 0.2419 | 0.2419 | 1.0219 | 1.0219 | 0.3715 | 0 |
| 1 | 53873555 | rs1288587 | T | C | 2 | 0.2421 | 0.2421 | 0.984 | 0.984 | 0.568 | 0 |
| 1 | 54589719 | rs671083 | G | C | 2 | 0.2421 | 0.2421 | 1.0161 | 1.0161 | 0.5481 | 0 |
| 1 | 54356647 | rs3586932 | G | T | 2 | 0.2423 | 0.2423 | 0.9636 | 0.9636 | 0.5191 | 0 |
| 1 | 54080347 | rs4927017 | G | A | 2 | 0.2426 | 0.2426 | 0.9819 | 0.9819 | 0.5019 | 0 |
| 1 | 54596842 | rs5768066 | C | T | 2 | 0.2428 | 0.2428 | 0.9839 | 0.9839 | 0.5147 | 0 |
| 1 | 54005423 | rs1273169 | T | C | 2 | 0.2434 | 0.2434 | 1.0164 | 1.0164 | 0.4345 | 0 |
| 1 | 54124942 | rs3006904 | A | T | 2 | 0.2434 | 0.3012 | 0.9256 | 0.9288 | 0.2912 | 10.25 |
| 1 | 54186786 | rs9803748 | A | G | 2 | 0.2436 | 0.2561 | 0.9769 | 0.9767 | 0.301 | 6.51 |
| 1 | 54124317 | rs7516195 | T | C | 2 | 0.2437 | 0.2437 | 1.1098 | 1.1098 | 0.3952 | 0 |
| 1 | 54098711 | rs3107571 | T | C | 2 | 0.2439 | 0.2439 | 1.018 | 1.018 | 0.3348 | 0 |
| 1 | 53679878 | rs1056438 | T | C | 2 | 0.2441 | 0.2441 | 1.017 | 1.017 | 0.5434 | 0 |
| 1 | 54463596 | rs666725 | A | G | 2 | 0.245 | 0.5109 | 1.0163 | 1.0137 | 0.1442 | 53.11 |
| 1 | 53639748 | rs6663288 | G | A | 2 | 0.2451 | 0.2451 | 1.0172 | 1.0172 | 0.5533 | 0 |
| 1 | 53526787 | rs1180069 | G | A | 2 | 0.2453 | 0.2453 | 1.0226 | 1.0226 | 0.434 | 0 |

| | | | | | | | | | | | |
|---|---|---|---|---|---|---|---|---|---|---|---|
| 1 | 54080842 | rs1088879 | C | G | 2 | 0.2453 | 0.2453 | 0.9819 | 0.9819 | 0.5191 | 0 |
| 1 | 53937182 | rs7586525 | G | C | 2 | 0.2454 | 0.2595 | 0.9713 | 0.9707 | 0.296 | 8.42 |
| 1 | 54338772 | rs6177482 | G | A | 2 | 0.2455 | 0.2455 | 0.9638 | 0.9638 | 0.5944 | 0 |
| 1 | 54186445 | rs1203170 | C | T | 2 | 0.2456 | 0.2456 | 0.9764 | 0.9764 | 0.6713 | 0 |
| 1 | 54323528 | rs7416528 | C | A | 2 | 0.2461 | 0.2461 | 0.9642 | 0.9642 | 0.5969 | 0 |
| 1 | 53874117 | rs1288586 | T | C | 2 | 0.2472 | 0.2472 | 0.9841 | 0.9841 | 0.5618 | 0 |
| 1 | 54619106 | rs1209697 | C | T | 2 | 0.2474 | 0.2474 | 1.0193 | 1.0193 | 0.9712 | 0 |
| 1 | 54240485 | rs7163781 | G | A | 2 | 0.2483 | 0.2483 | 0.9618 | 0.9618 | 0.4895 | 0 |
| 1 | 54006043 | rs7540317 | G | C | 2 | 0.2488 | 0.3033 | 1.0163 | 1.0178 | 0.2306 | 30.4 |
| 1 | 54339484 | rs6177482 | T | A | 2 | 0.2492 | 0.2492 | 0.9641 | 0.9641 | 0.5927 | 0 |
| 1 | 54641638 | rs7986711 | T | C | 2 | 0.2492 | 0.2492 | 1.062 | 1.062 | 0.5633 | 0 |
| 1 | 54694948 | rs5945598 | C | T | 2 | 0.2493 | 0.3052 | 1.0235 | 1.0269 | 0.2177 | 34.17 |
| 1 | 54410655 | rs1206802 | T | C | 2 | 0.2497 | 0.2497 | 1.043 | 1.043 | 0.3989 | 0 |
| 1 | 54549712 | rs3464968 | G | A | 2 | 0.2497 | 0.2497 | 1.0858 | 1.0858 | 0.8163 | 0 |
| 1 | 54021321 | rs1203564 | T | C | 2 | 0.2503 | 0.2503 | 0.9741 | 0.9741 | 0.7325 | 0 |
| 1 | 54348716 | rs3737835 | T | C | 2 | 0.2504 | 0.2504 | 0.9641 | 0.9641 | 0.534 | 0 |
| 1 | 54422901 | rs1209272 | C | G | 2 | 0.2505 | 0.2505 | 1.0433 | 1.0433 | 0.508 | 0 |
| 1 | 54588354 | rs7530816 | T | G | 2 | 0.2508 | 0.2508 | 0.9846 | 0.9846 | 0.496 | 0 |
| 1 | 53650094 | rs4244641 | C | T | 2 | 0.251 | 0.251 | 1.0176 | 1.0176 | 0.8931 | 0 |
| 1 | 54590341 | rs673866 | C | A | 2 | 0.2512 | 0.2512 | 1.0158 | 1.0158 | 0.5309 | 0 |
| 1 | 54359922 | rs2294510 | C | T | 2 | 0.2513 | 0.2513 | 0.9672 | 0.9672 | 0.5129 | 0 |
| 1 | 54592594 | rs638643 | A | G | 2 | 0.2515 | 0.2515 | 1.016 | 1.016 | 0.5945 | 0 |
| 1 | 53898807 | rs5953799 | C | T | 2 | 0.2516 | 0.4718 | 0.9761 | 0.9714 | 0.0596 | 71.82 |
| 1 | 54080606 | rs5625641 | T | C | 2 | 0.2516 | 0.2516 | 0.9821 | 0.9821 | 0.5245 | 0 |
| 1 | 53623538 | rs1296145 | T | C | 2 | 0.2519 | 0.5715 | 1.0201 | 1.0161 | 0.1119 | 60.43 |
| 1 | 53682106 | rs1736939 | A | C | 2 | 0.2522 | 0.2522 | 1.0167 | 1.0167 | 0.5428 | 0 |
| 1 | 54643974 | rs1387427 | G | C | 2 | 0.2525 | 0.2525 | 1.0652 | 1.0652 | 0.6337 | 0 |
| 1 | 53660773 | rs1120612 | A | G | 2 | 0.2527 | 0.2527 | 1.0167 | 1.0167 | 0.5261 | 0 |
| 1 | 54340621 | rs6177483 | T | G | 2 | 0.2528 | 0.2528 | 0.9643 | 0.9643 | 0.5854 | 0 |
| 1 | 53844161 | rs1274631 | G | A | 2 | 0.253 | 0.253 | 1.0157 | 1.0157 | 0.6287 | 0 |
| 1 | 54598713 | rs4483354 | G | A | 2 | 0.2531 | 0.2531 | 0.9698 | 0.9698 | 0.9594 | 0 |
| 1 | 54531604 | rs7892909 | G | A | 2 | 0.2534 | 0.2534 | 1.0852 | 1.0852 | 0.8032 | 0 |
| 1 | 53683921 | rs1214058 | C | G | 2 | 0.2537 | 0.2537 | 1.0167 | 1.0167 | 0.5572 | 0 |
| 1 | 53753896 | rs1002357 | G | C | 2 | 0.254 | 0.8096 | 1.0281 | 0.9817 | 0.058 | 72.17 |
| 1 | 53702871 | rs1134799 | T | A | 2 | 0.2551 | 0.2551 | 0.976 | 0.976 | 0.8302 | 0 |
| 1 | 53675532 | rs1273737 | T | C | 2 | 0.2558 | 0.2558 | 1.0165 | 1.0165 | 0.5509 | 0 |
| 1 | 53862594 | rs1288582 | C | T | 2 | 0.256 | 0.256 | 0.9847 | 0.9847 | 0.7901 | 0 |
| 1 | 53684324 | rs3736118 | A | G | 2 | 0.2565 | 0.2565 | 1.0166 | 1.0166 | 0.5579 | 0 |
| 1 | 53481814 | rs1120607 | G | T | 2 | 0.2567 | 0.2567 | 1.022 | 1.022 | 0.4552 | 0 |
| 1 | 53635011 | rs3125251 | A | T | 2 | 0.2567 | 0.2567 | 1.0178 | 1.0178 | 0.7367 | 0 |
| 1 | 53636616 | rs3125252 | C | T | 2 | 0.2571 | 0.2571 | 1.0169 | 1.0169 | 0.9469 | 0 |
| 1 | 53965066 | rs749378 | A | G | 2 | 0.2573 | 0.3314 | 1.0173 | 1.0191 | 0.2063 | 37.4 |
| 1 | 53898195 | rs7849315 | G | A | 2 | 0.2574 | 0.2574 | 1.0239 | 1.0239 | 0.7423 | 0 |
| 1 | 54272330 | rs1181145 | C | T | 2 | 0.2579 | 0.2579 | 0.9685 | 0.9685 | 0.7383 | 0 |
| 1 | 54624281 | rs675602 | A | G | 2 | 0.2582 | 0.2911 | 1.0345 | 1.0348 | 0.2811 | 13.94 |
| 1 | 53478528 | rs1158048 | T | C | 2 | 0.2584 | 0.2584 | 1.0222 | 1.0222 | 0.4571 | 0 |
| 1 | 53896599 | rs1202314 | C | T | 2 | 0.2586 | 0.4634 | 0.9764 | 0.972 | 0.0709 | 69.35 |
| 1 | 53677341 | rs1381999 | G | A | 2 | 0.2589 | 0.2589 | 1.0695 | 1.0695 | 0.9102 | 0 |

| | | | | | | | | | | | |
|---|---|---|---|---|---|---|---|---|---|---|---|
| 1 | 54076383 | rs1163665 | C | T | 2 | 0.2589 | 0.2589 | 0.9543 | 0.9543 | 0.481 | 0 |
| 1 | 53884654 | rs5992433 | T | A | 2 | 0.259 | 0.259 | 1.0238 | 1.0238 | 0.8651 | 0 |
| 1 | 53526247 | rs725510 | C | T | 2 | 0.2591 | 0.2591 | 1.0219 | 1.0219 | 0.4704 | 0 |
| 1 | 53494091 | rs4442320 | C | A | 2 | 0.2594 | 0.2594 | 1.0219 | 1.0219 | 0.4169 | 0 |
| 1 | 54633259 | rs1214349 | T | C | 2 | 0.2595 | 0.2595 | 1.0255 | 1.0255 | 0.4659 | 0 |
| 1 | 54353785 | rs1202819 | G | C | 2 | 0.2604 | 0.2604 | 0.9649 | 0.9649 | 0.4979 | 0 |
| 1 | 53512285 | rs1337650 | T | C | 2 | 0.2605 | 0.2605 | 1.0316 | 1.0316 | 0.5715 | 0 |
| 1 | 53865432 | rs5695943 | C | T | 2 | 0.2605 | 0.2605 | 1.059 | 1.059 | 0.3743 | 0 |
| 1 | 53967454 | rs1272704 | T | C | 2 | 0.2606 | 0.2606 | 1.0342 | 1.0342 | 0.3251 | 0 |
| 1 | 54382357 | rs3477980 | C | G | 2 | 0.2607 | 0.2607 | 0.9799 | 0.9799 | 0.9833 | 0 |
| 1 | 54353126 | rs3458911 | A | C | 2 | 0.2608 | 0.2608 | 0.9649 | 0.9649 | 0.4995 | 0 |
| 1 | 53917398 | rs2153944 | G | A | 2 | 0.261 | 0.3418 | 0.9711 | 0.9677 | 0.195 | 40.45 |
| 1 | 54005060 | rs1120616 | C | T | 2 | 0.2613 | 0.2932 | 1.0159 | 1.0168 | 0.266 | 19.17 |
| 1 | 54694254 | rs1120630 | C | G | 2 | 0.2613 | 0.2693 | 1.0226 | 1.023 | 0.3038 | 5.42 |
| 1 | 54647165 | rs1123055 | C | G | 2 | 0.2616 | 0.2616 | 0.9685 | 0.9685 | 0.7341 | 0 |
| 1 | 53952141 | rs1288617 | G | A | 2 | 0.2619 | 0.3901 | 0.9654 | 0.9681 | 0.2377 | 28.27 |
| 1 | 54241178 | rs1181194 | C | T | 2 | 0.2619 | 0.4398 | 0.9261 | 0.9207 | 0.1198 | 58.68 |
| 1 | 53476970 | rs6679819 | G | A | 2 | 0.262 | 0.262 | 1.0221 | 1.0221 | 0.5733 | 0 |
| 1 | 53684924 | rs6797463 | A | G | 2 | 0.262 | 0.262 | 1.0165 | 1.0165 | 0.5619 | 0 |
| 1 | 53910654 | rs5929390 | C | T | 2 | 0.2621 | 0.2621 | 0.9613 | 0.9613 | 0.454 | 0 |
| 1 | 53837269 | rs1158979 | C | T | 2 | 0.2624 | 0.2624 | 0.9822 | 0.9822 | 0.9706 | 0 |
| 1 | 54685855 | rs3398869 | G | A | 2 | 0.2624 | 0.2624 | 1.021 | 1.021 | 0.3861 | 0 |
| 1 | 53899711 | rs6176841 | A | T | 2 | 0.2625 | 0.2625 | 1.0236 | 1.0236 | 0.755 | 0 |
| 1 | 54084331 | rs3013745 | G | A | 2 | 0.2626 | 0.2626 | 1.017 | 1.017 | 0.6277 | 0 |
| 1 | 54238825 | rs3403784 | T | C | 2 | 0.2635 | 0.2635 | 0.9635 | 0.9635 | 0.5158 | 0 |
| 1 | 53525162 | rs1205993 | G | A | 2 | 0.2637 | 0.2637 | 1.0217 | 1.0217 | 0.4642 | 0 |
| 1 | 54687523 | rs1158588 | A | T | 2 | 0.2641 | 0.2641 | 1.0207 | 1.0207 | 0.4343 | 0 |
| 1 | 54592965 | rs640458 | C | T | 2 | 0.2643 | 0.2643 | 1.0153 | 1.0153 | 0.8755 | 0 |
| 1 | 54657181 | rs1078897 | A | C | 2 | 0.2644 | 0.2644 | 1.0386 | 1.0386 | 0.4085 | 0 |
| 1 | 54064240 | rs1015929 | C | T | 2 | 0.2656 | 0.6542 | 0.9823 | 0.9873 | 0.0876 | 65.72 |
| 1 | 54029642 | rs4927007 | C | T | 2 | 0.2658 | 0.5198 | 1.0153 | 1.0215 | 0.0183 | 82.04 |
| 1 | 54083646 | rs2950265 | A | G | 2 | 0.2659 | 0.2659 | 1.0169 | 1.0169 | 0.6339 | 0 |
| 1 | 54683014 | rs1410896 | C | G | 2 | 0.266 | 0.266 | 1.0208 | 1.0208 | 0.4188 | 0 |
| 1 | 54005821 | rs1275075 | G | A | 2 | 0.2665 | 0.3201 | 1.0159 | 1.0176 | 0.2288 | 30.96 |
| 1 | 53701581 | rs2297656 | A | G | 2 | 0.2668 | 0.2668 | 1.016 | 1.016 | 0.6019 | 0 |
| 1 | 53796222 | rs5850803 | A | G | 2 | 0.2668 | 0.2668 | 1.0787 | 1.0787 | 0.9085 | 0 |
| 1 | 53666108 | rs1275495 | T | C | 2 | 0.267 | 0.267 | 1.0162 | 1.0162 | 0.5284 | 0 |
| 1 | 53810076 | rs1207054 | T | C | 2 | 0.267 | 0.267 | 1.0637 | 1.0637 | 0.3719 | 0 |
| 1 | 54605520 | rs648573 | T | C | 2 | 0.2671 | 0.2671 | 1.0545 | 1.0545 | 0.6106 | 0 |
| 1 | 53483431 | rs1039993 | T | C | 2 | 0.2673 | 0.2673 | 1.0216 | 1.0216 | 0.4291 | 0 |
| 1 | 54296839 | rs1272524 | A | T | 2 | 0.2679 | 0.2679 | 0.9612 | 0.9612 | 0.5291 | 0 |
| 1 | 53799774 | rs1158364 | C | G | 2 | 0.2682 | 0.2682 | 1.0182 | 1.0182 | 0.6053 | 0 |
| 1 | 53890752 | rs1445559 | A | G | 2 | 0.2685 | 0.4844 | 0.9767 | 0.9715 | 0.0557 | 72.69 |
| 1 | 54602115 | rs688829 | T | C | 2 | 0.2685 | 0.2685 | 1.0521 | 1.0521 | 0.9104 | 0 |
| 1 | 54441853 | rs7408450 | A | G | 2 | 0.2686 | 0.6431 | 1.0403 | 1.0329 | 0.0527 | 73.36 |
| 1 | 54541664 | rs5627840 | G | A | 2 | 0.269 | 0.269 | 1.0822 | 1.0822 | 0.7713 | 0 |
| 1 | 54588398 | rs7516735 | C | A | 2 | 0.2691 | 0.2691 | 0.9851 | 0.9851 | 0.5388 | 0 |
| 1 | 53663128 | rs3766760 | G | T | 2 | 0.2694 | 0.2694 | 1.0161 | 1.0161 | 0.4723 | 0 |

| | | | | | | | | | | | |
|---|---|---|---|---|---|---|---|---|---|---|---|
| 1 | 54549291 | rs7844083 | G | A | 2 | 0.2699 | 0.2699 | 0.953 | 0.953 | 0.7692 | 0 |
| 1 | 53561879 | rs1209503 | C | T | 2 | 0.27 | 0.27 | 0.9533 | 0.9533 | 0.9898 | 0 |
| 1 | 53486745 | rs1180663 | T | A | 2 | 0.2701 | 0.2701 | 1.0214 | 1.0214 | 0.4396 | 0 |
| 1 | 53531449 | rs1043707 | T | A | 2 | 0.2704 | 0.2704 | 1.0221 | 1.0221 | 0.4667 | 0 |
| 1 | 53824808 | rs7408396 | A | C | 2 | 0.2706 | 0.2706 | 1.163 | 1.163 | 0.4661 | 0 |
| 1 | 53930551 | rs943514 | G | A | 2 | 0.2707 | 0.4789 | 0.966 | 0.9704 | 0.1876 | 42.4 |
| 1 | 54265866 | rs1181178 | T | C | 2 | 0.2707 | 0.2707 | 0.9693 | 0.9693 | 0.7434 | 0 |
| 1 | 53488564 | rs7546714 | T | A | 2 | 0.2711 | 0.2711 | 1.0214 | 1.0214 | 0.4384 | 0 |
| 1 | 53527601 | rs8034533 | G | T | 2 | 0.2711 | 0.5232 | 1.1549 | 1.1869 | 0.0419 | 75.83 |
| 1 | 54608848 | rs1137911 | A | C | 2 | 0.2713 | 0.2713 | 0.8872 | 0.8872 | 0.9517 | 0 |
| 1 | 53694065 | rs1211660 | C | T | 2 | 0.2719 | 0.2719 | 1.0161 | 1.0161 | 0.4595 | 0 |
| 1 | 54630030 | rs1150354 | T | C | 2 | 0.272 | 0.272 | 1.0346 | 1.0346 | 0.454 | 0 |
| 1 | 53891054 | rs1204622 | C | T | 2 | 0.2721 | 0.4959 | 0.977 | 0.9713 | 0.0475 | 74.54 |
| 1 | 53906995 | rs6176841 | C | T | 2 | 0.2727 | 0.2727 | 1.0231 | 1.0231 | 0.7781 | 0 |
| 1 | 54224706 | rs1185903 | C | T | 2 | 0.2732 | 0.6763 | 0.942 | 0.9631 | 0.1287 | 56.67 |
| 1 | 54031431 | rs1272804 | C | G | 2 | 0.2733 | 0.5143 | 1.0151 | 1.0209 | 0.0235 | 80.5 |
| 1 | 53973353 | rs1710865 | G | C | 2 | 0.2734 | 0.2734 | 0.9189 | 0.9189 | 0.3981 | 0 |
| 1 | 54630656 | rs1165487 | T | C | 2 | 0.2734 | 0.2734 | 1.0345 | 1.0345 | 0.4514 | 0 |
| 1 | 54154922 | rs3013774 | C | G | 2 | 0.2735 | 0.2735 | 0.9681 | 0.9681 | 0.9303 | 0 |
| 1 | 53870831 | rs1288588 | A | G | 2 | 0.274 | 0.274 | 0.985 | 0.985 | 0.5147 | 0 |
| 1 | 54630231 | rs1150363 | G | A | 2 | 0.2748 | 0.2748 | 1.0344 | 1.0344 | 0.4507 | 0 |
| 1 | 53510975 | rs6588458 | G | A | 2 | 0.2749 | 0.2749 | 1.0212 | 1.0212 | 0.3518 | 0 |
| 1 | 53495130 | rs1043706 | G | A | 2 | 0.275 | 0.275 | 1.0212 | 1.0212 | 0.4493 | 0 |
| 1 | 53486094 | rs6684171 | T | C | 2 | 0.2754 | 0.2754 | 1.0212 | 1.0212 | 0.4473 | 0 |
| 1 | 53638694 | rs1213048 | G | T | 2 | 0.2754 | 0.2754 | 1.0161 | 1.0161 | 0.9505 | 0 |
| 1 | 53494441 | rs4244640 | A | G | 2 | 0.2757 | 0.2757 | 1.0212 | 1.0212 | 0.4467 | 0 |
| 1 | 53838156 | rs1999899 | G | T | 2 | 0.2757 | 0.2757 | 0.9781 | 0.9781 | 0.7386 | 0 |
| 1 | 53481759 | rs1120607 | C | G | 2 | 0.2759 | 0.2759 | 1.0212 | 1.0212 | 0.4427 | 0 |
| 1 | 53930905 | rs1288635 | C | T | 2 | 0.276 | 0.4801 | 0.9663 | 0.9707 | 0.19 | 41.78 |
| 1 | 54594131 | rs4927056 | C | T | 2 | 0.276 | 0.276 | 0.985 | 0.985 | 0.5659 | 0 |
| 1 | 54576639 | rs1120628 | G | T | 2 | 0.2761 | 0.5238 | 0.9851 | 0.9873 | 0.1533 | 50.96 |
| 1 | 54563156 | rs6588502 | A | G | 2 | 0.2762 | 0.4822 | 0.9852 | 0.9869 | 0.1802 | 44.33 |
| 1 | 54252710 | rs3408813 | T | A | 2 | 0.2764 | 0.2764 | 0.9646 | 0.9646 | 0.5428 | 0 |
| 1 | 53502569 | rs1871749 | T | C | 2 | 0.2767 | 0.2767 | 1.0211 | 1.0211 | 0.4038 | 0 |
| 1 | 53694388 | rs2275087 | T | C | 2 | 0.2767 | 0.2767 | 1.0159 | 1.0159 | 0.4712 | 0 |
| 1 | 54259909 | rs6177480 | A | G | 2 | 0.2772 | 0.2772 | 0.9646 | 0.9646 | 0.5439 | 0 |
| 1 | 53855986 | rs1399016 | C | T | 2 | 0.2784 | 0.2784 | 0.9545 | 0.9545 | 0.74 | 0 |
| 1 | 54421202 | rs1204454 | C | T | 2 | 0.2794 | 0.2794 | 1.0407 | 1.0407 | 0.4967 | 0 |
| 1 | 54301458 | rs4130586 | G | T | 2 | 0.2799 | 0.2799 | 1.0225 | 1.0225 | 0.555 | 0 |
| 1 | 53810030 | rs1207518 | A | G | 2 | 0.2801 | 0.2801 | 1.0619 | 1.0619 | 0.4003 | 0 |
| 1 | 54683925 | rs6621 | G | A | 2 | 0.2801 | 0.2801 | 1.0202 | 1.0202 | 0.401 | 0 |
| 1 | 54207274 | rs797900 | G | A | 2 | 0.2805 | 0.2805 | 0.9762 | 0.9762 | 0.8608 | 0 |
| 1 | 54652687 | rs1120629 | T | C | 2 | 0.2807 | 0.2807 | 1.0624 | 1.0624 | 0.3191 | 0 |
| 1 | 53951204 | rs1288619 | G | A | 2 | 0.2808 | 0.6784 | 0.9685 | 0.9783 | 0.0853 | 66.23 |
| 1 | 54157022 | rs1158679 | C | A | 2 | 0.2808 | 0.2808 | 0.9774 | 0.9774 | 0.9156 | 0 |
| 1 | 54400344 | rs1120625 | G | C | 2 | 0.2808 | 0.2808 | 1.0402 | 1.0402 | 0.5548 | 0 |
| 1 | 54140639 | rs7436866 | G | A | 2 | 0.2809 | 0.2809 | 1.0406 | 1.0406 | 0.7103 | 0 |
| 1 | 53582321 | rs1679910 | G | A | 2 | 0.2811 | 0.3113 | 0.9848 | 0.9837 | 0.2643 | 19.74 |

| | | | | | | | | | | | |
|---|---|---|---|---|---|---|---|---|---|---|---|
| 1 | 54606522 | rs1207864 | T | C | 2 | 0.2815 | 0.2815 | 0.9667 | 0.9667 | 0.5953 | 0 |
| 1 | 53687589 | rs6663121 | T | C | 2 | 0.2821 | 0.2821 | 1.0147 | 1.0147 | 0.7595 | 0 |
| 1 | 53838168 | rs1213350 | T | A | 2 | 0.2821 | 0.2821 | 1.0149 | 1.0149 | 0.5332 | 0 |
| 1 | 53904746 | rs6176841 | C | T | 2 | 0.2828 | 0.2828 | 1.0226 | 1.0226 | 0.7371 | 0 |
| 1 | 54303469 | rs6177481 | T | G | 2 | 0.2829 | 0.2829 | 0.9632 | 0.9632 | 0.5057 | 0 |
| 1 | 54631365 | rs1157928 | C | T | 2 | 0.2829 | 0.2829 | 1.0338 | 1.0338 | 0.4611 | 0 |
| 1 | 54444609 | rs7408880 | A | G | 2 | 0.2834 | 0.6788 | 1.0391 | 1.0309 | 0.0422 | 75.76 |
| 1 | 53585723 | rs1769299 | G | T | 2 | 0.2835 | 0.2835 | 1.0149 | 1.0149 | 0.6907 | 0 |
| 1 | 53809900 | rs1206389 | G | T | 2 | 0.2836 | 0.2836 | 1.0616 | 1.0616 | 0.3725 | 0 |
| 1 | 54593036 | rs609198 | A | T | 2 | 0.2836 | 0.2836 | 1.0147 | 1.0147 | 0.8393 | 0 |
| 1 | 54548859 | rs3483455 | G | A | 2 | 0.2839 | 0.2839 | 0.9742 | 0.9742 | 0.8569 | 0 |
| 1 | 54593150 | rs641328 | C | G | 2 | 0.2839 | 0.2839 | 1.0147 | 1.0147 | 0.8261 | 0 |
| 1 | 54210787 | rs1183027 | C | G | 2 | 0.2843 | 0.2843 | 0.9761 | 0.9761 | 0.8835 | 0 |
| 1 | 54653112 | rs2275407 | A | C | 2 | 0.2843 | 0.2843 | 1.0619 | 1.0619 | 0.3206 | 0 |
| 1 | 54308631 | rs1273241 | A | G | 2 | 0.2844 | 0.2844 | 0.965 | 0.965 | 0.5104 | 0 |
| 1 | 53511146 | rs6588461 | A | G | 2 | 0.2849 | 0.2849 | 1.0209 | 1.0209 | 0.3454 | 0 |
| 1 | 53624259 | rs1288350 | G | A | 2 | 0.2851 | 0.2851 | 1.015 | 1.015 | 0.3892 | 0 |
| 1 | 53637636 | rs1159019 | C | T | 2 | 0.2852 | 0.2852 | 1.0158 | 1.0158 | 0.9264 | 0 |
| 1 | 54024840 | rs1203508 | G | A | 2 | 0.2852 | 0.2852 | 0.9758 | 0.9758 | 0.6886 | 0 |
| 1 | 54432317 | rs7163782 | A | G | 2 | 0.2852 | 0.2852 | 0.9319 | 0.9319 | 0.443 | 0 |
| 1 | 53753718 | rs1002358 | G | A | 2 | 0.2858 | 0.2858 | 0.9736 | 0.9736 | 0.7576 | 0 |
| 1 | 53616969 | rs1383867 | C | T | 2 | 0.286 | 0.3629 | 1.0327 | 1.0419 | 0.1596 | 49.44 |
| 1 | 53510129 | rs1120607 | A | G | 2 | 0.2862 | 0.2862 | 1.0359 | 1.0359 | 0.8912 | 0 |
| 1 | 53499512 | rs6131408 | T | C | 2 | 0.2863 | 0.2863 | 1.0207 | 1.0207 | 0.4309 | 0 |
| 1 | 54187349 | rs7582056 | G | A | 2 | 0.2864 | 0.2864 | 1.0378 | 1.0378 | 0.7713 | 0 |
| 1 | 54214435 | rs1500085 | C | T | 2 | 0.2865 | 0.2865 | 0.9313 | 0.9313 | 0.7581 | 0 |
| 1 | 53485839 | rs6588448 | A | G | 2 | 0.2869 | 0.2869 | 1.0209 | 1.0209 | 0.4521 | 0 |
| 1 | 53690097 | rs7443496 | G | A | 2 | 0.2869 | 0.2869 | 1.046 | 1.046 | 0.9041 | 0 |
| 1 | 54569981 | rs7785658 | C | G | 2 | 0.2869 | 0.2869 | 0.9525 | 0.9525 | 0.6394 | 0 |
| 1 | 54210713 | rs1185370 | C | T | 2 | 0.287 | 0.287 | 0.9762 | 0.9762 | 0.88 | 0 |
| 1 | 54661737 | rs6177522 | A | G | 2 | 0.2871 | 0.2871 | 1.0288 | 1.0288 | 0.6404 | 0 |
| 1 | 54231832 | rs4130275 | T | C | 2 | 0.2873 | 0.2873 | 0.9651 | 0.9651 | 0.5363 | 0 |
| 1 | 53848487 | rs6176838 | G | A | 2 | 0.2877 | 0.5075 | 0.9678 | 0.9581 | 0.0395 | 76.41 |
| 1 | 54139499 | rs7685785 | C | T | 2 | 0.2878 | 0.2878 | 1.0398 | 1.0398 | 0.7313 | 0 |
| 1 | 53757935 | rs1288499 | G | A | 2 | 0.2884 | 0.2884 | 0.9403 | 0.9403 | 0.377 | 0 |
| 1 | 53680090 | rs1056425 | G | A | 2 | 0.2885 | 0.2885 | 1.0154 | 1.0154 | 0.4881 | 0 |
| 1 | 53860219 | rs1157749 | A | G | 2 | 0.2889 | 0.2889 | 0.9557 | 0.9557 | 0.7327 | 0 |
| 1 | 53890586 | rs1434587 | G | A | 2 | 0.2889 | 0.5037 | 0.9779 | 0.973 | 0.0554 | 72.76 |
| 1 | 53506855 | rs1209446 | T | C | 2 | 0.289 | 0.289 | 1.0206 | 1.0206 | 0.4081 | 0 |
| 1 | 53932064 | rs1288636 | C | T | 2 | 0.2892 | 0.5201 | 0.9673 | 0.9722 | 0.1735 | 46.02 |
| 1 | 54653874 | rs7545435 | C | T | 2 | 0.2893 | 0.2893 | 1.0612 | 1.0612 | 0.3286 | 0 |
| 1 | 54439366 | rs1209423 | A | G | 2 | 0.2895 | 0.6266 | 1.0345 | 1.0291 | 0.0677 | 70.04 |
| 1 | 54663157 | rs1203848 | C | T | 2 | 0.2895 | 0.2895 | 1.0725 | 1.0725 | 0.7025 | 0 |
| 1 | 54153635 | rs2950245 | C | T | 2 | 0.2897 | 0.2897 | 0.969 | 0.969 | 0.645 | 0 |
| 1 | 54233511 | rs7163781 | T | C | 2 | 0.2897 | 0.2897 | 0.9653 | 0.9653 | 0.5314 | 0 |
| 1 | 53600350 | rs1288354 | A | G | 2 | 0.2898 | 0.3381 | 0.9722 | 0.9691 | 0.2313 | 30.22 |
| 1 | 53657567 | rs7549319 | C | T | 2 | 0.2901 | 0.2901 | 1.015 | 1.015 | 0.5194 | 0 |
| 1 | 53810601 | rs1207073 | T | C | 2 | 0.2902 | 0.2902 | 1.0606 | 1.0606 | 0.3418 | 0 |

| | | | | | | | | | | | |
|---|---|---|---|---|---|---|---|---|---|---|---|
| 1 | 53830764 | rs7289945 | G | T | 2 | 0.2902 | 0.4809 | 0.9 | 0.8981 | 0.126 | 57.29 |
| 1 | 54188904 | rs2839615 | T | C | 2 | 0.2902 | 0.2902 | 0.9791 | 0.9791 | 0.9661 | 0 |
| 1 | 53838085 | rs1088878 | G | C | 2 | 0.2903 | 0.2903 | 1.0147 | 1.0147 | 0.5259 | 0 |
| 1 | 53777543 | rs2788033 | T | C | 2 | 0.2904 | 0.3561 | 0.9768 | 0.9604 | 0.1869 | 42.6 |
| 1 | 53519218 | rs1088876 | C | T | 2 | 0.2907 | 0.2907 | 1.0205 | 1.0205 | 0.377 | 0 |
| 1 | 53519795 | rs1256927 | T | C | 2 | 0.2911 | 0.2911 | 1.0205 | 1.0205 | 0.3996 | 0 |
| 1 | 53652092 | rs7786214 | C | T | 2 | 0.2917 | 0.2917 | 0.9329 | 0.9329 | 0.5617 | 0 |
| 1 | 53651956 | rs7918948 | G | A | 2 | 0.2919 | 0.2919 | 0.9329 | 0.9329 | 0.5514 | 0 |
| 1 | 54534223 | rs7408514 | T | C | 2 | 0.292 | 0.292 | 1.0783 | 1.0783 | 0.7151 | 0 |
| 1 | 54651982 | rs1120629 | C | T | 2 | 0.292 | 0.292 | 1.061 | 1.061 | 0.3225 | 0 |
| 1 | 54105364 | rs941123 | G | C | 2 | 0.2922 | 0.2922 | 0.9838 | 0.9838 | 0.3613 | 0 |
| 1 | 54489544 | rs6104784 | A | C | 2 | 0.2923 | 0.2923 | 1.0768 | 1.0768 | 0.8799 | 0 |
| 1 | 54004170 | rs1274432 | G | A | 2 | 0.2926 | 0.3468 | 1.0149 | 1.0165 | 0.2244 | 32.24 |
| 1 | 53908625 | rs1475542 | C | T | 2 | 0.2931 | 0.2931 | 1.0221 | 1.0221 | 0.754 | 0 |
| 1 | 54651314 | rs955079 | T | A | 2 | 0.2932 | 0.2932 | 1.0608 | 1.0608 | 0.3221 | 0 |
| 1 | 54615232 | rs4927065 | C | T | 2 | 0.2934 | 0.5551 | 1.0165 | 1.0134 | 0.1577 | 49.9 |
| 1 | 53643674 | rs3102042 | A | G | 2 | 0.2935 | 0.2935 | 1.0161 | 1.0161 | 0.8295 | 0 |
| 1 | 53512941 | rs6670842 | A | G | 2 | 0.2937 | 0.2937 | 1.0204 | 1.0204 | 0.3752 | 0 |
| 1 | 53581972 | rs1288380 | C | G | 2 | 0.2937 | 0.2937 | 0.9818 | 0.9818 | 0.8118 | 0 |
| 1 | 53514728 | rs3766763 | A | G | 2 | 0.294 | 0.294 | 1.0204 | 1.0204 | 0.3748 | 0 |
| 1 | 54314844 | rs7570585 | G | T | 2 | 0.294 | 0.294 | 1.0385 | 1.0385 | 0.9909 | 0 |
| 1 | 54018483 | rs1120617 | G | A | 2 | 0.2952 | 0.5282 | 0.9808 | 0.9748 | 0.0319 | 78.29 |
| 1 | 53958603 | rs1298637 | C | T | 2 | 0.2953 | 0.3406 | 0.9844 | 0.9832 | 0.2454 | 25.9 |
| 1 | 54671695 | rs3397356 | C | T | 2 | 0.2955 | 0.2955 | 0.9849 | 0.9849 | 0.8082 | 0 |
| 1 | 53657595 | rs1241023 | T | A | 2 | 0.2957 | 0.2957 | 1.0149 | 1.0149 | 0.5256 | 0 |
| 1 | 54651358 | rs955080 | C | T | 2 | 0.2962 | 0.2962 | 1.0604 | 1.0604 | 0.3266 | 0 |
| 1 | 53505103 | rs2170301 | G | A | 2 | 0.2963 | 0.2963 | 1.0203 | 1.0203 | 0.4223 | 0 |
| 1 | 54586538 | rs1206505 | T | C | 2 | 0.2966 | 0.2966 | 0.9858 | 0.9858 | 0.3865 | 0 |
| 1 | 53891559 | rs1203245 | G | C | 2 | 0.2967 | 0.5057 | 0.9782 | 0.9735 | 0.0588 | 72 |
| 1 | 54273279 | rs3526545 | T | C | 2 | 0.297 | 0.5911 | 0.9418 | 0.9552 | 0.1545 | 50.67 |
| 1 | 54294766 | rs4927040 | G | A | 2 | 0.2971 | 0.2971 | 1.0217 | 1.0217 | 0.5774 | 0 |
| 1 | 53519270 | rs1088876 | C | T | 2 | 0.2976 | 0.2976 | 1.0202 | 1.0202 | 0.3865 | 0 |
| 1 | 54609773 | rs1710991 | C | T | 2 | 0.2977 | 0.2977 | 1.0494 | 1.0494 | 0.9081 | 0 |
| 1 | 54688211 | rs1120630 | C | T | 2 | 0.2977 | 0.2977 | 1.0196 | 1.0196 | 0.4604 | 0 |
| 1 | 53511764 | rs1120607 | G | T | 2 | 0.2984 | 0.2984 | 1.0223 | 1.0223 | 0.3817 | 0 |
| 1 | 54047792 | rs7408475 | G | A | 2 | 0.2984 | 0.2984 | 1.0574 | 1.0574 | 0.5157 | 0 |
| 1 | 54303780 | rs1181201 | C | T | 2 | 0.2984 | 0.2984 | 0.9707 | 0.9707 | 0.6163 | 0 |
| 1 | 54419293 | rs1467640 | A | G | 2 | 0.299 | 0.299 | 0.9601 | 0.9601 | 0.4589 | 0 |
| 1 | 54059264 | rs3013756 | T | C | 2 | 0.2994 | 0.2994 | 0.9469 | 0.9469 | 0.4089 | 0 |
| 1 | 53484234 | rs1120607 | C | T | 2 | 0.2995 | 0.2995 | 1.0202 | 1.0202 | 0.4135 | 0 |
| 1 | 53787673 | rs1288521 | C | T | 2 | 0.2996 | 0.2996 | 1.016 | 1.016 | 0.7306 | 0 |
| 1 | 53700108 | rs6671485 | C | T | 2 | 0.2997 | 0.2997 | 1.0151 | 1.0151 | 0.5038 | 0 |
| 1 | 53867207 | rs1207398 | C | T | 2 | 0.3005 | 0.391 | 0.9221 | 0.9197 | 0.2134 | 35.4 |
| 1 | 54504580 | rs7991733 | C | T | 2 | 0.3007 | 0.3007 | 1.0755 | 1.0755 | 0.9094 | 0 |
| 1 | 54293128 | rs7991306 | T | A | 2 | 0.3008 | 0.3008 | 0.9659 | 0.9659 | 0.474 | 0 |
| 1 | 53998923 | rs7651190 | G | A | 2 | 0.3009 | 0.5447 | 0.9551 | 0.9644 | 0.2041 | 37.99 |
| 1 | 54263332 | rs7556247 | T | C | 2 | 0.3012 | 0.3012 | 0.9665 | 0.9665 | 0.5019 | 0 |
| 1 | 54198562 | rs797913 | C | T | 2 | 0.3014 | 0.3014 | 0.9769 | 0.9769 | 0.9609 | 0 |

| | | | | | | | | | | | |
|---|---|---|---|---|---|---|---|---|---|---|---|
| 1 | 54592563 | rs638605 | G | A | 2 | 0.3014 | 0.3014 | 1.0142 | 1.0142 | 0.8003 | 0 |
| 1 | 54073994 | rs1208312 | C | T | 2 | 0.3015 | 0.3015 | 0.9838 | 0.9838 | 0.4942 | 0 |
| 1 | 53535499 | rs1710783 | C | T | 2 | 0.3021 | 0.3021 | 1.0327 | 1.0327 | 0.6424 | 0 |
| 1 | 53928642 | rs6176842 | C | T | 2 | 0.3023 | 0.3023 | 1.0216 | 1.0216 | 0.9011 | 0 |
| 1 | 54298297 | rs7266233 | T | C | 2 | 0.3024 | 0.3024 | 1.0214 | 1.0214 | 0.5821 | 0 |
| 1 | 54696319 | rs1241086 | A | C | 2 | 0.3024 | 0.3034 | 1.0203 | 1.0203 | 0.3156 | 0.72 |
| 1 | 53522317 | rs1120608 | A | G | 2 | 0.3026 | 0.3026 | 1.02 | 1.02 | 0.4189 | 0 |
| 1 | 53667209 | rs7892388 | A | G | 2 | 0.3026 | 0.3026 | 0.9316 | 0.9316 | 0.8336 | 0 |
| 1 | 53520126 | rs7550236 | T | C | 2 | 0.3037 | 0.3037 | 1.02 | 1.02 | 0.387 | 0 |
| 1 | 53521486 | rs1078894 | G | C | 2 | 0.3038 | 0.3038 | 1.02 | 1.02 | 0.4176 | 0 |
| 1 | 54295139 | rs7266233 | C | G | 2 | 0.3038 | 0.3038 | 1.0214 | 1.0214 | 0.5852 | 0 |
| 1 | 54271441 | rs5594931 | C | A | 2 | 0.3039 | 0.3039 | 0.9666 | 0.9666 | 0.4973 | 0 |
| 1 | 54290029 | rs1272976 | C | T | 2 | 0.3043 | 0.3043 | 0.9668 | 0.9668 | 0.4613 | 0 |
| 1 | 54647953 | rs7835573 | A | G | 2 | 0.3045 | 0.3045 | 1.0594 | 1.0594 | 0.3183 | 0 |
| 1 | 53622093 | rs1288347 | A | G | 2 | 0.3048 | 0.3048 | 1.0141 | 1.0141 | 0.4357 | 0 |
| 1 | 53505596 | rs2847657 | A | G | 2 | 0.3051 | 0.3051 | 1.0199 | 1.0199 | 0.4132 | 0 |
| 1 | 53841430 | rs4926987 | A | C | 2 | 0.3053 | 0.3053 | 1.0145 | 1.0145 | 0.9596 | 0 |
| 1 | 53497686 | rs4638058 | T | C | 2 | 0.3059 | 0.3059 | 1.0199 | 1.0199 | 0.4718 | 0 |
| 1 | 53838287 | rs1999900 | G | C | 2 | 0.3063 | 0.3063 | 0.9795 | 0.9795 | 0.7292 | 0 |
| 1 | 54304671 | rs4131272 | G | T | 2 | 0.3063 | 0.3063 | 1.0693 | 1.0693 | 0.9571 | 0 |
| 1 | 53810119 | rs1207520 | A | G | 2 | 0.3065 | 0.3065 | 1.0584 | 1.0584 | 0.433 | 0 |
| 1 | 53526104 | rs725509 | A | G | 2 | 0.3071 | 0.3071 | 1.0291 | 1.0291 | 0.4679 | 0 |
| 1 | 54161514 | rs1164165 | C | G | 2 | 0.3072 | 0.3072 | 1.0443 | 1.0443 | 0.7132 | 0 |
| 1 | 54268959 | rs1455637 | C | T | 2 | 0.3073 | 0.3073 | 1.0711 | 1.0711 | 0.8864 | 0 |
| 1 | 54690037 | rs2282332 | C | T | 2 | 0.3073 | 0.3073 | 1.0199 | 1.0199 | 0.3519 | 0 |
| 1 | 53639661 | rs6688253 | C | T | 2 | 0.3074 | 0.3074 | 1.015 | 1.015 | 0.5882 | 0 |
| 1 | 54307860 | rs1410480 | C | T | 2 | 0.3075 | 0.6138 | 1.0823 | 1.0791 | 0.0517 | 73.58 |
| 1 | 53594397 | rs768771 | G | T | 2 | 0.3078 | 0.3078 | 1.0179 | 1.0179 | 0.7347 | 0 |
| 1 | 53931771 | rs7773324 | A | T | 2 | 0.3088 | 0.3088 | 1.0213 | 1.0213 | 0.8831 | 0 |
| 1 | 54255719 | rs1181170 | C | T | 2 | 0.309 | 0.5268 | 0.9331 | 0.9278 | 0.0826 | 66.8 |
| 1 | 53538669 | rs1214561 | G | A | 2 | 0.3097 | 0.3097 | 0.9855 | 0.9855 | 0.9052 | 0 |
| 1 | 54688902 | rs1088884 | A | G | 2 | 0.3099 | 0.3099 | 1.0189 | 1.0189 | 0.4105 | 0 |
| 1 | 53869317 | rs1288589 | A | G | 2 | 0.3103 | 0.3103 | 0.9858 | 0.9858 | 0.5666 | 0 |
| 1 | 53711089 | rs6176962 | T | C | 2 | 0.3105 | 0.3105 | 1.0753 | 1.0753 | 0.784 | 0 |
| 1 | 53489169 | rs7824608 | C | T | 2 | 0.3108 | 0.4805 | 1.0908 | 1.0935 | 0.1397 | 54.15 |
| 1 | 53491455 | rs4926945 | G | A | 2 | 0.3109 | 0.3679 | 1.0196 | 1.0208 | 0.2376 | 28.3 |
| 1 | 53598015 | rs6176815 | G | A | 2 | 0.3109 | 0.3109 | 0.9823 | 0.9823 | 0.8374 | 0 |
| 1 | 54591442 | rs689239 | T | C | 2 | 0.3109 | 0.3109 | 1.0137 | 1.0137 | 0.6478 | 0 |
| 1 | 53502973 | rs1710771 | T | G | 2 | 0.311 | 0.4094 | 1.0228 | 1.0249 | 0.1843 | 43.26 |
| 1 | 54178288 | rs1718055 | A | G | 2 | 0.3111 | 0.3111 | 0.9686 | 0.9686 | 0.777 | 0 |
| 1 | 54592897 | rs608395 | C | A | 2 | 0.3113 | 0.3113 | 1.0139 | 1.0139 | 0.7775 | 0 |
| 1 | 53901450 | rs6176841 | C | G | 2 | 0.3117 | 0.3117 | 1.0211 | 1.0211 | 0.8447 | 0 |
| 1 | 54379443 | rs7611467 | C | G | 2 | 0.3121 | 0.4078 | 0.976 | 0.9724 | 0.1678 | 47.43 |
| 1 | 54487417 | rs1204716 | C | T | 2 | 0.3121 | 0.3121 | 0.986 | 0.986 | 0.342 | 0 |
| 1 | 54413516 | rs1138768 | C | T | 2 | 0.3123 | 0.3123 | 0.9752 | 0.9752 | 0.503 | 0 |
| 1 | 54280994 | rs1274781 | A | G | 2 | 0.3124 | 0.3124 | 0.9672 | 0.9672 | 0.4927 | 0 |
| 1 | 54393492 | rs1207365 | C | A | 2 | 0.3127 | 0.3127 | 1.0374 | 1.0374 | 0.5459 | 0 |
| 1 | 53969475 | rs5592103 | G | C | 2 | 0.313 | 0.5132 | 0.9659 | 0.9653 | 0.1169 | 59.31 |

| | | | | | | | | | | | |
|---|---|---|---|---|---|---|---|---|---|---|---|
| 1 | 54394849 | rs1204785 | G | A | 2 | 0.3134 | 0.3134 | 1.0374 | 1.0374 | 0.5406 | 0 |
| 1 | 54616599 | rs9326026 | T | C | 2 | 0.3135 | 0.3135 | 1.0251 | 1.0251 | 0.3471 | 0 |
| 1 | 53995515 | rs1202191 | G | A | 2 | 0.314 | 0.4188 | 0.959 | 0.9565 | 0.188 | 42.31 |
| 1 | 54279562 | rs1273808 | C | T | 2 | 0.3146 | 0.3146 | 0.9675 | 0.9675 | 0.4913 | 0 |
| 1 | 53786149 | rs1288519 | T | G | 2 | 0.3151 | 0.3151 | 1.015 | 1.015 | 0.6801 | 0 |
| 1 | 54285811 | rs7555869 | G | A | 2 | 0.3152 | 0.3152 | 0.9673 | 0.9673 | 0.4909 | 0 |
| 1 | 54009233 | rs3889128 | G | C | 2 | 0.3153 | 0.3153 | 0.9839 | 0.9839 | 0.4127 | 0 |
| 1 | 54170122 | rs1173593 | A | C | 2 | 0.3153 | 0.3153 | 0.9687 | 0.9687 | 0.9572 | 0 |
| 1 | 54081106 | rs4926606 | C | T | 2 | 0.316 | 0.316 | 0.9846 | 0.9846 | 0.4183 | 0 |
| 1 | 53499225 | rs5571313 | T | C | 2 | 0.3165 | 0.3165 | 1.0196 | 1.0196 | 0.47 | 0 |
| 1 | 53964341 | rs1203500 | G | A | 2 | 0.3165 | 0.5215 | 0.9665 | 0.9667 | 0.1201 | 58.62 |
| 1 | 54080806 | rs1088879 | T | C | 2 | 0.3166 | 0.3166 | 0.9842 | 0.9842 | 0.4009 | 0 |
| 1 | 53693090 | rs6670999 | G | T | 2 | 0.3168 | 0.3168 | 1.0145 | 1.0145 | 0.5722 | 0 |
| 1 | 54370264 | rs2294511 | A | T | 2 | 0.3169 | 0.4846 | 0.9854 | 0.9872 | 0.2221 | 32.93 |
| 1 | 54301944 | rs1272790 | G | A | 2 | 0.3172 | 0.3172 | 0.9673 | 0.9673 | 0.4448 | 0 |
| 1 | 53796921 | rs1413874 | G | T | 2 | 0.3173 | 0.3173 | 1.0717 | 1.0717 | 0.5232 | 0 |
| 1 | 54227433 | rs6166489 | G | A | 2 | 0.3178 | 0.5424 | 1.0901 | 1.0888 | 0.1057 | 61.8 |
| 1 | 54158303 | rs6177477 | G | A | 2 | 0.3183 | 0.3183 | 0.9801 | 0.9801 | 0.9902 | 0 |
| 1 | 53910631 | rs1159265 | C | T | 2 | 0.3186 | 0.3186 | 1.0209 | 1.0209 | 0.6655 | 0 |
| 1 | 54648533 | rs1213910 | G | A | 2 | 0.3191 | 0.3191 | 1.0228 | 1.0228 | 0.3558 | 0 |
| 1 | 54282325 | rs3596674 | A | C | 2 | 0.3199 | 0.3199 | 0.9677 | 0.9677 | 0.4959 | 0 |
| 1 | 54522355 | rs5784304 | C | T | 2 | 0.3206 | 0.3206 | 1.0723 | 1.0723 | 0.8924 | 0 |
| 1 | 53859670 | rs7267513 | A | G | 2 | 0.3207 | 0.3207 | 1.0574 | 1.0574 | 0.5866 | 0 |
| 1 | 53624820 | rs1288353 | T | C | 2 | 0.3208 | 0.3208 | 1.0139 | 1.0139 | 0.4122 | 0 |
| 1 | 54422004 | rs7266240 | G | A | 2 | 0.3208 | 0.3208 | 0.9778 | 0.9778 | 0.3552 | 0 |
| 1 | 53481733 | rs7869511 | T | A | 2 | 0.3211 | 0.3211 | 0.9568 | 0.9568 | 0.3897 | 0 |
| 1 | 54390427 | rs1120625 | G | A | 2 | 0.3212 | 0.3212 | 1.0366 | 1.0366 | 0.5919 | 0 |
| 1 | 54294835 | rs3564790 | C | G | 2 | 0.3213 | 0.3213 | 0.9686 | 0.9686 | 0.4388 | 0 |
| 1 | 54397240 | rs1120625 | G | A | 2 | 0.3217 | 0.3217 | 1.0366 | 1.0366 | 0.5047 | 0 |
| 1 | 53810499 | rs1207069 | T | C | 2 | 0.3225 | 0.3225 | 1.0565 | 1.0565 | 0.3821 | 0 |
| 1 | 54519731 | rs1710126 | A | G | 2 | 0.323 | 0.323 | 1.0885 | 1.0885 | 0.7645 | 0 |
| 1 | 54185821 | rs1513166 | C | T | 2 | 0.3233 | 0.3233 | 1.0624 | 1.0624 | 0.9784 | 0 |
| 1 | 54411468 | rs1402339 | C | A | 2 | 0.3237 | 0.3797 | 0.9752 | 0.9737 | 0.237 | 28.48 |
| 1 | 54628044 | rs7635092 | C | T | 2 | 0.3237 | 0.3237 | 1.0308 | 1.0308 | 0.595 | 0 |
| 1 | 54446927 | rs6697447 | T | C | 2 | 0.324 | 0.324 | 1.025 | 1.025 | 0.7796 | 0 |
| 1 | 53930881 | rs4142851 | T | C | 2 | 0.3243 | 0.3243 | 1.0207 | 1.0207 | 0.8881 | 0 |
| 1 | 53550640 | rs4129475 | C | T | 2 | 0.3244 | 0.3244 | 1.0464 | 1.0464 | 0.8661 | 0 |
| 1 | 53857675 | rs7267513 | A | G | 2 | 0.3253 | 0.3253 | 1.057 | 1.057 | 0.5822 | 0 |
| 1 | 54607180 | rs3766461 | A | T | 2 | 0.3257 | 0.3257 | 0.9125 | 0.9125 | 0.8989 | 0 |
| 1 | 53923833 | rs1288624 | A | G | 2 | 0.3259 | 0.3259 | 0.9853 | 0.9853 | 0.5459 | 0 |
| 1 | 53975436 | rs475322 | C | A | 2 | 0.3263 | 0.3263 | 0.9618 | 0.9618 | 0.9107 | 0 |
| 1 | 53952100 | rs1160434 | A | T | 2 | 0.3268 | 0.3268 | 0.9608 | 0.9608 | 0.3441 | 0 |
| 1 | 54687065 | rs1088884 | T | C | 2 | 0.3271 | 0.3271 | 1.0181 | 1.0181 | 0.4532 | 0 |
| 1 | 53988977 | rs7552778 | G | A | 2 | 0.3272 | 0.3272 | 0.9575 | 0.9575 | 0.4796 | 0 |
| 1 | 54655774 | rs7528517 | G | A | 2 | 0.3273 | 0.3741 | 0.9545 | 0.9553 | 0.2801 | 14.3 |
| 1 | 54431294 | rs2294513 | A | G | 2 | 0.3274 | 0.3274 | 1.0367 | 1.0367 | 0.3788 | 0 |
| 1 | 54160939 | rs1439284 | T | C | 2 | 0.3276 | 0.3276 | 1.0486 | 1.0486 | 0.5546 | 0 |
| 1 | 53625057 | rs4926961 | G | C | 2 | 0.3283 | 0.3283 | 1.0137 | 1.0137 | 0.3942 | 0 |

| | | | | | | | | | | | |
|---|---|---|---|---|---|---|---|---|---|---|---|
| 1 | 54204448 | rs761489 | A | C | 2 | 0.3283 | 0.3283 | 0.9769 | 0.9769 | 0.8578 | 0 |
| 1 | 53657883 | rs5959717 | G | T | 2 | 0.3286 | 0.7797 | 0.907 | 0.9351 | 0.0177 | 82.22 |
| 1 | 54660515 | rs1212028 | T | C | 2 | 0.3288 | 0.3288 | 0.9864 | 0.9864 | 0.9882 | 0 |
| 1 | 53852595 | rs7267513 | G | T | 2 | 0.329 | 0.329 | 1.0558 | 1.0558 | 0.6042 | 0 |
| 1 | 53810614 | rs1207538 | A | G | 2 | 0.3293 | 0.3293 | 1.0557 | 1.0557 | 0.3943 | 0 |
| 1 | 54276154 | rs7677615 | G | A | 2 | 0.3296 | 0.3296 | 0.9678 | 0.9678 | 0.5113 | 0 |
| 1 | 53936718 | rs1157957 | G | T | 2 | 0.3298 | 0.3298 | 1.0204 | 1.0204 | 0.901 | 0 |
| 1 | 53598989 | rs5604727 | G | A | 2 | 0.33 | 0.4515 | 1.0279 | 1.0396 | 0.0824 | 66.86 |
| 1 | 54208205 | rs797897 | A | G | 2 | 0.3301 | 0.3301 | 0.9786 | 0.9786 | 0.7151 | 0 |
| 1 | 54205185 | rs7534888 | G | C | 2 | 0.3302 | 0.3302 | 0.977 | 0.977 | 0.7146 | 0 |
| 1 | 54593256 | rs1572731 | A | T | 2 | 0.3307 | 0.3307 | 0.9867 | 0.9867 | 0.6249 | 0 |
| 1 | 53631029 | rs1214096 | A | G | 2 | 0.3308 | 0.3308 | 1.0139 | 1.0139 | 0.4365 | 0 |
| 1 | 53836899 | rs5914037 | C | T | 2 | 0.3308 | 0.3308 | 1.0143 | 1.0143 | 0.6131 | 0 |
| 1 | 53477900 | rs1144005 | C | T | 2 | 0.3318 | 0.4673 | 1.0874 | 1.089 | 0.1741 | 45.87 |
| 1 | 53592670 | rs1492583 | G | A | 2 | 0.3325 | 0.3325 | 0.8828 | 0.8828 | 0.7978 | 0 |
| 1 | 54475406 | rs1134611 | C | A | 2 | 0.3337 | 0.3337 | 0.976 | 0.976 | 0.8537 | 0 |
| 1 | 54157106 | rs1710911 | G | A | 2 | 0.3341 | 0.3341 | 1.0259 | 1.0259 | 0.896 | 0 |
| 1 | 53595255 | rs1679937 | G | T | 2 | 0.3342 | 0.3342 | 0.9845 | 0.9845 | 0.8509 | 0 |
| 1 | 54619337 | rs657783 | C | T | 2 | 0.3343 | 0.3343 | 1.0298 | 1.0298 | 0.3322 | 0 |
| 1 | 54072816 | rs2948040 | A | G | 2 | 0.3344 | 0.3344 | 1.0151 | 1.0151 | 0.5513 | 0 |
| 1 | 53657086 | rs7523531 | G | A | 2 | 0.3345 | 0.3345 | 1.0138 | 1.0138 | 0.5859 | 0 |
| 1 | 54399086 | rs1437499 | C | T | 2 | 0.3347 | 0.3347 | 1.0442 | 1.0442 | 0.7146 | 0 |
| 1 | 53586011 | rs6666264 | G | A | 2 | 0.3348 | 0.3348 | 0.9866 | 0.9866 | 0.4096 | 0 |
| 1 | 53625039 | rs4926960 | G | T | 2 | 0.3351 | 0.3351 | 1.013 | 1.013 | 0.794 | 0 |
| 1 | 53530617 | rs1710780 | A | C | 2 | 0.3364 | 0.3364 | 1.0274 | 1.0274 | 0.6288 | 0 |
| 1 | 53883126 | rs2000239 | G | A | 2 | 0.3364 | 0.3364 | 1.0145 | 1.0145 | 0.7463 | 0 |
| 1 | 53587179 | rs6694834 | C | T | 2 | 0.3368 | 0.3368 | 0.9867 | 0.9867 | 0.4058 | 0 |
| 1 | 54058831 | rs3013757 | T | C | 2 | 0.3371 | 0.3371 | 0.9511 | 0.9511 | 0.4454 | 0 |
| 1 | 53593913 | rs3766800 | A | G | 2 | 0.338 | 0.338 | 0.9832 | 0.9832 | 0.7762 | 0 |
| 1 | 53909020 | rs7512516 | G | C | 2 | 0.338 | 0.338 | 0.9655 | 0.9655 | 0.3915 | 0 |
| 1 | 53656676 | rs3531608 | A | G | 2 | 0.3382 | 0.3382 | 1.0136 | 1.0136 | 0.6042 | 0 |
| 1 | 54211693 | rs1146865 | C | T | 2 | 0.3384 | 0.3384 | 0.9773 | 0.9773 | 0.8375 | 0 |
| 1 | 54206278 | rs797901 | G | C | 2 | 0.3386 | 0.3386 | 0.9787 | 0.9787 | 0.6337 | 0 |
| 1 | 53658178 | rs7526732 | G | A | 2 | 0.3387 | 0.3387 | 1.0132 | 1.0132 | 0.8865 | 0 |
| 1 | 54536050 | rs1132196 | A | G | 2 | 0.3394 | 0.3394 | 1.0668 | 1.0668 | 0.3859 | 0 |
| 1 | 53696051 | rs1078894 | C | T | 2 | 0.3399 | 0.3399 | 1.0138 | 1.0138 | 0.5134 | 0 |
| 1 | 53907060 | rs1288600 | A | C | 2 | 0.34 | 0.3702 | 0.9858 | 0.9851 | 0.2688 | 18.21 |
| 1 | 54638374 | rs1213800 | T | C | 2 | 0.3401 | 0.4734 | 0.9822 | 0.9841 | 0.2487 | 24.86 |
| 1 | 53626467 | rs1211915 | C | T | 2 | 0.3406 | 0.3406 | 1.0129 | 1.0129 | 0.7698 | 0 |
| 1 | 53999509 | rs3887651 | C | T | 2 | 0.3406 | 0.4697 | 0.9651 | 0.9615 | 0.1485 | 52.1 |
| 1 | 54297113 | rs1181172 | G | A | 2 | 0.3406 | 0.4278 | 0.943 | 0.9407 | 0.2118 | 35.86 |
| 1 | 53975989 | rs580939 | G | A | 2 | 0.3415 | 0.3415 | 0.9632 | 0.9632 | 0.8623 | 0 |
| 1 | 53693831 | rs1214553 | C | T | 2 | 0.3416 | 0.3416 | 1.0143 | 1.0143 | 0.5008 | 0 |
| 1 | 53995955 | rs1203687 | C | A | 2 | 0.3428 | 0.431 | 0.9611 | 0.9587 | 0.2025 | 38.44 |
| 1 | 53515305 | rs3766764 | G | A | 2 | 0.3437 | 0.3437 | 1.0203 | 1.0203 | 0.4493 | 0 |
| 1 | 54679912 | rs4927071 | T | C | 2 | 0.3438 | 0.3438 | 0.9849 | 0.9849 | 0.3764 | 0 |
| 1 | 53861570 | rs1288583 | A | G | 2 | 0.3439 | 0.3439 | 0.987 | 0.987 | 0.5817 | 0 |
| 1 | 54198206 | rs797910 | G | A | 2 | 0.3439 | 0.416 | 1.0313 | 1.0319 | 0.2368 | 28.54 |

| | | | | | | | | | | | |
|---|---|---|---|---|---|---|---|---|---|---|---|
| 1 | 53944451 | rs1157829 | G | A | 2 | 0.3446 | 0.3446 | 1.0198 | 1.0198 | 0.8843 | 0 |
| 1 | 53945201 | rs6052385 | G | A | 2 | 0.3446 | 0.3446 | 1.0198 | 1.0198 | 0.8843 | 0 |
| 1 | 53966330 | rs6034300 | A | G | 2 | 0.3446 | 0.3446 | 1.019 | 1.019 | 0.9515 | 0 |
| 1 | 54172592 | rs7266067 | T | G | 2 | 0.3449 | 0.3449 | 1.0466 | 1.0466 | 0.7857 | 0 |
| 1 | 54486829 | rs6681465 | C | T | 2 | 0.345 | 0.4213 | 1.0337 | 1.0324 | 0.261 | 20.86 |
| 1 | 53526696 | rs1120608 | C | G | 2 | 0.3457 | 0.3457 | 1.0265 | 1.0265 | 0.3965 | 0 |
| 1 | 53576030 | rs4926586 | C | A | 2 | 0.3458 | 0.3458 | 0.9845 | 0.9845 | 0.9291 | 0 |
| 1 | 54072759 | rs3013749 | A | T | 2 | 0.3462 | 0.3462 | 1.0147 | 1.0147 | 0.5823 | 0 |
| 1 | 53536920 | rs4129474 | G | A | 2 | 0.3466 | 0.3466 | 1.0292 | 1.0292 | 0.6732 | 0 |
| 1 | 54014761 | rs7289526 | C | T | 2 | 0.3477 | 0.3477 | 0.9846 | 0.9846 | 0.3775 | 0 |
| 1 | 54105636 | rs7520808 | C | G | 2 | 0.3477 | 0.3477 | 0.9857 | 0.9857 | 0.5373 | 0 |
| 1 | 54630948 | rs1120629 | C | T | 2 | 0.3477 | 0.5155 | 0.981 | 0.9767 | 0.0807 | 67.22 |
| 1 | 54587634 | rs1209445 | G | T | 2 | 0.3487 | 0.3669 | 0.9562 | 0.9558 | 0.2949 | 8.84 |
| 1 | 53850061 | rs7267513 | G | A | 2 | 0.3488 | 0.3488 | 1.0539 | 1.0539 | 0.5509 | 0 |
| 1 | 54485787 | rs7266412 | T | C | 2 | 0.3504 | 0.3601 | 0.9872 | 0.9872 | 0.3113 | 2.47 |
| 1 | 53556625 | rs7408414 | C | T | 2 | 0.3515 | 0.3515 | 1.0372 | 1.0372 | 0.9881 | 0 |
| 1 | 53807582 | rs2782492 | G | C | 2 | 0.3521 | 0.3521 | 1.0131 | 1.0131 | 0.8608 | 0 |
| 1 | 54018427 | rs1120617 | G | A | 2 | 0.3522 | 0.5773 | 0.9831 | 0.9759 | 0.0197 | 81.61 |
| 1 | 54186457 | rs1409492 | A | T | 2 | 0.3529 | 0.3529 | 1.0579 | 1.0579 | 0.9394 | 0 |
| 1 | 53707088 | rs6176962 | C | T | 2 | 0.3536 | 0.3536 | 1.061 | 1.061 | 0.9174 | 0 |
| 1 | 53597481 | rs5781416 | A | G | 2 | 0.3538 | 0.3538 | 0.9838 | 0.9838 | 0.7835 | 0 |
| 1 | 54038536 | rs1049317 | C | A | 2 | 0.355 | 0.7038 | 0.9846 | 0.9892 | 0.0976 | 63.55 |
| 1 | 53578890 | rs1710801 | G | A | 2 | 0.357 | 0.5833 | 1.0473 | 1.0568 | 0.046 | 74.9 |
| 1 | 53692437 | rs1213346 | T | A | 2 | 0.3573 | 0.3573 | 1.0135 | 1.0135 | 0.6166 | 0 |
| 1 | 54689962 | rs1120630 | G | C | 2 | 0.3575 | 0.3575 | 1.0177 | 1.0177 | 0.4004 | 0 |
| 1 | 54384502 | rs7922187 | C | T | 2 | 0.358 | 0.4335 | 0.9782 | 0.9751 | 0.1871 | 42.54 |
| 1 | 54355804 | rs7266237 | C | A | 2 | 0.3581 | 0.3581 | 1.0384 | 1.0384 | 0.3703 | 0 |
| 1 | 53718060 | rs6176962 | T | C | 2 | 0.3583 | 0.3583 | 1.0605 | 1.0605 | 0.9089 | 0 |
| 1 | 53703072 | rs7267313 | C | A | 2 | 0.3587 | 0.3587 | 1.0133 | 1.0133 | 0.4868 | 0 |
| 1 | 53777964 | rs1044974 | C | G | 2 | 0.3589 | 0.9428 | 0.9732 | 1.0054 | 0.1931 | 40.96 |
| 1 | 53785864 | rs945184 | A | G | 2 | 0.3595 | 0.3595 | 1.0136 | 1.0136 | 0.5814 | 0 |
| 1 | 54078562 | rs2950261 | A | G | 2 | 0.3596 | 0.3596 | 1.0144 | 1.0144 | 0.6319 | 0 |
| 1 | 53705904 | rs6176962 | C | T | 2 | 0.3602 | 0.3602 | 1.0601 | 1.0601 | 0.9284 | 0 |
| 1 | 53692488 | rs1213683 | A | G | 2 | 0.3603 | 0.3603 | 1.0133 | 1.0133 | 0.5204 | 0 |
| 1 | 53537833 | rs1256897 | G | T | 2 | 0.3605 | 0.3605 | 1.0205 | 1.0205 | 0.3663 | 0 |
| 1 | 53807340 | rs2095409 | T | C | 2 | 0.3608 | 0.3608 | 1.0128 | 1.0128 | 0.8244 | 0 |
| 1 | 53557944 | rs1158832 | C | T | 2 | 0.361 | 0.361 | 1.0367 | 1.0367 | 0.9945 | 0 |
| 1 | 53536261 | rs7290517 | C | T | 2 | 0.362 | 0.362 | 1.0282 | 1.0282 | 0.6488 | 0 |
| 1 | 53695467 | rs6176961 | C | T | 2 | 0.3625 | 0.3625 | 1.0599 | 1.0599 | 0.9178 | 0 |
| 1 | 53999758 | rs561664 | T | C | 2 | 0.3633 | 0.5214 | 0.9672 | 0.9623 | 0.1052 | 61.91 |
| 1 | 54615994 | rs780515 | G | A | 2 | 0.3635 | 0.4139 | 1.0226 | 1.0259 | 0.2151 | 34.93 |
| 1 | 53770124 | rs2788036 | T | C | 2 | 0.3645 | 0.3645 | 1.0153 | 1.0153 | 0.5146 | 0 |
| 1 | 53515596 | rs7534827 | A | G | 2 | 0.3647 | 0.3647 | 1.0255 | 1.0255 | 0.5286 | 0 |
| 1 | 54417254 | rs1145709 | C | T | 2 | 0.365 | 0.365 | 1.0448 | 1.0448 | 0.4246 | 0 |
| 1 | 54585049 | rs7526507 | G | A | 2 | 0.3651 | 0.3759 | 0.9519 | 0.9509 | 0.2982 | 7.6 |
| 1 | 53851029 | rs4926989 | C | T | 2 | 0.3656 | 0.3656 | 1.0124 | 1.0124 | 0.5993 | 0 |
| 1 | 53843155 | rs5603403 | C | T | 2 | 0.366 | 0.366 | 1.0521 | 1.0521 | 0.575 | 0 |
| 1 | 53807682 | rs2788029 | A | G | 2 | 0.3661 | 0.3661 | 1.0127 | 1.0127 | 0.8441 | 0 |

| | | | | | | | | | | | |
|---|---|---|---|---|---|---|---|---|---|---|---|
| 1 | 54229971 | rs1406557 | G | A | 2 | 0.3664 | 0.3664 | 1.0618 | 1.0618 | 0.9945 | 0 |
| 1 | 53843322 | rs5575413 | T | C | 2 | 0.3666 | 0.3666 | 1.052 | 1.052 | 0.5781 | 0 |
| 1 | 53660362 | rs1207254 | C | T | 2 | 0.3667 | 0.3667 | 0.9396 | 0.9396 | 0.7744 | 0 |
| 1 | 54408551 | rs7784091 | C | T | 2 | 0.3667 | 0.3667 | 0.9778 | 0.9778 | 0.4719 | 0 |
| 1 | 54487776 | rs1138923 | C | A | 2 | 0.3668 | 0.468 | 1.0323 | 1.0305 | 0.2436 | 26.44 |
| 1 | 53640636 | rs1411046 | C | T | 2 | 0.3672 | 0.3672 | 0.9414 | 0.9414 | 0.6226 | 0 |
| 1 | 54289215 | rs1181189 | A | G | 2 | 0.3674 | 0.3878 | 0.9462 | 0.9457 | 0.2919 | 9.99 |
| 1 | 54424448 | rs1088882 | G | A | 2 | 0.3674 | 0.7179 | 1.017 | 1.0121 | 0.0808 | 67.2 |
| 1 | 54113628 | rs1384133 | T | C | 2 | 0.3693 | 0.3693 | 1.0235 | 1.0235 | 0.854 | 0 |
| 1 | 53995030 | rs3855967 | G | A | 2 | 0.3695 | 0.5772 | 0.9674 | 0.961 | 0.0557 | 72.68 |
| 1 | 54361239 | rs2268180 | C | T | 2 | 0.3703 | 0.3703 | 0.9739 | 0.9739 | 0.4899 | 0 |
| 1 | 53807803 | rs2782493 | T | C | 2 | 0.3705 | 0.3705 | 1.0126 | 1.0126 | 0.8359 | 0 |
| 1 | 53523698 | rs4926947 | G | C | 2 | 0.3713 | 0.3713 | 1.0253 | 1.0253 | 0.5025 | 0 |
| 1 | 54422984 | rs5637007 | G | A | 2 | 0.3714 | 0.4451 | 1.0325 | 1.0313 | 0.2611 | 20.82 |
| 1 | 53615701 | rs1120611 | T | C | 2 | 0.3715 | 0.3715 | 1.0122 | 1.0122 | 0.6019 | 0 |
| 1 | 54657070 | rs6177522 | T | C | 2 | 0.3715 | 0.3715 | 1.023 | 1.023 | 0.8087 | 0 |
| 1 | 53784098 | rs6694764 | A | G | 2 | 0.372 | 0.372 | 0.9867 | 0.9867 | 0.8666 | 0 |
| 1 | 54604602 | rs634218 | G | T | 2 | 0.3723 | 0.3723 | 1.0406 | 1.0406 | 0.8911 | 0 |
| 1 | 53527948 | rs7513730 | T | G | 2 | 0.3724 | 0.3724 | 1.0174 | 1.0174 | 0.5685 | 0 |
| 1 | 53596663 | rs5574761 | C | G | 2 | 0.3725 | 0.4304 | 1.0254 | 1.0312 | 0.1834 | 43.5 |
| 1 | 53691287 | rs6176961 | C | G | 2 | 0.3726 | 0.3726 | 1.0587 | 1.0587 | 0.9298 | 0 |
| 1 | 54593248 | rs7163783 | G | A | 2 | 0.3729 | 0.3729 | 1.0912 | 1.0912 | 0.9033 | 0 |
| 1 | 53528528 | rs1383598 | G | C | 2 | 0.3732 | 0.3732 | 1.0174 | 1.0174 | 0.416 | 0 |
| 1 | 54537151 | rs1385808 | A | G | 2 | 0.3732 | 0.3732 | 1.063 | 1.063 | 0.7942 | 0 |
| 1 | 53846253 | rs1710840 | A | G | 2 | 0.3736 | 0.3736 | 1.0486 | 1.0486 | 0.6558 | 0 |
| 1 | 54685195 | rs2026045 | T | A | 2 | 0.3736 | 0.3736 | 1.0165 | 1.0165 | 0.4991 | 0 |
| 1 | 53782720 | rs5994599 | C | T | 2 | 0.3737 | 0.3737 | 1.0585 | 1.0585 | 0.6852 | 0 |
| 1 | 54199218 | rs1209593 | G | C | 2 | 0.374 | 0.374 | 0.9791 | 0.9791 | 0.9522 | 0 |
| 1 | 54590961 | rs7163782 | T | A | 2 | 0.3749 | 0.3749 | 1.0906 | 1.0906 | 0.9249 | 0 |
| 1 | 53846403 | rs1202793 | C | A | 2 | 0.3753 | 0.3753 | 1.0356 | 1.0356 | 0.3604 | 0 |
| 1 | 53969197 | rs5632427 | A | G | 2 | 0.3757 | 0.3757 | 1.0189 | 1.0189 | 0.7696 | 0 |
| 1 | 53595729 | rs3766808 | C | T | 2 | 0.376 | 0.376 | 0.9845 | 0.9845 | 0.7488 | 0 |
| 1 | 54361433 | rs2268182 | G | T | 2 | 0.3762 | 0.3762 | 0.9744 | 0.9744 | 0.4834 | 0 |
| 1 | 54546623 | rs1480064 | A | T | 2 | 0.3764 | 0.3764 | 1.0626 | 1.0626 | 0.3892 | 0 |
| 1 | 53799390 | rs1120614 | C | T | 2 | 0.3765 | 0.3765 | 0.924 | 0.924 | 0.8982 | 0 |
| 1 | 53859781 | rs7267514 | G | A | 2 | 0.3765 | 0.3765 | 1.0507 | 1.0507 | 0.5029 | 0 |
| 1 | 53715795 | rs7585412 | T | C | 2 | 0.3766 | 0.3766 | 0.9263 | 0.9263 | 0.474 | 0 |
| 1 | 54660252 | rs1485205 | C | T | 2 | 0.3768 | 0.3768 | 0.9793 | 0.9793 | 0.8338 | 0 |
| 1 | 54075349 | rs2948049 | A | G | 2 | 0.377 | 0.377 | 1.0139 | 1.0139 | 0.6127 | 0 |
| 1 | 54608460 | rs1710990 | C | T | 2 | 0.377 | 0.377 | 1.0414 | 1.0414 | 0.857 | 0 |
| 1 | 53519716 | rs5831395 | T | C | 2 | 0.3771 | 0.3771 | 1.0251 | 1.0251 | 0.5084 | 0 |
| 1 | 54373185 | rs3766467 | G | T | 2 | 0.3772 | 0.3772 | 1.0322 | 1.0322 | 0.3982 | 0 |
| 1 | 53513423 | rs1120607 | C | T | 2 | 0.3775 | 0.3775 | 1.025 | 1.025 | 0.5091 | 0 |
| 1 | 54072278 | rs1212211 | T | C | 2 | 0.3775 | 0.3775 | 1.0419 | 1.0419 | 0.5884 | 0 |
| 1 | 54357645 | rs1088881 | G | A | 2 | 0.3776 | 0.3776 | 0.9736 | 0.9736 | 0.5722 | 0 |
| 1 | 54693536 | rs1157766 | C | T | 2 | 0.3777 | 0.3777 | 1.0171 | 1.0171 | 0.4731 | 0 |
| 1 | 54075005 | rs3006893 | C | G | 2 | 0.3779 | 0.3779 | 1.0138 | 1.0138 | 0.6218 | 0 |
| 1 | 53790007 | rs5831618 | A | G | 2 | 0.378 | 0.378 | 1.0594 | 1.0594 | 0.84 | 0 |

| | | | | | | | | | | | |
|---|---|---|---|---|---|---|---|---|---|---|---|
| 1 | 54314043 | rs4354491 | C | T | 2 | 0.3784 | 0.3784 | 0.9856 | 0.9856 | 0.8044 | 0 |
| 1 | 54637202 | rs3795360 | C | T | 2 | 0.3784 | 0.3784 | 1.0503 | 1.0503 | 0.4075 | 0 |
| 1 | 53512519 | rs1158820 | G | T | 2 | 0.3785 | 0.3785 | 1.0249 | 1.0249 | 0.5101 | 0 |
| 1 | 54606148 | rs7995012 | C | T | 2 | 0.3787 | 0.3787 | 0.9826 | 0.9826 | 0.7807 | 0 |
| 1 | 54634941 | rs686106 | A | C | 2 | 0.3787 | 0.3787 | 1.0119 | 1.0119 | 0.4899 | 0 |
| 1 | 54008032 | rs6177026 | G | A | 2 | 0.3793 | 0.4365 | 1.0152 | 1.0145 | 0.2874 | 11.66 |
| 1 | 53514066 | rs1120608 | C | G | 2 | 0.3794 | 0.3794 | 1.0247 | 1.0247 | 0.5046 | 0 |
| 1 | 53944037 | rs734908 | T | G | 2 | 0.3794 | 0.3797 | 0.9869 | 0.9869 | 0.3168 | 0.19 |
| 1 | 53808693 | rs1416095 | A | G | 2 | 0.3796 | 0.3796 | 1.0124 | 1.0124 | 0.8145 | 0 |
| 1 | 53513168 | rs1078894 | G | A | 2 | 0.38 | 0.38 | 1.0248 | 1.0248 | 0.5058 | 0 |
| 1 | 53513837 | rs4348672 | G | C | 2 | 0.3802 | 0.3802 | 1.0247 | 1.0247 | 0.5054 | 0 |
| 1 | 53514350 | rs6664654 | C | T | 2 | 0.3802 | 0.3802 | 1.0248 | 1.0248 | 0.5019 | 0 |
| 1 | 54645048 | rs7536714 | A | G | 2 | 0.3802 | 0.3802 | 1.0506 | 1.0506 | 0.3936 | 0 |
| 1 | 53514628 | rs6677541 | A | G | 2 | 0.3803 | 0.3803 | 1.0247 | 1.0247 | 0.5055 | 0 |
| 1 | 54075325 | rs7918437 | C | T | 2 | 0.3803 | 0.3803 | 0.9659 | 0.9659 | 0.4878 | 0 |
| 1 | 53515101 | rs4926946 | C | A | 2 | 0.3804 | 0.3804 | 1.0247 | 1.0247 | 0.5056 | 0 |
| 1 | 53519436 | rs1157765 | A | G | 2 | 0.3808 | 0.3808 | 1.0247 | 1.0247 | 0.4999 | 0 |
| 1 | 53519464 | rs6053790 | C | T | 2 | 0.3809 | 0.3809 | 1.0246 | 1.0246 | 0.5023 | 0 |
| 1 | 53519530 | rs6088595 | G | A | 2 | 0.3809 | 0.3809 | 1.0246 | 1.0246 | 0.5023 | 0 |
| 1 | 53519646 | rs6090587 | G | A | 2 | 0.3809 | 0.3809 | 1.0246 | 1.0246 | 0.5023 | 0 |
| 1 | 53514497 | rs6664845 | C | G | 2 | 0.3811 | 0.3811 | 1.0247 | 1.0247 | 0.5063 | 0 |
| 1 | 53615541 | rs1088876 | C | T | 2 | 0.3811 | 0.3811 | 1.0119 | 1.0119 | 0.5809 | 0 |
| 1 | 54298716 | rs5915657 | C | G | 2 | 0.3811 | 0.3811 | 1.0182 | 1.0182 | 0.5872 | 0 |
| 1 | 54689765 | rs1120630 | C | G | 2 | 0.3811 | 0.3811 | 1.0169 | 1.0169 | 0.415 | 0 |
| 1 | 53570644 | rs1288421 | G | C | 2 | 0.3812 | 0.3812 | 1.0842 | 1.0842 | 0.7141 | 0 |
| 1 | 53600078 | rs3737989 | A | G | 2 | 0.3813 | 0.3813 | 0.9849 | 0.9849 | 0.7106 | 0 |
| 1 | 54644557 | rs1205959 | A | G | 2 | 0.3813 | 0.3813 | 1.0505 | 1.0505 | 0.3931 | 0 |
| 1 | 53518235 | rs1207049 | C | T | 2 | 0.3817 | 0.3817 | 1.0245 | 1.0245 | 0.503 | 0 |
| 1 | 53517401 | rs1209484 | T | C | 2 | 0.3818 | 0.3818 | 1.0245 | 1.0245 | 0.5032 | 0 |
| 1 | 53517654 | rs4282747 | A | G | 2 | 0.3819 | 0.3819 | 1.0245 | 1.0245 | 0.5033 | 0 |
| 1 | 53517997 | rs1710776 | T | C | 2 | 0.3819 | 0.3819 | 1.0245 | 1.0245 | 0.5032 | 0 |
| 1 | 53518049 | rs1209616 | T | G | 2 | 0.3819 | 0.3819 | 1.0245 | 1.0245 | 0.5032 | 0 |
| 1 | 53517698 | rs1710776 | G | A | 2 | 0.382 | 0.382 | 1.0245 | 1.0245 | 0.5033 | 0 |
| 1 | 54616304 | rs7290800 | A | G | 2 | 0.382 | 0.4261 | 1.0216 | 1.0247 | 0.2204 | 33.41 |
| 1 | 54585415 | rs1206342 | T | C | 2 | 0.3822 | 0.3822 | 0.9882 | 0.9882 | 0.4036 | 0 |
| 1 | 53647574 | rs7523046 | T | G | 2 | 0.3825 | 0.3825 | 0.9437 | 0.9437 | 0.555 | 0 |
| 1 | 54129668 | rs1149915 | C | T | 2 | 0.3829 | 0.3829 | 0.9632 | 0.9632 | 0.9603 | 0 |
| 1 | 53619482 | rs7556421 | C | T | 2 | 0.3834 | 0.3834 | 1.0118 | 1.0118 | 0.914 | 0 |
| 1 | 53616916 | rs1088877 | A | G | 2 | 0.3844 | 0.3844 | 1.012 | 1.012 | 0.6167 | 0 |
| 1 | 54298315 | rs1274484 | C | T | 2 | 0.3844 | 0.3844 | 0.9761 | 0.9761 | 0.35 | 0 |
| 1 | 53946829 | rs5984619 | G | A | 2 | 0.3845 | 0.3845 | 1.0182 | 1.0182 | 0.8629 | 0 |
| 1 | 53901342 | rs6672280 | T | C | 2 | 0.3847 | 0.3847 | 0.9871 | 0.9871 | 0.3889 | 0 |
| 1 | 53531275 | rs1043706 | G | A | 2 | 0.3848 | 0.3848 | 1.0249 | 1.0249 | 0.9969 | 0 |
| 1 | 54373668 | rs5611818 | A | G | 2 | 0.3848 | 0.3848 | 1.0325 | 1.0325 | 0.5693 | 0 |
| 1 | 54366817 | rs7621696 | G | A | 2 | 0.385 | 0.385 | 0.9657 | 0.9657 | 0.3927 | 0 |
| 1 | 53835299 | rs6687108 | T | C | 2 | 0.3856 | 0.4683 | 0.9854 | 0.9863 | 0.2655 | 19.35 |
| 1 | 54003207 | rs1273894 | G | A | 2 | 0.3857 | 0.3857 | 1.013 | 1.013 | 0.3412 | 0 |
| 1 | 54171212 | rs1173586 | C | G | 2 | 0.386 | 0.386 | 0.973 | 0.973 | 0.972 | 0 |

| | | | | | | | | | | | |
|---|---|---|---|---|---|---|---|---|---|---|---|
| 1 | 53513768 | rs4599997 | C | G | 2 | 0.3862 | 0.3862 | 1.0244 | 1.0244 | 0.5112 | 0 |
| 1 | 54414500 | rs1207367 | T | C | 2 | 0.3862 | 0.7735 | 1.0163 | 1.0106 | 0.0544 | 72.98 |
| 1 | 54494504 | rs1150029 | T | C | 2 | 0.387 | 0.387 | 1.0632 | 1.0632 | 0.6628 | 0 |
| 1 | 54519158 | rs1120627 | G | A | 2 | 0.3872 | 0.3872 | 0.9668 | 0.9668 | 0.4681 | 0 |
| 1 | 53912522 | rs1202860 | C | A | 2 | 0.3873 | 0.3963 | 0.9688 | 0.9689 | 0.3086 | 3.55 |
| 1 | 54483694 | rs3476495 | C | T | 2 | 0.3873 | 0.3873 | 0.9882 | 0.9882 | 0.3962 | 0 |
| 1 | 54413422 | rs7526168 | C | T | 2 | 0.3877 | 0.7681 | 1.0162 | 1.0107 | 0.0582 | 72.13 |
| 1 | 53610852 | rs2486634 | C | G | 2 | 0.3882 | 0.3882 | 0.9882 | 0.9882 | 0.423 | 0 |
| 1 | 53924150 | rs1288627 | C | T | 2 | 0.3885 | 0.3885 | 0.9871 | 0.9871 | 0.6811 | 0 |
| 1 | 54357986 | rs1078896 | C | T | 2 | 0.3886 | 0.3886 | 0.9741 | 0.9741 | 0.5024 | 0 |
| 1 | 53498221 | rs1710768 | T | C | 2 | 0.3887 | 0.6168 | 1.0566 | 1.0431 | 0.2116 | 35.91 |
| 1 | 53656136 | rs6176959 | T | G | 2 | 0.389 | 0.6013 | 1.0418 | 1.0503 | 0.0497 | 74.04 |
| 1 | 54153829 | rs1158210 | T | A | 2 | 0.389 | 0.389 | 0.9828 | 0.9828 | 0.9359 | 0 |
| 1 | 54210899 | rs1780382 | T | C | 2 | 0.3891 | 0.3891 | 0.9809 | 0.9809 | 0.8288 | 0 |
| 1 | 53775436 | rs1181148 | A | G | 2 | 0.3894 | 0.3894 | 1.0627 | 1.0627 | 0.6096 | 0 |
| 1 | 53652951 | rs1159025 | T | C | 2 | 0.39 | 0.39 | 0.9868 | 0.9868 | 0.9219 | 0 |
| 1 | 54358070 | rs1078897 | G | A | 2 | 0.3903 | 0.3903 | 0.974 | 0.974 | 0.5127 | 0 |
| 1 | 54153169 | rs1240511 | T | C | 2 | 0.3908 | 0.3908 | 0.9828 | 0.9828 | 0.9336 | 0 |
| 1 | 54076977 | rs2948051 | A | G | 2 | 0.3912 | 0.3912 | 1.0136 | 1.0136 | 0.5812 | 0 |
| 1 | 54372124 | rs1208424 | T | A | 2 | 0.3913 | 0.4008 | 0.9802 | 0.9798 | 0.2991 | 7.26 |
| 1 | 53923927 | rs3501530 | G | A | 2 | 0.3914 | 0.3914 | 1.0178 | 1.0178 | 0.9053 | 0 |
| 1 | 54076130 | rs2948050 | A | G | 2 | 0.3915 | 0.3915 | 1.0135 | 1.0135 | 0.5811 | 0 |
| 1 | 54260668 | rs7408441 | C | T | 2 | 0.3917 | 0.701 | 1.0685 | 1.0684 | 0.026 | 79.84 |
| 1 | 54683413 | rs1088884 | A | T | 2 | 0.3917 | 0.3917 | 1.0159 | 1.0159 | 0.484 | 0 |
| 1 | 53938784 | rs6176842 | C | T | 2 | 0.3919 | 0.3919 | 1.0181 | 1.0181 | 0.8023 | 0 |
| 1 | 54077470 | rs2950258 | A | G | 2 | 0.3925 | 0.3925 | 1.0134 | 1.0134 | 0.5739 | 0 |
| 1 | 54078203 | rs2950260 | G | A | 2 | 0.3927 | 0.3927 | 1.0134 | 1.0134 | 0.5693 | 0 |
| 1 | 54016353 | rs1710878 | C | T | 2 | 0.3928 | 0.6285 | 1.0141 | 1.0117 | 0.1485 | 52.1 |
| 1 | 54635702 | rs1710998 | C | T | 2 | 0.393 | 0.393 | 1.0487 | 1.0487 | 0.3762 | 0 |
| 1 | 53531278 | rs1043706 | C | G | 2 | 0.3937 | 0.3937 | 1.0244 | 1.0244 | 0.9958 | 0 |
| 1 | 53995366 | rs6004120 | G | A | 2 | 0.3939 | 0.5859 | 0.9689 | 0.9628 | 0.0622 | 71.25 |
| 1 | 53812211 | rs6675333 | C | T | 2 | 0.394 | 0.394 | 0.9881 | 0.9881 | 0.7915 | 0 |
| 1 | 53601241 | rs4926957 | A | G | 2 | 0.3943 | 0.3943 | 1.0126 | 1.0126 | 0.9271 | 0 |
| 1 | 54207814 | rs797899 | G | C | 2 | 0.3944 | 0.3944 | 0.981 | 0.981 | 0.673 | 0 |
| 1 | 54083213 | rs3006897 | T | C | 2 | 0.3945 | 0.3945 | 1.0132 | 1.0132 | 0.492 | 0 |
| 1 | 53808358 | rs2782495 | C | T | 2 | 0.3947 | 0.3947 | 1.0136 | 1.0136 | 0.6141 | 0 |
| 1 | 53486581 | rs1180495 | G | T | 2 | 0.3956 | 0.3956 | 1.024 | 1.024 | 0.4161 | 0 |
| 1 | 54078584 | rs3006895 | C | T | 2 | 0.3973 | 0.3973 | 1.0132 | 1.0132 | 0.571 | 0 |
| 1 | 53613018 | rs1679936 | G | C | 2 | 0.3976 | 0.3976 | 1.0115 | 1.0115 | 0.5566 | 0 |
| 1 | 54079908 | rs2950264 | G | A | 2 | 0.398 | 0.398 | 1.0134 | 1.0134 | 0.5672 | 0 |
| 1 | 54515041 | rs1088883 | T | C | 2 | 0.3981 | 0.3981 | 0.9675 | 0.9675 | 0.6028 | 0 |
| 1 | 54668779 | rs1203134 | A | G | 2 | 0.3988 | 0.3988 | 1.0156 | 1.0156 | 0.4565 | 0 |
| 1 | 53796783 | rs1235419 | C | T | 2 | 0.3992 | 0.3992 | 0.9269 | 0.9269 | 0.8722 | 0 |
| 1 | 54636277 | rs1181065 | G | A | 2 | 0.3996 | 0.3996 | 1.0483 | 1.0483 | 0.3756 | 0 |
| 1 | 54649589 | rs1204394 | G | A | 2 | 0.3998 | 0.3998 | 1.0142 | 1.0142 | 0.6022 | 0 |
| 1 | 53621628 | rs1564485 | A | T | 2 | 0.3999 | 0.3999 | 1.0113 | 1.0113 | 0.8005 | 0 |
| 1 | 54383336 | rs1152630 | C | T | 2 | 0.4003 | 0.4032 | 0.9808 | 0.9807 | 0.3117 | 2.29 |
| 1 | 54364997 | rs7514037 | T | G | 2 | 0.4004 | 0.4004 | 0.9864 | 0.9864 | 0.9776 | 0 |

| | | | | | | | | | | | |
|---|---|---|---|---|---|---|---|---|---|---|---|
| 1 | 54364809 | rs6588496 | G | A | 2 | 0.401 | 0.401 | 0.9864 | 0.9864 | 0.9188 | 0 |
| 1 | 54364827 | rs6588497 | C | T | 2 | 0.401 | 0.401 | 0.9864 | 0.9864 | 0.9188 | 0 |
| 1 | 53995757 | rs1202561 | T | C | 2 | 0.4013 | 0.4514 | 0.9655 | 0.9641 | 0.2468 | 25.45 |
| 1 | 53938536 | rs1296439 | G | T | 2 | 0.4022 | 0.4022 | 0.9876 | 0.9876 | 0.5144 | 0 |
| 1 | 54543320 | rs1144410 | T | G | 2 | 0.4023 | 0.4023 | 1.0562 | 1.0562 | 0.3629 | 0 |
| 1 | 54074803 | rs3013747 | A | G | 2 | 0.4024 | 0.4024 | 1.0132 | 1.0132 | 0.5644 | 0 |
| 1 | 53513982 | rs2896792 | C | T | 2 | 0.4025 | 0.4025 | 1.0234 | 1.0234 | 0.5434 | 0 |
| 1 | 53528955 | rs7408241 | G | A | 2 | 0.4027 | 0.4027 | 1.0288 | 1.0288 | 0.8431 | 0 |
| 1 | 53817060 | rs1421922 | G | C | 2 | 0.4027 | 0.6898 | 0.9547 | 0.9691 | 0.1867 | 42.64 |
| 1 | 54111511 | rs6177659 | C | T | 2 | 0.4027 | 0.4027 | 0.9572 | 0.9572 | 0.442 | 0 |
| 1 | 54369730 | rs4926616 | T | C | 2 | 0.4027 | 0.4027 | 1.0124 | 1.0124 | 0.4672 | 0 |
| 1 | 53808721 | rs2782496 | G | A | 2 | 0.4029 | 0.4029 | 1.0129 | 1.0129 | 0.5808 | 0 |
| 1 | 53740567 | rs7963897 | A | G | 2 | 0.403 | 0.403 | 1.0561 | 1.0561 | 0.9233 | 0 |
| 1 | 54365396 | rs2284453 | G | A | 2 | 0.403 | 0.403 | 0.9864 | 0.9864 | 0.979 | 0 |
| 1 | 53944180 | rs1288645 | A | G | 2 | 0.4032 | 0.4032 | 0.9876 | 0.9876 | 0.6019 | 0 |
| 1 | 54356721 | rs3546375 | A | T | 2 | 0.4038 | 0.4038 | 0.9751 | 0.9751 | 0.5756 | 0 |
| 1 | 54205759 | rs797902 | T | C | 2 | 0.404 | 0.404 | 0.9814 | 0.9814 | 0.9075 | 0 |
| 1 | 54601021 | rs4927057 | G | A | 2 | 0.4042 | 0.4042 | 1.0185 | 1.0185 | 0.4877 | 0 |
| 1 | 53811942 | rs6698595 | G | T | 2 | 0.4044 | 0.4044 | 0.9884 | 0.9884 | 0.8003 | 0 |
| 1 | 53530945 | rs1043706 | C | T | 2 | 0.4049 | 0.4049 | 1.0238 | 1.0238 | 0.9805 | 0 |
| 1 | 53595504 | rs3766807 | G | A | 2 | 0.4053 | 0.4053 | 1.0154 | 1.0154 | 0.5626 | 0 |
| 1 | 53866793 | rs1205989 | G | C | 2 | 0.4053 | 0.4053 | 0.9848 | 0.9848 | 0.8807 | 0 |
| 1 | 53652962 | rs1157690 | A | G | 2 | 0.4056 | 0.4056 | 0.9872 | 0.9872 | 0.8945 | 0 |
| 1 | 54362985 | rs1883456 | C | T | 2 | 0.4056 | 0.4056 | 0.9867 | 0.9867 | 0.8399 | 0 |
| 1 | 54153757 | rs1120619 | C | T | 2 | 0.4059 | 0.4059 | 0.9834 | 0.9834 | 0.975 | 0 |
| 1 | 53531193 | rs1043705 | T | C | 2 | 0.406 | 0.406 | 1.0238 | 1.0238 | 0.986 | 0 |
| 1 | 53530487 | rs1158464 | C | T | 2 | 0.4064 | 0.4064 | 1.0238 | 1.0238 | 0.9198 | 0 |
| 1 | 54365679 | rs1088881 | G | A | 2 | 0.4065 | 0.4065 | 0.9865 | 0.9865 | 0.9792 | 0 |
| 1 | 54559794 | rs1748182 | T | C | 2 | 0.4075 | 0.4075 | 0.9827 | 0.9827 | 0.4168 | 0 |
| 1 | 54362300 | rs6684508 | C | T | 2 | 0.4076 | 0.4076 | 0.9868 | 0.9868 | 0.8424 | 0 |
| 1 | 54043703 | rs1240855 | T | C | 2 | 0.4077 | 0.5565 | 0.9864 | 0.9881 | 0.2278 | 31.25 |
| 1 | 53772897 | rs7529264 | A | G | 2 | 0.4078 | 0.4078 | 1.0575 | 1.0575 | 0.7501 | 0 |
| 1 | 53531698 | rs1208169 | C | T | 2 | 0.4079 | 0.4079 | 1.0283 | 1.0283 | 0.8203 | 0 |
| 1 | 53643797 | rs1180982 | G | A | 2 | 0.4079 | 0.4079 | 0.9464 | 0.9464 | 0.603 | 0 |
| 1 | 54599426 | rs645980 | G | A | 2 | 0.4079 | 0.4079 | 1.0394 | 1.0394 | 0.9517 | 0 |
| 1 | 54366902 | rs7520300 | T | A | 2 | 0.4081 | 0.4081 | 0.9865 | 0.9865 | 0.9776 | 0 |
| 1 | 54076357 | rs3006894 | C | T | 2 | 0.4084 | 0.4084 | 1.0129 | 1.0129 | 0.5586 | 0 |
| 1 | 54079685 | rs2950263 | A | T | 2 | 0.4085 | 0.4085 | 1.0128 | 1.0128 | 0.5547 | 0 |
| 1 | 53528929 | rs1256570 | C | T | 2 | 0.4087 | 0.4087 | 1.0161 | 1.0161 | 0.622 | 0 |
| 1 | 53938695 | rs1288640 | T | C | 2 | 0.409 | 0.409 | 0.9877 | 0.9877 | 0.5242 | 0 |
| 1 | 54353744 | rs1429951 | C | T | 2 | 0.4091 | 0.4091 | 1.0225 | 1.0225 | 0.843 | 0 |
| 1 | 54208626 | rs702487 | C | T | 2 | 0.4093 | 0.4093 | 0.9817 | 0.9817 | 0.8705 | 0 |
| 1 | 54073480 | rs2950257 | G | C | 2 | 0.4095 | 0.4095 | 1.013 | 1.013 | 0.5783 | 0 |
| 1 | 54360819 | rs6662835 | T | C | 2 | 0.4095 | 0.4095 | 0.9866 | 0.9866 | 0.8794 | 0 |
| 1 | 54208046 | rs797898 | C | T | 2 | 0.4099 | 0.4099 | 0.9818 | 0.9818 | 0.8828 | 0 |
| 1 | 53937177 | rs1296438 | A | G | 2 | 0.4106 | 0.4106 | 0.9878 | 0.9878 | 0.6013 | 0 |
| 1 | 53741896 | rs6176962 | A | G | 2 | 0.4107 | 0.4107 | 1.0552 | 1.0552 | 0.9275 | 0 |
| 1 | 54345595 | rs1274357 | C | T | 2 | 0.4107 | 0.4107 | 0.9524 | 0.9524 | 0.4307 | 0 |

| | | | | | | | | | | | |
|---|---|---|---|---|---|---|---|---|---|---|---|
| 1 | 54166530 | rs1173592 | T | A | 2 | 0.4108 | 0.4108 | 0.9745 | 0.9745 | 0.8564 | 0 |
| 1 | 54074729 | rs2948048 | G | A | 2 | 0.4109 | 0.4109 | 1.0129 | 1.0129 | 0.5703 | 0 |
| 1 | 54595926 | rs8011081 | C | T | 2 | 0.411 | 0.411 | 1.0846 | 1.0846 | 0.859 | 0 |
| 1 | 54365530 | rs1120624 | A | G | 2 | 0.4115 | 0.4115 | 0.9867 | 0.9867 | 0.9933 | 0 |
| 1 | 54074362 | rs3013748 | A | T | 2 | 0.4125 | 0.4125 | 1.0128 | 1.0128 | 0.5873 | 0 |
| 1 | 54360957 | rs6671330 | G | T | 2 | 0.4129 | 0.4129 | 0.9867 | 0.9867 | 0.8722 | 0 |
| 1 | 54361022 | rs1207069 | G | A | 2 | 0.4132 | 0.4132 | 0.9867 | 0.9867 | 0.8721 | 0 |
| 1 | 53617071 | rs1088877 | G | A | 2 | 0.4134 | 0.4134 | 1.0113 | 1.0113 | 0.5865 | 0 |
| 1 | 54373366 | rs1202321 | C | A | 2 | 0.4136 | 0.4136 | 1.0296 | 1.0296 | 0.4023 | 0 |
| 1 | 54672275 | rs1337445 | G | T | 2 | 0.4142 | 0.4142 | 1.0118 | 1.0118 | 0.8778 | 0 |
| 1 | 53534257 | rs4926949 | T | G | 2 | 0.4147 | 0.4147 | 1.0278 | 1.0278 | 0.9925 | 0 |
| 1 | 53606392 | rs4926959 | T | C | 2 | 0.4149 | 0.4149 | 1.0113 | 1.0113 | 0.6249 | 0 |
| 1 | 53638218 | rs4926962 | T | C | 2 | 0.4153 | 0.4153 | 1.0121 | 1.0121 | 0.8046 | 0 |
| 1 | 53992515 | rs554798 | C | T | 2 | 0.4155 | 0.5885 | 0.9618 | 0.9665 | 0.1928 | 41.05 |
| 1 | 53592123 | rs1383599 | C | G | 2 | 0.4162 | 0.4162 | 1.0462 | 1.0462 | 0.7855 | 0 |
| 1 | 53711735 | rs5177 | G | C | 2 | 0.4162 | 0.4162 | 1.0112 | 1.0112 | 0.8087 | 0 |
| 1 | 54522830 | rs7602787 | A | G | 2 | 0.4162 | 0.4162 | 0.9688 | 0.9688 | 0.5362 | 0 |
| 1 | 53532609 | rs1158009 | T | C | 2 | 0.4164 | 0.4164 | 1.0278 | 1.0278 | 0.8509 | 0 |
| 1 | 54563522 | rs7538004 | G | A | 2 | 0.4165 | 0.4165 | 0.9689 | 0.9689 | 0.4791 | 0 |
| 1 | 54363088 | rs1883458 | G | A | 2 | 0.4166 | 0.4166 | 0.987 | 0.987 | 0.8527 | 0 |
| 1 | 54508394 | rs1443102 | A | G | 2 | 0.4167 | 0.4167 | 1.059 | 1.059 | 0.681 | 0 |
| 1 | 54021144 | rs525568 | G | A | 2 | 0.417 | 0.5561 | 0.9888 | 0.9834 | 0.0465 | 74.78 |
| 1 | 54441275 | rs3765404 | C | T | 2 | 0.4171 | 0.6348 | 0.9119 | 0.9331 | 0.2363 | 28.71 |
| 1 | 53810084 | rs1088877 | A | G | 2 | 0.4172 | 0.4172 | 0.9882 | 0.9882 | 0.5507 | 0 |
| 1 | 53576497 | rs1288370 | G | A | 2 | 0.4175 | 0.4175 | 1.0134 | 1.0134 | 0.425 | 0 |
| 1 | 54492493 | rs5618647 | C | T | 2 | 0.4177 | 0.4177 | 0.9688 | 0.9688 | 0.6344 | 0 |
| 1 | 54362771 | rs1883455 | C | T | 2 | 0.4178 | 0.4178 | 0.987 | 0.987 | 0.8584 | 0 |
| 1 | 54015479 | rs6177026 | A | T | 2 | 0.4179 | 0.4179 | 1.0135 | 1.0135 | 0.414 | 0 |
| 1 | 54361639 | rs2268184 | T | A | 2 | 0.4181 | 0.4181 | 0.9869 | 0.9869 | 0.8672 | 0 |
| 1 | 54605095 | rs625370 | G | A | 2 | 0.4186 | 0.4186 | 1.0364 | 1.0364 | 0.9718 | 0 |
| 1 | 53885963 | rs7555252 | T | G | 2 | 0.4192 | 0.4192 | 0.9879 | 0.9879 | 0.5301 | 0 |
| 1 | 53955249 | rs1204524 | C | T | 2 | 0.4195 | 0.6398 | 0.9725 | 0.9736 | 0.0982 | 63.43 |
| 1 | 54354935 | rs3817871 | G | C | 2 | 0.4199 | 0.4199 | 0.9869 | 0.9869 | 0.9198 | 0 |
| 1 | 54361619 | rs2268183 | T | C | 2 | 0.4201 | 0.4201 | 0.9869 | 0.9869 | 0.8651 | 0 |
| 1 | 54503729 | rs1209534 | T | A | 2 | 0.4207 | 0.4207 | 0.9689 | 0.9689 | 0.6148 | 0 |
| 1 | 54365962 | rs2284454 | A | G | 2 | 0.4209 | 0.4209 | 0.9869 | 0.9869 | 0.9936 | 0 |
| 1 | 54203608 | rs797905 | C | T | 2 | 0.4211 | 0.4211 | 0.9821 | 0.9821 | 0.7614 | 0 |
| 1 | 53764602 | rs7289536 | C | G | 2 | 0.4218 | 0.4218 | 0.9792 | 0.9792 | 0.3423 | 0 |
| 1 | 53813087 | rs7922790 | A | G | 2 | 0.4218 | 0.4218 | 1.0215 | 1.0215 | 0.8465 | 0 |
| 1 | 54366479 | rs7527713 | G | A | 2 | 0.4219 | 0.4219 | 0.987 | 0.987 | 0.9906 | 0 |
| 1 | 53834375 | rs4926983 | C | A | 2 | 0.422 | 0.422 | 0.9862 | 0.9862 | 0.3886 | 0 |
| 1 | 53740854 | rs7447644 | A | G | 2 | 0.4222 | 0.4798 | 0.9681 | 0.9568 | 0.1415 | 53.74 |
| 1 | 53900842 | rs1766642 | C | T | 2 | 0.4231 | 0.4231 | 0.9598 | 0.9598 | 0.5504 | 0 |
| 1 | 53808588 | rs1416097 | T | C | 2 | 0.4233 | 0.4233 | 1.0112 | 1.0112 | 0.7286 | 0 |
| 1 | 53636177 | rs4543741 | G | C | 2 | 0.4234 | 0.4234 | 1.0122 | 1.0122 | 0.7405 | 0 |
| 1 | 53840245 | rs7267512 | C | T | 2 | 0.4238 | 0.4238 | 1.045 | 1.045 | 0.8192 | 0 |
| 1 | 54602648 | rs913811 | C | T | 2 | 0.424 | 0.424 | 1.0174 | 1.0174 | 0.4674 | 0 |
| 1 | 53728448 | rs1154533 | G | A | 2 | 0.4241 | 0.4241 | 0.9592 | 0.9592 | 0.7246 | 0 |

| | | | | | | | | | | | |
|---|---|---|---|---|---|---|---|---|---|---|---|
| 1 | 54509786 | rs1444730 | C | T | 2 | 0.4241 | 0.4241 | 1.058 | 1.058 | 0.6919 | 0 |
| 1 | 54417386 | rs6588499 | A | C | 2 | 0.4242 | 0.8073 | 1.0149 | 1.0091 | 0.0489 | 74.22 |
| 1 | 53790538 | rs1889541 | G | C | 2 | 0.4247 | 0.4247 | 1.0117 | 1.0117 | 0.673 | 0 |
| 1 | 54363434 | rs6663024 | G | A | 2 | 0.4247 | 0.4247 | 0.987 | 0.987 | 0.8924 | 0 |
| 1 | 54399825 | rs7980010 | C | T | 2 | 0.4248 | 0.4248 | 0.9818 | 0.9818 | 0.3177 | 0 |
| 1 | 54683856 | rs15921 | C | G | 2 | 0.4248 | 0.4455 | 1.0126 | 1.0134 | 0.2736 | 16.58 |
| 1 | 54608323 | rs7616215 | G | T | 2 | 0.425 | 0.425 | 0.9105 | 0.9105 | 0.687 | 0 |
| 1 | 53527996 | rs5605492 | A | G | 2 | 0.4253 | 0.4253 | 1.0155 | 1.0155 | 0.6398 | 0 |
| 1 | 54018932 | rs501339 | G | A | 2 | 0.4253 | 0.5018 | 1.0707 | 1.1408 | 0.0366 | 77.12 |
| 1 | 54400259 | rs1161249 | C | T | 2 | 0.4255 | 0.4255 | 0.9671 | 0.9671 | 0.9004 | 0 |
| 1 | 54364100 | rs1275023 | G | A | 2 | 0.4256 | 0.4256 | 0.9871 | 0.9871 | 0.954 | 0 |
| 1 | 54487146 | rs1203316 | G | A | 2 | 0.4258 | 0.63 | 0.989 | 0.9909 | 0.179 | 44.62 |
| 1 | 53934091 | rs1288637 | A | G | 2 | 0.426 | 0.426 | 0.9882 | 0.9882 | 0.621 | 0 |
| 1 | 53473973 | rs6694402 | C | A | 2 | 0.4268 | 0.5404 | 1.0191 | 1.0216 | 0.1452 | 52.88 |
| 1 | 53877199 | rs1120615 | G | T | 2 | 0.4268 | 0.4268 | 1.0149 | 1.0149 | 0.8934 | 0 |
| 1 | 53810840 | rs4926594 | A | G | 2 | 0.4271 | 0.4271 | 0.9885 | 0.9885 | 0.525 | 0 |
| 1 | 54341967 | rs7610391 | G | A | 2 | 0.4274 | 0.5186 | 0.9612 | 0.9584 | 0.189 | 42.04 |
| 1 | 54617611 | rs7862911 | C | T | 2 | 0.4277 | 0.4536 | 0.9759 | 0.9757 | 0.2852 | 12.45 |
| 1 | 54156255 | rs7786023 | G | A | 2 | 0.4292 | 0.4292 | 1.0261 | 1.0261 | 0.9975 | 0 |
| 1 | 54039864 | rs3013759 | C | T | 2 | 0.4293 | 0.4293 | 0.9331 | 0.9331 | 0.4268 | 0 |
| 1 | 53932181 | rs7807058 | A | G | 2 | 0.4294 | 0.4294 | 1.0164 | 1.0164 | 0.9487 | 0 |
| 1 | 53864086 | rs1275872 | C | T | 2 | 0.4296 | 0.4296 | 1.0264 | 1.0264 | 0.3352 | 0 |
| 1 | 54421597 | rs1120625 | G | A | 2 | 0.43 | 0.7979 | 1.0149 | 1.0092 | 0.0614 | 71.41 |
| 1 | 54321013 | rs7551454 | G | A | 2 | 0.4302 | 0.4934 | 0.9816 | 0.9764 | 0.1529 | 51.04 |
| 1 | 54512583 | rs1412824 | G | A | 2 | 0.4302 | 0.4302 | 1.0573 | 1.0573 | 0.6996 | 0 |
| 1 | 54590416 | rs3426308 | C | T | 2 | 0.4303 | 0.4303 | 1.0614 | 1.0614 | 0.6397 | 0 |
| 1 | 54400674 | rs1120625 | A | G | 2 | 0.4306 | 0.4306 | 0.982 | 0.982 | 0.3183 | 0 |
| 1 | 53793170 | rs1933534 | C | T | 2 | 0.4309 | 0.4309 | 1.0115 | 1.0115 | 0.6523 | 0 |
| 1 | 54040118 | rs3006896 | G | A | 2 | 0.4309 | 0.4309 | 0.9334 | 0.9334 | 0.4218 | 0 |
| 1 | 54156204 | rs1138494 | C | T | 2 | 0.4316 | 0.4316 | 1.0259 | 1.0259 | 0.9952 | 0 |
| 1 | 53874444 | rs7267515 | G | A | 2 | 0.4321 | 0.4321 | 1.0144 | 1.0144 | 0.9236 | 0 |
| 1 | 53677119 | rs8033267 | A | C | 2 | 0.4322 | 0.4322 | 1.0671 | 1.0671 | 0.7558 | 0 |
| 1 | 53944665 | rs1288646 | A | G | 2 | 0.4342 | 0.4342 | 0.9884 | 0.9884 | 0.6442 | 0 |
| 1 | 53611040 | rs2806272 | T | A | 2 | 0.4345 | 0.4345 | 0.9893 | 0.9893 | 0.3861 | 0 |
| 1 | 53702840 | rs6176961 | T | C | 2 | 0.4349 | 0.4349 | 1.0496 | 1.0496 | 0.9341 | 0 |
| 1 | 53676401 | rs2229291 | T | G | 2 | 0.435 | 0.435 | 0.9346 | 0.9346 | 0.7694 | 0 |
| 1 | 53772544 | rs1288489 | C | G | 2 | 0.435 | 0.435 | 1.0132 | 1.0132 | 0.6939 | 0 |
| 1 | 54641579 | rs611903 | C | T | 2 | 0.4351 | 0.4351 | 1.0106 | 1.0106 | 0.4791 | 0 |
| 1 | 54617654 | rs5577792 | T | G | 2 | 0.436 | 0.436 | 1.0538 | 1.0538 | 0.8437 | 0 |
| 1 | 54456151 | rs1459454 | T | A | 2 | 0.4362 | 0.4362 | 1.0349 | 1.0349 | 0.5284 | 0 |
| 1 | 54522280 | rs5748696 | G | A | 2 | 0.4362 | 0.4362 | 0.97 | 0.97 | 0.5168 | 0 |
| 1 | 53813085 | rs1381839 | T | C | 2 | 0.4367 | 0.4367 | 1.0209 | 1.0209 | 0.9187 | 0 |
| 1 | 53626626 | rs7354975 | A | T | 2 | 0.4369 | 0.4369 | 0.9407 | 0.9407 | 0.7843 | 0 |
| 1 | 54528944 | rs7266413 | C | T | 2 | 0.4374 | 0.4374 | 0.9698 | 0.9698 | 0.5054 | 0 |
| 1 | 53775740 | rs1288486 | C | T | 2 | 0.4378 | 0.4378 | 1.0131 | 1.0131 | 0.6487 | 0 |
| 1 | 54173953 | rs2950246 | A | G | 2 | 0.4378 | 0.4378 | 0.9754 | 0.9754 | 0.9119 | 0 |
| 1 | 54662035 | rs6668490 | T | A | 2 | 0.4384 | 0.4384 | 1.0495 | 1.0495 | 0.4041 | 0 |
| 1 | 54426808 | rs1205732 | C | T | 2 | 0.4385 | 0.8007 | 1.0146 | 1.009 | 0.0612 | 71.46 |

| | | | | | | | | | | | |
|---|---|---|---|---|---|---|---|---|---|---|---|
| 1 | 53594011 | rs3766802 | T | C | 2 | 0.4392 | 0.4392 | 0.9114 | 0.9114 | 0.8403 | 0 |
| 1 | 54672542 | rs1121537 | G | A | 2 | 0.4395 | 0.4395 | 1.04 | 1.04 | 0.9281 | 0 |
| 1 | 54588110 | rs7516442 | C | T | 2 | 0.4399 | 0.4399 | 0.9893 | 0.9893 | 0.6169 | 0 |
| 1 | 54354658 | rs2294508 | T | G | 2 | 0.4403 | 0.4403 | 0.9875 | 0.9875 | 0.9639 | 0 |
| 1 | 53537266 | rs1769316 | G | A | 2 | 0.4409 | 0.5721 | 1.0107 | 1.0093 | 0.2461 | 25.68 |
| 1 | 53489448 | rs7512539 | G | A | 2 | 0.441 | 0.506 | 1.0154 | 1.0171 | 0.2019 | 38.61 |
| 1 | 54359022 | rs7418835 | A | G | 2 | 0.441 | 0.441 | 0.9875 | 0.9875 | 0.932 | 0 |
| 1 | 54355208 | rs1120623 | G | A | 2 | 0.4422 | 0.4422 | 0.9875 | 0.9875 | 0.955 | 0 |
| 1 | 54356106 | rs6656722 | G | A | 2 | 0.4422 | 0.4422 | 0.9875 | 0.9875 | 0.9547 | 0 |
| 1 | 54355840 | rs2272931 | C | T | 2 | 0.4423 | 0.4423 | 0.9875 | 0.9875 | 0.9548 | 0 |
| 1 | 54355943 | rs6702765 | G | A | 2 | 0.4423 | 0.4423 | 0.9875 | 0.9875 | 0.9548 | 0 |
| 1 | 54356002 | rs6702862 | G | T | 2 | 0.4423 | 0.4423 | 0.9875 | 0.9875 | 0.9547 | 0 |
| 1 | 53860680 | rs7699888 | T | C | 2 | 0.4427 | 0.4427 | 0.988 | 0.988 | 0.9178 | 0 |
| 1 | 54163854 | rs1780394 | T | C | 2 | 0.4427 | 0.4427 | 0.9761 | 0.9761 | 0.8702 | 0 |
| 1 | 54355041 | rs1088881 | A | G | 2 | 0.443 | 0.443 | 0.9876 | 0.9876 | 0.9575 | 0 |
| 1 | 54161679 | rs3013764 | C | G | 2 | 0.4432 | 0.4432 | 0.9762 | 0.9762 | 0.9083 | 0 |
| 1 | 53720723 | rs1211650 | T | C | 2 | 0.4434 | 0.4004 | 0.9693 | 0.8571 | 0.0679 | 69.99 |
| 1 | 54087127 | rs3006901 | C | T | 2 | 0.4438 | 0.4438 | 1.0391 | 1.0391 | 0.4577 | 0 |
| 1 | 54583318 | rs6656731 | A | T | 2 | 0.4439 | 0.4439 | 0.975 | 0.975 | 0.358 | 0 |
| 1 | 53748888 | rs8027222 | G | A | 2 | 0.4441 | 0.4441 | 1.0447 | 1.0447 | 0.6427 | 0 |
| 1 | 54198318 | rs797911 | A | G | 2 | 0.4444 | 0.4444 | 0.9481 | 0.9481 | 0.4021 | 0 |
| 1 | 54019131 | rs503166 | A | G | 2 | 0.4462 | 0.5618 | 0.9895 | 0.9838 | 0.0508 | 73.78 |
| 1 | 53576221 | rs7408559 | G | T | 2 | 0.4465 | 0.4465 | 0.9102 | 0.9102 | 0.877 | 0 |
| 1 | 53797580 | rs2782491 | A | C | 2 | 0.4467 | 0.4467 | 1.0109 | 1.0109 | 0.7598 | 0 |
| 1 | 54336046 | rs1212907 | A | C | 2 | 0.4471 | 0.4471 | 1.0105 | 1.0105 | 0.4382 | 0 |
| 1 | 54619846 | rs644790 | T | A | 2 | 0.4472 | 0.4857 | 1.0235 | 1.0237 | 0.2713 | 17.36 |
| 1 | 53940255 | rs1288642 | T | C | 2 | 0.4476 | 0.7642 | 0.9782 | 0.986 | 0.1181 | 59.06 |
| 1 | 54046556 | rs941129 | T | C | 2 | 0.4476 | 0.5657 | 0.9876 | 0.9889 | 0.2473 | 25.29 |
| 1 | 53598161 | rs1288357 | A | G | 2 | 0.4481 | 0.4851 | 0.9792 | 0.9748 | 0.2013 | 38.75 |
| 1 | 53793511 | rs4926972 | T | C | 2 | 0.4481 | 0.4481 | 1.0108 | 1.0108 | 0.688 | 0 |
| 1 | 53662823 | rs7267312 | C | A | 2 | 0.4485 | 0.4485 | 0.9874 | 0.9874 | 0.8445 | 0 |
| 1 | 53941519 | rs1288644 | A | G | 2 | 0.4492 | 0.4492 | 0.9888 | 0.9888 | 0.5784 | 0 |
| 1 | 53998139 | rs1203361 | C | A | 2 | 0.4492 | 0.6119 | 0.9722 | 0.9664 | 0.0728 | 68.92 |
| 1 | 53776341 | rs1288484 | A | G | 2 | 0.4494 | 0.4494 | 1.0128 | 1.0128 | 0.7681 | 0 |
| 1 | 54473616 | rs7266411 | A | T | 2 | 0.4494 | 0.4494 | 0.9712 | 0.9712 | 0.5155 | 0 |
| 1 | 53713549 | rs1120612 | G | A | 2 | 0.4497 | 0.4497 | 1.0103 | 1.0103 | 0.986 | 0 |
| 1 | 54600710 | rs1120628 | C | T | 2 | 0.4498 | 0.4498 | 1.0124 | 1.0124 | 0.8445 | 0 |
| 1 | 53775770 | rs1288485 | G | C | 2 | 0.4503 | 0.4503 | 1.0128 | 1.0128 | 0.6972 | 0 |
| 1 | 53602831 | rs1212402 | T | C | 2 | 0.4511 | 0.4511 | 1.0105 | 1.0105 | 0.6197 | 0 |
| 1 | 54155404 | rs3013773 | C | T | 2 | 0.4515 | 0.4515 | 0.9777 | 0.9777 | 0.7383 | 0 |
| 1 | 53952532 | rs7552078 | G | A | 2 | 0.452 | 0.452 | 0.9281 | 0.9281 | 0.897 | 0 |
| 1 | 54358971 | rs7412364 | G | A | 2 | 0.4526 | 0.4526 | 0.9878 | 0.9878 | 0.9198 | 0 |
| 1 | 54496480 | rs7526578 | G | C | 2 | 0.4526 | 0.4526 | 0.971 | 0.971 | 0.5976 | 0 |
| 1 | 53611451 | rs1679931 | G | A | 2 | 0.4528 | 0.4528 | 0.9897 | 0.9897 | 0.4283 | 0 |
| 1 | 53590871 | rs1769309 | G | A | 2 | 0.4531 | 0.4531 | 0.9892 | 0.9892 | 0.3681 | 0 |
| 1 | 53516017 | rs1710775 | G | C | 2 | 0.4534 | 0.4534 | 1.0254 | 1.0254 | 0.963 | 0 |
| 1 | 54039019 | rs3108389 | A | G | 2 | 0.4534 | 0.4534 | 0.9366 | 0.9366 | 0.4083 | 0 |
| 1 | 53929940 | rs1288633 | C | G | 2 | 0.4535 | 0.4535 | 0.9889 | 0.9889 | 0.6398 | 0 |

| | | | | | | | | | | | |
|---|---|---|---|---|---|---|---|---|---|---|---|
| 1 | 53929463 | rs1108989 | T | A | 2 | 0.4543 | 0.4543 | 0.9889 | 0.9889 | 0.6442 | 0 |
| 1 | 54093506 | rs4927020 | G | C | 2 | 0.4543 | 0.4543 | 0.9898 | 0.9898 | 0.4346 | 0 |
| 1 | 53738386 | rs4129475 | G | A | 2 | 0.4549 | 0.5066 | 0.9702 | 0.9584 | 0.1328 | 55.74 |
| 1 | 54333594 | rs3766468 | C | T | 2 | 0.4549 | 0.4549 | 1.0103 | 1.0103 | 0.4316 | 0 |
| 1 | 54574676 | rs5973978 | G | C | 2 | 0.4552 | 0.4552 | 1.0249 | 1.0249 | 0.9546 | 0 |
| 1 | 54357659 | rs1074969 | G | A | 2 | 0.4555 | 0.4555 | 0.9879 | 0.9879 | 0.9214 | 0 |
| 1 | 53519508 | rs5979807 | C | T | 2 | 0.4558 | 0.4558 | 1.0174 | 1.0174 | 0.6052 | 0 |
| 1 | 53947568 | rs1202709 | T | C | 2 | 0.4559 | 0.6354 | 0.9747 | 0.9755 | 0.1274 | 56.96 |
| 1 | 53929536 | rs1288632 | C | G | 2 | 0.4561 | 0.4561 | 0.989 | 0.989 | 0.6423 | 0 |
| 1 | 54011053 | rs6177026 | G | T | 2 | 0.4565 | 0.4662 | 1.0129 | 1.0127 | 0.3125 | 1.98 |
| 1 | 53864019 | rs1737103 | G | A | 2 | 0.4566 | 0.4566 | 0.9853 | 0.9853 | 0.6193 | 0 |
| 1 | 54199484 | rs1207552 | C | T | 2 | 0.4567 | 0.4567 | 0.9824 | 0.9824 | 0.9328 | 0 |
| 1 | 54337972 | rs1158199 | C | A | 2 | 0.4567 | 0.4567 | 1.0102 | 1.0102 | 0.4348 | 0 |
| 1 | 53637106 | rs3125253 | T | C | 2 | 0.458 | 0.458 | 1.0111 | 1.0111 | 0.8315 | 0 |
| 1 | 54588140 | rs7528149 | A | G | 2 | 0.4581 | 0.4581 | 0.9899 | 0.9899 | 0.6136 | 0 |
| 1 | 54432522 | rs2294515 | T | C | 2 | 0.4583 | 0.8374 | 1.0139 | 1.0078 | 0.0472 | 74.6 |
| 1 | 53603072 | rs2885094 | A | G | 2 | 0.459 | 0.459 | 1.0106 | 1.0106 | 0.4405 | 0 |
| 1 | 53479032 | rs1132279 | C | A | 2 | 0.4602 | 0.5938 | 1.0178 | 1.0226 | 0.0834 | 66.63 |
| 1 | 53574076 | rs1679938 | G | A | 2 | 0.4604 | 0.4604 | 0.9881 | 0.9881 | 0.6958 | 0 |
| 1 | 53942583 | rs1158109 | A | G | 2 | 0.4605 | 0.4605 | 1.015 | 1.015 | 0.9826 | 0 |
| 1 | 53808658 | rs1416096 | C | T | 2 | 0.4606 | 0.4606 | 1.0106 | 1.0106 | 0.5089 | 0 |
| 1 | 53955109 | rs1459808 | G | A | 2 | 0.4608 | 0.4608 | 0.9292 | 0.9292 | 0.9111 | 0 |
| 1 | 54604553 | rs633861 | G | A | 2 | 0.461 | 0.461 | 1.0333 | 1.0333 | 0.8044 | 0 |
| 1 | 53534588 | rs4926950 | C | T | 2 | 0.4612 | 0.4612 | 1.0252 | 1.0252 | 0.9466 | 0 |
| 1 | 53573320 | rs2790428 | G | C | 2 | 0.4613 | 0.4613 | 0.9881 | 0.9881 | 0.7445 | 0 |
| 1 | 53849881 | rs7549726 | C | T | 2 | 0.4615 | 0.4916 | 1.0313 | 1.0312 | 0.2871 | 11.76 |
| 1 | 54358462 | rs1207191 | T | G | 2 | 0.4615 | 0.4615 | 0.988 | 0.988 | 0.9157 | 0 |
| 1 | 54041799 | rs7267519 | G | A | 2 | 0.4616 | 0.6457 | 0.9879 | 0.9901 | 0.2027 | 38.37 |
| 1 | 54303630 | rs7408442 | C | A | 2 | 0.4619 | 0.7277 | 1.0579 | 1.061 | 0.0262 | 79.77 |
| 1 | 53611144 | rs1679930 | C | T | 2 | 0.4627 | 0.4627 | 0.9899 | 0.9899 | 0.4065 | 0 |
| 1 | 53916576 | rs1204666 | G | A | 2 | 0.4632 | 0.4632 | 1.0139 | 1.0139 | 0.4265 | 0 |
| 1 | 53611453 | rs1679932 | C | T | 2 | 0.4634 | 0.4634 | 0.9899 | 0.9899 | 0.4362 | 0 |
| 1 | 53904986 | rs7461654 | C | G | 2 | 0.4641 | 0.4641 | 0.9738 | 0.9738 | 0.4186 | 0 |
| 1 | 53776386 | rs1288483 | C | T | 2 | 0.4644 | 0.4644 | 1.0124 | 1.0124 | 0.7537 | 0 |
| 1 | 53856277 | rs1288585 | A | G | 2 | 0.4645 | 0.4645 | 0.99 | 0.99 | 0.6413 | 0 |
| 1 | 53997336 | rs4927001 | G | A | 2 | 0.4654 | 0.6251 | 0.9732 | 0.9671 | 0.0679 | 70 |
| 1 | 53946718 | rs1202773 | G | A | 2 | 0.4655 | 0.628 | 0.9751 | 0.9758 | 0.1432 | 53.34 |
| 1 | 53855223 | rs1766635 | A | C | 2 | 0.466 | 0.466 | 0.9901 | 0.9901 | 0.8126 | 0 |
| 1 | 53927649 | rs943521 | C | A | 2 | 0.4661 | 0.4661 | 0.9892 | 0.9892 | 0.6572 | 0 |
| 1 | 53637740 | rs3125254 | T | C | 2 | 0.4663 | 0.4663 | 1.011 | 1.011 | 0.9684 | 0 |
| 1 | 53594949 | rs7809757 | G | A | 2 | 0.4664 | 0.4664 | 0.9165 | 0.9165 | 0.8534 | 0 |
| 1 | 53958663 | rs1162865 | T | G | 2 | 0.4664 | 0.5445 | 0.9667 | 0.9595 | 0.149 | 51.99 |
| 1 | 54640833 | rs1385041 | C | T | 2 | 0.4665 | 0.4665 | 1.0618 | 1.0618 | 0.8571 | 0 |
| 1 | 54547324 | rs1146359 | C | T | 2 | 0.4679 | 0.4679 | 1.0522 | 1.0522 | 0.6485 | 0 |
| 1 | 53874063 | rs1203984 | C | T | 2 | 0.4682 | 0.4682 | 1.0134 | 1.0134 | 0.8312 | 0 |
| 1 | 54208835 | rs797896 | C | T | 2 | 0.4682 | 0.4682 | 0.9839 | 0.9839 | 0.8399 | 0 |
| 1 | 53874346 | rs1203994 | C | A | 2 | 0.4684 | 0.4684 | 1.0134 | 1.0134 | 0.8518 | 0 |
| 1 | 54281558 | rs1206994 | T | C | 2 | 0.4687 | 0.7292 | 1.0559 | 1.0585 | 0.0287 | 79.11 |

| | | | | | | | | | | | |
|---|---|---|---|---|---|---|---|---|---|---|---|
| 1 | 54076442 | rs7793983 | G | A | 2 | 0.4691 | 0.4691 | 1.0366 | 1.0366 | 0.4364 | 0 |
| 1 | 53808497 | rs1416098 | A | G | 2 | 0.4694 | 0.4694 | 1.0097 | 1.0097 | 0.4972 | 0 |
| 1 | 54661234 | rs7974381 | C | T | 2 | 0.4694 | 0.4694 | 1.0388 | 1.0388 | 0.6991 | 0 |
| 1 | 54368253 | rs731828 | A | C | 2 | 0.4696 | 0.4696 | 1.0103 | 1.0103 | 0.6751 | 0 |
| 1 | 54511349 | rs5589703 | C | G | 2 | 0.4699 | 0.9202 | 1.0481 | 1.014 | 0.0431 | 75.55 |
| 1 | 54010026 | rs6654482 | G | C | 2 | 0.4709 | 0.4709 | 0.9886 | 0.9886 | 0.3636 | 0 |
| 1 | 54080244 | rs4927016 | A | G | 2 | 0.4712 | 0.7982 | 0.9883 | 0.993 | 0.1061 | 61.71 |
| 1 | 53602687 | rs1120610 | A | G | 2 | 0.4721 | 0.4721 | 1.0098 | 1.0098 | 0.6013 | 0 |
| 1 | 54331715 | rs1883452 | C | G | 2 | 0.4722 | 0.4722 | 1.01 | 1.01 | 0.4649 | 0 |
| 1 | 54583692 | rs1120628 | T | C | 2 | 0.4722 | 0.4722 | 0.9787 | 0.9787 | 0.5932 | 0 |
| 1 | 54210123 | rs797895 | G | C | 2 | 0.4726 | 0.4726 | 0.9841 | 0.9841 | 0.82 | 0 |
| 1 | 53797690 | rs2788038 | C | A | 2 | 0.4728 | 0.4728 | 1.011 | 1.011 | 0.6126 | 0 |
| 1 | 54247271 | rs1569783 | C | G | 2 | 0.4728 | 0.4728 | 1.0145 | 1.0145 | 0.7767 | 0 |
| 1 | 53930847 | rs1288634 | G | A | 2 | 0.4732 | 0.4732 | 0.9894 | 0.9894 | 0.6225 | 0 |
| 1 | 54209984 | rs809522 | C | T | 2 | 0.4738 | 0.4738 | 0.9841 | 0.9841 | 0.8195 | 0 |
| 1 | 54375348 | rs1710958 | T | C | 2 | 0.4743 | 0.4743 | 0.9839 | 0.9839 | 0.4707 | 0 |
| 1 | 53792651 | rs3820198 | A | C | 2 | 0.4744 | 0.4744 | 1.0102 | 1.0102 | 0.6503 | 0 |
| 1 | 54010112 | rs6693495 | T | C | 2 | 0.4744 | 0.4744 | 0.9886 | 0.9886 | 0.3618 | 0 |
| 1 | 54425204 | rs6660975 | T | C | 2 | 0.4744 | 0.9014 | 1.0133 | 1.0056 | 0.017 | 82.43 |
| 1 | 54531888 | rs1447647 | C | T | 2 | 0.4745 | 0.4745 | 1.0514 | 1.0514 | 0.6704 | 0 |
| 1 | 53722752 | rs2297663 | A | C | 2 | 0.4756 | 0.4756 | 1.0098 | 1.0098 | 0.916 | 0 |
| 1 | 54583847 | rs1710980 | C | T | 2 | 0.4757 | 0.4848 | 0.9595 | 0.9573 | 0.2869 | 11.82 |
| 1 | 54333474 | rs3766469 | A | T | 2 | 0.4768 | 0.4768 | 1.0098 | 1.0098 | 0.3995 | 0 |
| 1 | 53576592 | rs1288371 | T | C | 2 | 0.4778 | 0.4778 | 1.0107 | 1.0107 | 0.9899 | 0 |
| 1 | 53674542 | rs1157883 | G | A | 2 | 0.4778 | 0.4778 | 0.9883 | 0.9883 | 0.746 | 0 |
| 1 | 54373323 | rs1208037 | G | A | 2 | 0.4785 | 0.4785 | 0.9839 | 0.9839 | 0.51 | 0 |
| 1 | 53707953 | rs1778538 | A | G | 2 | 0.4789 | 0.4789 | 1.0098 | 1.0098 | 0.8283 | 0 |
| 1 | 54341947 | rs1078896 | G | A | 2 | 0.479 | 0.479 | 1.0097 | 1.0097 | 0.438 | 0 |
| 1 | 54540731 | rs1748161 | G | C | 2 | 0.479 | 0.479 | 0.9792 | 0.9792 | 0.8342 | 0 |
| 1 | 53672028 | rs1158151 | A | G | 2 | 0.4797 | 0.4797 | 0.9884 | 0.9884 | 0.7808 | 0 |
| 1 | 53610196 | rs1120611 | A | T | 2 | 0.4802 | 0.4802 | 1.0097 | 1.0097 | 0.3641 | 0 |
| 1 | 54519292 | rs4129477 | G | A | 2 | 0.4818 | 0.4818 | 1.0506 | 1.0506 | 0.6895 | 0 |
| 1 | 53611606 | rs1769320 | G | A | 2 | 0.4819 | 0.4819 | 0.9904 | 0.9904 | 0.4414 | 0 |
| 1 | 53947125 | rs5623825 | T | C | 2 | 0.4821 | 0.6592 | 0.9765 | 0.9763 | 0.1065 | 61.63 |
| 1 | 53611902 | rs7553128 | G | A | 2 | 0.4823 | 0.4823 | 1.012 | 1.012 | 0.698 | 0 |
| 1 | 53851038 | rs5706478 | A | G | 2 | 0.4827 | 0.4827 | 0.9892 | 0.9892 | 0.942 | 0 |
| 1 | 54514065 | rs7573779 | G | T | 2 | 0.4829 | 0.4829 | 1.0497 | 1.0497 | 0.6552 | 0 |
| 1 | 53793248 | rs1933533 | T | G | 2 | 0.4833 | 0.4833 | 1.0101 | 1.0101 | 0.6237 | 0 |
| 1 | 53611699 | rs1679933 | A | G | 2 | 0.4835 | 0.4835 | 0.9904 | 0.9904 | 0.4387 | 0 |
| 1 | 54393004 | rs1206301 | A | G | 2 | 0.4838 | 0.4838 | 0.9845 | 0.9845 | 0.5072 | 0 |
| 1 | 53609535 | rs1120611 | G | A | 2 | 0.4848 | 0.4848 | 1.012 | 1.012 | 0.7435 | 0 |
| 1 | 54672492 | rs1134526 | C | T | 2 | 0.4856 | 0.4856 | 1.0356 | 1.0356 | 0.9926 | 0 |
| 1 | 54356341 | rs6682112 | C | G | 2 | 0.486 | 0.486 | 0.9886 | 0.9886 | 0.9745 | 0 |
| 1 | 54356347 | rs6694562 | A | T | 2 | 0.486 | 0.486 | 0.9886 | 0.9886 | 0.9745 | 0 |
| 1 | 54266678 | rs1181177 | C | G | 2 | 0.4869 | 0.5613 | 0.9581 | 0.9553 | 0.2023 | 38.5 |
| 1 | 54200662 | rs4927033 | C | T | 2 | 0.4875 | 0.4875 | 1.0154 | 1.0154 | 0.9656 | 0 |
| 1 | 54534800 | rs1454834 | T | C | 2 | 0.4879 | 0.4879 | 1.0498 | 1.0498 | 0.6833 | 0 |
| 1 | 54470321 | rs1391278 | C | T | 2 | 0.4882 | 0.4882 | 1.0323 | 1.0323 | 0.8601 | 0 |

| | | | | | | | | | | | |
|---|---|---|---|---|---|---|---|---|---|---|---|
| 1 | 53998448 | rs3855968 | A | T | 2 | 0.4892 | 0.6324 | 0.9746 | 0.9691 | 0.0801 | 67.36 |
| 1 | 54307295 | rs7538888 | T | C | 2 | 0.4894 | 0.4894 | 0.9802 | 0.9802 | 0.6177 | 0 |
| 1 | 54334021 | rs7526676 | T | C | 2 | 0.4894 | 0.4894 | 1.0096 | 1.0096 | 0.4637 | 0 |
| 1 | 53813973 | rs1088878 | A | G | 2 | 0.4895 | 0.4895 | 0.9904 | 0.9904 | 0.8873 | 0 |
| 1 | 54347427 | rs7516530 | G | T | 2 | 0.4898 | 0.4898 | 1.0095 | 1.0095 | 0.4465 | 0 |
| 1 | 53575575 | rs1288369 | C | T | 2 | 0.4909 | 0.4909 | 1.0113 | 1.0113 | 0.566 | 0 |
| 1 | 54373528 | rs1208048 | G | A | 2 | 0.4911 | 0.4911 | 0.9844 | 0.9844 | 0.4778 | 0 |
| 1 | 54608352 | rs1571541 | T | C | 2 | 0.4919 | 0.4919 | 0.9877 | 0.9877 | 0.7163 | 0 |
| 1 | 53492657 | rs6588455 | A | C | 2 | 0.4921 | 0.618 | 1.0164 | 1.0215 | 0.0748 | 68.5 |
| 1 | 54626640 | rs1240846 | G | A | 2 | 0.4921 | 0.5063 | 0.9784 | 0.9782 | 0.297 | 8.05 |
| 1 | 53695882 | rs1337574 | T | C | 2 | 0.4926 | 0.4926 | 0.9889 | 0.9889 | 0.8003 | 0 |
| 1 | 53796099 | rs2788039 | C | G | 2 | 0.4929 | 0.4929 | 1.0101 | 1.0101 | 0.7565 | 0 |
| 1 | 53531298 | rs7290516 | C | T | 2 | 0.493 | 0.493 | 0.962 | 0.962 | 0.9498 | 0 |
| 1 | 53738183 | rs7546246 | A | G | 2 | 0.4931 | 0.4931 | 1.0093 | 1.0093 | 0.7898 | 0 |
| 1 | 53611540 | rs7531069 | C | T | 2 | 0.4954 | 0.4954 | 0.9907 | 0.9907 | 0.4207 | 0 |
| 1 | 53714139 | rs1203115 | C | T | 2 | 0.4956 | 0.4956 | 1.0094 | 1.0094 | 0.8817 | 0 |
| 1 | 53906343 | rs1464708 | G | A | 2 | 0.4956 | 0.4956 | 1.0302 | 1.0302 | 0.3565 | 0 |
| 1 | 54377977 | rs1120624 | T | C | 2 | 0.4957 | 0.4957 | 0.9847 | 0.9847 | 0.6287 | 0 |
| 1 | 53738975 | rs1078895 | C | A | 2 | 0.496 | 0.496 | 1.0093 | 1.0093 | 0.7468 | 0 |
| 1 | 53994981 | rs3855966 | C | T | 2 | 0.496 | 0.6465 | 0.975 | 0.9688 | 0.0651 | 70.62 |
| 1 | 53835096 | rs4926984 | A | G | 2 | 0.4963 | 0.4963 | 0.9904 | 0.9904 | 0.6757 | 0 |
| 1 | 54668562 | rs1711008 | G | A | 2 | 0.4966 | 0.4966 | 0.9417 | 0.9417 | 0.6201 | 0 |
| 1 | 53602788 | rs1274102 | A | G | 2 | 0.497 | 0.497 | 1.0097 | 1.0097 | 0.5289 | 0 |
| 1 | 53626014 | rs6682324 | G | A | 2 | 0.4971 | 0.4971 | 0.9483 | 0.9483 | 0.6566 | 0 |
| 1 | 53601701 | rs1120610 | A | G | 2 | 0.4974 | 0.4974 | 1.0093 | 1.0093 | 0.6829 | 0 |
| 1 | 54380172 | rs1014722 | C | T | 2 | 0.498 | 0.498 | 0.9847 | 0.9847 | 0.5398 | 0 |
| 1 | 54290398 | rs1181187 | T | C | 2 | 0.4984 | 0.4984 | 1.0115 | 1.0115 | 0.9877 | 0 |
| 1 | 54341369 | rs6678296 | A | C | 2 | 0.4984 | 0.4984 | 1.0093 | 1.0093 | 0.3825 | 0 |
| 1 | 53739377 | rs1120613 | A | G | 2 | 0.4985 | 0.4985 | 1.0092 | 1.0092 | 0.7948 | 0 |
| 1 | 54021166 | rs525626 | A | G | 2 | 0.4989 | 0.587 | 0.9907 | 0.9859 | 0.0688 | 69.81 |
| 1 | 53813961 | rs1088878 | T | C | 2 | 0.4993 | 0.4993 | 0.9906 | 0.9906 | 0.902 | 0 |
| 1 | 53836465 | rs1442288 | G | T | 2 | 0.4994 | 0.4994 | 1.0286 | 1.0286 | 0.5084 | 0 |
| 1 | 53631049 | rs1288331 | T | C | 2 | 0.5006 | 0.5006 | 1.0092 | 1.0092 | 0.5501 | 0 |
| 1 | 53791998 | rs6691468 | C | G | 2 | 0.5006 | 0.5006 | 1.0162 | 1.0162 | 0.4535 | 0 |
| 1 | 53851500 | rs4926990 | T | C | 2 | 0.5011 | 0.5011 | 1.0092 | 1.0092 | 0.8087 | 0 |
| 1 | 53625449 | rs6588466 | G | T | 2 | 0.5015 | 0.5015 | 0.9487 | 0.9487 | 0.654 | 0 |
| 1 | 53624516 | rs6588465 | T | C | 2 | 0.5016 | 0.5016 | 0.9489 | 0.9489 | 0.6506 | 0 |
| 1 | 53967080 | rs1710861 | G | A | 2 | 0.5017 | 0.5017 | 1.0141 | 1.0141 | 0.5728 | 0 |
| 1 | 54378026 | rs1120624 | T | C | 2 | 0.5017 | 0.5017 | 0.9849 | 0.9849 | 0.6418 | 0 |
| 1 | 53776407 | rs1288482 | C | T | 2 | 0.5018 | 0.5018 | 1.0114 | 1.0114 | 0.7661 | 0 |
| 1 | 53853533 | rs6588478 | C | T | 2 | 0.502 | 0.502 | 1.0092 | 1.0092 | 0.7014 | 0 |
| 1 | 53857444 | rs6660818 | T | A | 2 | 0.5022 | 0.5022 | 0.9894 | 0.9894 | 0.9429 | 0 |
| 1 | 54343718 | rs1078896 | G | T | 2 | 0.5025 | 0.5025 | 1.0093 | 1.0093 | 0.4269 | 0 |
| 1 | 54513132 | rs4129614 | T | C | 2 | 0.5025 | 0.5614 | 0.975 | 0.9764 | 0.279 | 14.67 |
| 1 | 53588494 | rs1679977 | G | A | 2 | 0.5027 | 0.5027 | 0.9903 | 0.9903 | 0.3433 | 0 |
| 1 | 53736249 | rs3446825 | A | G | 2 | 0.5027 | 0.5027 | 1.0093 | 1.0093 | 0.9847 | 0 |
| 1 | 53486573 | rs7517773 | C | T | 2 | 0.503 | 0.6264 | 1.0162 | 1.0192 | 0.1048 | 61.99 |
| 1 | 53739185 | rs1120613 | G | A | 2 | 0.5033 | 0.5033 | 1.0091 | 1.0091 | 0.8209 | 0 |

| | | | | | | | | | | | |
|---|---|---|---|---|---|---|---|---|---|---|---|
| 1 | 54380900 | rs1208322 | G | A | 2 | 0.5034 | 0.5034 | 0.9849 | 0.9849 | 0.541 | 0 |
| 1 | 54529526 | rs950397 | G | A | 2 | 0.5034 | 0.5034 | 0.9742 | 0.9742 | 0.4905 | 0 |
| 1 | 53552265 | rs1710789 | A | G | 2 | 0.5038 | 0.6786 | 1.03 | 1.0274 | 0.1416 | 53.71 |
| 1 | 54244626 | rs1780412 | A | G | 2 | 0.505 | 0.505 | 1.0135 | 1.0135 | 0.7866 | 0 |
| 1 | 53765814 | rs2782498 | C | G | 2 | 0.5052 | 0.5052 | 1.0113 | 1.0113 | 0.7723 | 0 |
| 1 | 54387010 | rs1180335 | T | C | 2 | 0.5074 | 0.5074 | 0.9717 | 0.9717 | 0.46 | 0 |
| 1 | 54538418 | rs7988443 | T | G | 2 | 0.5075 | 0.5075 | 0.9745 | 0.9745 | 0.535 | 0 |
| 1 | 54241811 | rs5842148 | G | A | 2 | 0.5084 | 0.5997 | 0.9309 | 0.9342 | 0.2321 | 29.96 |
| 1 | 53658460 | rs7688067 | G | A | 2 | 0.509 | 0.509 | 0.9458 | 0.9458 | 0.8272 | 0 |
| 1 | 54658541 | rs1135304 | C | T | 2 | 0.5093 | 0.5093 | 0.9608 | 0.9608 | 0.3356 | 0 |
| 1 | 53657416 | rs7993677 | G | A | 2 | 0.5094 | 0.7937 | 0.9423 | 0.9557 | 0.0561 | 72.6 |
| 1 | 54532130 | rs7688274 | T | C | 2 | 0.5094 | 0.5094 | 1.043 | 1.043 | 0.8687 | 0 |
| 1 | 53736136 | rs1202846 | G | A | 2 | 0.5095 | 0.5095 | 1.0091 | 1.0091 | 0.978 | 0 |
| 1 | 54529997 | rs1180162 | G | A | 2 | 0.51 | 0.51 | 0.9746 | 0.9746 | 0.4848 | 0 |
| 1 | 54335640 | rs6687200 | G | A | 2 | 0.5103 | 0.5103 | 1.009 | 1.009 | 0.5413 | 0 |
| 1 | 53808068 | rs2782494 | G | A | 2 | 0.5109 | 0.5109 | 1.0094 | 1.0094 | 0.6261 | 0 |
| 1 | 53630960 | rs1288332 | G | A | 2 | 0.5116 | 0.5116 | 1.009 | 1.009 | 0.4786 | 0 |
| 1 | 53705174 | rs1778537 | A | T | 2 | 0.5121 | 0.5121 | 1.0098 | 1.0098 | 0.4613 | 0 |
| 1 | 53657106 | rs1137558 | G | A | 2 | 0.5125 | 0.5125 | 0.9591 | 0.9591 | 0.5585 | 0 |
| 1 | 53807897 | rs5862722 | G | A | 2 | 0.5126 | 0.5042 | 1.0434 | 1.0545 | 0.2567 | 22.28 |
| 1 | 54352197 | rs1208575 | C | T | 2 | 0.5129 | 0.5129 | 1.009 | 1.009 | 0.3961 | 0 |
| 1 | 54276021 | rs7408441 | G | A | 2 | 0.5135 | 0.7494 | 1.0513 | 1.0538 | 0.0323 | 78.17 |
| 1 | 54376730 | rs1209508 | A | G | 2 | 0.5146 | 0.5146 | 0.9854 | 0.9854 | 0.6276 | 0 |
| 1 | 54340672 | rs1088881 | G | A | 2 | 0.5147 | 0.5147 | 1.009 | 1.009 | 0.4054 | 0 |
| 1 | 54435755 | rs1203947 | G | A | 2 | 0.5148 | 0.8036 | 1.0123 | 1.0079 | 0.097 | 63.69 |
| 1 | 54278286 | rs1181154 | T | A | 2 | 0.5149 | 0.5149 | 1.011 | 1.011 | 0.8977 | 0 |
| 1 | 53635244 | rs6684231 | A | C | 2 | 0.515 | 0.515 | 1.0101 | 1.0101 | 0.8913 | 0 |
| 1 | 53630914 | rs1288333 | G | C | 2 | 0.5154 | 0.5154 | 1.009 | 1.009 | 0.5062 | 0 |
| 1 | 53763851 | rs2297824 | G | A | 2 | 0.5156 | 0.5156 | 0.9831 | 0.9831 | 0.5008 | 0 |
| 1 | 54669970 | rs3491215 | G | A | 2 | 0.5158 | 0.5158 | 0.9439 | 0.9439 | 0.5907 | 0 |
| 1 | 53527132 | rs6118138 | C | T | 2 | 0.5161 | 0.5332 | 1.0152 | 1.0157 | 0.2836 | 13.02 |
| 1 | 53628085 | rs1114940 | A | G | 2 | 0.5165 | 0.5165 | 0.9503 | 0.9503 | 0.6858 | 0 |
| 1 | 53856091 | rs7852581 | C | A | 2 | 0.5165 | 0.5165 | 0.9898 | 0.9898 | 0.9735 | 0 |
| 1 | 53814038 | rs7535922 | C | T | 2 | 0.517 | 0.517 | 0.9912 | 0.9912 | 0.6459 | 0 |
| 1 | 54359030 | rs7731351 | C | T | 2 | 0.5172 | 0.5375 | 0.9849 | 0.9821 | 0.2243 | 32.29 |
| 1 | 54690954 | rs3088379 | T | C | 2 | 0.5172 | 0.5172 | 1.0089 | 1.0089 | 0.5961 | 0 |
| 1 | 54034869 | rs1949943 | T | C | 2 | 0.5177 | 0.6546 | 1.0098 | 1.0086 | 0.2082 | 36.86 |
| 1 | 53624431 | rs6588464 | G | A | 2 | 0.5179 | 0.5179 | 0.9507 | 0.9507 | 0.6681 | 0 |
| 1 | 53689077 | rs8017882 | C | T | 2 | 0.5181 | 0.5346 | 0.9744 | 0.966 | 0.1846 | 43.19 |
| 1 | 54098459 | rs6775211 | G | A | 2 | 0.5183 | 0.5183 | 0.9901 | 0.9901 | 0.5341 | 0 |
| 1 | 53525481 | rs1737627 | C | T | 2 | 0.5188 | 0.5188 | 0.9773 | 0.9773 | 0.4976 | 0 |
| 1 | 53855711 | rs1539500 | G | A | 2 | 0.5191 | 0.5191 | 0.9898 | 0.9898 | 0.9685 | 0 |
| 1 | 53626511 | rs7778496 | T | G | 2 | 0.5193 | 0.5193 | 0.9506 | 0.9506 | 0.6683 | 0 |
| 1 | 53735026 | rs1088877 | C | T | 2 | 0.5193 | 0.5193 | 1.0088 | 1.0088 | 0.9846 | 0 |
| 1 | 54670159 | rs1711009 | C | T | 2 | 0.5193 | 0.5193 | 0.9447 | 0.9447 | 0.5857 | 0 |
| 1 | 54309582 | rs1181015 | A | G | 2 | 0.5195 | 0.5195 | 0.9816 | 0.9816 | 0.6828 | 0 |
| 1 | 53486561 | rs6695341 | G | A | 2 | 0.5198 | 0.6581 | 1.0158 | 1.0182 | 0.0965 | 63.81 |
| 1 | 53716416 | rs3737983 | G | A | 2 | 0.5198 | 0.5198 | 1.0088 | 1.0088 | 0.9269 | 0 |

| | | | | | | | | | | | |
|---|---|---|---|---|---|---|---|---|---|---|---|
| 1 | 54688529 | rs5944088 | A | G | 2 | 0.5203 | 0.8287 | 1.0124 | 1.0066 | 0.1299 | 56.4 |
| 1 | 53880799 | rs1157890 | G | A | 2 | 0.5205 | 0.5205 | 0.9885 | 0.9885 | 0.5684 | 0 |
| 1 | 53513022 | rs8006970 | G | A | 2 | 0.5216 | 0.5216 | 0.9775 | 0.9775 | 0.4991 | 0 |
| 1 | 54328988 | rs6588493 | C | A | 2 | 0.5216 | 0.5216 | 1.0089 | 1.0089 | 0.5142 | 0 |
| 1 | 54536086 | rs1710971 | C | T | 2 | 0.5218 | 0.5218 | 0.9753 | 0.9753 | 0.5057 | 0 |
| 1 | 54602977 | rs1207884 | A | C | 2 | 0.5218 | 0.5218 | 1.0105 | 1.0105 | 0.8646 | 0 |
| 1 | 53681699 | rs1134688 | T | G | 2 | 0.522 | 0.522 | 0.9894 | 0.9894 | 0.8928 | 0 |
| 1 | 54681768 | rs2236555 | A | G | 2 | 0.5222 | 0.5222 | 0.9453 | 0.9453 | 0.5643 | 0 |
| 1 | 54669984 | rs6693202 | A | G | 2 | 0.5223 | 0.5223 | 1.0116 | 1.0116 | 0.5796 | 0 |
| 1 | 53720211 | rs7732410 | T | C | 2 | 0.5225 | 0.5225 | 0.9461 | 0.9461 | 0.495 | 0 |
| 1 | 53497596 | rs1207598 | G | T | 2 | 0.5237 | 0.5567 | 1.018 | 1.0175 | 0.2946 | 8.97 |
| 1 | 54162232 | rs1181059 | C | T | 2 | 0.5237 | 0.5237 | 1.0205 | 1.0205 | 0.9483 | 0 |
| 1 | 53624161 | rs6588463 | C | G | 2 | 0.5238 | 0.5238 | 0.9514 | 0.9514 | 0.6725 | 0 |
| 1 | 54588056 | rs1088883 | G | C | 2 | 0.524 | 0.524 | 0.9916 | 0.9916 | 0.6247 | 0 |
| 1 | 54279124 | rs1436906 | T | C | 2 | 0.5242 | 0.5242 | 1.0458 | 1.0458 | 0.7086 | 0 |
| 1 | 54214077 | rs1181163 | G | A | 2 | 0.5244 | 0.5244 | 0.9573 | 0.9573 | 0.7107 | 0 |
| 1 | 54523689 | rs5757687 | T | C | 2 | 0.5244 | 0.5244 | 0.9756 | 0.9756 | 0.4327 | 0 |
| 1 | 54672112 | rs1450792 | T | C | 2 | 0.5249 | 0.5249 | 0.9456 | 0.9456 | 0.5509 | 0 |
| 1 | 54684622 | rs3556587 | A | G | 2 | 0.5253 | 0.5667 | 1.0115 | 1.0191 | 0.0831 | 66.7 |
| 1 | 54281836 | rs1181152 | A | G | 2 | 0.5254 | 0.5254 | 1.0107 | 1.0107 | 0.9535 | 0 |
| 1 | 54681920 | rs13571 | G | C | 2 | 0.5254 | 0.5336 | 0.9901 | 0.9897 | 0.29 | 10.69 |
| 1 | 54527829 | rs1710969 | G | A | 2 | 0.5268 | 0.5268 | 0.9757 | 0.9757 | 0.4695 | 0 |
| 1 | 53625313 | rs7267089 | T | C | 2 | 0.5273 | 0.5273 | 0.9727 | 0.9727 | 0.658 | 0 |
| 1 | 54249029 | rs6657928 | T | C | 2 | 0.5275 | 0.5275 | 1.0107 | 1.0107 | 0.9441 | 0 |
| 1 | 54008488 | rs3889185 | G | C | 2 | 0.5276 | 0.5276 | 0.9899 | 0.9899 | 0.3676 | 0 |
| 1 | 54672468 | rs1141781 | A | T | 2 | 0.529 | 0.529 | 0.946 | 0.946 | 0.5486 | 0 |
| 1 | 53735514 | rs7349164 | G | C | 2 | 0.5291 | 0.5291 | 1.0087 | 1.0087 | 0.9625 | 0 |
| 1 | 53997093 | rs6673003 | T | C | 2 | 0.5295 | 0.6531 | 0.9769 | 0.9717 | 0.088 | 65.64 |
| 1 | 53532117 | rs1015891 | A | G | 2 | 0.5296 | 0.5296 | 1.0148 | 1.0148 | 0.3211 | 0 |
| 1 | 53567492 | rs7528784 | G | A | 2 | 0.5305 | 0.5305 | 0.9729 | 0.9729 | 0.4898 | 0 |
| 1 | 54328814 | rs6681549 | G | A | 2 | 0.5307 | 0.5307 | 1.0086 | 1.0086 | 0.4901 | 0 |
| 1 | 54579655 | rs1329779 | A | G | 2 | 0.5307 | 0.5307 | 0.9755 | 0.9755 | 0.4355 | 0 |
| 1 | 53611537 | rs7552743 | G | A | 2 | 0.5324 | 0.5324 | 0.9914 | 0.9914 | 0.3804 | 0 |
| 1 | 54494702 | rs1120627 | G | T | 2 | 0.5324 | 0.5324 | 1.0173 | 1.0173 | 0.8405 | 0 |
| 1 | 54344690 | rs2316992 | C | T | 2 | 0.5325 | 0.5325 | 1.0086 | 1.0086 | 0.3978 | 0 |
| 1 | 54029776 | rs1738581 | G | A | 2 | 0.5327 | 0.5327 | 0.9804 | 0.9804 | 0.5698 | 0 |
| 1 | 54186792 | rs1849608 | T | C | 2 | 0.5329 | 0.9376 | 0.9535 | 0.9892 | 0.0947 | 64.19 |
| 1 | 53596328 | rs1288360 | A | G | 2 | 0.5333 | 0.5541 | 0.9829 | 0.9776 | 0.1825 | 43.73 |
| 1 | 54564485 | rs7547352 | G | A | 2 | 0.5336 | 0.5336 | 1.0555 | 1.0555 | 0.8845 | 0 |
| 1 | 53778252 | rs1181039 | A | C | 2 | 0.5343 | 0.5754 | 0.9611 | 0.9608 | 0.2628 | 20.24 |
| 1 | 54518725 | rs4129477 | G | A | 2 | 0.5354 | 0.5354 | 1.044 | 1.044 | 0.6595 | 0 |
| 1 | 53610122 | rs1120611 | G | A | 2 | 0.5355 | 0.5355 | 1.0085 | 1.0085 | 0.3579 | 0 |
| 1 | 54522645 | rs1118935 | C | G | 2 | 0.5359 | 0.5359 | 0.9763 | 0.9763 | 0.4195 | 0 |
| 1 | 53526136 | rs1452501 | C | G | 2 | 0.536 | 0.7454 | 1.04 | 1.0272 | 0.2188 | 33.86 |
| 1 | 54066774 | rs1147142 | A | C | 2 | 0.5361 | 0.5361 | 0.9735 | 0.9735 | 0.5649 | 0 |
| 1 | 54334315 | rs7526784 | A | T | 2 | 0.5364 | 0.5364 | 1.0085 | 1.0085 | 0.4699 | 0 |
| 1 | 53787298 | rs872316 | G | A | 2 | 0.5367 | 0.5367 | 1.0434 | 1.0434 | 0.7776 | 0 |
| 1 | 53572065 | rs1288425 | G | A | 2 | 0.537 | 0.537 | 0.9874 | 0.9874 | 0.8021 | 0 |

| | | | | | | | | | | | |
|---|---|---|---|---|---|---|---|---|---|---|---|
| 1 | 53728048 | rs1156075 | A | G | 2 | 0.5376 | 0.5376 | 0.951 | 0.951 | 0.5599 | 0 |
| 1 | 53787633 | rs1288520 | G | A | 2 | 0.5377 | 0.5377 | 1.009 | 1.009 | 0.769 | 0 |
| 1 | 53958280 | rs5632733 | C | A | 2 | 0.5381 | 0.7094 | 0.9788 | 0.9798 | 0.1148 | 59.79 |
| 1 | 53787175 | rs5880280 | T | C | 2 | 0.5382 | 0.5382 | 1.0431 | 1.0431 | 0.8762 | 0 |
| 1 | 53786710 | rs871452 | T | C | 2 | 0.5385 | 0.5385 | 1.0431 | 1.0431 | 0.8749 | 0 |
| 1 | 53623735 | rs1203902 | T | C | 2 | 0.5388 | 0.5388 | 1.0102 | 1.0102 | 0.6634 | 0 |
| 1 | 53792936 | rs1144589 | A | G | 2 | 0.539 | 0.539 | 1.0464 | 1.0464 | 0.7224 | 0 |
| 1 | 54330717 | rs6588494 | T | A | 2 | 0.5395 | 0.5395 | 1.0085 | 1.0085 | 0.4474 | 0 |
| 1 | 53786872 | rs871453 | A | C | 2 | 0.5396 | 0.5396 | 1.043 | 1.043 | 0.8745 | 0 |
| 1 | 53810549 | rs7547624 | G | A | 2 | 0.5399 | 0.5399 | 1.0181 | 1.0181 | 0.423 | 0 |
| 1 | 54333808 | rs1142737 | G | C | 2 | 0.5399 | 0.5399 | 1.035 | 1.035 | 0.5539 | 0 |
| 1 | 53611401 | rs1769318 | C | G | 2 | 0.5402 | 0.5402 | 0.9916 | 0.9916 | 0.3583 | 0 |
| 1 | 53481477 | rs1710756 | G | A | 2 | 0.5405 | 0.6447 | 1.0144 | 1.0183 | 0.0962 | 63.86 |
| 1 | 53610094 | rs1120611 | A | G | 2 | 0.5406 | 0.5406 | 1.0084 | 1.0084 | 0.4632 | 0 |
| 1 | 54584403 | rs7713220 | C | T | 2 | 0.5406 | 0.5406 | 1.044 | 1.044 | 0.7888 | 0 |
| 1 | 53767474 | rs7529821 | A | C | 2 | 0.5417 | 0.5417 | 1.0103 | 1.0103 | 0.5803 | 0 |
| 1 | 53729210 | rs6668172 | C | T | 2 | 0.5419 | 0.5419 | 1.0083 | 1.0083 | 0.9083 | 0 |
| 1 | 53717656 | rs1240890 | G | A | 2 | 0.5425 | 0.5425 | 1.0084 | 1.0084 | 0.909 | 0 |
| 1 | 53767650 | rs7518153 | C | T | 2 | 0.5426 | 0.5426 | 1.0102 | 1.0102 | 0.5661 | 0 |
| 1 | 53486262 | rs6684285 | T | C | 2 | 0.5436 | 0.6423 | 1.0144 | 1.0188 | 0.0934 | 64.48 |
| 1 | 53487047 | rs7540852 | T | C | 2 | 0.5436 | 0.6423 | 1.0144 | 1.0188 | 0.0934 | 64.48 |
| 1 | 53766972 | rs867884 | A | G | 2 | 0.5439 | 0.5439 | 1.0102 | 1.0102 | 0.5975 | 0 |
| 1 | 54019668 | rs490868 | A | G | 2 | 0.5445 | 0.5445 | 0.939 | 0.939 | 0.5504 | 0 |
| 1 | 53486512 | rs6686963 | T | C | 2 | 0.5446 | 0.6453 | 1.0143 | 1.0184 | 0.0959 | 63.92 |
| 1 | 53739367 | rs1181197 | G | A | 2 | 0.5447 | 0.5447 | 1.0083 | 1.0083 | 0.7151 | 0 |
| 1 | 53723190 | rs869987 | G | T | 2 | 0.545 | 0.545 | 1.0082 | 1.0082 | 0.8201 | 0 |
| 1 | 53730607 | rs1073638 | C | T | 2 | 0.545 | 0.545 | 1.0082 | 1.0082 | 0.9023 | 0 |
| 1 | 53916077 | rs5588143 | G | A | 2 | 0.5453 | 0.5453 | 1.0114 | 1.0114 | 0.3963 | 0 |
| 1 | 54025213 | rs3900873 | C | T | 2 | 0.5454 | 0.6284 | 0.9918 | 0.9868 | 0.0514 | 73.64 |
| 1 | 53479298 | rs5814049 | G | A | 2 | 0.5455 | 0.6459 | 1.0144 | 1.018 | 0.1033 | 62.32 |
| 1 | 53784785 | rs5879853 | T | C | 2 | 0.5462 | 0.5462 | 1.0428 | 1.0428 | 0.9082 | 0 |
| 1 | 54527461 | rs1710968 | G | A | 2 | 0.5462 | 0.5462 | 0.9764 | 0.9764 | 0.4895 | 0 |
| 1 | 53493072 | rs7290313 | G | C | 2 | 0.5467 | 0.6469 | 1.0142 | 1.0182 | 0.0975 | 63.59 |
| 1 | 53808589 | rs7456467 | G | A | 2 | 0.5468 | 0.5454 | 1.0397 | 1.0407 | 0.3115 | 2.38 |
| 1 | 53919491 | rs5589971 | C | T | 2 | 0.5472 | 0.6199 | 0.9903 | 0.9872 | 0.1144 | 59.88 |
| 1 | 53776827 | rs1180235 | C | T | 2 | 0.5473 | 0.5473 | 0.9838 | 0.9838 | 0.5507 | 0 |
| 1 | 53492867 | rs6674545 | C | T | 2 | 0.5477 | 0.6461 | 1.0142 | 1.0183 | 0.0964 | 63.83 |
| 1 | 53679229 | rs1799822 | A | G | 2 | 0.5477 | 0.5477 | 0.9901 | 0.9901 | 0.8659 | 0 |
| 1 | 53721864 | rs1074968 | G | C | 2 | 0.5477 | 0.5477 | 1.0082 | 1.0082 | 0.9917 | 0 |
| 1 | 53789711 | rs7289540 | A | C | 2 | 0.5478 | 0.5478 | 1.0415 | 1.0415 | 0.9914 | 0 |
| 1 | 54504112 | rs1157266 | G | A | 2 | 0.5479 | 0.5479 | 1.0289 | 1.0289 | 0.7121 | 0 |
| 1 | 53723349 | rs869988 | T | C | 2 | 0.548 | 0.548 | 1.0082 | 1.0082 | 0.8864 | 0 |
| 1 | 53931983 | rs5825960 | G | T | 2 | 0.5481 | 0.5481 | 1.0112 | 1.0112 | 0.5289 | 0 |
| 1 | 54217535 | rs7266069 | G | A | 2 | 0.5485 | 0.5485 | 0.9849 | 0.9849 | 0.398 | 0 |
| 1 | 53488916 | rs4378148 | A | G | 2 | 0.5491 | 0.6393 | 1.0143 | 1.0189 | 0.0969 | 63.71 |
| 1 | 53488917 | rs4130006 | C | T | 2 | 0.5491 | 0.6395 | 1.0143 | 1.0188 | 0.0984 | 63.39 |
| 1 | 54612220 | rs6704344 | T | C | 2 | 0.5492 | 0.5492 | 1.0251 | 1.0251 | 0.3927 | 0 |
| 1 | 53922844 | rs1204743 | G | T | 2 | 0.5498 | 0.5798 | 0.9906 | 0.9893 | 0.2246 | 32.19 |

| | | | | | | | | | | | |
|---|---|---|---|---|---|---|---|---|---|---|---|
| 1 | 53776413 | rs1288481 | T | C | 2 | 0.5505 | 0.5505 | 1.01 | 1.01 | 0.7614 | 0 |
| 1 | 54523344 | rs7408514 | C | A | 2 | 0.5508 | 0.5508 | 0.9771 | 0.9771 | 0.4246 | 0 |
| 1 | 53950226 | rs6082008 | G | A | 2 | 0.5513 | 0.7064 | 0.9795 | 0.9803 | 0.127 | 57.06 |
| 1 | 53630003 | rs1288337 | A | G | 2 | 0.5514 | 0.5514 | 1.0083 | 1.0083 | 0.4281 | 0 |
| 1 | 54565650 | rs1209574 | A | G | 2 | 0.5518 | 0.5518 | 0.9777 | 0.9777 | 0.3893 | 0 |
| 1 | 54532638 | rs1240392 | G | A | 2 | 0.552 | 0.552 | 1.0526 | 1.0526 | 0.7581 | 0 |
| 1 | 53488402 | rs6588450 | G | C | 2 | 0.5522 | 0.6483 | 1.0141 | 1.0181 | 0.0982 | 63.42 |
| 1 | 54509616 | rs1207054 | C | T | 2 | 0.5523 | 0.5523 | 1.0418 | 1.0418 | 0.5853 | 0 |
| 1 | 53623662 | rs1204929 | C | T | 2 | 0.5524 | 0.5524 | 1.0098 | 1.0098 | 0.5483 | 0 |
| 1 | 54511200 | rs4153804 | A | G | 2 | 0.5524 | 0.5524 | 0.9771 | 0.9771 | 0.7432 | 0 |
| 1 | 53486378 | rs6686866 | T | C | 2 | 0.5525 | 0.6492 | 1.014 | 1.0179 | 0.1002 | 63 |
| 1 | 53486613 | rs7290139 | G | A | 2 | 0.5526 | 0.6499 | 1.014 | 1.0179 | 0.0997 | 63.11 |
| 1 | 53494599 | rs6049254 | C | A | 2 | 0.553 | 0.6467 | 1.0141 | 1.0183 | 0.0968 | 63.73 |
| 1 | 54042650 | rs9651200 | T | C | 2 | 0.553 | 0.6101 | 0.9479 | 0.9505 | 0.2723 | 17.01 |
| 1 | 53488092 | rs6690936 | T | G | 2 | 0.5534 | 0.6498 | 1.014 | 1.0177 | 0.1017 | 62.67 |
| 1 | 53641937 | rs6176958 | C | T | 2 | 0.5536 | 0.5715 | 1.0557 | 1.0543 | 0.3117 | 2.29 |
| 1 | 54681600 | rs3471302 | C | G | 2 | 0.5538 | 0.5538 | 0.9491 | 0.9491 | 0.5893 | 0 |
| 1 | 54041190 | rs7267519 | G | C | 2 | 0.5541 | 0.8286 | 0.9902 | 0.9943 | 0.1216 | 58.27 |
| 1 | 53479940 | rs6678994 | A | G | 2 | 0.5543 | 0.6509 | 1.0139 | 1.0177 | 0.1012 | 62.79 |
| 1 | 53485391 | rs7544876 | G | A | 2 | 0.555 | 0.6512 | 1.0139 | 1.0176 | 0.1016 | 62.69 |
| 1 | 53493540 | rs1710763 | T | C | 2 | 0.5554 | 0.651 | 1.0139 | 1.0178 | 0.1007 | 62.9 |
| 1 | 53993862 | rs7289334 | C | T | 2 | 0.5554 | 0.6826 | 0.9784 | 0.9722 | 0.0647 | 70.7 |
| 1 | 53675049 | rs370493 | G | A | 2 | 0.5557 | 0.5557 | 1.0087 | 1.0087 | 0.3637 | 0 |
| 1 | 54342837 | rs1120623 | C | T | 2 | 0.5558 | 0.5558 | 1.0081 | 1.0081 | 0.4764 | 0 |
| 1 | 53531201 | rs7880879 | T | C | 2 | 0.5562 | 0.5562 | 1.0139 | 1.0139 | 0.3332 | 0 |
| 1 | 53735446 | rs1088877 | G | T | 2 | 0.5563 | 0.5563 | 1.008 | 1.008 | 0.8991 | 0 |
| 1 | 53489244 | rs7555162 | G | A | 2 | 0.5568 | 0.6509 | 1.0139 | 1.0177 | 0.103 | 62.39 |
| 1 | 53490179 | rs5746093 | T | A | 2 | 0.5569 | 0.6513 | 1.0139 | 1.0177 | 0.1009 | 62.85 |
| 1 | 53480745 | rs7529265 | G | A | 2 | 0.557 | 0.653 | 1.0138 | 1.0176 | 0.1004 | 62.94 |
| 1 | 53489815 | rs6041192 | G | A | 2 | 0.5573 | 0.6511 | 1.0138 | 1.0175 | 0.1038 | 62.21 |
| 1 | 53489949 | rs5865163 | A | C | 2 | 0.5573 | 0.6511 | 1.0138 | 1.0175 | 0.1038 | 62.21 |
| 1 | 53773248 | rs7884883 | G | A | 2 | 0.5573 | 0.5573 | 0.9832 | 0.9832 | 0.8546 | 0 |
| 1 | 53837187 | rs6588477 | A | G | 2 | 0.5575 | 0.5575 | 0.986 | 0.986 | 0.9305 | 0 |
| 1 | 54513867 | rs7408514 | G | T | 2 | 0.5575 | 0.5575 | 0.977 | 0.977 | 0.5133 | 0 |
| 1 | 54690271 | rs6174441 | C | A | 2 | 0.5579 | 0.5579 | 0.9491 | 0.9491 | 0.6154 | 0 |
| 1 | 53812854 | rs7289746 | A | G | 2 | 0.558 | 0.558 | 0.9372 | 0.9372 | 0.5349 | 0 |
| 1 | 54043269 | rs7530524 | G | C | 2 | 0.5581 | 0.6233 | 0.9485 | 0.9516 | 0.2655 | 19.33 |
| 1 | 53486314 | rs6684182 | A | G | 2 | 0.5584 | 0.6526 | 1.0138 | 1.0176 | 0.1018 | 62.66 |
| 1 | 53486885 | rs7526661 | C | T | 2 | 0.5584 | 0.6526 | 1.0138 | 1.0176 | 0.1018 | 62.66 |
| 1 | 53488129 | rs6675803 | C | G | 2 | 0.5584 | 0.6526 | 1.0138 | 1.0176 | 0.1018 | 62.66 |
| 1 | 53488202 | rs6675908 | C | T | 2 | 0.5584 | 0.6526 | 1.0138 | 1.0176 | 0.1018 | 62.66 |
| 1 | 53488291 | rs6678556 | C | T | 2 | 0.5584 | 0.6526 | 1.0138 | 1.0176 | 0.1018 | 62.66 |
| 1 | 54233542 | rs4131115 | A | G | 2 | 0.5584 | 0.5584 | 0.9739 | 0.9739 | 0.7426 | 0 |
| 1 | 53787483 | rs872314 | A | G | 2 | 0.5585 | 0.5585 | 1.0409 | 1.0409 | 0.9089 | 0 |
| 1 | 53489534 | rs7535959 | C | T | 2 | 0.5587 | 0.6523 | 1.0138 | 1.0176 | 0.1029 | 62.4 |
| 1 | 53491334 | rs7290312 | C | T | 2 | 0.5587 | 0.6523 | 1.0138 | 1.0176 | 0.1029 | 62.4 |
| 1 | 53678554 | rs737464 | A | G | 2 | 0.5589 | 0.5589 | 1.0079 | 1.0079 | 0.4208 | 0 |
| 1 | 53614976 | rs1679941 | A | G | 2 | 0.559 | 0.559 | 1.0078 | 1.0078 | 0.8138 | 0 |

| | | | | | | | | | | | |
|---|---|---|---|---|---|---|---|---|---|---|---|
| 1 | 53492434 | rs6588453 | A | T | 2 | 0.5594 | 0.6524 | 1.0137 | 1.0174 | 0.1048 | 61.98 |
| 1 | 53489296 | rs7549603 | T | G | 2 | 0.5596 | 0.6531 | 1.0138 | 1.0176 | 0.1022 | 62.56 |
| 1 | 53765419 | rs2782499 | A | G | 2 | 0.5597 | 0.5597 | 1.0099 | 1.0099 | 0.725 | 0 |
| 1 | 53490763 | rs7290312 | C | T | 2 | 0.5598 | 0.6532 | 1.0138 | 1.0175 | 0.1026 | 62.47 |
| 1 | 53563234 | rs1288416 | A | G | 2 | 0.56 | 0.5947 | 0.9905 | 0.985 | 0.0966 | 63.77 |
| 1 | 53489601 | rs7536048 | C | T | 2 | 0.5601 | 0.6534 | 1.0138 | 1.0175 | 0.1026 | 62.48 |
| 1 | 53489717 | rs1710762 | A | G | 2 | 0.5601 | 0.6534 | 1.0138 | 1.0175 | 0.1026 | 62.48 |
| 1 | 53489759 | rs1710763 | T | A | 2 | 0.5601 | 0.6534 | 1.0138 | 1.0175 | 0.1026 | 62.48 |
| 1 | 53489983 | rs5828379 | G | A | 2 | 0.5601 | 0.6534 | 1.0138 | 1.0175 | 0.1026 | 62.48 |
| 1 | 53490989 | rs7290312 | A | G | 2 | 0.5601 | 0.6534 | 1.0138 | 1.0175 | 0.1026 | 62.48 |
| 1 | 53491013 | rs7290312 | G | C | 2 | 0.5601 | 0.6534 | 1.0138 | 1.0175 | 0.1026 | 62.48 |
| 1 | 53491039 | rs7736877 | A | G | 2 | 0.5601 | 0.6534 | 1.0138 | 1.0175 | 0.1026 | 62.48 |
| 1 | 53491183 | rs7290312 | C | T | 2 | 0.5601 | 0.6534 | 1.0138 | 1.0175 | 0.1026 | 62.48 |
| 1 | 53491292 | rs7290312 | C | A | 2 | 0.5601 | 0.6534 | 1.0138 | 1.0175 | 0.1026 | 62.48 |
| 1 | 54343462 | rs7541170 | T | C | 2 | 0.5601 | 0.5601 | 1.008 | 1.008 | 0.4043 | 0 |
| 1 | 54588636 | rs1209591 | G | A | 2 | 0.5602 | 0.5602 | 1.0228 | 1.0228 | 0.9143 | 0 |
| 1 | 53922155 | rs7543581 | C | T | 2 | 0.5603 | 0.5871 | 0.9908 | 0.9896 | 0.2287 | 30.97 |
| 1 | 54611295 | rs617067 | C | T | 2 | 0.5605 | 0.5605 | 0.9904 | 0.9904 | 0.406 | 0 |
| 1 | 54165440 | rs1159107 | C | G | 2 | 0.561 | 0.561 | 1.0202 | 1.0202 | 0.5803 | 0 |
| 1 | 54357981 | rs6701185 | T | G | 2 | 0.5614 | 0.5614 | 0.9905 | 0.9905 | 0.9193 | 0 |
| 1 | 53500734 | rs1420944 | G | A | 2 | 0.5615 | 0.5615 | 1.0184 | 1.0184 | 0.8947 | 0 |
| 1 | 54124935 | rs1159115 | G | A | 2 | 0.5615 | 0.5965 | 0.9871 | 0.9852 | 0.2136 | 35.35 |
| 1 | 53492366 | rs6588451 | C | T | 2 | 0.5621 | 0.6541 | 1.0137 | 1.0175 | 0.1031 | 62.36 |
| 1 | 53492412 | rs6588452 | C | T | 2 | 0.5621 | 0.6541 | 1.0137 | 1.0175 | 0.1031 | 62.36 |
| 1 | 53789344 | rs7289539 | T | G | 2 | 0.5625 | 0.5625 | 1.04 | 1.04 | 0.9944 | 0 |
| 1 | 54568531 | rs6168793 | C | A | 2 | 0.5625 | 0.5625 | 0.9778 | 0.9778 | 0.6359 | 0 |
| 1 | 53809030 | rs2782497 | T | C | 2 | 0.5626 | 0.5626 | 1.0084 | 1.0084 | 0.5851 | 0 |
| 1 | 54043487 | rs7289929 | G | A | 2 | 0.5628 | 0.6306 | 0.9491 | 0.9523 | 0.263 | 20.17 |
| 1 | 53494879 | rs1710765 | C | G | 2 | 0.5629 | 0.6538 | 1.0136 | 1.0173 | 0.1059 | 61.75 |
| 1 | 53967392 | rs6176845 | G | A | 2 | 0.5632 | 0.5632 | 1.0128 | 1.0128 | 0.7382 | 0 |
| 1 | 53492917 | rs6698146 | G | A | 2 | 0.5635 | 0.655 | 1.0136 | 1.0174 | 0.1038 | 62.2 |
| 1 | 53492965 | rs6698224 | G | A | 2 | 0.5635 | 0.655 | 1.0136 | 1.0174 | 0.1038 | 62.2 |
| 1 | 54611730 | rs1418304 | A | T | 2 | 0.5639 | 0.5639 | 1.0311 | 1.0311 | 0.4378 | 0 |
| 1 | 53538231 | rs1710784 | C | T | 2 | 0.5644 | 0.5644 | 0.9796 | 0.9796 | 0.816 | 0 |
| 1 | 53672270 | rs7539949 | G | A | 2 | 0.5644 | 0.5644 | 1.0078 | 1.0078 | 0.4307 | 0 |
| 1 | 54341646 | rs6678606 | A | C | 2 | 0.5644 | 0.5644 | 1.0079 | 1.0079 | 0.3824 | 0 |
| 1 | 54007863 | rs1240331 | T | C | 2 | 0.5651 | 0.5651 | 0.9908 | 0.9908 | 0.3468 | 0 |
| 1 | 53493426 | rs7996929 | C | G | 2 | 0.5652 | 0.6557 | 1.0136 | 1.0173 | 0.1043 | 62.1 |
| 1 | 54538322 | rs1128630 | A | C | 2 | 0.5653 | 0.5653 | 0.9779 | 0.9779 | 0.4843 | 0 |
| 1 | 53842152 | rs7267512 | A | G | 2 | 0.5654 | 0.5654 | 1.0316 | 1.0316 | 0.696 | 0 |
| 1 | 54547459 | rs6040636 | C | G | 2 | 0.5655 | 0.5655 | 0.9779 | 0.9779 | 0.5513 | 0 |
| 1 | 54343294 | rs7541049 | T | C | 2 | 0.5661 | 0.5661 | 1.0079 | 1.0079 | 0.394 | 0 |
| 1 | 54680058 | rs4927072 | A | G | 2 | 0.5661 | 0.595 | 1.0103 | 1.0182 | 0.0707 | 69.4 |
| 1 | 53493625 | rs1710764 | A | G | 2 | 0.5662 | 0.6559 | 1.0136 | 1.0173 | 0.1045 | 62.06 |
| 1 | 53572574 | rs3516105 | C | G | 2 | 0.5662 | 0.5662 | 1.0115 | 1.0115 | 0.3373 | 0 |
| 1 | 53493822 | rs1710764 | G | T | 2 | 0.5663 | 0.656 | 1.0136 | 1.0173 | 0.1045 | 62.06 |
| 1 | 54505756 | rs4133134 | G | A | 2 | 0.5666 | 0.5666 | 1.0403 | 1.0403 | 0.5716 | 0 |
| 1 | 54320905 | rs5632084 | C | T | 2 | 0.567 | 0.9829 | 0.9916 | 0.9992 | 0.0215 | 81.08 |

| | | | | | | | | | | | |
|---|---|---|---|---|---|---|---|---|---|---|---|
| 1 | 54542802 | rs5896659 | A | G | 2 | 0.5673 | 0.6228 | 0.9891 | 0.9897 | 0.2748 | 16.17 |
| 1 | 53790907 | rs7289740 | T | C | 2 | 0.5675 | 0.5675 | 1.0418 | 1.0418 | 0.7367 | 0 |
| 1 | 53609793 | rs1275062 | G | A | 2 | 0.5676 | 0.5676 | 1.0079 | 1.0079 | 0.3547 | 0 |
| 1 | 54338639 | rs1214295 | G | A | 2 | 0.568 | 0.568 | 1.0078 | 1.0078 | 0.4728 | 0 |
| 1 | 53494345 | rs6092613 | A | C | 2 | 0.5681 | 0.6569 | 1.0135 | 1.0172 | 0.1052 | 61.9 |
| 1 | 53789431 | rs7289539 | G | T | 2 | 0.5684 | 0.5684 | 1.0394 | 1.0394 | 0.9891 | 0 |
| 1 | 53614880 | rs1628297 | T | C | 2 | 0.5685 | 0.5685 | 1.0077 | 1.0077 | 0.8135 | 0 |
| 1 | 53609355 | rs1120611 | T | A | 2 | 0.5687 | 0.5687 | 1.0077 | 1.0077 | 0.5613 | 0 |
| 1 | 54341762 | rs6689665 | G | A | 2 | 0.5687 | 0.5687 | 1.0078 | 1.0078 | 0.377 | 0 |
| 1 | 54571630 | rs4498751 | A | G | 2 | 0.5691 | 0.5691 | 0.9783 | 0.9783 | 0.402 | 0 |
| 1 | 53494781 | rs1710764 | C | T | 2 | 0.5693 | 0.6575 | 1.0134 | 1.0172 | 0.1057 | 61.8 |
| 1 | 54612981 | rs7487442 | T | A | 2 | 0.5697 | 0.5697 | 1.0237 | 1.0237 | 0.4241 | 0 |
| 1 | 54549019 | rs6129138 | C | A | 2 | 0.5698 | 0.5698 | 0.9782 | 0.9782 | 0.5478 | 0 |
| 1 | 53495529 | rs1710766 | T | G | 2 | 0.5699 | 0.6578 | 1.0134 | 1.0171 | 0.1057 | 61.8 |
| 1 | 54043390 | rs1710883 | G | A | 2 | 0.5699 | 0.6446 | 0.95 | 0.9536 | 0.2569 | 22.2 |
| 1 | 53623564 | rs6666580 | T | G | 2 | 0.5703 | 0.5703 | 1.0094 | 1.0094 | 0.5067 | 0 |
| 1 | 53824428 | rs1213518 | T | C | 2 | 0.5706 | 0.5706 | 0.9901 | 0.9901 | 0.7023 | 0 |
| 1 | 53788279 | rs4136004 | T | G | 2 | 0.5707 | 0.5707 | 1.0396 | 1.0396 | 0.8974 | 0 |
| 1 | 54159359 | rs7800048 | A | T | 2 | 0.571 | 0.571 | 1.0145 | 1.0145 | 0.766 | 0 |
| 1 | 53625626 | rs1273112 | A | G | 2 | 0.5711 | 0.5711 | 1.0087 | 1.0087 | 0.8288 | 0 |
| 1 | 53962592 | rs6176845 | G | T | 2 | 0.5713 | 0.5713 | 1.0121 | 1.0121 | 0.6936 | 0 |
| 1 | 53931929 | rs1202744 | T | G | 2 | 0.5715 | 0.5715 | 0.9789 | 0.9789 | 0.3368 | 0 |
| 1 | 53493246 | rs7290313 | G | T | 2 | 0.5716 | 0.6583 | 1.0134 | 1.0171 | 0.1063 | 61.67 |
| 1 | 53493267 | rs1120244 | C | T | 2 | 0.5716 | 0.6583 | 1.0134 | 1.0171 | 0.1063 | 61.67 |
| 1 | 53491839 | rs5697822 | C | A | 2 | 0.5717 | 0.657 | 1.0134 | 1.0173 | 0.1051 | 61.92 |
| 1 | 53771861 | rs1288491 | C | T | 2 | 0.5722 | 0.5722 | 1.0096 | 1.0096 | 0.7279 | 0 |
| 1 | 53616952 | rs1120611 | C | T | 2 | 0.5725 | 0.5725 | 1.0081 | 1.0081 | 0.7298 | 0 |
| 1 | 54670060 | rs3608673 | C | G | 2 | 0.573 | 0.8635 | 1.0108 | 1.0052 | 0.1306 | 56.25 |
| 1 | 54519800 | rs7408514 | G | A | 2 | 0.5732 | 0.5732 | 0.9782 | 0.9782 | 0.5356 | 0 |
| 1 | 54442907 | rs4475694 | A | G | 2 | 0.5733 | 0.5733 | 1.0138 | 1.0138 | 0.7551 | 0 |
| 1 | 53805882 | rs1021881 | A | G | 2 | 0.5737 | 0.5737 | 1.0084 | 1.0084 | 0.5054 | 0 |
| 1 | 54542362 | rs1048944 | G | A | 2 | 0.5737 | 0.5737 | 0.9771 | 0.9771 | 0.8705 | 0 |
| 1 | 53577284 | rs1288372 | G | A | 2 | 0.5741 | 0.5741 | 1.0084 | 1.0084 | 0.9421 | 0 |
| 1 | 54216464 | rs991352 | T | A | 2 | 0.5743 | 0.7224 | 0.9657 | 0.9695 | 0.1616 | 48.95 |
| 1 | 53486737 | rs8014670 | G | A | 2 | 0.5747 | 0.6597 | 1.0133 | 1.017 | 0.1072 | 61.45 |
| 1 | 53821618 | rs1256479 | G | C | 2 | 0.575 | 0.575 | 1.0152 | 1.0152 | 0.5721 | 0 |
| 1 | 53494225 | rs6150484 | G | A | 2 | 0.5751 | 0.6561 | 1.0133 | 1.0175 | 0.104 | 62.17 |
| 1 | 54581371 | rs1206244 | A | G | 2 | 0.5754 | 0.5754 | 0.9778 | 0.9778 | 0.5027 | 0 |
| 1 | 54498824 | rs1136166 | G | T | 2 | 0.5763 | 0.5763 | 1.0395 | 1.0395 | 0.5019 | 0 |
| 1 | 53930864 | rs1203613 | C | T | 2 | 0.5766 | 0.5766 | 0.9792 | 0.9792 | 0.3414 | 0 |
| 1 | 54592618 | rs7526909 | T | C | 2 | 0.5771 | 0.6724 | 0.9707 | 0.9663 | 0.1302 | 56.34 |
| 1 | 53631805 | rs874348 | C | T | 2 | 0.5773 | 0.5773 | 1.0076 | 1.0076 | 0.5793 | 0 |
| 1 | 53610372 | rs1320594 | C | A | 2 | 0.5777 | 0.5777 | 1.0075 | 1.0075 | 0.5759 | 0 |
| 1 | 53814294 | rs4129476 | A | G | 2 | 0.5779 | 0.5779 | 0.9923 | 0.9923 | 0.7881 | 0 |
| 1 | 54112148 | rs1240848 | G | A | 2 | 0.5784 | 0.917 | 0.9911 | 0.9969 | 0.0789 | 67.61 |
| 1 | 53836391 | rs1437878 | G | A | 2 | 0.5786 | 0.5786 | 1.0232 | 1.0232 | 0.3854 | 0 |
| 1 | 53473680 | rs1393118 | C | T | 2 | 0.5788 | 0.6523 | 1.0178 | 1.0244 | 0.0962 | 63.86 |
| 1 | 53732315 | rs2297660 | G | T | 2 | 0.5788 | 0.5788 | 1.0076 | 1.0076 | 0.9451 | 0 |

| | | | | | | | | | | | |
|---|---|---|---|---|---|---|---|---|---|---|---|
| 1 | 53924164 | rs1203514 | C | T | 2 | 0.5791 | 0.5791 | 0.9793 | 0.9793 | 0.3359 | 0 |
| 1 | 54111136 | rs7603920 | C | T | 2 | 0.5791 | 0.5791 | 0.9688 | 0.9688 | 0.7774 | 0 |
| 1 | 54678305 | rs2275408 | C | G | 2 | 0.5792 | 0.8436 | 0.9592 | 1.0425 | 0.2248 | 32.13 |
| 1 | 53561589 | rs1288411 | C | T | 2 | 0.5798 | 0.6029 | 0.991 | 0.9861 | 0.1137 | 60.03 |
| 1 | 53756519 | rs1288506 | C | T | 2 | 0.5801 | 0.5801 | 0.9911 | 0.9911 | 0.4209 | 0 |
| 1 | 54060378 | rs7619769 | G | A | 2 | 0.5802 | 0.5802 | 1.0281 | 1.0281 | 0.3399 | 0 |
| 1 | 54525951 | rs7484855 | G | A | 2 | 0.5802 | 0.5802 | 0.9785 | 0.9785 | 0.4608 | 0 |
| 1 | 54553030 | rs6121049 | T | C | 2 | 0.5802 | 0.5802 | 0.9788 | 0.9788 | 0.5392 | 0 |
| 1 | 54279837 | rs1181153 | C | T | 2 | 0.5812 | 0.5812 | 1.0093 | 1.0093 | 0.9457 | 0 |
| 1 | 54580012 | rs1329780 | T | C | 2 | 0.5812 | 0.5812 | 0.9777 | 0.9777 | 0.8421 | 0 |
| 1 | 53593925 | rs6667002 | T | C | 2 | 0.5814 | 0.5814 | 1.0099 | 1.0099 | 0.8565 | 0 |
| 1 | 54576912 | rs1209397 | G | A | 2 | 0.5816 | 0.6036 | 0.9794 | 0.9781 | 0.2597 | 21.27 |
| 1 | 53587718 | rs2806271 | C | T | 2 | 0.5817 | 0.5817 | 0.9922 | 0.9922 | 0.4586 | 0 |
| 1 | 54578100 | rs9919295 | T | C | 2 | 0.5822 | 0.5822 | 0.9788 | 0.9788 | 0.5018 | 0 |
| 1 | 54490779 | rs1205816 | C | T | 2 | 0.5824 | 0.5824 | 1.0387 | 1.0387 | 0.5632 | 0 |
| 1 | 54340300 | rs1088881 | G | A | 2 | 0.5826 | 0.5826 | 1.0075 | 1.0075 | 0.4286 | 0 |
| 1 | 53516931 | rs3766765 | G | T | 2 | 0.5827 | 0.5827 | 1.0905 | 1.0905 | 0.8675 | 0 |
| 1 | 53686383 | rs1679913 | C | G | 2 | 0.5827 | 0.5827 | 1.0081 | 1.0081 | 0.3595 | 0 |
| 1 | 54047367 | rs7289930 | G | A | 2 | 0.5831 | 0.6687 | 0.952 | 0.9564 | 0.2489 | 24.79 |
| 1 | 53736613 | rs7542607 | A | G | 2 | 0.5833 | 0.5833 | 1.0075 | 1.0075 | 0.8986 | 0 |
| 1 | 53627986 | rs1203850 | T | C | 2 | 0.5834 | 0.5834 | 1.0091 | 1.0091 | 0.6168 | 0 |
| 1 | 53561414 | rs1288410 | A | G | 2 | 0.5835 | 0.6067 | 0.9911 | 0.9859 | 0.1065 | 61.63 |
| 1 | 54577508 | rs5585001 | G | A | 2 | 0.5835 | 0.5835 | 0.9789 | 0.9789 | 0.5004 | 0 |
| 1 | 54686044 | rs7534187 | C | T | 2 | 0.5835 | 0.5835 | 0.9534 | 0.9534 | 0.5632 | 0 |
| 1 | 53613907 | rs6696614 | A | T | 2 | 0.584 | 0.584 | 1.0083 | 1.0083 | 0.9581 | 0 |
| 1 | 53731265 | rs7528745 | T | A | 2 | 0.5841 | 0.5841 | 1.0074 | 1.0074 | 0.9202 | 0 |
| 1 | 54355646 | rs2272930 | T | C | 2 | 0.5841 | 0.5841 | 1.0077 | 1.0077 | 0.4508 | 0 |
| 1 | 53601803 | rs3486630 | C | T | 2 | 0.5843 | 0.5843 | 1.0083 | 1.0083 | 0.6642 | 0 |
| 1 | 53763537 | rs1572262 | C | T | 2 | 0.5845 | 0.6106 | 1.0105 | 0.9642 | 0.0253 | 80.02 |
| 1 | 53593930 | rs1679963 | C | T | 2 | 0.5847 | 0.5847 | 0.9926 | 0.9926 | 0.5813 | 0 |
| 1 | 53827004 | rs6659925 | C | T | 2 | 0.5849 | 0.5849 | 0.9906 | 0.9906 | 0.6808 | 0 |
| 1 | 53968968 | rs1240591 | G | A | 2 | 0.5853 | 0.5853 | 1.0087 | 1.0087 | 0.5744 | 0 |
| 1 | 53790373 | rs5731533 | T | C | 2 | 0.5854 | 0.5854 | 1.0398 | 1.0398 | 0.711 | 0 |
| 1 | 54578668 | rs1710978 | G | C | 2 | 0.5854 | 0.5854 | 0.979 | 0.979 | 0.5 | 0 |
| 1 | 53802455 | rs1207152 | A | G | 2 | 0.5856 | 0.5856 | 1.0337 | 1.0337 | 0.4211 | 0 |
| 1 | 54355361 | rs7830967 | G | A | 2 | 0.5861 | 0.6096 | 0.9737 | 0.9732 | 0.2779 | 15.04 |
| 1 | 53795895 | rs1133882 | C | A | 2 | 0.5865 | 0.5865 | 1.0395 | 1.0395 | 0.6199 | 0 |
| 1 | 53831346 | rs1933537 | C | T | 2 | 0.5865 | 0.5865 | 0.9908 | 0.9908 | 0.4987 | 0 |
| 1 | 54488424 | rs1209618 | C | T | 2 | 0.5868 | 0.5868 | 0.9687 | 0.9687 | 0.6379 | 0 |
| 1 | 54553608 | rs6096337 | G | A | 2 | 0.5868 | 0.5868 | 0.9791 | 0.9791 | 0.5333 | 0 |
| 1 | 54546449 | rs7556423 | G | A | 2 | 0.5871 | 0.5871 | 0.9792 | 0.9792 | 0.534 | 0 |
| 1 | 53938542 | rs1202667 | A | T | 2 | 0.5895 | 0.5895 | 1.0102 | 1.0102 | 0.4243 | 0 |
| 1 | 54549625 | rs5608169 | A | T | 2 | 0.5898 | 0.5898 | 0.9793 | 0.9793 | 0.5305 | 0 |
| 1 | 54670033 | rs3507077 | C | T | 2 | 0.5898 | 0.5898 | 0.9535 | 0.9535 | 0.6379 | 0 |
| 1 | 54685258 | rs3400832 | T | C | 2 | 0.59 | 0.9296 | 1.0103 | 1.003 | 0.0925 | 64.67 |
| 1 | 53845892 | rs6050796 | T | C | 2 | 0.5905 | 0.5905 | 0.9918 | 0.9918 | 0.7798 | 0 |
| 1 | 54629073 | rs7946078 | C | A | 2 | 0.5914 | 0.6226 | 0.9829 | 0.9823 | 0.2596 | 21.32 |
| 1 | 53494487 | rs5699826 | A | G | 2 | 0.592 | 0.663 | 1.0128 | 1.0169 | 0.1091 | 61.04 |

| | | | | | | | | | | | |
|---|---|---|---|---|---|---|---|---|---|---|---|
| 1 | 54345590 | rs1120623 | C | T | 2 | 0.5921 | 0.5921 | 1.0074 | 1.0074 | 0.4404 | 0 |
| 1 | 54276304 | rs1206473 | G | A | 2 | 0.5924 | 0.7971 | 1.0411 | 1.0449 | 0.0231 | 80.62 |
| 1 | 53734998 | rs1078895 | T | A | 2 | 0.5926 | 0.5926 | 1.0073 | 1.0073 | 0.9437 | 0 |
| 1 | 54690622 | rs1158864 | A | T | 2 | 0.5926 | 0.713 | 0.9704 | 0.9755 | 0.2399 | 27.6 |
| 1 | 53994733 | rs7528586 | G | A | 2 | 0.5929 | 0.8945 | 1.0192 | 0.9876 | 0.0151 | 83.05 |
| 1 | 53526879 | rs6131263 | T | C | 2 | 0.5932 | 0.6713 | 1.0126 | 1.0161 | 0.1129 | 60.22 |
| 1 | 53771202 | rs1049317 | A | G | 2 | 0.5943 | 0.7886 | 1.0138 | 0.9834 | 0.1325 | 55.81 |
| 1 | 54373632 | rs1205992 | C | T | 2 | 0.5943 | 0.5943 | 0.9881 | 0.9881 | 0.5939 | 0 |
| 1 | 54525150 | rs1208576 | C | A | 2 | 0.5945 | 0.5945 | 1.0372 | 1.0372 | 0.6073 | 0 |
| 1 | 53476182 | rs1134933 | G | A | 2 | 0.5948 | 0.6107 | 1.013 | 1.0135 | 0.2783 | 14.91 |
| 1 | 53593704 | rs946591 | A | G | 2 | 0.5948 | 0.5948 | 0.9928 | 0.9928 | 0.5975 | 0 |
| 1 | 54491369 | rs1118193 | G | A | 2 | 0.5954 | 0.5954 | 1.0372 | 1.0372 | 0.555 | 0 |
| 1 | 54344619 | rs2316993 | C | T | 2 | 0.5955 | 0.5955 | 1.0073 | 1.0073 | 0.4746 | 0 |
| 1 | 54610834 | rs621833 | T | C | 2 | 0.5963 | 0.5963 | 0.9906 | 0.9906 | 0.4783 | 0 |
| 1 | 53784478 | rs5894057 | T | C | 2 | 0.5964 | 0.5964 | 1.0375 | 1.0375 | 0.862 | 0 |
| 1 | 53803189 | rs7289744 | C | G | 2 | 0.5965 | 0.5965 | 0.9505 | 0.9505 | 0.6699 | 0 |
| 1 | 53731772 | rs1120612 | C | G | 2 | 0.597 | 0.597 | 1.0072 | 1.0072 | 0.9158 | 0 |
| 1 | 54314921 | rs1120622 | G | A | 2 | 0.5972 | 0.6373 | 1.0083 | 1.0128 | 0.0927 | 64.61 |
| 1 | 53633248 | rs1298238 | A | G | 2 | 0.5973 | 0.5973 | 1.0079 | 1.0079 | 0.5157 | 0 |
| 1 | 54261054 | rs5564649 | C | A | 2 | 0.5975 | 0.5975 | 1.0089 | 1.0089 | 0.9729 | 0 |
| 1 | 53992630 | rs553854 | C | T | 2 | 0.5981 | 0.7071 | 0.9807 | 0.9739 | 0.0589 | 71.97 |
| 1 | 54345822 | rs1120623 | T | C | 2 | 0.5986 | 0.5986 | 1.0072 | 1.0072 | 0.4642 | 0 |
| 1 | 53489668 | rs1710762 | C | T | 2 | 0.5989 | 0.662 | 1.0124 | 1.0151 | 0.1456 | 52.79 |
| 1 | 54342977 | rs1120623 | A | G | 2 | 0.5994 | 0.5994 | 1.0072 | 1.0072 | 0.4353 | 0 |
| 1 | 54685352 | rs3502609 | G | A | 2 | 0.5996 | 0.5996 | 0.9549 | 0.9549 | 0.549 | 0 |
| 1 | 53867792 | rs4926991 | T | C | 2 | 0.6002 | 0.6002 | 0.9902 | 0.9902 | 0.5323 | 0 |
| 1 | 54543416 | rs7408515 | C | G | 2 | 0.6002 | 0.6002 | 0.98 | 0.98 | 0.5619 | 0 |
| 1 | 54314930 | rs1120622 | T | G | 2 | 0.6005 | 0.6408 | 1.0082 | 1.0129 | 0.0882 | 65.61 |
| 1 | 53554305 | rs2306459 | G | A | 2 | 0.6007 | 0.6007 | 1.0145 | 1.0145 | 0.8402 | 0 |
| 1 | 53801793 | rs7289743 | C | T | 2 | 0.6007 | 0.6007 | 0.951 | 0.951 | 0.6732 | 0 |
| 1 | 54578136 | rs9919296 | T | C | 2 | 0.6007 | 0.6243 | 0.9805 | 0.9789 | 0.2502 | 24.35 |
| 1 | 54677862 | rs3766457 | A | G | 2 | 0.6017 | 0.6146 | 1.0094 | 1.0177 | 0.064 | 70.85 |
| 1 | 53832114 | rs6658952 | C | G | 2 | 0.6019 | 0.6019 | 0.9912 | 0.9912 | 0.462 | 0 |
| 1 | 53697139 | rs6673692 | A | G | 2 | 0.602 | 0.602 | 1.0071 | 1.0071 | 0.3668 | 0 |
| 1 | 53525266 | rs965097 | A | C | 2 | 0.6021 | 0.6781 | 1.0123 | 1.0158 | 0.1119 | 60.44 |
| 1 | 53934667 | rs1128123 | G | A | 2 | 0.6021 | 0.6099 | 0.9806 | 0.9807 | 0.3081 | 3.73 |
| 1 | 54161166 | rs1143697 | C | A | 2 | 0.6022 | 0.6022 | 1.0245 | 1.0245 | 0.5371 | 0 |
| 1 | 53495410 | rs1710766 | C | T | 2 | 0.6025 | 0.6771 | 1.0123 | 1.0157 | 0.1131 | 60.16 |
| 1 | 54569582 | rs1209381 | A | G | 2 | 0.6025 | 0.6025 | 0.9796 | 0.9796 | 0.3858 | 0 |
| 1 | 54571644 | rs4631638 | C | G | 2 | 0.6027 | 0.6027 | 0.979 | 0.979 | 0.6848 | 0 |
| 1 | 53490682 | rs7290311 | G | A | 2 | 0.6029 | 0.6672 | 1.0124 | 1.0165 | 0.1123 | 60.35 |
| 1 | 54295292 | rs1181174 | T | C | 2 | 0.6032 | 0.6032 | 1.0088 | 1.0088 | 0.9126 | 0 |
| 1 | 53532535 | rs1710780 | G | C | 2 | 0.6034 | 0.6034 | 1.0185 | 1.0185 | 0.516 | 0 |
| 1 | 54505703 | rs8025559 | G | A | 2 | 0.6041 | 0.6041 | 1.0364 | 1.0364 | 0.5385 | 0 |
| 1 | 53993341 | rs3893165 | C | A | 2 | 0.6047 | 0.7244 | 1.0382 | 1.0356 | 0.171 | 46.65 |
| 1 | 53825758 | rs3414683 | C | T | 2 | 0.6055 | 0.6055 | 0.9914 | 0.9914 | 0.6505 | 0 |
| 1 | 53852449 | rs6677153 | C | T | 2 | 0.6058 | 0.6058 | 1.0072 | 1.0072 | 0.6698 | 0 |
| 1 | 54633847 | rs1710998 | C | T | 2 | 0.606 | 0.5413 | 0.9736 | 0.9382 | 0.0807 | 67.22 |

| | | | | | | | | | | | |
|---|---|---|---|---|---|---|---|---|---|---|---|
| 1 | 53478189 | rs7956629 | G | C | 2 | 0.6064 | 0.6805 | 1.0163 | 1.0257 | 0.0537 | 73.14 |
| 1 | 53829820 | rs1999898 | C | T | 2 | 0.6067 | 0.6067 | 0.9913 | 0.9913 | 0.4892 | 0 |
| 1 | 53607004 | rs1273822 | C | T | 2 | 0.6069 | 0.8563 | 1.007 | 1.0039 | 0.1212 | 58.37 |
| 1 | 54547293 | rs1120628 | G | A | 2 | 0.6069 | 0.5882 | 0.9517 | 0.9418 | 0.2759 | 15.77 |
| 1 | 53490221 | rs1710763 | T | C | 2 | 0.6073 | 0.6735 | 1.0122 | 1.0158 | 0.1179 | 59.11 |
| 1 | 54046682 | rs7289929 | G | A | 2 | 0.6073 | 0.6787 | 0.9546 | 0.9582 | 0.2569 | 22.2 |
| 1 | 54112410 | rs1379373 | T | C | 2 | 0.6077 | 0.6077 | 1.0301 | 1.0301 | 0.8459 | 0 |
| 1 | 54558276 | rs7530819 | T | G | 2 | 0.6078 | 0.6078 | 0.9803 | 0.9803 | 0.515 | 0 |
| 1 | 54680135 | rs3561024 | C | T | 2 | 0.6081 | 0.6081 | 0.955 | 0.955 | 0.6301 | 0 |
| 1 | 53813684 | rs1088878 | A | G | 2 | 0.6091 | 0.6091 | 0.9926 | 0.9926 | 0.5237 | 0 |
| 1 | 54284814 | rs1208920 | C | T | 2 | 0.6094 | 0.6094 | 1.0086 | 1.0086 | 0.8743 | 0 |
| 1 | 53486558 | rs7463739 | C | T | 2 | 0.6095 | 0.712 | 1.0123 | 1.0159 | 0.0773 | 67.95 |
| 1 | 53846850 | rs7514050 | T | G | 2 | 0.6097 | 0.6097 | 1.007 | 1.007 | 0.552 | 0 |
| 1 | 54584679 | rs7508455 | C | G | 2 | 0.6098 | 0.6098 | 1.0436 | 1.0436 | 0.7983 | 0 |
| 1 | 54637151 | rs8029568 | A | G | 2 | 0.6102 | 0.6102 | 0.9666 | 0.9666 | 0.4115 | 0 |
| 1 | 54512553 | rs4152874 | A | G | 2 | 0.611 | 0.611 | 0.9764 | 0.9764 | 0.9848 | 0 |
| 1 | 54343342 | rs7548793 | G | A | 2 | 0.6111 | 0.6111 | 1.007 | 1.007 | 0.4761 | 0 |
| 1 | 53609496 | rs1120611 | C | T | 2 | 0.6112 | 0.6112 | 1.007 | 1.007 | 0.3729 | 0 |
| 1 | 53786174 | rs915192 | C | T | 2 | 0.6112 | 0.6112 | 1.0358 | 1.0358 | 0.9383 | 0 |
| 1 | 53795290 | rs7289741 | C | T | 2 | 0.6119 | 0.6119 | 1.037 | 1.037 | 0.6561 | 0 |
| 1 | 53551703 | rs1202872 | G | A | 2 | 0.6123 | 0.6123 | 0.9864 | 0.9864 | 0.8008 | 0 |
| 1 | 54558819 | rs7528941 | A | G | 2 | 0.6123 | 0.6123 | 0.9803 | 0.9803 | 0.4261 | 0 |
| 1 | 54289699 | rs1208881 | C | A | 2 | 0.6124 | 0.8214 | 1.0377 | 1.0383 | 0.0223 | 80.84 |
| 1 | 53544439 | rs1288388 | G | A | 2 | 0.6125 | 0.6125 | 0.9923 | 0.9912 | 0.2612 | 20.8 |
| 1 | 54096591 | rs4926607 | C | T | 2 | 0.6125 | 0.8262 | 0.992 | 0.995 | 0.167 | 47.64 |
| 1 | 53970912 | rs1207819 | G | A | 2 | 0.6126 | 0.6593 | 0.9882 | 0.9801 | 0.0599 | 71.76 |
| 1 | 54684786 | rs3457639 | C | A | 2 | 0.613 | 0.613 | 0.9565 | 0.9565 | 0.5592 | 0 |
| 1 | 53603796 | rs2404210 | G | A | 2 | 0.6131 | 0.6131 | 1.0076 | 1.0076 | 0.6408 | 0 |
| 1 | 54608400 | rs1571542 | T | G | 2 | 0.6131 | 0.6131 | 0.9907 | 0.9907 | 0.6638 | 0 |
| 1 | 53795619 | rs7289741 | A | G | 2 | 0.6132 | 0.6132 | 1.0368 | 1.0368 | 0.6515 | 0 |
| 1 | 53500429 | rs1710768 | A | G | 2 | 0.6136 | 0.6136 | 1.0742 | 1.0742 | 0.9371 | 0 |
| 1 | 54608399 | rs4333802 | A | C | 2 | 0.6136 | 0.6136 | 0.9908 | 0.9908 | 0.668 | 0 |
| 1 | 54024808 | rs4575030 | C | T | 2 | 0.6138 | 0.658 | 0.9931 | 0.9888 | 0.0694 | 69.66 |
| 1 | 54500221 | rs1206485 | C | T | 2 | 0.6138 | 0.6138 | 1.0353 | 1.0353 | 0.5303 | 0 |
| 1 | 53548295 | rs899973 | C | G | 2 | 0.6139 | 0.6138 | 0.9923 | 0.9909 | 0.2479 | 25.1 |
| 1 | 53835594 | rs4926986 | A | G | 2 | 0.6141 | 0.6163 | 0.9928 | 0.9924 | 0.2932 | 9.47 |
| 1 | 54520603 | rs1157839 | T | C | 2 | 0.6141 | 0.6141 | 0.9851 | 0.9851 | 0.8488 | 0 |
| 1 | 53633748 | rs3125248 | G | A | 2 | 0.6147 | 0.6147 | 0.9903 | 0.9903 | 0.425 | 0 |
| 1 | 54412181 | rs1063162 | T | C | 2 | 0.6147 | 0.9637 | 1.0094 | 1.002 | 0.0182 | 82.07 |
| 1 | 53562806 | rs1288414 | C | T | 2 | 0.6151 | 0.6257 | 0.9918 | 0.9868 | 0.1098 | 60.88 |
| 1 | 53784484 | rs5932734 | T | C | 2 | 0.6153 | 0.6153 | 1.0355 | 1.0355 | 0.8448 | 0 |
| 1 | 53933534 | rs5616139 | G | A | 2 | 0.6153 | 0.6273 | 0.9813 | 0.9814 | 0.3028 | 5.84 |
| 1 | 53865066 | rs6711785 | C | T | 2 | 0.6154 | 0.6154 | 1.0225 | 1.0225 | 0.9952 | 0 |
| 1 | 54686413 | rs7709752 | C | G | 2 | 0.6155 | 0.6155 | 0.9568 | 0.9568 | 0.5583 | 0 |
| 1 | 53582325 | rs1679911 | A | C | 2 | 0.6156 | 0.6156 | 0.9928 | 0.9928 | 0.6462 | 0 |
| 1 | 53496675 | rs1710766 | A | C | 2 | 0.6159 | 0.6962 | 1.0118 | 1.016 | 0.0851 | 66.27 |
| 1 | 53635094 | rs1209693 | A | G | 2 | 0.6159 | 0.6159 | 0.9748 | 0.9748 | 0.8826 | 0 |
| 1 | 53494516 | rs6144128 | T | C | 2 | 0.616 | 0.6782 | 1.0119 | 1.0154 | 0.1215 | 58.31 |

| | | | | | | | | | | | |
|---|---|---|---|---|---|---|---|---|---|---|---|
| 1 | 54558101 | rs4926619 | T | C | 2 | 0.6162 | 0.6162 | 0.9931 | 0.9931 | 0.632 | 0 |
| 1 | 54048595 | rs6132649 | G | T | 2 | 0.6165 | 0.6323 | 0.9558 | 0.9566 | 0.3043 | 5.22 |
| 1 | 53852469 | rs1120615 | A | G | 2 | 0.6168 | 0.6168 | 1.0067 | 1.0067 | 0.6376 | 0 |
| 1 | 54633543 | rs1710996 | C | T | 2 | 0.6169 | 0.5444 | 0.9744 | 0.9365 | 0.0706 | 69.41 |
| 1 | 53490614 | rs7290311 | T | C | 2 | 0.617 | 0.6745 | 1.0119 | 1.0159 | 0.118 | 59.08 |
| 1 | 53567361 | rs1120609 | C | A | 2 | 0.6173 | 0.618 | 0.9924 | 0.9893 | 0.1725 | 46.28 |
| 1 | 54450490 | rs638749 | A | G | 2 | 0.6173 | 0.6173 | 1.0068 | 1.0068 | 0.5596 | 0 |
| 1 | 54492569 | rs1154 | T | C | 2 | 0.618 | 0.618 | 0.9805 | 0.9805 | 0.5679 | 0 |
| 1 | 53627891 | rs6665477 | C | T | 2 | 0.6181 | 0.6181 | 1.0078 | 1.0078 | 0.8695 | 0 |
| 1 | 54633630 | rs1710997 | C | G | 2 | 0.6181 | 0.6181 | 0.9779 | 0.9779 | 0.4965 | 0 |
| 1 | 54552238 | rs7408516 | A | G | 2 | 0.6184 | 0.6184 | 0.9808 | 0.9808 | 0.5623 | 0 |
| 1 | 53738100 | rs1078895 | C | T | 2 | 0.6186 | 0.6186 | 1.0068 | 1.0068 | 0.8911 | 0 |
| 1 | 53494927 | rs1710765 | G | C | 2 | 0.6187 | 0.6821 | 1.0117 | 1.0149 | 0.1264 | 57.19 |
| 1 | 53763374 | rs1288495 | T | C | 2 | 0.619 | 0.5517 | 1.0096 | 0.9491 | 0.0073 | 86.09 |
| 1 | 53632424 | rs2046140 | G | A | 2 | 0.6192 | 0.6192 | 1.0085 | 1.0085 | 0.7258 | 0 |
| 1 | 53825618 | rs4926978 | T | C | 2 | 0.6193 | 0.6193 | 0.9918 | 0.9918 | 0.6435 | 0 |
| 1 | 53490399 | rs5787169 | A | G | 2 | 0.6195 | 0.675 | 1.0118 | 1.0159 | 0.1182 | 59.04 |
| 1 | 53490705 | rs7290311 | G | A | 2 | 0.6195 | 0.6756 | 1.0118 | 1.0157 | 0.1202 | 58.59 |
| 1 | 53490345 | rs7751434 | C | T | 2 | 0.6197 | 0.6742 | 1.0119 | 1.016 | 0.1174 | 59.2 |
| 1 | 54480903 | rs6691549 | A | G | 2 | 0.6197 | 0.6197 | 0.9932 | 0.9932 | 0.5626 | 0 |
| 1 | 53794969 | rs5857286 | A | G | 2 | 0.6198 | 0.6198 | 1.0362 | 1.0362 | 0.6527 | 0 |
| 1 | 54412164 | rs1905593 | G | A | 2 | 0.6198 | 0.6198 | 0.9795 | 0.9795 | 0.8845 | 0 |
| 1 | 54685854 | rs3467337 | C | T | 2 | 0.6198 | 0.6198 | 0.9573 | 0.9573 | 0.5604 | 0 |
| 1 | 53794258 | rs7289741 | A | C | 2 | 0.6199 | 0.6199 | 1.0361 | 1.0361 | 0.6599 | 0 |
| 1 | 53791873 | rs1710833 | C | G | 2 | 0.62 | 0.62 | 1.0363 | 1.0363 | 0.7624 | 0 |
| 1 | 54598299 | rs1389334 | A | G | 2 | 0.6204 | 0.6204 | 0.974 | 0.974 | 0.4797 | 0 |
| 1 | 53606080 | rs1120610 | C | T | 2 | 0.6217 | 0.6217 | 1.0074 | 1.0074 | 0.6581 | 0 |
| 1 | 54235082 | rs1181199 | T | C | 2 | 0.6217 | 0.6217 | 1.0084 | 1.0084 | 0.8599 | 0 |
| 1 | 53489087 | rs4130007 | C | G | 2 | 0.6218 | 0.7029 | 1.0117 | 1.0157 | 0.0862 | 66.03 |
| 1 | 53490369 | rs5709363 | G | A | 2 | 0.6219 | 0.6767 | 1.0118 | 1.0157 | 0.1197 | 58.71 |
| 1 | 54579134 | rs9919314 | T | C | 2 | 0.6219 | 0.6219 | 0.9811 | 0.9811 | 0.5695 | 0 |
| 1 | 54678927 | rs3766456 | G | A | 2 | 0.6222 | 0.6222 | 0.9568 | 0.9568 | 0.6337 | 0 |
| 1 | 53723965 | rs2297661 | C | G | 2 | 0.6223 | 0.6223 | 0.9664 | 0.9664 | 0.8907 | 0 |
| 1 | 53796395 | rs7289742 | C | T | 2 | 0.6226 | 0.6226 | 1.0358 | 1.0358 | 0.6371 | 0 |
| 1 | 53860821 | rs6176838 | G | A | 2 | 0.6232 | 0.6232 | 0.9896 | 0.9896 | 0.5306 | 0 |
| 1 | 53794114 | rs7535616 | A | C | 2 | 0.6235 | 0.6235 | 1.0358 | 1.0358 | 0.6585 | 0 |
| 1 | 54670047 | rs6680918 | C | T | 2 | 0.6235 | 0.6235 | 1.0118 | 1.0118 | 0.9192 | 0 |
| 1 | 53516999 | rs8871 | A | C | 2 | 0.6237 | 0.6967 | 1.0116 | 1.0152 | 0.101 | 62.81 |
| 1 | 53868891 | rs1288590 | A | G | 2 | 0.6237 | 0.6237 | 0.9933 | 0.9933 | 0.8827 | 0 |
| 1 | 54432019 | rs2294514 | G | A | 2 | 0.6237 | 0.967 | 1.0091 | 1.0018 | 0.0224 | 80.82 |
| 1 | 54260903 | rs3546096 | T | C | 2 | 0.624 | 0.624 | 1.0083 | 1.0083 | 0.862 | 0 |
| 1 | 53517349 | rs6057666 | G | A | 2 | 0.6241 | 0.696 | 1.0116 | 1.0153 | 0.1008 | 62.86 |
| 1 | 53796237 | rs7289742 | C | T | 2 | 0.6241 | 0.6241 | 1.0357 | 1.0357 | 0.6375 | 0 |
| 1 | 54677343 | rs3486679 | C | T | 2 | 0.6241 | 0.6241 | 0.9572 | 0.9572 | 0.6413 | 0 |
| 1 | 53516742 | rs13496 | G | A | 2 | 0.6243 | 0.6961 | 1.0116 | 1.0153 | 0.1008 | 62.87 |
| 1 | 53518658 | rs7290513 | G | A | 2 | 0.6243 | 0.6961 | 1.0116 | 1.0153 | 0.1008 | 62.87 |
| 1 | 53801359 | rs7289743 | A | C | 2 | 0.6245 | 0.6245 | 0.9538 | 0.9538 | 0.6407 | 0 |
| 1 | 54351135 | rs1158612 | C | T | 2 | 0.6247 | 0.7755 | 0.9853 | 0.9886 | 0.1893 | 41.96 |

| | | | | | | | | | | | |
|---|---|---|---|---|---|---|---|---|---|---|---|
| 1 | 53566818 | rs1273741 | C | T | 2 | 0.6249 | 0.6249 | 0.9905 | 0.9905 | 0.7685 | 0 |
| 1 | 53488234 | rs6588449 | G | A | 2 | 0.625 | 0.7055 | 1.0116 | 1.0156 | 0.0848 | 66.33 |
| 1 | 53797058 | rs7289742 | C | T | 2 | 0.6251 | 0.6251 | 0.9452 | 0.9452 | 0.9505 | 0 |
| 1 | 53731019 | rs7526226 | T | A | 2 | 0.6254 | 0.6254 | 1.0066 | 1.0066 | 0.9521 | 0 |
| 1 | 53826343 | rs4926596 | C | T | 2 | 0.6255 | 0.6255 | 0.992 | 0.992 | 0.6351 | 0 |
| 1 | 54339727 | rs1120623 | T | A | 2 | 0.6256 | 0.6256 | 1.0067 | 1.0067 | 0.3587 | 0 |
| 1 | 53494514 | rs6139805 | C | A | 2 | 0.6257 | 0.6832 | 1.0116 | 1.015 | 0.1254 | 57.43 |
| 1 | 53562221 | rs6693952 | C | T | 2 | 0.626 | 0.8166 | 1.0214 | 1.0172 | 0.092 | 64.78 |
| 1 | 53792961 | rs7289741 | T | G | 2 | 0.6264 | 0.6264 | 1.0355 | 1.0355 | 0.6683 | 0 |
| 1 | 54669050 | rs1180412 | A | G | 2 | 0.6264 | 0.9408 | 1.0093 | 1.0024 | 0.0996 | 63.13 |
| 1 | 53868471 | rs1288591 | C | G | 2 | 0.6265 | 0.6265 | 0.9933 | 0.9933 | 0.9081 | 0 |
| 1 | 54286763 | rs1181190 | C | G | 2 | 0.6269 | 0.6269 | 1.0082 | 1.0082 | 0.9087 | 0 |
| 1 | 54271830 | rs1753650 | T | C | 2 | 0.627 | 0.627 | 1.0146 | 1.0146 | 0.7363 | 0 |
| 1 | 54267199 | rs1181176 | C | T | 2 | 0.6271 | 0.6271 | 1.0082 | 1.0082 | 0.8942 | 0 |
| 1 | 53765250 | rs3542382 | C | A | 2 | 0.6273 | 0.4835 | 0.9905 | 1.0872 | 0.0047 | 87.47 |
| 1 | 54268873 | rs1181148 | C | G | 2 | 0.6274 | 0.6274 | 1.0082 | 1.0082 | 0.898 | 0 |
| 1 | 54562966 | rs7527305 | T | C | 2 | 0.6275 | 0.6275 | 0.9814 | 0.9814 | 0.4097 | 0 |
| 1 | 53792821 | rs7289740 | C | T | 2 | 0.6279 | 0.6279 | 1.0353 | 1.0353 | 0.6681 | 0 |
| 1 | 53579583 | rs1288375 | G | A | 2 | 0.6283 | 0.5962 | 1.0081 | 1.0118 | 0.2773 | 15.29 |
| 1 | 54356113 | rs6177483 | C | T | 2 | 0.6292 | 0.9825 | 0.993 | 1.0007 | 0.0266 | 79.66 |
| 1 | 53802279 | rs1125206 | A | G | 2 | 0.6293 | 0.6293 | 1.0364 | 1.0364 | 0.6227 | 0 |
| 1 | 54461263 | rs664758 | C | T | 2 | 0.6299 | 0.6299 | 1.0066 | 1.0066 | 0.5 | 0 |
| 1 | 53553069 | rs1047741 | G | A | 2 | 0.6302 | 0.7727 | 1.019 | 1.0162 | 0.1568 | 50.13 |
| 1 | 53567546 | rs3766775 | A | G | 2 | 0.6302 | 0.6275 | 0.9927 | 0.9892 | 0.1599 | 49.36 |
| 1 | 53792456 | rs1271842 | G | C | 2 | 0.6302 | 0.6302 | 1.0351 | 1.0351 | 0.6699 | 0 |
| 1 | 54092573 | rs1240526 | G | A | 2 | 0.6303 | 0.8123 | 0.9924 | 0.9949 | 0.1906 | 41.63 |
| 1 | 54668427 | rs946445 | G | T | 2 | 0.631 | 0.9484 | 1.0092 | 1.0022 | 0.096 | 63.91 |
| 1 | 54653291 | rs6177522 | C | T | 2 | 0.6311 | 0.6311 | 1.0113 | 1.0113 | 0.3557 | 0 |
| 1 | 53562220 | rs1288412 | C | G | 2 | 0.6312 | 0.6314 | 0.9922 | 0.9885 | 0.1558 | 50.36 |
| 1 | 53793452 | rs1137594 | T | C | 2 | 0.6314 | 0.6314 | 1.0344 | 1.0344 | 0.6868 | 0 |
| 1 | 54677366 | rs3407857 | A | T | 2 | 0.6319 | 0.6319 | 0.958 | 0.958 | 0.6482 | 0 |
| 1 | 54216654 | rs761490 | C | G | 2 | 0.6321 | 0.6321 | 0.9923 | 0.9923 | 0.822 | 0 |
| 1 | 53815537 | rs4926976 | G | A | 2 | 0.6323 | 0.6323 | 0.9935 | 0.9935 | 0.9596 | 0 |
| 1 | 53503918 | rs1394221 | G | A | 2 | 0.6327 | 0.6327 | 0.9758 | 0.9758 | 0.4951 | 0 |
| 1 | 54488250 | rs3510929 | A | G | 2 | 0.6328 | 0.6328 | 0.9935 | 0.9935 | 0.3969 | 0 |
| 1 | 53545763 | rs1288393 | G | A | 2 | 0.6333 | 0.631 | 0.9927 | 0.9919 | 0.2751 | 16.04 |
| 1 | 53693627 | rs1208230 | T | G | 2 | 0.6333 | 0.6333 | 0.9672 | 0.9672 | 0.8293 | 0 |
| 1 | 53696784 | rs6676003 | T | G | 2 | 0.6334 | 0.6334 | 1.0064 | 1.0064 | 0.4548 | 0 |
| 1 | 54683164 | rs3596292 | A | C | 2 | 0.6337 | 0.6337 | 0.9588 | 0.9588 | 0.5742 | 0 |
| 1 | 54400627 | rs1120625 | G | A | 2 | 0.6339 | 0.6339 | 0.9893 | 0.9893 | 0.5005 | 0 |
| 1 | 53742219 | rs1202793 | T | C | 2 | 0.6342 | 0.6342 | 0.9671 | 0.9671 | 0.8386 | 0 |
| 1 | 54009376 | rs9700109 | G | A | 2 | 0.6347 | 0.6347 | 0.9921 | 0.9921 | 0.5069 | 0 |
| 1 | 54578016 | rs5890801 | C | T | 2 | 0.6348 | 0.6348 | 0.9808 | 0.9808 | 0.8321 | 0 |
| 1 | 53554736 | rs5860078 | C | T | 2 | 0.6349 | 0.7749 | 1.0187 | 1.0158 | 0.1612 | 49.05 |
| 1 | 54665124 | rs946447 | T | C | 2 | 0.636 | 0.636 | 1.011 | 1.011 | 0.7429 | 0 |
| 1 | 54677696 | rs3766458 | G | T | 2 | 0.6367 | 0.6329 | 1.0085 | 1.0168 | 0.0636 | 70.94 |
| 1 | 53879615 | rs1158033 | T | A | 2 | 0.6369 | 0.6369 | 0.9918 | 0.9918 | 0.4278 | 0 |
| 1 | 54646921 | rs6177522 | T | A | 2 | 0.6369 | 0.6369 | 1.0111 | 1.0111 | 0.3676 | 0 |

| | | | | | | | | | | | |
|---|---|---|---|---|---|---|---|---|---|---|---|
| 1 | 53757154 | rs1288502 | C | A | 2 | 0.6378 | 0.6378 | 0.9924 | 0.9924 | 0.4922 | 0 |
| 1 | 54587121 | rs1710980 | T | C | 2 | 0.6378 | 0.6325 | 0.9738 | 0.9684 | 0.244 | 26.34 |
| 1 | 53972132 | rs2296759 | A | G | 2 | 0.6379 | 0.6379 | 0.9906 | 0.9906 | 0.3849 | 0 |
| 1 | 53502301 | rs8023263 | A | G | 2 | 0.639 | 0.6945 | 1.0146 | 1.0244 | 0.0511 | 73.71 |
| 1 | 53616509 | rs1214390 | C | A | 2 | 0.6394 | 0.6394 | 1.0071 | 1.0071 | 0.9813 | 0 |
| 1 | 54326301 | rs1181096 | C | A | 2 | 0.6395 | 0.6395 | 1.0065 | 1.0065 | 0.4449 | 0 |
| 1 | 53921571 | rs4144364 | T | C | 2 | 0.6403 | 0.6731 | 0.9925 | 0.9899 | 0.1432 | 53.35 |
| 1 | 54519107 | rs2236512 | G | A | 2 | 0.6403 | 0.6403 | 0.9824 | 0.9824 | 0.614 | 0 |
| 1 | 54257376 | rs1710937 | G | C | 2 | 0.6408 | 0.6408 | 1.0079 | 1.0079 | 0.7419 | 0 |
| 1 | 54532435 | rs1710970 | T | C | 2 | 0.6409 | 0.6409 | 0.982 | 0.982 | 0.3552 | 0 |
| 1 | 53784688 | rs1120614 | A | C | 2 | 0.641 | 0.773 | 0.9553 | 0.9628 | 0.183 | 43.59 |
| 1 | 54644859 | rs6173885 | C | T | 2 | 0.641 | 0.641 | 1.0109 | 1.0109 | 0.3531 | 0 |
| 1 | 53545479 | rs1288391 | C | A | 2 | 0.6413 | 0.6371 | 0.9929 | 0.9915 | 0.246 | 25.7 |
| 1 | 54660979 | rs1120629 | T | C | 2 | 0.6414 | 0.8186 | 1.0278 | 1.0174 | 0.2227 | 32.75 |
| 1 | 54224072 | rs2316321 | A | G | 2 | 0.6415 | 0.6308 | 1.0092 | 1.0115 | 0.2375 | 28.34 |
| 1 | 54337886 | rs1088881 | G | A | 2 | 0.642 | 0.642 | 1.0062 | 1.0062 | 0.3268 | 0 |
| 1 | 53495624 | rs1421513 | G | A | 2 | 0.6421 | 0.672 | 1.0147 | 1.0276 | 0.0486 | 74.28 |
| 1 | 53520889 | rs4431784 | T | G | 2 | 0.6421 | 0.7016 | 1.011 | 1.0144 | 0.116 | 59.53 |
| 1 | 53992842 | rs521604 | G | A | 2 | 0.6425 | 0.7118 | 0.9831 | 0.9778 | 0.1015 | 62.71 |
| 1 | 53676448 | rs1799821 | G | A | 2 | 0.6438 | 0.6438 | 1.0062 | 1.0062 | 0.3219 | 0 |
| 1 | 54372232 | rs7586151 | C | T | 2 | 0.6441 | 0.6441 | 1.0375 | 1.0375 | 0.972 | 0 |
| 1 | 53527619 | rs1120608 | T | A | 2 | 0.6446 | 0.6446 | 1.0139 | 1.0139 | 0.5838 | 0 |
| 1 | 54049333 | rs6588481 | T | C | 2 | 0.6447 | 0.6685 | 0.9592 | 0.9604 | 0.297 | 8.05 |
| 1 | 54538700 | rs7537946 | A | G | 2 | 0.6448 | 0.6448 | 0.9938 | 0.9938 | 0.6354 | 0 |
| 1 | 54375570 | rs2235544 | C | A | 2 | 0.6452 | 0.6452 | 1.0062 | 1.0062 | 0.7928 | 0 |
| 1 | 53767951 | rs1203795 | T | C | 2 | 0.6454 | 0.4881 | 0.9911 | 1.0824 | 0.0037 | 88.15 |
| 1 | 53800114 | rs7289743 | G | A | 2 | 0.6455 | 0.6455 | 1.0327 | 1.0327 | 0.7182 | 0 |
| 1 | 54482236 | rs1537321 | C | A | 2 | 0.6461 | 0.6461 | 0.9936 | 0.9936 | 0.7242 | 0 |
| 1 | 53520736 | rs1710777 | C | T | 2 | 0.6464 | 0.7049 | 1.0108 | 1.0142 | 0.1148 | 59.79 |
| 1 | 53720574 | rs7289532 | G | A | 2 | 0.6466 | 0.6466 | 0.9912 | 0.9912 | 0.4914 | 0 |
| 1 | 54324579 | rs1120622 | G | A | 2 | 0.6466 | 0.6466 | 1.0063 | 1.0063 | 0.3884 | 0 |
| 1 | 53845390 | rs1157899 | C | G | 2 | 0.6469 | 0.6469 | 0.9931 | 0.9931 | 0.8611 | 0 |
| 1 | 53626734 | rs7516971 | C | G | 2 | 0.6472 | 0.6472 | 1.0071 | 1.0071 | 0.7671 | 0 |
| 1 | 54270883 | rs1472632 | G | A | 2 | 0.6475 | 0.6475 | 0.9687 | 0.9687 | 0.5359 | 0 |
| 1 | 53764060 | rs1158374 | G | T | 2 | 0.6485 | 0.6485 | 1.0131 | 1.0131 | 0.4761 | 0 |
| 1 | 53617475 | rs1202928 | G | A | 2 | 0.6493 | 0.6493 | 1.0075 | 1.0075 | 0.4742 | 0 |
| 1 | 54264996 | rs1181180 | T | C | 2 | 0.6497 | 0.6497 | 1.0077 | 1.0077 | 0.9104 | 0 |
| 1 | 53633353 | rs4500254 | A | G | 2 | 0.6499 | 0.6499 | 0.9931 | 0.9931 | 0.5837 | 0 |
| 1 | 53495634 | rs1507791 | C | T | 2 | 0.6502 | 0.6755 | 1.0144 | 1.027 | 0.0509 | 73.77 |
| 1 | 54262163 | rs9436477 | A | G | 2 | 0.6509 | 0.6509 | 1.0077 | 1.0077 | 0.8928 | 0 |
| 1 | 53607666 | rs2404211 | T | C | 2 | 0.6514 | 0.6514 | 1.0064 | 1.0064 | 0.9892 | 0 |
| 1 | 54269087 | rs1181147 | G | A | 2 | 0.6516 | 0.6516 | 1.0076 | 1.0076 | 0.8599 | 0 |
| 1 | 54567371 | rs7494118 | A | G | 2 | 0.6519 | 0.6519 | 0.9818 | 0.9818 | 0.7248 | 0 |
| 1 | 53489498 | rs1710761 | C | G | 2 | 0.652 | 0.6973 | 1.014 | 1.0234 | 0.0589 | 71.97 |
| 1 | 53488641 | rs7750977 | A | C | 2 | 0.6521 | 0.6969 | 1.014 | 1.0234 | 0.0595 | 71.85 |
| 1 | 53567207 | rs3432262 | G | A | 2 | 0.6521 | 0.6436 | 0.9931 | 0.991 | 0.2167 | 34.47 |
| 1 | 53819026 | rs2027261 | G | A | 2 | 0.6523 | 0.6523 | 0.9936 | 0.9936 | 0.4788 | 0 |
| 1 | 54484622 | rs1141148 | T | C | 2 | 0.6529 | 0.6529 | 1.0216 | 1.0216 | 0.7908 | 0 |

| | | | | | | | | | | | |
|---|---|---|---|---|---|---|---|---|---|---|---|
| 1 | 54547953 | rs7408516 | A | G | 2 | 0.6532 | 0.6532 | 0.9818 | 0.9818 | 0.7744 | 0 |
| 1 | 54568488 | rs4492560 | G | A | 2 | 0.6533 | 0.6533 | 0.9939 | 0.9939 | 0.534 | 0 |
| 1 | 53485330 | rs8014580 | C | T | 2 | 0.6534 | 0.6979 | 1.0139 | 1.0233 | 0.0593 | 71.88 |
| 1 | 54468633 | rs1167265 | T | C | 2 | 0.6534 | 0.6534 | 0.9806 | 0.9806 | 0.349 | 0 |
| 1 | 53552319 | rs1288398 | C | T | 2 | 0.6536 | 0.6478 | 0.9932 | 0.992 | 0.2609 | 20.88 |
| 1 | 53557528 | rs1148298 | A | G | 2 | 0.6537 | 0.6537 | 0.9794 | 0.9794 | 0.8651 | 0 |
| 1 | 53565087 | rs1769287 | C | T | 2 | 0.6537 | 0.6452 | 0.9931 | 0.9913 | 0.2273 | 31.4 |
| 1 | 53487808 | rs7547031 | C | T | 2 | 0.6539 | 0.6977 | 1.0139 | 1.0234 | 0.0593 | 71.89 |
| 1 | 53489320 | rs7690015 | C | T | 2 | 0.6549 | 0.6989 | 1.0139 | 1.0233 | 0.0585 | 72.06 |
| 1 | 53878602 | rs7894697 | G | A | 2 | 0.6553 | 0.6553 | 0.9923 | 0.9923 | 0.417 | 0 |
| 1 | 53708850 | rs3737984 | G | T | 2 | 0.6555 | 0.6555 | 1.006 | 1.006 | 0.6562 | 0 |
| 1 | 53477323 | rs7912892 | T | C | 2 | 0.656 | 0.701 | 1.0141 | 1.022 | 0.0757 | 68.31 |
| 1 | 53581670 | rs3820201 | A | G | 2 | 0.6562 | 0.6562 | 1.0065 | 1.0065 | 0.5095 | 0 |
| 1 | 54636689 | rs7552121 | A | G | 2 | 0.6566 | 0.8824 | 1.009 | 1.0048 | 0.1138 | 60 |
| 1 | 53671036 | rs1120612 | A | C | 2 | 0.6574 | 0.6574 | 1.006 | 1.006 | 0.3436 | 0 |
| 1 | 53713336 | rs1078895 | A | G | 2 | 0.6574 | 0.6574 | 1.0064 | 1.0064 | 0.5666 | 0 |
| 1 | 53630304 | rs6588467 | G | A | 2 | 0.6578 | 0.6578 | 1.006 | 1.006 | 0.6675 | 0 |
| 1 | 54566758 | rs5580654 | T | C | 2 | 0.6582 | 0.6582 | 0.9831 | 0.9831 | 0.4866 | 0 |
| 1 | 54557263 | rs1209425 | C | A | 2 | 0.6585 | 0.6585 | 1.0308 | 1.0308 | 0.7052 | 0 |
| 1 | 54675001 | rs3546524 | G | C | 2 | 0.6593 | 0.9827 | 1.0084 | 1.0008 | 0.0816 | 67.03 |
| 1 | 53582419 | rs7556017 | G | A | 2 | 0.6594 | 0.7173 | 1.0436 | 1.0429 | 0.2317 | 30.09 |
| 1 | 53587612 | rs3820202 | G | T | 2 | 0.6595 | 0.6506 | 1.0082 | 1.0097 | 0.2559 | 22.52 |
| 1 | 53540071 | rs7516625 | T | G | 2 | 0.6597 | 0.8057 | 1.0188 | 1.0153 | 0.1463 | 52.61 |
| 1 | 54540084 | rs1118709 | C | T | 2 | 0.6606 | 0.6606 | 1.0305 | 1.0305 | 0.5458 | 0 |
| 1 | 53482078 | rs1710757 | G | A | 2 | 0.6609 | 0.6991 | 1.0137 | 1.0232 | 0.0613 | 71.45 |
| 1 | 53491502 | rs1710763 | T | C | 2 | 0.6609 | 0.7015 | 1.0136 | 1.0229 | 0.061 | 71.5 |
| 1 | 53627373 | rs5599767 | G | A | 2 | 0.661 | 0.661 | 1.0074 | 1.0074 | 0.9992 | 0 |
| 1 | 53553754 | rs1288401 | T | C | 2 | 0.6612 | 0.6559 | 0.9933 | 0.9925 | 0.2767 | 15.49 |
| 1 | 54298702 | rs3495312 | G | A | 2 | 0.6612 | 0.6612 | 1.0079 | 1.0079 | 0.7595 | 0 |
| 1 | 54629734 | rs6177516 | C | T | 2 | 0.6612 | 0.6875 | 0.9886 | 0.9829 | 0.1092 | 61.02 |
| 1 | 54252278 | rs1181182 | G | A | 2 | 0.6615 | 0.6615 | 1.0074 | 1.0074 | 0.8831 | 0 |
| 1 | 54491847 | rs7879851 | C | T | 2 | 0.6616 | 0.6616 | 0.9865 | 0.9865 | 0.7989 | 0 |
| 1 | 54005538 | rs4927006 | T | A | 2 | 0.6618 | 0.9295 | 1.0061 | 1.002 | 0.1249 | 57.54 |
| 1 | 53802696 | rs7632466 | T | C | 2 | 0.662 | 0.662 | 1.0282 | 1.0282 | 0.4963 | 0 |
| 1 | 54320383 | rs1181094 | G | T | 2 | 0.6622 | 0.6622 | 1.0061 | 1.0061 | 0.4047 | 0 |
| 1 | 54312425 | rs7266234 | G | C | 2 | 0.6626 | 0.6791 | 1.0069 | 1.0128 | 0.0584 | 72.08 |
| 1 | 53997760 | rs6134792 | G | A | 2 | 0.6627 | 0.6627 | 1.0324 | 1.0324 | 0.4851 | 0 |
| 1 | 53517283 | rs3820200 | C | G | 2 | 0.6632 | 0.715 | 1.0127 | 1.0232 | 0.0342 | 77.71 |
| 1 | 54254093 | rs7544308 | T | C | 2 | 0.6636 | 0.8537 | 0.9615 | 0.9716 | 0.084 | 66.5 |
| 1 | 54634135 | rs1209062 | G | A | 2 | 0.6637 | 0.5767 | 0.9777 | 0.9505 | 0.1251 | 57.49 |
| 1 | 53568101 | rs1120609 | G | A | 2 | 0.6646 | 0.6415 | 1.0073 | 1.0147 | 0.0794 | 67.51 |
| 1 | 54100837 | rs1450705 | A | G | 2 | 0.6647 | 0.6647 | 1.0145 | 1.0145 | 0.83 | 0 |
| 1 | 53829404 | rs1157099 | C | G | 2 | 0.6649 | 0.6649 | 1.0167 | 1.0167 | 0.9141 | 0 |
| 1 | 53513437 | rs1288362 | T | C | 2 | 0.6652 | 0.6652 | 1.006 | 1.006 | 0.9869 | 0 |
| 1 | 53796901 | rs2225725 | G | A | 2 | 0.6652 | 0.6652 | 1.0313 | 1.0313 | 0.5718 | 0 |
| 1 | 54501432 | rs1121421 | C | T | 2 | 0.6661 | 0.6661 | 1.0299 | 1.0299 | 0.4543 | 0 |
| 1 | 53829729 | rs1999897 | T | C | 2 | 0.6664 | 0.6664 | 0.994 | 0.994 | 0.8591 | 0 |
| 1 | 53490887 | rs8016253 | G | C | 2 | 0.6668 | 0.7022 | 1.0134 | 1.0228 | 0.0627 | 71.14 |

| | | | | | | | | | | | |
|---|---|---|---|---|---|---|---|---|---|---|---|
| 1 | 53677563 | rs6692897 | A | G | 2 | 0.6678 | 0.6678 | 1.0058 | 1.0058 | 0.585 | 0 |
| 1 | 53935740 | rs6671214 | C | T | 2 | 0.668 | 0.6837 | 0.9932 | 0.9911 | 0.1729 | 46.16 |
| 1 | 54532322 | rs1710969 | T | C | 2 | 0.6682 | 0.6682 | 0.9833 | 0.9833 | 0.3862 | 0 |
| 1 | 53533513 | rs1256681 | C | A | 2 | 0.6684 | 0.7913 | 0.9875 | 0.9904 | 0.2185 | 33.96 |
| 1 | 53763728 | rs1288494 | C | G | 2 | 0.6688 | 0.6688 | 1.0073 | 1.0073 | 0.6044 | 0 |
| 1 | 53773594 | rs7289537 | T | C | 2 | 0.6688 | 0.7822 | 1.0111 | 0.9838 | 0.1461 | 52.67 |
| 1 | 53846598 | rs1158721 | G | A | 2 | 0.6692 | 0.6692 | 0.9935 | 0.9935 | 0.8598 | 0 |
| 1 | 53603071 | rs7821413 | C | T | 2 | 0.6695 | 0.6695 | 1.0122 | 1.0122 | 0.4813 | 0 |
| 1 | 54517324 | rs1048944 | A | G | 2 | 0.6717 | 0.6717 | 1.0274 | 1.0274 | 0.7805 | 0 |
| 1 | 53536361 | rs1710783 | C | T | 2 | 0.6719 | 0.6719 | 1.0142 | 1.0142 | 0.5653 | 0 |
| 1 | 53788874 | rs1288523 | G | A | 2 | 0.6719 | 0.6719 | 1.006 | 1.006 | 0.6875 | 0 |
| 1 | 53550187 | rs899974 | C | T | 2 | 0.6722 | 0.6722 | 1.0109 | 1.0109 | 0.8363 | 0 |
| 1 | 54379193 | rs7878859 | G | A | 2 | 0.6723 | 0.6723 | 1.0214 | 1.0214 | 0.5029 | 0 |
| 1 | 53840769 | rs1398847 | T | C | 2 | 0.6729 | 0.6729 | 0.9656 | 0.9656 | 0.8459 | 0 |
| 1 | 53545744 | rs1288392 | A | G | 2 | 0.6732 | 0.6732 | 1.0113 | 1.0113 | 0.9884 | 0 |
| 1 | 54589045 | rs4295836 | G | A | 2 | 0.6732 | 0.6732 | 1.0165 | 1.0165 | 0.9184 | 0 |
| 1 | 54482915 | rs1180483 | G | A | 2 | 0.6734 | 0.6734 | 0.9943 | 0.9943 | 0.5879 | 0 |
| 1 | 53831211 | rs1933535 | A | T | 2 | 0.674 | 0.674 | 0.9941 | 0.9941 | 0.8474 | 0 |
| 1 | 54519295 | rs4129478 | C | T | 2 | 0.6744 | 0.6744 | 0.9829 | 0.9829 | 0.813 | 0 |
| 1 | 54562444 | rs7524477 | A | G | 2 | 0.6747 | 0.6747 | 1.014 | 1.014 | 0.8867 | 0 |
| 1 | 53843539 | rs5734018 | C | G | 2 | 0.6748 | 0.6748 | 0.9935 | 0.9935 | 0.6987 | 0 |
| 1 | 54029893 | rs1207664 | G | A | 2 | 0.6753 | 0.6421 | 1.0265 | 1.0423 | 0.1747 | 45.72 |
| 1 | 53534357 | rs1710781 | T | C | 2 | 0.6755 | 0.6855 | 1.0131 | 1.0207 | 0.1118 | 60.46 |
| 1 | 54676110 | rs3448453 | C | T | 2 | 0.6756 | 0.6756 | 1.014 | 1.014 | 0.8685 | 0 |
| 1 | 53629901 | rs6672276 | C | A | 2 | 0.6757 | 0.6757 | 1.0058 | 1.0058 | 0.7891 | 0 |
| 1 | 53493820 | rs7733057 | G | A | 2 | 0.676 | 0.676 | 1.0298 | 1.0298 | 0.9335 | 0 |
| 1 | 53877512 | rs7570905 | C | T | 2 | 0.676 | 0.6739 | 0.9927 | 0.9922 | 0.2929 | 9.59 |
| 1 | 53607560 | rs3505817 | C | G | 2 | 0.6761 | 0.6761 | 0.9927 | 0.9927 | 0.9414 | 0 |
| 1 | 53755117 | rs1330653 | G | A | 2 | 0.6763 | 0.5779 | 1.0105 | 0.9474 | 0.02 | 81.52 |
| 1 | 53660993 | rs7554022 | A | C | 2 | 0.6767 | 0.6767 | 1.0056 | 1.0056 | 0.358 | 0 |
| 1 | 54234215 | rs3753420 | T | C | 2 | 0.6769 | 0.6769 | 1.0071 | 1.0071 | 0.7922 | 0 |
| 1 | 53839995 | rs7267512 | G | A | 2 | 0.6783 | 0.6783 | 0.9901 | 0.9901 | 0.6573 | 0 |
| 1 | 53578174 | rs3766779 | C | T | 2 | 0.6784 | 0.6784 | 1.0367 | 1.0367 | 0.6234 | 0 |
| 1 | 53830487 | rs1088878 | T | C | 2 | 0.6794 | 0.6794 | 0.9942 | 0.9942 | 0.8712 | 0 |
| 1 | 53796857 | rs7676654 | G | C | 2 | 0.6806 | 0.6806 | 1.0298 | 1.0298 | 0.5884 | 0 |
| 1 | 54485779 | rs6767600 | G | A | 2 | 0.6812 | 0.6812 | 0.9945 | 0.9945 | 0.4664 | 0 |
| 1 | 53585568 | rs1158404 | G | C | 2 | 0.6815 | 0.6815 | 1.0482 | 1.0482 | 0.7166 | 0 |
| 1 | 54682493 | rs3400178 | C | G | 2 | 0.6815 | 0.6815 | 0.9645 | 0.9645 | 0.4845 | 0 |
| 1 | 53652273 | rs5956104 | A | T | 2 | 0.6823 | 0.6823 | 1.0168 | 1.0168 | 0.8158 | 0 |
| 1 | 53829511 | rs4926981 | C | G | 2 | 0.6824 | 0.6824 | 0.9943 | 0.9943 | 0.8596 | 0 |
| 1 | 54193922 | rs1120621 | G | C | 2 | 0.6831 | 0.6831 | 0.9692 | 0.9692 | 0.9434 | 0 |
| 1 | 53560301 | rs3766767 | T | C | 2 | 0.6838 | 0.6838 | 0.9935 | 0.9935 | 0.4182 | 0 |
| 1 | 54548798 | rs1120628 | T | C | 2 | 0.6838 | 0.6838 | 1.0283 | 1.0283 | 0.6895 | 0 |
| 1 | 53543112 | rs946590 | C | G | 2 | 0.684 | 0.684 | 1.0108 | 1.0108 | 0.9983 | 0 |
| 1 | 53494825 | rs1710765 | A | G | 2 | 0.6845 | 0.7131 | 1.0126 | 1.0214 | 0.0677 | 70.04 |
| 1 | 53706829 | rs6677126 | C | T | 2 | 0.6846 | 0.6846 | 1.006 | 1.006 | 0.3877 | 0 |
| 1 | 53970536 | rs1205728 | C | T | 2 | 0.6847 | 0.7808 | 1.0078 | 1.0062 | 0.2556 | 22.63 |
| 1 | 53500694 | rs7290317 | A | G | 2 | 0.6857 | 0.6857 | 0.9857 | 0.9857 | 0.8887 | 0 |

| | | | | | | | | | | | |
|---|---|---|---|---|---|---|---|---|---|---|---|
| 1 | 53845985 | rs1710839 | G | A | 2 | 0.6858 | 0.6858 | 0.9939 | 0.9939 | 0.8561 | 0 |
| 1 | 53566088 | rs1273629 | C | T | 2 | 0.6863 | 0.6863 | 0.9921 | 0.9921 | 0.7492 | 0 |
| 1 | 53572058 | rs1288424 | G | C | 2 | 0.6866 | 0.6866 | 0.9917 | 0.9917 | 0.7299 | 0 |
| 1 | 53534143 | rs5665738 | C | G | 2 | 0.6871 | 0.6919 | 1.0126 | 1.0201 | 0.1153 | 59.68 |
| 1 | 53993395 | rs495835 | G | A | 2 | 0.6872 | 0.751 | 0.9853 | 0.9792 | 0.0739 | 68.69 |
| 1 | 54326490 | rs1181097 | T | A | 2 | 0.6873 | 0.6873 | 1.0056 | 1.0056 | 0.4631 | 0 |
| 1 | 54360467 | rs1158811 | C | G | 2 | 0.6875 | 0.6875 | 1.0056 | 1.0056 | 0.4974 | 0 |
| 1 | 53534248 | rs1710781 | A | G | 2 | 0.6876 | 0.6923 | 1.0126 | 1.0201 | 0.1146 | 59.83 |
| 1 | 53658039 | rs5739040 | A | G | 2 | 0.6879 | 0.6879 | 1.0083 | 1.0083 | 0.4856 | 0 |
| 1 | 53873140 | rs6176838 | G | A | 2 | 0.688 | 0.688 | 1.0185 | 1.0185 | 0.5197 | 0 |
| 1 | 53555071 | rs1288402 | G | A | 2 | 0.6881 | 0.6714 | 0.9939 | 0.9918 | 0.2214 | 33.11 |
| 1 | 53500244 | rs7434419 | A | G | 2 | 0.6887 | 0.7171 | 1.0124 | 1.0214 | 0.0639 | 70.87 |
| 1 | 53631902 | rs874349 | C | T | 2 | 0.6887 | 0.6887 | 1.0064 | 1.0064 | 0.8408 | 0 |
| 1 | 54698610 | rs6676841 | C | A | 2 | 0.6887 | 0.6887 | 0.9656 | 0.9656 | 0.3922 | 0 |
| 1 | 54572243 | rs1078897 | C | A | 2 | 0.6893 | 0.6893 | 0.9946 | 0.9946 | 0.5979 | 0 |
| 1 | 53816317 | rs5853983 | A | T | 2 | 0.69 | 0.69 | 1.0237 | 1.0237 | 0.687 | 0 |
| 1 | 53877480 | rs1737980 | C | T | 2 | 0.6901 | 0.6885 | 0.993 | 0.9928 | 0.3048 | 5.05 |
| 1 | 53500513 | rs6100536 | C | G | 2 | 0.6903 | 0.7175 | 1.0124 | 1.0213 | 0.064 | 70.85 |
| 1 | 53477558 | rs7498477 | T | G | 2 | 0.691 | 0.691 | 1.0142 | 1.0142 | 0.8181 | 0 |
| 1 | 54219484 | rs1181160 | A | G | 2 | 0.6913 | 0.6913 | 1.0058 | 1.0058 | 0.4707 | 0 |
| 1 | 54578658 | rs1120628 | T | C | 2 | 0.6927 | 0.6927 | 1.0129 | 1.0129 | 0.8476 | 0 |
| 1 | 53586380 | rs936642 | C | A | 2 | 0.6939 | 0.6929 | 1.0073 | 1.0074 | 0.3138 | 1.44 |
| 1 | 53831342 | rs1933536 | C | A | 2 | 0.6941 | 0.6941 | 0.9945 | 0.9945 | 0.9113 | 0 |
| 1 | 53661966 | rs1088877 | G | A | 2 | 0.6942 | 0.7018 | 1.0053 | 1.0052 | 0.3116 | 2.33 |
| 1 | 53849975 | rs1150539 | C | T | 2 | 0.6944 | 0.6944 | 0.9914 | 0.9914 | 0.7903 | 0 |
| 1 | 53665151 | rs2062015 | C | G | 2 | 0.6945 | 0.7196 | 1.0053 | 1.0051 | 0.2986 | 7.45 |
| 1 | 53505568 | rs5694357 | G | A | 2 | 0.6947 | 0.72 | 1.0122 | 1.0211 | 0.0642 | 70.81 |
| 1 | 53741816 | rs945183 | A | G | 2 | 0.6957 | 0.6957 | 1.0058 | 1.0058 | 0.3415 | 0 |
| 1 | 54263989 | rs1181181 | T | C | 2 | 0.6959 | 0.6959 | 1.0066 | 1.0066 | 0.8127 | 0 |
| 1 | 54482523 | rs1537323 | T | G | 2 | 0.6962 | 0.6962 | 0.9948 | 0.9948 | 0.5918 | 0 |
| 1 | 53812473 | rs1115362 | G | A | 2 | 0.6963 | 0.6963 | 0.9839 | 0.9839 | 0.6928 | 0 |
| 1 | 54627748 | rs581542 | A | G | 2 | 0.6965 | 0.6965 | 1.0056 | 1.0056 | 0.6915 | 0 |
| 1 | 53797939 | rs7289742 | C | T | 2 | 0.6968 | 0.6968 | 1.0282 | 1.0282 | 0.6008 | 0 |
| 1 | 53695628 | rs1859251 | A | G | 2 | 0.6971 | 0.6971 | 0.9665 | 0.9665 | 0.5544 | 0 |
| 1 | 54661262 | rs1154419 | C | T | 2 | 0.6984 | 0.6984 | 1.0154 | 1.0154 | 0.4449 | 0 |
| 1 | 53505072 | rs6135553 | A | T | 2 | 0.6987 | 0.7223 | 1.012 | 1.021 | 0.0636 | 70.95 |
| 1 | 53506670 | rs1710772 | A | C | 2 | 0.6988 | 0.7223 | 1.012 | 1.021 | 0.0636 | 70.94 |
| 1 | 53562770 | rs1288413 | C | T | 2 | 0.6995 | 0.6841 | 0.9941 | 0.9926 | 0.2449 | 26.04 |
| 1 | 53496507 | rs1503116 | G | A | 2 | 0.6996 | 0.7225 | 1.0119 | 1.0204 | 0.07 | 69.55 |
| 1 | 53798745 | rs6017524 | T | C | 2 | 0.6999 | 0.6999 | 1.0279 | 1.0279 | 0.6002 | 0 |
| 1 | 53970656 | rs4926999 | C | T | 2 | 0.7 | 0.6966 | 1.0062 | 1.0066 | 0.2991 | 7.26 |
| 1 | 53776616 | rs1288480 | C | T | 2 | 0.7003 | 0.7003 | 1.0067 | 1.0067 | 0.9959 | 0 |
| 1 | 54522307 | rs3820112 | A | G | 2 | 0.7007 | 0.7007 | 1.0115 | 1.0115 | 0.4921 | 0 |
| 1 | 53629585 | rs6695567 | G | A | 2 | 0.7009 | 0.7009 | 1.0052 | 1.0052 | 0.7101 | 0 |
| 1 | 53605727 | rs7757722 | A | G | 2 | 0.701 | 0.701 | 1.0107 | 1.0107 | 0.5977 | 0 |
| 1 | 53654542 | rs3014991 | A | G | 2 | 0.7019 | 0.6543 | 0.968 | 0.9351 | 0.0973 | 63.62 |
| 1 | 54362947 | rs1088881 | A | G | 2 | 0.7019 | 0.7019 | 1.0054 | 1.0054 | 0.5153 | 0 |
| 1 | 53970728 | rs1120616 | C | T | 2 | 0.7022 | 0.6982 | 1.0062 | 1.0066 | 0.2959 | 8.49 |

| | | | | | | | | | | | |
|---|---|---|---|---|---|---|---|---|---|---|---|
| 1 | 54360647 | rs5713260 | C | A | 2 | 0.7023 | 0.7023 | 0.993 | 0.993 | 0.5402 | 0 |
| 1 | 54124833 | rs1078896 | G | C | 2 | 0.7028 | 0.7028 | 0.9876 | 0.9876 | 0.8985 | 0 |
| 1 | 54469989 | rs1507769 | G | A | 2 | 0.7028 | 0.7028 | 0.9829 | 0.9829 | 0.3216 | 0 |
| 1 | 53502018 | rs1710770 | A | C | 2 | 0.703 | 0.703 | 0.9863 | 0.9863 | 0.8653 | 0 |
| 1 | 53817135 | rs6658926 | T | C | 2 | 0.7034 | 0.7034 | 0.9947 | 0.9947 | 0.9007 | 0 |
| 1 | 53817220 | rs6588474 | T | C | 2 | 0.7034 | 0.7034 | 0.9946 | 0.9946 | 0.473 | 0 |
| 1 | 53756870 | rs1288503 | C | T | 2 | 0.7035 | 0.6735 | 0.9935 | 0.9905 | 0.294 | 9.19 |
| 1 | 53798397 | rs7515071 | A | C | 2 | 0.7036 | 0.7036 | 1.0275 | 1.0275 | 0.6082 | 0 |
| 1 | 53798423 | rs7289743 | T | C | 2 | 0.7037 | 0.7037 | 1.0275 | 1.0275 | 0.6077 | 0 |
| 1 | 53617840 | rs1203310 | T | C | 2 | 0.7038 | 0.7038 | 1.0063 | 1.0063 | 0.417 | 0 |
| 1 | 53592335 | rs7446385 | G | A | 2 | 0.7039 | 0.7039 | 0.9537 | 0.9537 | 0.8166 | 0 |
| 1 | 53607227 | rs1207190 | A | C | 2 | 0.704 | 0.704 | 1.0054 | 1.0054 | 0.7972 | 0 |
| 1 | 54476323 | rs1203653 | T | C | 2 | 0.704 | 0.704 | 1.0051 | 1.0051 | 0.9828 | 0 |
| 1 | 53798162 | rs8027619 | T | C | 2 | 0.7042 | 0.7042 | 1.0275 | 1.0275 | 0.6015 | 0 |
| 1 | 53944430 | rs1120616 | C | A | 2 | 0.7042 | 0.7042 | 1.0068 | 1.0068 | 0.6429 | 0 |
| 1 | 53826846 | rs1211862 | C | T | 2 | 0.7051 | 0.7051 | 0.9936 | 0.9936 | 0.6079 | 0 |
| 1 | 54579153 | rs9919142 | A | G | 2 | 0.7051 | 0.7051 | 1.0127 | 1.0127 | 0.8651 | 0 |
| 1 | 53816524 | rs1214327 | C | G | 2 | 0.7056 | 0.7056 | 0.9946 | 0.9946 | 0.5402 | 0 |
| 1 | 53605035 | rs7867959 | C | T | 2 | 0.7057 | 0.7789 | 1.016 | 1.0132 | 0.2785 | 14.86 |
| 1 | 53654573 | rs2929074 | A | C | 2 | 0.706 | 0.6312 | 0.9706 | 0.9372 | 0.1176 | 59.16 |
| 1 | 54476967 | rs4136684 | C | G | 2 | 0.7065 | 0.7065 | 1.0051 | 1.0051 | 0.9307 | 0 |
| 1 | 53798164 | rs7986581 | C | A | 2 | 0.707 | 0.707 | 1.0272 | 1.0272 | 0.5993 | 0 |
| 1 | 53600988 | rs1212617 | A | G | 2 | 0.7074 | 0.7074 | 1.0058 | 1.0058 | 0.6853 | 0 |
| 1 | 54525807 | rs7533726 | G | A | 2 | 0.7075 | 0.7075 | 1.032 | 1.032 | 0.6655 | 0 |
| 1 | 53763106 | rs2297826 | A | G | 2 | 0.7081 | 0.7081 | 0.9939 | 0.9939 | 0.3444 | 0 |
| 1 | 53709447 | rs1778538 | T | A | 2 | 0.7092 | 0.7092 | 0.9794 | 0.9794 | 0.4763 | 0 |
| 1 | 54679518 | rs1180952 | C | T | 2 | 0.7093 | 0.9781 | 1.0071 | 0.999 | 0.0725 | 68.99 |
| 1 | 53501596 | rs1256629 | G | C | 2 | 0.7095 | 0.7246 | 1.0116 | 1.0207 | 0.066 | 70.42 |
| 1 | 54482306 | rs1537322 | G | T | 2 | 0.71 | 0.71 | 0.995 | 0.995 | 0.6024 | 0 |
| 1 | 53537396 | rs3410752 | G | A | 2 | 0.7104 | 0.7104 | 0.9882 | 0.9882 | 0.4046 | 0 |
| 1 | 53729536 | rs6588473 | T | C | 2 | 0.7105 | 0.7275 | 1.0055 | 1.0053 | 0.306 | 4.58 |
| 1 | 53993663 | rs493163 | G | A | 2 | 0.7105 | 0.7633 | 0.9865 | 0.9804 | 0.0764 | 68.16 |
| 1 | 54497040 | rs1547467 | T | C | 2 | 0.7106 | 0.7106 | 0.995 | 0.995 | 0.4936 | 0 |
| 1 | 54505096 | rs1209788 | T | A | 2 | 0.7106 | 0.7106 | 0.9875 | 0.9875 | 0.7391 | 0 |
| 1 | 53628658 | rs3535266 | G | A | 2 | 0.711 | 0.711 | 1.0058 | 1.0058 | 0.8678 | 0 |
| 1 | 53843997 | rs7289948 | G | A | 2 | 0.711 | 0.711 | 0.9941 | 0.9941 | 0.7907 | 0 |
| 1 | 53510237 | rs7567485 | G | A | 2 | 0.7116 | 0.7294 | 1.0115 | 1.0205 | 0.0623 | 71.23 |
| 1 | 54626529 | rs646914 | G | C | 2 | 0.7116 | 0.7116 | 1.005 | 1.005 | 0.6124 | 0 |
| 1 | 54484682 | rs7540270 | A | G | 2 | 0.7117 | 0.7117 | 0.995 | 0.995 | 0.4694 | 0 |
| 1 | 54366741 | rs7527992 | G | A | 2 | 0.712 | 0.712 | 0.9931 | 0.9931 | 0.725 | 0 |
| 1 | 53586053 | rs6666283 | G | C | 2 | 0.7121 | 0.9949 | 1.0066 | 1.0002 | 0.1409 | 53.88 |
| 1 | 54206980 | rs1145731 | T | C | 2 | 0.7128 | 0.7128 | 0.9849 | 0.9849 | 0.7604 | 0 |
| 1 | 53630705 | rs6588469 | C | T | 2 | 0.7131 | 0.7131 | 1.0051 | 1.0051 | 0.7369 | 0 |
| 1 | 53970693 | rs7290694 | C | T | 2 | 0.7131 | 0.8228 | 0.9868 | 0.9859 | 0.0796 | 67.46 |
| 1 | 54513323 | rs1206578 | A | G | 2 | 0.7132 | 0.7132 | 0.9875 | 0.9875 | 0.9153 | 0 |
| 1 | 53782812 | rs7289538 | G | A | 2 | 0.7138 | 0.7138 | 1.023 | 1.023 | 0.7215 | 0 |
| 1 | 53537699 | rs1710783 | A | G | 2 | 0.7142 | 0.7142 | 1.0123 | 1.0123 | 0.5552 | 0 |
| 1 | 54500487 | rs7539402 | G | A | 2 | 0.7146 | 0.7146 | 0.9876 | 0.9876 | 0.8476 | 0 |

| | | | | | | | | | | | |
|---|---|---|---|---|---|---|---|---|---|---|---|
| 1 | 53630648 | rs6588468 | A | G | 2 | 0.7155 | 0.7155 | 1.005 | 1.005 | 0.7272 | 0 |
| 1 | 54208890 | rs2316320 | C | T | 2 | 0.7163 | 0.7163 | 1.0056 | 1.0056 | 0.5889 | 0 |
| 1 | 53684525 | rs436887 | C | G | 2 | 0.7179 | 0.7179 | 1.0055 | 1.0055 | 0.4397 | 0 |
| 1 | 54592941 | rs1273945 | C | T | 2 | 0.7181 | 0.7181 | 1.0173 | 1.0173 | 0.6078 | 0 |
| 1 | 54676025 | rs3737833 | C | G | 2 | 0.7182 | 0.9722 | 1.0069 | 0.9987 | 0.0725 | 68.99 |
| 1 | 53604780 | rs5992551 | G | T | 2 | 0.7183 | 0.7183 | 1.01 | 1.01 | 0.5921 | 0 |
| 1 | 53568878 | rs1120610 | T | A | 2 | 0.7184 | 0.6661 | 1.0063 | 1.0134 | 0.0959 | 63.93 |
| 1 | 53530023 | rs1710779 | G | A | 2 | 0.7188 | 0.7188 | 1.0131 | 1.0131 | 0.9281 | 0 |
| 1 | 54240170 | rs1181196 | T | C | 2 | 0.7191 | 0.7191 | 1.0061 | 1.0061 | 0.9181 | 0 |
| 1 | 53672380 | rs7871767 | C | T | 2 | 0.7193 | 0.7193 | 1.0076 | 1.0076 | 0.5611 | 0 |
| 1 | 53815565 | rs4926977 | G | T | 2 | 0.7194 | 0.7194 | 0.9949 | 0.9949 | 0.4868 | 0 |
| 1 | 53665612 | rs2046138 | G | A | 2 | 0.7202 | 0.7202 | 1.0049 | 1.0049 | 0.4448 | 0 |
| 1 | 53544597 | rs1288389 | T | C | 2 | 0.7205 | 0.7205 | 0.993 | 0.993 | 0.8212 | 0 |
| 1 | 53608638 | rs7549922 | A | C | 2 | 0.7209 | 0.7209 | 1.005 | 1.005 | 0.9345 | 0 |
| 1 | 53565128 | rs1629126 | C | T | 2 | 0.7212 | 0.7212 | 0.9949 | 0.9949 | 0.3663 | 0 |
| 1 | 53666965 | rs3766759 | T | G | 2 | 0.7214 | 0.7214 | 1.0048 | 1.0048 | 0.4456 | 0 |
| 1 | 54613731 | rs3915626 | C | G | 2 | 0.7214 | 0.7214 | 1.0056 | 1.0056 | 0.5364 | 0 |
| 1 | 53777882 | rs2788030 | T | C | 2 | 0.7217 | 0.7217 | 0.98 | 0.98 | 0.3884 | 0 |
| 1 | 53832824 | rs3207053 | C | T | 2 | 0.722 | 0.722 | 0.995 | 0.995 | 0.9324 | 0 |
| 1 | 54487596 | rs3571946 | G | A | 2 | 0.7231 | 0.7231 | 0.9952 | 0.9952 | 0.4038 | 0 |
| 1 | 54215812 | rs6588488 | C | T | 2 | 0.7232 | 0.7232 | 0.9944 | 0.9944 | 0.6009 | 0 |
| 1 | 53678728 | rs3766757 | C | T | 2 | 0.7233 | 0.7233 | 1.0163 | 1.0163 | 0.5315 | 0 |
| 1 | 54457657 | rs1141682 | T | G | 2 | 0.7233 | 0.7233 | 1.0161 | 1.0161 | 0.3863 | 0 |
| 1 | 54620197 | rs1407081 | G | A | 2 | 0.7234 | 0.8131 | 1.0175 | 1.0141 | 0.2325 | 29.85 |
| 1 | 54130244 | rs7727941 | T | A | 2 | 0.7236 | 0.7236 | 1.016 | 1.016 | 0.7587 | 0 |
| 1 | 54318725 | rs6749734 | G | C | 2 | 0.724 | 0.724 | 1.0048 | 1.0048 | 0.3275 | 0 |
| 1 | 53536335 | rs5961175 | A | G | 2 | 0.7243 | 0.7115 | 1.0111 | 1.018 | 0.1318 | 55.97 |
| 1 | 53567419 | rs1088876 | T | C | 2 | 0.7255 | 0.7255 | 1.0094 | 1.0094 | 0.6672 | 0 |
| 1 | 54318411 | rs7519869 | G | A | 2 | 0.7261 | 0.7261 | 1.0049 | 1.0049 | 0.4802 | 0 |
| 1 | 53809442 | rs7467979 | A | G | 2 | 0.7266 | 0.7266 | 1.0236 | 1.0236 | 0.5171 | 0 |
| 1 | 54318765 | rs1936835 | T | C | 2 | 0.7274 | 0.7274 | 1.0048 | 1.0048 | 0.3257 | 0 |
| 1 | 54581781 | rs7505815 | A | T | 2 | 0.7274 | 0.7274 | 1.0298 | 1.0298 | 0.8013 | 0 |
| 1 | 54487125 | rs6696554 | A | G | 2 | 0.7275 | 0.7275 | 0.9953 | 0.9953 | 0.4016 | 0 |
| 1 | 54630044 | rs1137806 | T | C | 2 | 0.728 | 0.7237 | 0.9909 | 0.9868 | 0.1585 | 49.71 |
| 1 | 53808916 | rs1710835 | C | A | 2 | 0.7284 | 0.7284 | 1.0234 | 1.0234 | 0.5272 | 0 |
| 1 | 53548606 | rs1180439 | A | G | 2 | 0.7295 | 0.6843 | 1.0067 | 1.011 | 0.1845 | 43.22 |
| 1 | 53682678 | rs1710814 | G | A | 2 | 0.7295 | 0.7295 | 1.0073 | 1.0073 | 0.6214 | 0 |
| 1 | 54687307 | rs4926624 | A | C | 2 | 0.7303 | 0.6836 | 1.0062 | 1.011 | 0.1523 | 51.2 |
| 1 | 54581566 | rs7946908 | G | T | 2 | 0.7304 | 0.7304 | 1.0295 | 1.0295 | 0.8037 | 0 |
| 1 | 53628911 | rs3603824 | T | C | 2 | 0.7305 | 0.7305 | 1.0054 | 1.0054 | 0.7787 | 0 |
| 1 | 54601281 | rs2026111 | G | A | 2 | 0.7305 | 0.7305 | 1.0053 | 1.0053 | 0.3275 | 0 |
| 1 | 53530368 | rs1710779 | C | T | 2 | 0.7307 | 0.7396 | 0.9666 | 0.9671 | 0.3078 | 3.87 |
| 1 | 53972764 | rs1120616 | C | T | 2 | 0.7307 | 0.9908 | 1.0063 | 0.9996 | 0.1034 | 62.31 |
| 1 | 53629190 | rs6692772 | G | A | 2 | 0.7316 | 0.7316 | 1.0046 | 1.0046 | 0.7321 | 0 |
| 1 | 53521242 | rs3599460 | G | A | 2 | 0.7318 | 0.7318 | 1.0113 | 1.0113 | 0.7763 | 0 |
| 1 | 54674983 | rs1088883 | C | G | 2 | 0.7323 | 0.7323 | 1.0082 | 1.0082 | 0.71 | 0 |
| 1 | 53596148 | rs1299859 | T | G | 2 | 0.7326 | 0.714 | 0.9905 | 0.9887 | 0.2735 | 16.59 |
| 1 | 53492087 | rs5960475 | A | G | 2 | 0.7332 | 0.7332 | 1.0111 | 1.0111 | 0.8754 | 0 |

| | | | | | | | | | | | |
|---|---|---|---|---|---|---|---|---|---|---|---|
| 1 | 53822103 | rs1274946 | G | A | 2 | 0.734 | 0.734 | 0.9943 | 0.9943 | 0.5942 | 0 |
| 1 | 54574610 | rs1204451 | G | A | 2 | 0.734 | 0.734 | 0.9954 | 0.9954 | 0.5276 | 0 |
| 1 | 53875848 | rs1125627 | C | T | 2 | 0.7343 | 0.7324 | 0.9941 | 0.9939 | 0.3097 | 3.09 |
| 1 | 53517863 | rs7989163 | C | A | 2 | 0.7349 | 0.7349 | 1.0124 | 1.0124 | 0.9258 | 0 |
| 1 | 53536452 | rs1710783 | A | G | 2 | 0.7352 | 0.7172 | 1.0106 | 1.0177 | 0.1293 | 56.54 |
| 1 | 53490068 | rs4926944 | C | T | 2 | 0.7359 | 0.7359 | 1.011 | 1.011 | 0.8744 | 0 |
| 1 | 54005331 | rs1143886 | G | A | 2 | 0.7361 | 0.7361 | 1.0143 | 1.0143 | 0.3885 | 0 |
| 1 | 54471189 | rs681660 | G | A | 2 | 0.7366 | 0.7894 | 1.0046 | 1.004 | 0.2743 | 16.34 |
| 1 | 54632364 | rs1209794 | A | G | 2 | 0.7372 | 0.7372 | 0.9885 | 0.9885 | 0.7369 | 0 |
| 1 | 53817480 | rs6695236 | C | T | 2 | 0.7381 | 0.7381 | 0.9954 | 0.9954 | 0.9047 | 0 |
| 1 | 54671859 | rs1074969 | T | C | 2 | 0.7381 | 0.7381 | 1.0083 | 1.0083 | 0.7516 | 0 |
| 1 | 54175540 | rs1710919 | G | C | 2 | 0.7382 | 0.7382 | 0.9748 | 0.9748 | 0.8562 | 0 |
| 1 | 54599517 | rs3425284 | T | G | 2 | 0.7387 | 0.7387 | 1.0124 | 1.0124 | 0.7733 | 0 |
| 1 | 54493060 | rs1180086 | C | T | 2 | 0.739 | 0.739 | 0.9887 | 0.9887 | 0.8063 | 0 |
| 1 | 53765434 | rs1120614 | G | A | 2 | 0.741 | 0.478 | 0.9936 | 1.0824 | 0.0046 | 87.55 |
| 1 | 54183742 | rs7928378 | C | A | 2 | 0.7411 | 0.7411 | 0.9751 | 0.9751 | 0.9709 | 0 |
| 1 | 53629849 | rs6672178 | C | T | 2 | 0.7414 | 0.7414 | 1.0045 | 1.0045 | 0.7277 | 0 |
| 1 | 53604717 | rs5872268 | C | T | 2 | 0.7422 | 0.7422 | 1.0092 | 1.0092 | 0.5715 | 0 |
| 1 | 53568350 | rs3606261 | C | T | 2 | 0.7424 | 0.6732 | 1.0057 | 1.0151 | 0.0557 | 72.68 |
| 1 | 53613779 | rs7585522 | C | T | 2 | 0.7424 | 0.7424 | 1.0054 | 1.0054 | 0.4656 | 0 |
| 1 | 53564586 | rs1288418 | T | C | 2 | 0.7425 | 0.7425 | 1.0089 | 1.0089 | 0.9189 | 0 |
| 1 | 54591294 | rs1458224 | C | T | 2 | 0.743 | 0.743 | 0.9847 | 0.9847 | 0.5338 | 0 |
| 1 | 53712727 | rs5174 | C | T | 2 | 0.7438 | 0.7438 | 1.0046 | 1.0046 | 0.8862 | 0 |
| 1 | 53937254 | rs8179476 | G | T | 2 | 0.7439 | 0.7439 | 0.9857 | 0.9857 | 0.6953 | 0 |
| 1 | 53602874 | rs1149217 | G | A | 2 | 0.7454 | 0.7454 | 1.0091 | 1.0091 | 0.5962 | 0 |
| 1 | 54495254 | rs1208591 | G | C | 2 | 0.7459 | 0.7459 | 0.989 | 0.989 | 0.8209 | 0 |
| 1 | 53502916 | rs7422400 | T | A | 2 | 0.7464 | 0.7706 | 1.0077 | 1.0113 | 0.1065 | 61.62 |
| 1 | 53536315 | rs5992696 | A | G | 2 | 0.7466 | 0.7255 | 1.0101 | 1.0174 | 0.1258 | 57.34 |
| 1 | 53547188 | rs7862438 | G | T | 2 | 0.7466 | 0.7221 | 1.0103 | 1.0135 | 0.2468 | 25.45 |
| 1 | 53808580 | rs7827977 | C | T | 2 | 0.7468 | 0.7468 | 1.0215 | 1.0215 | 0.5907 | 0 |
| 1 | 53605884 | rs1288367 | T | C | 2 | 0.747 | 0.747 | 0.9946 | 0.9946 | 0.3805 | 0 |
| 1 | 53536231 | rs7834413 | A | G | 2 | 0.7474 | 0.7262 | 1.0101 | 1.0173 | 0.1253 | 57.44 |
| 1 | 53499226 | rs1465569 | G | A | 2 | 0.7475 | 0.7343 | 1.0101 | 1.0194 | 0.0761 | 68.22 |
| 1 | 54366364 | rs1213146 | G | A | 2 | 0.7478 | 0.7478 | 0.9941 | 0.9941 | 0.6787 | 0 |
| 1 | 53536236 | rs7809640 | T | C | 2 | 0.7481 | 0.7264 | 1.0101 | 1.0173 | 0.1254 | 57.42 |
| 1 | 53551880 | rs1204267 | C | T | 2 | 0.7491 | 0.7491 | 0.9914 | 0.9914 | 0.9419 | 0 |
| 1 | 53489132 | rs7555050 | G | A | 2 | 0.7493 | 0.7493 | 1.0116 | 1.0116 | 0.861 | 0 |
| 1 | 54062619 | rs7267520 | G | T | 2 | 0.7493 | 0.9171 | 1.0122 | 0.9916 | 0.0438 | 75.39 |
| 1 | 54009583 | rs4136304 | C | T | 2 | 0.7498 | 0.7877 | 0.985 | 0.9863 | 0.2821 | 13.55 |
| 1 | 53605304 | rs7543108 | C | T | 2 | 0.7501 | 0.7501 | 1.0089 | 1.0089 | 0.5957 | 0 |
| 1 | 53602464 | rs1119859 | G | A | 2 | 0.7505 | 0.7505 | 1.0089 | 1.0089 | 0.6048 | 0 |
| 1 | 53536025 | rs2275456 | T | C | 2 | 0.7506 | 0.7279 | 1.01 | 1.0173 | 0.1244 | 57.65 |
| 1 | 54491048 | rs9651201 | T | C | 2 | 0.7515 | 0.7515 | 0.9893 | 0.9893 | 0.815 | 0 |
| 1 | 53545071 | rs1180383 | T | C | 2 | 0.7517 | 0.7008 | 1.0061 | 1.0103 | 0.1875 | 42.43 |
| 1 | 54360879 | rs6662856 | T | C | 2 | 0.752 | 0.752 | 0.9957 | 0.9957 | 0.4689 | 0 |
| 1 | 53514516 | rs3766762 | T | C | 2 | 0.7521 | 0.751 | 1.0098 | 1.0185 | 0.0661 | 70.39 |
| 1 | 53511507 | rs5933535 | G | A | 2 | 0.7532 | 0.748 | 1.0098 | 1.0189 | 0.0657 | 70.49 |
| 1 | 54598275 | rs630538 | C | T | 2 | 0.7532 | 0.7532 | 1.0045 | 1.0045 | 0.8925 | 0 |

| | | | | | | | | | | | |
|---|---|---|---|---|---|---|---|---|---|---|---|
| 1 | 53806223 | rs1206706 | G | A | 2 | 0.7534 | 0.7534 | 0.9695 | 0.9695 | 0.7152 | 0 |
| 1 | 53604599 | rs6694665 | G | A | 2 | 0.754 | 0.754 | 1.0087 | 1.0087 | 0.6374 | 0 |
| 1 | 54484482 | rs914720 | C | T | 2 | 0.7543 | 0.86 | 0.9958 | 0.9971 | 0.224 | 32.37 |
| 1 | 53620833 | rs6689888 | C | T | 2 | 0.7547 | 0.7547 | 1.0052 | 1.0052 | 0.5477 | 0 |
| 1 | 54222667 | rs1426898 | T | C | 2 | 0.7553 | 0.7553 | 0.9848 | 0.9848 | 0.4685 | 0 |
| 1 | 53512354 | rs7544825 | T | G | 2 | 0.7556 | 0.7527 | 1.0096 | 1.0184 | 0.0657 | 70.49 |
| 1 | 54477726 | rs5576612 | T | G | 2 | 0.7558 | 0.7558 | 1.0042 | 1.0042 | 0.9442 | 0 |
| 1 | 53566341 | rs6176811 | C | T | 2 | 0.756 | 0.6843 | 1.0054 | 1.0132 | 0.0763 | 68.17 |
| 1 | 53831095 | rs1120615 | G | T | 2 | 0.756 | 0.756 | 0.9957 | 0.9957 | 0.9062 | 0 |
| 1 | 54127745 | rs1710904 | A | G | 2 | 0.7564 | 0.7398 | 0.9931 | 0.9909 | 0.2284 | 31.08 |
| 1 | 54505437 | rs3766466 | C | A | 2 | 0.7568 | 0.7568 | 0.9895 | 0.9895 | 0.8069 | 0 |
| 1 | 54493420 | rs6699257 | A | G | 2 | 0.7576 | 0.7576 | 0.9896 | 0.9896 | 0.8097 | 0 |
| 1 | 53499893 | rs6673572 | G | A | 2 | 0.7577 | 0.7773 | 1.0073 | 1.0105 | 0.1195 | 58.73 |
| 1 | 53918872 | rs4142165 | T | C | 2 | 0.7581 | 0.8079 | 0.9882 | 0.9888 | 0.2301 | 30.58 |
| 1 | 53577650 | rs1737713 | G | A | 2 | 0.7585 | 0.7585 | 1.0079 | 1.0079 | 0.6132 | 0 |
| 1 | 53605608 | rs7724666 | C | T | 2 | 0.7589 | 0.7589 | 1.0085 | 1.0085 | 0.6019 | 0 |
| 1 | 53810749 | rs7940777 | G | T | 2 | 0.7589 | 0.7589 | 1.0116 | 1.0116 | 0.9135 | 0 |
| 1 | 54542989 | rs1158329 | A | C | 2 | 0.759 | 0.759 | 0.9859 | 0.9859 | 0.8236 | 0 |
| 1 | 53805413 | rs7289745 | C | T | 2 | 0.7594 | 0.7594 | 0.9703 | 0.9703 | 0.6827 | 0 |
| 1 | 53510671 | rs1159087 | A | G | 2 | 0.7607 | 0.7607 | 1.0093 | 1.0093 | 0.6285 | 0 |
| 1 | 53759072 | rs958825 | G | A | 2 | 0.7616 | 0.5362 | 0.995 | 1.0461 | 0.0075 | 86.02 |
| 1 | 54500185 | rs7528837 | A | G | 2 | 0.7616 | 0.7616 | 0.9897 | 0.9897 | 0.8052 | 0 |
| 1 | 53551293 | rs1288397 | A | G | 2 | 0.7619 | 0.7619 | 1.0045 | 1.0045 | 0.6748 | 0 |
| 1 | 54476931 | rs7266411 | A | G | 2 | 0.7623 | 0.7623 | 0.9958 | 0.9958 | 0.4882 | 0 |
| 1 | 53568783 | rs1209744 | C | T | 2 | 0.7626 | 0.6864 | 1.0052 | 1.0145 | 0.0519 | 73.53 |
| 1 | 54615599 | rs1158778 | A | C | 2 | 0.7628 | 0.7479 | 1.0104 | 1.0125 | 0.2635 | 20.03 |
| 1 | 53822114 | rs1240551 | G | C | 2 | 0.763 | 0.763 | 0.995 | 0.995 | 0.5641 | 0 |
| 1 | 53535918 | rs2275454 | A | C | 2 | 0.7636 | 0.7473 | 1.0096 | 1.0165 | 0.1177 | 59.14 |
| 1 | 53825310 | rs4926595 | A | G | 2 | 0.7637 | 0.7637 | 0.9958 | 0.9958 | 0.9792 | 0 |
| 1 | 54567906 | rs7979393 | C | T | 2 | 0.7638 | 0.7638 | 1.0214 | 1.0214 | 0.6589 | 0 |
| 1 | 54548069 | rs1163211 | T | C | 2 | 0.7652 | 0.7073 | 1.016 | 1.0305 | 0.1517 | 51.33 |
| 1 | 54595626 | rs592650 | C | T | 2 | 0.7652 | 0.7652 | 1.012 | 1.012 | 0.4495 | 0 |
| 1 | 53568980 | rs2242274 | G | A | 2 | 0.7656 | 0.6865 | 1.0052 | 1.0149 | 0.0471 | 74.64 |
| 1 | 53666878 | rs7811715 | C | T | 2 | 0.7656 | 0.7656 | 1.0063 | 1.0063 | 0.5699 | 0 |
| 1 | 54364094 | rs1274984 | A | C | 2 | 0.7662 | 0.7662 | 0.9959 | 0.9959 | 0.4705 | 0 |
| 1 | 54504955 | rs1209251 | G | T | 2 | 0.7665 | 0.7665 | 0.9899 | 0.9899 | 0.7984 | 0 |
| 1 | 53777513 | rs2788034 | A | T | 2 | 0.7668 | 0.8733 | 0.9815 | 0.9747 | 0.0109 | 84.58 |
| 1 | 53511094 | rs6588459 | C | T | 2 | 0.7669 | 0.7669 | 1.0071 | 1.0071 | 0.9784 | 0 |
| 1 | 54363897 | rs7541876 | C | G | 2 | 0.7674 | 0.7674 | 0.996 | 0.996 | 0.4709 | 0 |
| 1 | 53535924 | rs2275455 | A | C | 2 | 0.7675 | 0.75 | 1.0094 | 1.0164 | 0.1153 | 59.68 |
| 1 | 53501578 | rs1180696 | G | A | 2 | 0.7681 | 0.7681 | 1.0091 | 1.0091 | 0.6168 | 0 |
| 1 | 53512914 | rs1256782 | G | A | 2 | 0.7681 | 0.7566 | 1.0091 | 1.018 | 0.0679 | 70.01 |
| 1 | 54487691 | rs3529939 | A | C | 2 | 0.7695 | 0.7695 | 0.996 | 0.996 | 0.328 | 0 |
| 1 | 54502749 | rs1206813 | C | T | 2 | 0.7701 | 0.7701 | 0.99 | 0.99 | 0.7926 | 0 |
| 1 | 53501409 | rs1710768 | G | A | 2 | 0.7702 | 0.7702 | 0.9897 | 0.9897 | 0.7686 | 0 |
| 1 | 53523904 | rs1049317 | A | G | 2 | 0.7711 | 0.7585 | 1.009 | 1.0182 | 0.063 | 71.08 |
| 1 | 53574050 | rs7534682 | A | G | 2 | 0.7712 | 0.7712 | 0.9917 | 0.9917 | 0.8407 | 0 |
| 1 | 54057905 | rs1209712 | A | G | 2 | 0.7718 | 0.7718 | 0.9824 | 0.9824 | 0.8317 | 0 |

| | | | | | | | | | | | |
|---|---|---|---|---|---|---|---|---|---|---|---|
| 1 | 53559503 | rs1203138 | G | A | 2 | 0.7719 | 0.7719 | 0.9919 | 0.9919 | 0.601 | 0 |
| 1 | 54629425 | rs1710996 | A | T | 2 | 0.7724 | 0.7442 | 1.0154 | 1.0251 | 0.1624 | 48.76 |
| 1 | 54176081 | rs1120620 | A | G | 2 | 0.7727 | 0.7727 | 0.9781 | 0.9781 | 0.974 | 0 |
| 1 | 54240555 | rs7266230 | A | C | 2 | 0.7735 | 0.7735 | 1.0049 | 1.0049 | 0.9449 | 0 |
| 1 | 53866272 | rs1208015 | C | T | 2 | 0.7736 | 0.7736 | 1.0123 | 1.0123 | 0.7993 | 0 |
| 1 | 54596128 | rs653948 | A | C | 2 | 0.7736 | 0.7832 | 1.0114 | 1.0112 | 0.3046 | 5.14 |
| 1 | 53752640 | rs1288510 | G | A | 2 | 0.7739 | 0.7739 | 1.0041 | 1.0041 | 0.3685 | 0 |
| 1 | 53511321 | rs6421491 | A | G | 2 | 0.7748 | 0.7748 | 1.0069 | 1.0069 | 0.9674 | 0 |
| 1 | 53756465 | rs1288507 | A | G | 2 | 0.7749 | 0.7673 | 0.9951 | 0.9947 | 0.314 | 1.37 |
| 1 | 53510831 | rs7552336 | C | A | 2 | 0.776 | 0.776 | 1.0087 | 1.0087 | 0.6374 | 0 |
| 1 | 53623654 | rs1021883 | G | A | 2 | 0.7762 | 0.7762 | 0.9766 | 0.9766 | 0.8465 | 0 |
| 1 | 54144396 | rs6177476 | A | G | 2 | 0.7766 | 0.8619 | 0.9832 | 0.9882 | 0.263 | 20.18 |
| 1 | 53815420 | rs4926975 | G | A | 2 | 0.7768 | 0.7768 | 0.996 | 0.996 | 0.4418 | 0 |
| 1 | 53955093 | rs5633442 | T | C | 2 | 0.777 | 0.777 | 0.9847 | 0.9847 | 0.6194 | 0 |
| 1 | 53826484 | rs4926597 | T | C | 2 | 0.7775 | 0.7775 | 0.996 | 0.996 | 0.9375 | 0 |
| 1 | 53970223 | rs1141224 | G | A | 2 | 0.7775 | 0.6928 | 0.9904 | 0.9764 | 0.0968 | 63.73 |
| 1 | 53553657 | rs4131204 | G | A | 2 | 0.7778 | 0.7658 | 1.009 | 1.0101 | 0.2916 | 10.08 |
| 1 | 53551065 | rs2282340 | G | A | 2 | 0.7786 | 0.744 | 1.009 | 1.0128 | 0.2341 | 29.38 |
| 1 | 54651384 | rs955081 | A | G | 2 | 0.7786 | 0.9838 | 1.0146 | 0.9985 | 0.1762 | 45.33 |
| 1 | 54608119 | rs1571540 | G | A | 2 | 0.7796 | 0.7796 | 1.0042 | 1.0042 | 0.576 | 0 |
| 1 | 54104363 | rs1206263 | C | G | 2 | 0.7798 | 0.9742 | 0.9788 | 1.0041 | 0.1181 | 59.06 |
| 1 | 53550543 | rs2282339 | A | G | 2 | 0.78 | 0.78 | 0.9925 | 0.9925 | 0.6499 | 0 |
| 1 | 53923890 | rs1412621 | C | T | 2 | 0.7801 | 0.8293 | 0.9911 | 0.984 | 0.021 | 81.22 |
| 1 | 53505358 | rs7290319 | G | A | 2 | 0.7806 | 0.7806 | 0.9902 | 0.9902 | 0.899 | 0 |
| 1 | 53588988 | rs3766791 | G | A | 2 | 0.7816 | 0.7763 | 1.0051 | 1.0054 | 0.307 | 4.18 |
| 1 | 53640300 | rs1150762 | G | A | 2 | 0.782 | 0.782 | 1.0134 | 1.0134 | 0.5444 | 0 |
| 1 | 54117669 | rs7266063 | C | T | 2 | 0.7824 | 0.9558 | 1.0103 | 0.9966 | 0.1186 | 58.95 |
| 1 | 53556122 | rs1146186 | C | T | 2 | 0.7833 | 0.7833 | 0.9904 | 0.9904 | 0.9656 | 0 |
| 1 | 53544807 | rs1710786 | C | T | 2 | 0.786 | 0.7395 | 1.0086 | 1.0142 | 0.196 | 40.18 |
| 1 | 53757331 | rs1288501 | A | C | 2 | 0.786 | 0.5584 | 1.0067 | 0.9463 | 0.0162 | 82.71 |
| 1 | 54363021 | rs1883457 | C | A | 2 | 0.7868 | 0.7868 | 0.9951 | 0.9951 | 0.5857 | 0 |
| 1 | 53762781 | rs1120613 | G | A | 2 | 0.787 | 0.5579 | 0.9956 | 1.0365 | 0.0296 | 78.87 |
| 1 | 54419526 | rs1212796 | T | A | 2 | 0.7872 | 0.7872 | 1.0036 | 1.0036 | 0.9884 | 0 |
| 1 | 54630325 | rs613910 | G | C | 2 | 0.7872 | 0.7872 | 1.0037 | 1.0037 | 0.5849 | 0 |
| 1 | 53501402 | rs7441970 | A | G | 2 | 0.7873 | 0.7873 | 0.9907 | 0.9907 | 0.7548 | 0 |
| 1 | 53501451 | rs1710769 | A | C | 2 | 0.7873 | 0.7873 | 0.9907 | 0.9907 | 0.7548 | 0 |
| 1 | 53499784 | rs6662474 | A | G | 2 | 0.7874 | 0.7894 | 1.0064 | 1.0098 | 0.1281 | 56.81 |
| 1 | 53600471 | rs5672586 | T | C | 2 | 0.7878 | 0.9146 | 1.0066 | 1.0036 | 0.1806 | 44.22 |
| 1 | 53785654 | rs1120614 | T | C | 2 | 0.7884 | 0.8217 | 0.9749 | 0.977 | 0.2767 | 15.47 |
| 1 | 53993909 | rs7267517 | C | T | 2 | 0.7888 | 0.7888 | 0.9872 | 0.9872 | 0.5396 | 0 |
| 1 | 53921278 | rs6159283 | C | A | 2 | 0.7891 | 0.773 | 0.9957 | 0.9932 | 0.1518 | 51.31 |
| 1 | 53555783 | rs1288403 | A | C | 2 | 0.7898 | 0.7898 | 0.9948 | 0.9948 | 0.9777 | 0 |
| 1 | 53575181 | rs1469383 | T | C | 2 | 0.7898 | 0.7898 | 0.9875 | 0.9875 | 0.7471 | 0 |
| 1 | 53771564 | rs1288492 | C | T | 2 | 0.7898 | 0.7898 | 1.0045 | 1.0045 | 0.7276 | 0 |
| 1 | 54673943 | rs1120630 | C | T | 2 | 0.7905 | 0.7905 | 1.0064 | 1.0064 | 0.7067 | 0 |
| 1 | 54215327 | rs1154830 | G | A | 2 | 0.7909 | 0.7909 | 1.0046 | 1.0046 | 0.5988 | 0 |
| 1 | 54361288 | rs2268181 | T | C | 2 | 0.791 | 0.791 | 0.9953 | 0.9953 | 0.4623 | 0 |
| 1 | 54438180 | rs1468868 | C | G | 2 | 0.7915 | 0.7915 | 1.0172 | 1.0172 | 0.4217 | 0 |

| | | | | | | | | | | | |
|---|---|---|---|---|---|---|---|---|---|---|---|
| 1 | 54497983 | rs1209251 | A | C | 2 | 0.7915 | 0.7915 | 1.0188 | 1.0188 | 0.5949 | 0 |
| 1 | 53501472 | rs1710769 | G | A | 2 | 0.7916 | 0.7916 | 0.991 | 0.991 | 0.7507 | 0 |
| 1 | 54390019 | rs1088882 | A | T | 2 | 0.7926 | 0.7755 | 0.9949 | 0.9923 | 0.1725 | 46.26 |
| 1 | 53568099 | rs1120609 | T | C | 2 | 0.7927 | 0.6996 | 1.0046 | 1.014 | 0.0501 | 73.96 |
| 1 | 54299440 | rs4927041 | G | A | 2 | 0.7928 | 0.7928 | 1.0045 | 1.0045 | 0.8979 | 0 |
| 1 | 54619404 | rs7519887 | C | T | 2 | 0.7928 | 0.7928 | 1.0044 | 1.0044 | 0.6863 | 0 |
| 1 | 53543312 | rs1274597 | C | T | 2 | 0.793 | 0.7405 | 1.0051 | 1.0083 | 0.2155 | 34.82 |
| 1 | 54538699 | rs1208230 | C | T | 2 | 0.7932 | 0.7932 | 1.0186 | 1.0186 | 0.6578 | 0 |
| 1 | 54251865 | rs7488769 | T | A | 2 | 0.7935 | 0.9603 | 0.9764 | 0.9913 | 0.056 | 72.61 |
| 1 | 54648888 | rs1206836 | C | A | 2 | 0.7935 | 0.7935 | 1.017 | 1.017 | 0.5181 | 0 |
| 1 | 53564199 | rs3766772 | G | A | 2 | 0.7938 | 0.6922 | 1.0045 | 1.0139 | 0.0603 | 71.66 |
| 1 | 54366695 | rs7266239 | G | T | 2 | 0.7948 | 0.7948 | 0.9952 | 0.9952 | 0.6119 | 0 |
| 1 | 53634222 | rs7415149 | A | C | 2 | 0.795 | 0.7804 | 1.0263 | 1.0295 | 0.3065 | 4.36 |
| 1 | 53821821 | rs1120614 | C | T | 2 | 0.7956 | 0.7956 | 0.9957 | 0.9957 | 0.5398 | 0 |
| 1 | 53651529 | rs5676052 | G | A | 2 | 0.7957 | 0.7957 | 1.0105 | 1.0105 | 0.8515 | 0 |
| 1 | 53826464 | rs4926979 | A | G | 2 | 0.7958 | 0.7958 | 0.9964 | 0.9964 | 0.7997 | 0 |
| 1 | 54648889 | rs1206837 | C | A | 2 | 0.7959 | 0.7959 | 1.0168 | 1.0168 | 0.5152 | 0 |
| 1 | 53573600 | rs3766778 | T | G | 2 | 0.7962 | 0.7962 | 0.9926 | 0.9926 | 0.8309 | 0 |
| 1 | 53619412 | rs7523016 | A | G | 2 | 0.7965 | 0.7965 | 0.9803 | 0.9803 | 0.4195 | 0 |
| 1 | 53564198 | rs3766771 | T | C | 2 | 0.7968 | 0.6935 | 1.0045 | 1.0138 | 0.06 | 71.74 |
| 1 | 53506814 | rs7290320 | G | A | 2 | 0.7981 | 0.7981 | 0.9912 | 0.9912 | 0.7702 | 0 |
| 1 | 53544513 | rs1256702 | C | T | 2 | 0.7981 | 0.7981 | 1.0131 | 1.0131 | 0.4311 | 0 |
| 1 | 54522240 | rs1206641 | T | C | 2 | 0.7981 | 0.7981 | 0.9899 | 0.9899 | 0.6936 | 0 |
| 1 | 53549671 | rs3416325 | C | T | 2 | 0.7982 | 0.7277 | 1.005 | 1.0098 | 0.1693 | 47.08 |
| 1 | 53614403 | rs7902299 | C | G | 2 | 0.7987 | 0.8504 | 1.0058 | 1.0047 | 0.2691 | 18.13 |
| 1 | 53901838 | rs1766644 | G | A | 2 | 0.7987 | 0.7987 | 0.9955 | 0.9955 | 0.5947 | 0 |
| 1 | 53528264 | rs6694494 | C | A | 2 | 0.799 | 0.799 | 1.0087 | 1.0087 | 0.6365 | 0 |
| 1 | 53593813 | rs899976 | G | A | 2 | 0.7997 | 0.7997 | 1.0041 | 1.0041 | 0.4586 | 0 |
| 1 | 53821781 | rs3537886 | T | C | 2 | 0.7998 | 0.7998 | 0.9958 | 0.9958 | 0.5349 | 0 |
| 1 | 53525644 | rs1710778 | C | T | 2 | 0.8001 | 0.8001 | 1.0099 | 1.0099 | 0.6352 | 0 |
| 1 | 53566795 | rs1275782 | T | C | 2 | 0.8002 | 0.7052 | 1.0044 | 1.0127 | 0.0677 | 70.04 |
| 1 | 53799408 | rs1207732 | C | A | 2 | 0.8004 | 0.8004 | 1.0131 | 1.0131 | 0.5626 | 0 |
| 1 | 53597998 | rs1159081 | G | A | 2 | 0.8005 | 0.8005 | 0.993 | 0.993 | 0.735 | 0 |
| 1 | 53799410 | rs1206715 | A | G | 2 | 0.8005 | 0.8005 | 1.0131 | 1.0131 | 0.562 | 0 |
| 1 | 54314924 | rs1120622 | A | C | 2 | 0.8006 | 0.9849 | 0.9963 | 0.9996 | 0.1244 | 57.65 |
| 1 | 53630300 | rs7892511 | C | A | 2 | 0.801 | 0.801 | 0.9795 | 0.9795 | 0.8062 | 0 |
| 1 | 54299674 | rs2064457 | A | G | 2 | 0.8011 | 0.8011 | 1.0043 | 1.0043 | 0.8933 | 0 |
| 1 | 53530786 | rs2790427 | A | G | 2 | 0.8012 | 0.8012 | 0.9964 | 0.9964 | 0.8019 | 0 |
| 1 | 53593874 | rs899975 | C | T | 2 | 0.8015 | 0.8015 | 1.004 | 1.004 | 0.4007 | 0 |
| 1 | 54030132 | rs1465034 | A | G | 2 | 0.8017 | 0.8017 | 1.0039 | 1.0039 | 0.5535 | 0 |
| 1 | 53544578 | rs1120608 | T | C | 2 | 0.8022 | 0.7493 | 1.0049 | 1.0079 | 0.2194 | 33.69 |
| 1 | 53907942 | rs1288602 | C | G | 2 | 0.8038 | 0.8038 | 0.9959 | 0.9959 | 0.8915 | 0 |
| 1 | 53578293 | rs3766780 | A | G | 2 | 0.8039 | 0.8039 | 0.9803 | 0.9803 | 0.5591 | 0 |
| 1 | 53511139 | rs6588460 | A | G | 2 | 0.8042 | 0.8042 | 1.0061 | 1.0061 | 0.9655 | 0 |
| 1 | 54320986 | rs1509834 | C | T | 2 | 0.8042 | 0.8042 | 1.0116 | 1.0116 | 0.5573 | 0 |
| 1 | 54575599 | rs1710976 | T | A | 2 | 0.8047 | 0.8047 | 1.0175 | 1.0175 | 0.6595 | 0 |
| 1 | 53827481 | rs1214367 | C | A | 2 | 0.805 | 0.805 | 0.9958 | 0.9958 | 0.4908 | 0 |
| 1 | 54105063 | rs1120618 | G | A | 2 | 0.8052 | 0.9736 | 0.9813 | 1.004 | 0.1353 | 55.17 |

| | | | | | | | | | | | |
|---|---|---|---|---|---|---|---|---|---|---|---|
| 1 | 53611605 | rs1240873 | C | T | 2 | 0.8053 | 0.8053 | 1.007 | 1.007 | 0.4295 | 0 |
| 1 | 53567721 | rs1120609 | T | C | 2 | 0.8054 | 0.7066 | 1.0043 | 1.0132 | 0.0575 | 72.28 |
| 1 | 53828651 | rs1120615 | G | T | 2 | 0.8062 | 0.8062 | 0.9958 | 0.9958 | 0.4925 | 0 |
| 1 | 54328358 | rs6681067 | G | A | 2 | 0.8064 | 0.8064 | 0.9955 | 0.9955 | 0.6781 | 0 |
| 1 | 53535876 | rs2275453 | T | C | 2 | 0.8068 | 0.7802 | 1.0078 | 1.0145 | 0.1131 | 60.15 |
| 1 | 53567229 | rs5698088 | C | T | 2 | 0.8069 | 0.8069 | 1.0079 | 1.0079 | 0.3973 | 0 |
| 1 | 53567864 | rs1120609 | A | G | 2 | 0.8075 | 0.7077 | 1.0042 | 1.0132 | 0.0571 | 72.36 |
| 1 | 53556532 | rs2306461 | G | A | 2 | 0.8083 | 0.8756 | 1.0101 | 1.0082 | 0.2088 | 36.7 |
| 1 | 53821507 | rs1120614 | A | G | 2 | 0.8085 | 0.8085 | 0.996 | 0.996 | 0.5271 | 0 |
| 1 | 53832697 | rs1204196 | T | C | 2 | 0.8089 | 0.8089 | 0.9966 | 0.9966 | 0.3238 | 0 |
| 1 | 54207709 | rs1113699 | C | T | 2 | 0.8089 | 0.8089 | 0.9909 | 0.9909 | 0.7469 | 0 |
| 1 | 53832748 | rs1202416 | C | G | 2 | 0.8092 | 0.8092 | 0.9966 | 0.9966 | 0.3211 | 0 |
| 1 | 54178200 | rs1710921 | A | G | 2 | 0.8092 | 0.8092 | 0.9813 | 0.9813 | 0.8289 | 0 |
| 1 | 53681020 | rs7460383 | T | C | 2 | 0.8093 | 0.8093 | 1.0051 | 1.0051 | 0.6441 | 0 |
| 1 | 54527011 | rs1206035 | G | A | 2 | 0.8094 | 0.8094 | 1.0171 | 1.0171 | 0.6179 | 0 |
| 1 | 53535175 | rs1531699 | T | C | 2 | 0.8096 | 0.7743 | 1.0075 | 1.0122 | 0.1823 | 43.79 |
| 1 | 53762226 | rs4926968 | C | A | 2 | 0.8099 | 0.9388 | 1.006 | 1.0023 | 0.268 | 18.51 |
| 1 | 53827661 | rs1088878 | C | T | 2 | 0.8105 | 0.8105 | 0.9966 | 0.9966 | 0.8694 | 0 |
| 1 | 54609689 | rs3398151 | A | G | 2 | 0.8108 | 0.8108 | 0.9967 | 0.9967 | 0.5971 | 0 |
| 1 | 53558366 | rs1212965 | T | C | 2 | 0.8116 | 0.8116 | 0.9932 | 0.9932 | 0.8579 | 0 |
| 1 | 53544289 | rs1120608 | C | T | 2 | 0.8117 | 0.8117 | 0.9934 | 0.9934 | 0.7517 | 0 |
| 1 | 53628740 | rs1126097 | G | A | 2 | 0.8117 | 0.8117 | 0.9808 | 0.9808 | 0.7846 | 0 |
| 1 | 53502905 | rs7290318 | T | A | 2 | 0.8123 | 0.8123 | 0.9919 | 0.9919 | 0.7481 | 0 |
| 1 | 53568162 | rs1120609 | C | T | 2 | 0.8125 | 0.7068 | 1.0041 | 1.014 | 0.0444 | 75.25 |
| 1 | 54573946 | rs7408519 | T | C | 2 | 0.813 | 0.8114 | 0.991 | 0.99 | 0.2714 | 17.34 |
| 1 | 54627097 | rs7528248 | C | T | 2 | 0.8134 | 0.8134 | 1.0045 | 1.0045 | 0.3734 | 0 |
| 1 | 53817020 | rs6176965 | A | G | 2 | 0.8135 | 0.8135 | 0.9961 | 0.9961 | 0.5382 | 0 |
| 1 | 53535478 | rs1288386 | G | A | 2 | 0.814 | 0.814 | 0.9967 | 0.9967 | 0.3323 | 0 |
| 1 | 53635392 | rs1209228 | T | C | 2 | 0.8143 | 0.8143 | 0.988 | 0.988 | 0.8573 | 0 |
| 1 | 53806783 | rs7749183 | G | A | 2 | 0.8164 | 0.8164 | 1.0087 | 1.0087 | 0.9878 | 0 |
| 1 | 53500537 | rs8007719 | G | A | 2 | 0.8166 | 0.8166 | 0.9918 | 0.9918 | 0.7439 | 0 |
| 1 | 53511341 | rs6421492 | T | A | 2 | 0.8173 | 0.8173 | 1.0055 | 1.0055 | 0.8932 | 0 |
| 1 | 53557976 | rs6096188 | C | T | 2 | 0.8177 | 0.8125 | 1.0074 | 1.0078 | 0.3094 | 3.2 |
| 1 | 53807341 | rs1427831 | G | A | 2 | 0.8179 | 0.8179 | 1.0087 | 1.0087 | 0.9939 | 0 |
| 1 | 54479491 | rs1147259 | C | T | 2 | 0.818 | 0.818 | 1.0103 | 1.0103 | 0.568 | 0 |
| 1 | 53899583 | rs1776416 | G | A | 2 | 0.8183 | 0.815 | 0.9867 | 0.9863 | 0.3127 | 1.88 |
| 1 | 53490746 | rs7290311 | A | T | 2 | 0.8185 | 0.8185 | 1.0078 | 1.0078 | 0.6395 | 0 |
| 1 | 53535269 | rs1531702 | G | A | 2 | 0.8189 | 0.7817 | 1.0071 | 1.0119 | 0.1788 | 44.68 |
| 1 | 53501433 | rs1180863 | T | C | 2 | 0.8192 | 0.8192 | 1.007 | 1.007 | 0.5644 | 0 |
| 1 | 53511141 | rs1116818 | C | T | 2 | 0.8194 | 0.8194 | 1.0055 | 1.0055 | 0.9654 | 0 |
| 1 | 53547800 | rs6687909 | A | G | 2 | 0.8194 | 0.8912 | 1.0089 | 1.0067 | 0.2103 | 36.29 |
| 1 | 54200987 | rs797915 | A | G | 2 | 0.8197 | 0.8197 | 0.9826 | 0.9826 | 0.745 | 0 |
| 1 | 53908908 | rs5911345 | C | A | 2 | 0.8198 | 0.8198 | 0.9913 | 0.9913 | 0.3214 | 0 |
| 1 | 53854860 | rs7505724 | G | A | 2 | 0.82 | 0.82 | 0.98 | 0.98 | 0.3498 | 0 |
| 1 | 53817481 | rs7524966 | G | A | 2 | 0.8202 | 0.8202 | 0.9962 | 0.9962 | 0.5386 | 0 |
| 1 | 53535205 | rs1531700 | C | T | 2 | 0.8217 | 0.7836 | 1.007 | 1.0118 | 0.1778 | 44.94 |
| 1 | 53761249 | rs1120613 | G | C | 2 | 0.8224 | 0.6226 | 0.9964 | 1.0236 | 0.0816 | 67.02 |
| 1 | 53972205 | rs2296758 | T | C | 2 | 0.8224 | 0.7462 | 0.9962 | 0.9905 | 0.0923 | 64.72 |

| | | | | | | | | | | | |
|---|---|---|---|---|---|---|---|---|---|---|---|
| 1 | 53563287 | rs3497899 | G | A | 2 | 0.8226 | 0.7272 | 1.0044 | 1.0111 | 0.1284 | 56.75 |
| 1 | 53564777 | rs7827967 | C | T | 2 | 0.8236 | 0.8236 | 1.0071 | 1.0071 | 0.3336 | 0 |
| 1 | 53933225 | rs4130616 | T | C | 2 | 0.8238 | 0.7957 | 0.9964 | 0.9941 | 0.1585 | 49.7 |
| 1 | 53535055 | rs7817527 | G | A | 2 | 0.8241 | 0.7856 | 1.0069 | 1.0117 | 0.1769 | 45.15 |
| 1 | 53519423 | rs1159283 | T | G | 2 | 0.8243 | 0.8243 | 1.0079 | 1.0079 | 0.9618 | 0 |
| 1 | 53566635 | rs2854502 | A | G | 2 | 0.8246 | 0.721 | 1.0038 | 1.0114 | 0.0822 | 66.9 |
| 1 | 53547371 | rs1710787 | A | G | 2 | 0.8247 | 0.7673 | 1.0043 | 1.0073 | 0.2203 | 33.45 |
| 1 | 54615566 | rs1157635 | C | T | 2 | 0.8253 | 0.794 | 1.0076 | 1.0108 | 0.2382 | 28.12 |
| 1 | 53851165 | rs1224006 | T | C | 2 | 0.8254 | 0.8254 | 0.9807 | 0.9807 | 0.3488 | 0 |
| 1 | 53557484 | rs1288404 | T | C | 2 | 0.8255 | 0.8255 | 1.0037 | 1.0037 | 0.3926 | 0 |
| 1 | 54339657 | rs1211817 | T | C | 2 | 0.826 | 0.826 | 0.9959 | 0.9959 | 0.7137 | 0 |
| 1 | 53620759 | rs1125356 | C | A | 2 | 0.8263 | 0.8263 | 1.0178 | 1.0178 | 0.4858 | 0 |
| 1 | 53625090 | rs3595957 | G | C | 2 | 0.8265 | 0.8851 | 1.0049 | 1.0037 | 0.2589 | 21.55 |
| 1 | 53820667 | rs6176965 | T | C | 2 | 0.8268 | 0.8268 | 0.9963 | 0.9963 | 0.4139 | 0 |
| 1 | 53535222 | rs1531701 | G | A | 2 | 0.8272 | 0.7862 | 1.0068 | 1.0119 | 0.168 | 47.4 |
| 1 | 54581474 | rs7657923 | C | T | 2 | 0.8274 | 0.8274 | 1.0187 | 1.0187 | 0.8051 | 0 |
| 1 | 53891296 | rs1203235 | G | A | 2 | 0.8279 | 0.7927 | 0.9966 | 0.9933 | 0.1119 | 60.43 |
| 1 | 53746473 | rs2297658 | T | C | 2 | 0.828 | 0.828 | 0.9957 | 0.9957 | 0.8213 | 0 |
| 1 | 53778904 | rs1288479 | T | C | 2 | 0.8282 | 0.8282 | 0.9959 | 0.9959 | 0.7034 | 0 |
| 1 | 53831391 | rs1933538 | C | T | 2 | 0.8285 | 0.8285 | 0.997 | 0.997 | 0.3268 | 0 |
| 1 | 53829345 | rs4926980 | C | T | 2 | 0.8296 | 0.8296 | 0.997 | 0.997 | 0.3324 | 0 |
| 1 | 53904579 | rs1288598 | T | G | 2 | 0.8298 | 0.8298 | 0.9963 | 0.9963 | 0.5677 | 0 |
| 1 | 53612267 | rs7825053 | C | T | 2 | 0.8299 | 0.8758 | 1.0048 | 1.0039 | 0.2724 | 16.99 |
| 1 | 54030702 | rs7522812 | C | A | 2 | 0.8299 | 0.8299 | 0.9891 | 0.9891 | 0.8592 | 0 |
| 1 | 54588898 | rs5799600 | C | T | 2 | 0.8301 | 0.945 | 1.013 | 1.0074 | 0.078 | 67.8 |
| 1 | 53832151 | rs1157744 | C | T | 2 | 0.8302 | 0.8302 | 0.997 | 0.997 | 0.3309 | 0 |
| 1 | 54141867 | rs1165330 | C | A | 2 | 0.8305 | 0.8305 | 1.0069 | 1.0069 | 0.8122 | 0 |
| 1 | 53782093 | rs884586 | A | C | 2 | 0.8308 | 0.888 | 0.9833 | 0.9811 | 0.0856 | 66.17 |
| 1 | 53742081 | rs6699794 | G | A | 2 | 0.8312 | 0.8312 | 1.0029 | 1.0029 | 0.543 | 0 |
| 1 | 53754361 | rs6685032 | C | G | 2 | 0.8316 | 0.8316 | 1.0037 | 1.0037 | 0.5189 | 0 |
| 1 | 53560428 | rs1295456 | C | T | 2 | 0.8317 | 0.717 | 0.9964 | 0.9872 | 0.0513 | 73.67 |
| 1 | 53936383 | rs1157952 | G | A | 2 | 0.8317 | 0.7997 | 0.9966 | 0.994 | 0.1463 | 52.61 |
| 1 | 54278471 | rs3565548 | C | T | 2 | 0.8317 | 0.8317 | 1.0037 | 1.0037 | 0.9506 | 0 |
| 1 | 54450192 | rs7548146 | C | T | 2 | 0.8323 | 0.8323 | 1.0115 | 1.0115 | 0.6962 | 0 |
| 1 | 53535085 | rs1531698 | C | T | 2 | 0.8328 | 0.7914 | 1.0066 | 1.0114 | 0.1753 | 45.57 |
| 1 | 53501159 | rs7290317 | A | G | 2 | 0.8345 | 0.8345 | 0.9928 | 0.9928 | 0.711 | 0 |
| 1 | 53569432 | rs2242275 | C | T | 2 | 0.8349 | 0.8349 | 1.0121 | 1.0121 | 0.4073 | 0 |
| 1 | 53622050 | rs6657387 | A | G | 2 | 0.835 | 0.835 | 0.9832 | 0.9832 | 0.7264 | 0 |
| 1 | 53755679 | rs1203902 | A | C | 2 | 0.8351 | 0.5368 | 0.9966 | 1.042 | 0.0117 | 84.26 |
| 1 | 54621887 | rs7570910 | C | T | 2 | 0.8351 | 0.8351 | 0.9903 | 0.9903 | 0.4607 | 0 |
| 1 | 54219241 | rs1181161 | C | T | 2 | 0.8354 | 0.8354 | 1.0031 | 1.0031 | 0.6762 | 0 |
| 1 | 53593017 | rs1679965 | T | A | 2 | 0.8363 | 0.8363 | 0.9972 | 0.9972 | 0.5306 | 0 |
| 1 | 54474472 | rs6672989 | T | C | 2 | 0.8367 | 0.8367 | 0.9972 | 0.9972 | 0.535 | 0 |
| 1 | 53559846 | rs7875130 | G | A | 2 | 0.837 | 0.837 | 1.0066 | 1.0066 | 0.3232 | 0 |
| 1 | 54001820 | rs1212149 | T | C | 2 | 0.8371 | 0.9729 | 1.0029 | 0.9993 | 0.1538 | 50.83 |
| 1 | 53524715 | rs1580356 | G | A | 2 | 0.8373 | 0.8373 | 1.007 | 1.007 | 0.6994 | 0 |
| 1 | 53902833 | rs1288595 | G | A | 2 | 0.8379 | 0.8379 | 0.9882 | 0.9882 | 0.3433 | 0 |
| 1 | 53535860 | rs2275452 | G | A | 2 | 0.8381 | 0.7975 | 1.0066 | 1.0131 | 0.1208 | 58.44 |

| | | | | | | | | | | | |
|---|---|---|---|---|---|---|---|---|---|---|---|
| 1 | 53535443 | rs1531703 | G | A | 2 | 0.8388 | 0.7892 | 1.0064 | 1.0122 | 0.1609 | 49.14 |
| 1 | 54646015 | rs1206227 | A | C | 2 | 0.8389 | 0.7796 | 1.0111 | 0.9663 | 0.0481 | 74.39 |
| 1 | 53566351 | rs6752406 | C | T | 2 | 0.8392 | 0.7276 | 1.0035 | 1.0112 | 0.0786 | 67.67 |
| 1 | 53556695 | rs7699026 | A | C | 2 | 0.8394 | 0.8394 | 0.9929 | 0.9929 | 0.995 | 0 |
| 1 | 53622147 | rs6668684 | G | A | 2 | 0.8394 | 0.8394 | 0.9837 | 0.9837 | 0.7302 | 0 |
| 1 | 53903238 | rs1288596 | A | T | 2 | 0.8398 | 0.8398 | 0.9883 | 0.9883 | 0.3432 | 0 |
| 1 | 53509949 | rs7535299 | T | G | 2 | 0.8406 | 0.8406 | 0.9931 | 0.9931 | 0.6975 | 0 |
| 1 | 53565197 | rs6177079 | C | T | 2 | 0.8406 | 0.7275 | 1.0035 | 1.0113 | 0.0776 | 67.89 |
| 1 | 53518803 | rs7290513 | G | A | 2 | 0.8407 | 0.8407 | 1.0068 | 1.0068 | 0.6352 | 0 |
| 1 | 54180994 | rs1256692 | G | A | 2 | 0.8408 | 0.8408 | 0.9845 | 0.9845 | 0.9884 | 0 |
| 1 | 53608711 | rs1274475 | C | T | 2 | 0.8413 | 0.9164 | 1.0045 | 1.0028 | 0.2385 | 28.04 |
| 1 | 53509589 | rs6099040 | C | T | 2 | 0.8414 | 0.8414 | 0.993 | 0.993 | 0.6948 | 0 |
| 1 | 53564273 | rs3766774 | G | A | 2 | 0.8417 | 0.7286 | 1.0034 | 1.0113 | 0.0767 | 68.09 |
| 1 | 53608133 | rs3737990 | C | A | 2 | 0.8421 | 0.9186 | 1.0045 | 1.0028 | 0.2362 | 28.73 |
| 1 | 53600917 | rs1240677 | C | T | 2 | 0.8435 | 0.8318 | 0.997 | 0.9965 | 0.2759 | 15.76 |
| 1 | 54122797 | rs1918176 | T | G | 2 | 0.8438 | 0.8438 | 0.9951 | 0.9951 | 0.7192 | 0 |
| 1 | 53748286 | rs1288516 | G | A | 2 | 0.8439 | 0.8439 | 0.9971 | 0.9971 | 0.9373 | 0 |
| 1 | 53566254 | rs6176811 | C | A | 2 | 0.844 | 0.7296 | 1.0034 | 1.0112 | 0.0782 | 67.76 |
| 1 | 53554680 | rs3816744 | T | C | 2 | 0.8445 | 0.8445 | 0.9945 | 0.9945 | 0.7816 | 0 |
| 1 | 53628789 | rs3565825 | T | A | 2 | 0.8446 | 0.8461 | 1.0045 | 1.0045 | 0.3161 | 0.49 |
| 1 | 54691465 | rs9012 | C | T | 2 | 0.8454 | 0.8454 | 0.9971 | 0.9971 | 0.9575 | 0 |
| 1 | 53751610 | rs4623641 | C | T | 2 | 0.8455 | 0.8455 | 0.9972 | 0.9972 | 0.732 | 0 |
| 1 | 53501202 | rs7290317 | C | G | 2 | 0.8457 | 0.8457 | 0.9933 | 0.9933 | 0.6994 | 0 |
| 1 | 54345258 | rs1710952 | G | A | 2 | 0.8458 | 0.8458 | 0.9964 | 0.9964 | 0.6537 | 0 |
| 1 | 53509852 | rs7542983 | G | C | 2 | 0.846 | 0.846 | 0.9933 | 0.9933 | 0.6918 | 0 |
| 1 | 53566167 | rs7743370 | C | T | 2 | 0.8463 | 0.8238 | 1.0059 | 1.0074 | 0.2816 | 13.75 |
| 1 | 53504411 | rs1180828 | G | A | 2 | 0.847 | 0.847 | 1.0059 | 1.0059 | 0.5531 | 0 |
| 1 | 53567237 | rs1148558 | C | T | 2 | 0.8473 | 0.7311 | 1.0033 | 1.0111 | 0.0777 | 67.87 |
| 1 | 53620672 | rs7769492 | A | G | 2 | 0.8478 | 0.8478 | 1.0155 | 1.0155 | 0.4715 | 0 |
| 1 | 53611275 | rs3454901 | T | C | 2 | 0.8484 | 0.907 | 1.0043 | 1.003 | 0.256 | 22.48 |
| 1 | 53607194 | rs1273884 | G | A | 2 | 0.8488 | 0.9492 | 1.0028 | 1.0012 | 0.197 | 39.92 |
| 1 | 53565946 | rs6177079 | T | C | 2 | 0.8496 | 0.7325 | 1.0033 | 1.0111 | 0.078 | 67.81 |
| 1 | 53565625 | rs5750499 | C | G | 2 | 0.8498 | 0.7326 | 1.0033 | 1.011 | 0.0788 | 67.64 |
| 1 | 53566793 | rs1273727 | A | C | 2 | 0.8502 | 0.7329 | 1.0033 | 1.011 | 0.0786 | 67.68 |
| 1 | 53845440 | rs2185077 | T | C | 2 | 0.8503 | 0.8503 | 1.0037 | 1.0037 | 0.7397 | 0 |
| 1 | 54356588 | rs1214135 | C | T | 2 | 0.8503 | 0.8503 | 0.9965 | 0.9965 | 0.6567 | 0 |
| 1 | 53503428 | rs9662321 | A | G | 2 | 0.8506 | 0.8506 | 0.9936 | 0.9936 | 0.6695 | 0 |
| 1 | 53561670 | rs1167843 | C | T | 2 | 0.8513 | 0.8513 | 1.0198 | 1.0198 | 0.9178 | 0 |
| 1 | 53565528 | rs6177079 | T | C | 2 | 0.8515 | 0.7337 | 1.0032 | 1.011 | 0.0791 | 67.57 |
| 1 | 53928180 | rs732385 | G | A | 2 | 0.8519 | 0.8204 | 0.997 | 0.9951 | 0.1845 | 43.21 |
| 1 | 54357315 | rs1240806 | C | A | 2 | 0.8521 | 0.8521 | 0.9966 | 0.9966 | 0.6357 | 0 |
| 1 | 53567421 | rs1088876 | C | T | 2 | 0.8529 | 0.8529 | 1.005 | 1.005 | 0.8059 | 0 |
| 1 | 53500778 | rs7594884 | A | G | 2 | 0.8531 | 0.8531 | 0.9937 | 0.9937 | 0.7079 | 0 |
| 1 | 54345014 | rs1212030 | G | A | 2 | 0.8531 | 0.8531 | 0.9966 | 0.9966 | 0.6542 | 0 |
| 1 | 53946997 | rs5568621 | A | G | 2 | 0.8539 | 0.8597 | 0.9943 | 0.9939 | 0.2673 | 18.73 |
| 1 | 53627978 | rs1383365 | T | C | 2 | 0.8542 | 0.8542 | 1.0048 | 1.0048 | 0.5798 | 0 |
| 1 | 54585461 | rs1112813 | C | G | 2 | 0.8555 | 0.8652 | 0.9895 | 0.9882 | 0.2323 | 29.91 |
| 1 | 53804985 | rs1141346 | C | G | 2 | 0.8558 | 0.8558 | 0.9921 | 0.9921 | 0.4925 | 0 |

| | | | | | | | | | | | |
|---|---|---|---|---|---|---|---|---|---|---|---|
| 1 | 53499822 | rs6673465 | G | A | 2 | 0.8561 | 0.8561 | 0.9938 | 0.9938 | 0.7123 | 0 |
| 1 | 53761755 | rs5590113 | G | T | 2 | 0.8575 | 0.5526 | 0.9971 | 1.0353 | 0.036 | 77.26 |
| 1 | 54673965 | rs1120630 | C | T | 2 | 0.8575 | 0.8575 | 1.0043 | 1.0043 | 0.614 | 0 |
| 1 | 53553522 | rs1158105 | G | A | 2 | 0.8579 | 0.7713 | 1.0035 | 1.008 | 0.1792 | 44.57 |
| 1 | 53567318 | rs1275201 | G | A | 2 | 0.8583 | 0.7362 | 1.0031 | 1.011 | 0.0762 | 68.19 |
| 1 | 53566186 | rs6177079 | A | G | 2 | 0.8584 | 0.7362 | 1.0031 | 1.011 | 0.0762 | 68.2 |
| 1 | 54313553 | rs6145645 | A | G | 2 | 0.8586 | 0.7863 | 1.0028 | 1.0072 | 0.0964 | 63.82 |
| 1 | 53754104 | rs1039982 | G | T | 2 | 0.8587 | 0.4861 | 1.0029 | 1.0438 | 0.0209 | 81.25 |
| 1 | 54322418 | rs1710945 | T | C | 2 | 0.8588 | 0.8588 | 0.9968 | 0.9968 | 0.6363 | 0 |
| 1 | 53596228 | rs946593 | T | C | 2 | 0.8592 | 0.9895 | 1.0026 | 1.0003 | 0.151 | 51.51 |
| 1 | 53598160 | rs7267088 | C | T | 2 | 0.8595 | 0.8595 | 0.9971 | 0.9971 | 0.5303 | 0 |
| 1 | 53771643 | rs1273237 | G | A | 2 | 0.8595 | 0.445 | 0.9965 | 1.1018 | 0.0029 | 88.72 |
| 1 | 53551940 | rs6588462 | G | T | 2 | 0.8597 | 0.7724 | 1.0034 | 1.008 | 0.1778 | 44.93 |
| 1 | 53643272 | rs1120612 | A | G | 2 | 0.8597 | 0.8597 | 0.9912 | 0.9912 | 0.6016 | 0 |
| 1 | 54590188 | rs6683426 | T | C | 2 | 0.8598 | 0.8598 | 0.9972 | 0.9972 | 0.9598 | 0 |
| 1 | 53566032 | rs1156446 | T | C | 2 | 0.8608 | 0.8608 | 0.9938 | 0.9938 | 0.8995 | 0 |
| 1 | 53566191 | rs6176811 | G | A | 2 | 0.8623 | 0.7413 | 1.003 | 1.0104 | 0.0856 | 66.16 |
| 1 | 53531019 | rs7833974 | C | T | 2 | 0.8628 | 0.8089 | 1.0054 | 1.0118 | 0.1276 | 56.91 |
| 1 | 53566571 | rs2845155 | A | G | 2 | 0.8631 | 0.7418 | 1.003 | 1.0103 | 0.0873 | 65.8 |
| 1 | 53494738 | rs7290314 | G | C | 2 | 0.8633 | 0.8633 | 1.0059 | 1.0059 | 0.6822 | 0 |
| 1 | 53901293 | rs6657140 | C | T | 2 | 0.8634 | 0.8634 | 0.997 | 0.997 | 0.5464 | 0 |
| 1 | 53938221 | rs8179495 | G | C | 2 | 0.8634 | 0.8168 | 0.9973 | 0.9941 | 0.115 | 59.75 |
| 1 | 54123267 | rs1405479 | G | A | 2 | 0.8637 | 0.8637 | 0.9958 | 0.9958 | 0.6768 | 0 |
| 1 | 53492437 | rs6588454 | A | T | 2 | 0.8642 | 0.8642 | 1.0058 | 1.0058 | 0.6826 | 0 |
| 1 | 53564656 | rs1046582 | G | A | 2 | 0.8644 | 0.7414 | 1.0029 | 1.0106 | 0.0806 | 67.25 |
| 1 | 53494069 | rs7290314 | T | G | 2 | 0.8647 | 0.8647 | 1.0058 | 1.0058 | 0.6809 | 0 |
| 1 | 53551619 | rs1157996 | G | A | 2 | 0.8647 | 0.7747 | 1.0033 | 1.0079 | 0.1762 | 45.35 |
| 1 | 53545533 | rs1120608 | C | T | 2 | 0.8648 | 0.8173 | 1.0033 | 1.0053 | 0.2547 | 22.93 |
| 1 | 53506250 | rs7520302 | A | G | 2 | 0.8649 | 0.8649 | 0.9941 | 0.9941 | 0.7044 | 0 |
| 1 | 53564637 | rs1046582 | A | G | 2 | 0.8651 | 0.7407 | 1.0029 | 1.0107 | 0.079 | 67.59 |
| 1 | 53509581 | rs6042662 | G | A | 2 | 0.8652 | 0.8652 | 0.9941 | 0.9941 | 0.738 | 0 |
| 1 | 53564605 | rs1046582 | C | T | 2 | 0.8655 | 0.7408 | 1.0029 | 1.0107 | 0.0791 | 67.57 |
| 1 | 53557612 | rs1288405 | A | C | 2 | 0.866 | 0.9911 | 1.0032 | 1.0003 | 0.2205 | 33.38 |
| 1 | 53633554 | rs5565535 | A | G | 2 | 0.8662 | 0.8662 | 0.9967 | 0.9967 | 0.9518 | 0 |
| 1 | 53618841 | rs1088877 | A | C | 2 | 0.8666 | 0.8666 | 0.9976 | 0.9976 | 0.6686 | 0 |
| 1 | 53799050 | rs7689711 | C | T | 2 | 0.8667 | 0.8667 | 1.0061 | 1.0061 | 0.9722 | 0 |
| 1 | 53570972 | rs7422401 | A | C | 2 | 0.8669 | 0.8669 | 1.0054 | 1.0054 | 0.4473 | 0 |
| 1 | 53564686 | rs1046582 | T | C | 2 | 0.8671 | 0.7417 | 1.0029 | 1.0106 | 0.0785 | 67.69 |
| 1 | 53563744 | rs3502428 | T | C | 2 | 0.8673 | 0.7427 | 1.0029 | 1.0105 | 0.0812 | 67.11 |
| 1 | 53549039 | rs1001512 | G | A | 2 | 0.8675 | 0.8675 | 0.9955 | 0.9955 | 0.7085 | 0 |
| 1 | 54580585 | rs7290621 | T | A | 2 | 0.8675 | 0.8892 | 0.9899 | 0.9887 | 0.1832 | 43.54 |
| 1 | 53798393 | rs7532838 | A | G | 2 | 0.8676 | 0.8676 | 0.9947 | 0.9947 | 0.8878 | 0 |
| 1 | 53900967 | rs6678150 | G | A | 2 | 0.8677 | 0.8677 | 0.9971 | 0.9971 | 0.3987 | 0 |
| 1 | 53608425 | rs3441827 | C | A | 2 | 0.8679 | 0.8979 | 1.0025 | 1.0021 | 0.2845 | 12.69 |
| 1 | 53502660 | rs7290318 | A | G | 2 | 0.869 | 0.869 | 1.0057 | 1.0057 | 0.6419 | 0 |
| 1 | 53508707 | rs6588456 | C | T | 2 | 0.8691 | 0.8691 | 0.9943 | 0.9943 | 0.6844 | 0 |
| 1 | 54386402 | rs1088882 | G | T | 2 | 0.8691 | 0.8736 | 1.0033 | 1.0032 | 0.3128 | 1.83 |
| 1 | 53852154 | rs1710842 | G | A | 2 | 0.8694 | 0.8694 | 1.0157 | 1.0157 | 0.3296 | 0 |

| | | | | | | | | | | | |
|---|---|---|---|---|---|---|---|---|---|---|---|
| 1 | 54331763 | rs1883453 | A | G | 2 | 0.87 | 0.87 | 1.0022 | 1.0022 | 0.5486 | 0 |
| 1 | 54373313 | rs1120624 | A | G | 2 | 0.8709 | 0.8366 | 0.9968 | 0.9948 | 0.2043 | 37.94 |
| 1 | 54639490 | rs1738996 | G | A | 2 | 0.872 | 0.9446 | 1.0067 | 0.9939 | 0.0356 | 77.35 |
| 1 | 53503996 | rs1710771 | C | A | 2 | 0.8721 | 0.8721 | 0.9945 | 0.9945 | 0.6884 | 0 |
| 1 | 53558270 | rs1288406 | A | G | 2 | 0.8721 | 0.9978 | 1.0031 | 1.0001 | 0.2191 | 33.79 |
| 1 | 53504307 | rs7290319 | G | A | 2 | 0.8725 | 0.8725 | 0.9945 | 0.9945 | 0.6888 | 0 |
| 1 | 53505490 | rs7290319 | A | G | 2 | 0.8725 | 0.8725 | 0.9945 | 0.9945 | 0.6877 | 0 |
| 1 | 53558792 | rs1288408 | G | A | 2 | 0.8728 | 0.9927 | 1.0027 | 0.9998 | 0.1895 | 41.9 |
| 1 | 53474202 | rs6658468 | A | G | 2 | 0.8739 | 0.8739 | 1.0059 | 1.0059 | 0.8174 | 0 |
| 1 | 53904040 | rs1288597 | A | G | 2 | 0.8751 | 0.8751 | 0.9973 | 0.9973 | 0.5323 | 0 |
| 1 | 53507681 | rs6682799 | G | A | 2 | 0.8753 | 0.8753 | 0.9946 | 0.9946 | 0.6903 | 0 |
| 1 | 53756892 | rs8013268 | C | T | 2 | 0.8766 | 0.8734 | 1.0038 | 0.994 | 0.1864 | 42.71 |
| 1 | 53558521 | rs1214050 | C | T | 2 | 0.8768 | 0.8768 | 0.9957 | 0.9957 | 0.7707 | 0 |
| 1 | 53747126 | rs1240210 | G | A | 2 | 0.8768 | 0.8768 | 1.0021 | 1.0021 | 0.8748 | 0 |
| 1 | 53564146 | rs1272548 | A | G | 2 | 0.8775 | 0.768 | 1.003 | 1.0088 | 0.1521 | 51.24 |
| 1 | 54349366 | rs1018402 | T | A | 2 | 0.8775 | 0.8775 | 1.0021 | 1.0021 | 0.3607 | 0 |
| 1 | 54350292 | rs7515771 | A | T | 2 | 0.8776 | 0.8776 | 1.0021 | 1.0021 | 0.3205 | 0 |
| 1 | 54013084 | rs1202918 | G | A | 2 | 0.8777 | 0.8916 | 0.9928 | 0.9934 | 0.3015 | 6.34 |
| 1 | 53502952 | rs7290318 | C | T | 2 | 0.8779 | 0.8779 | 0.9947 | 0.9947 | 0.6835 | 0 |
| 1 | 53505807 | rs7290319 | C | T | 2 | 0.8783 | 0.8783 | 0.9948 | 0.9948 | 0.693 | 0 |
| 1 | 54222119 | rs1181193 | A | G | 2 | 0.8787 | 0.8787 | 0.9978 | 0.9978 | 0.4694 | 0 |
| 1 | 54381406 | rs1206377 | C | T | 2 | 0.8787 | 0.8338 | 0.997 | 0.9941 | 0.1543 | 50.72 |
| 1 | 53482662 | rs1710757 | A | C | 2 | 0.8794 | 0.8794 | 1.0052 | 1.0052 | 0.686 | 0 |
| 1 | 54350164 | rs7517900 | T | C | 2 | 0.8795 | 0.8795 | 1.0021 | 1.0021 | 0.3197 | 0 |
| 1 | 54646167 | rs1205742 | T | C | 2 | 0.8797 | 0.7608 | 1.0083 | 0.9634 | 0.0479 | 74.44 |
| 1 | 53528502 | rs7798021 | C | T | 2 | 0.8798 | 0.8283 | 1.0047 | 1.0108 | 0.1158 | 59.56 |
| 1 | 53612165 | rs7165498 | G | A | 2 | 0.8799 | 0.945 | 1.0034 | 1.0018 | 0.241 | 27.25 |
| 1 | 53535859 | rs2275451 | T | C | 2 | 0.8819 | 0.8267 | 1.0048 | 1.0108 | 0.1325 | 55.82 |
| 1 | 53802995 | rs1500473 | T | C | 2 | 0.8821 | 0.8821 | 1.0056 | 1.0056 | 0.86 | 0 |
| 1 | 54349600 | rs2015011 | T | C | 2 | 0.8821 | 0.8821 | 1.002 | 1.002 | 0.3233 | 0 |
| 1 | 53775786 | rs1710827 | T | A | 2 | 0.8829 | 0.435 | 0.9971 | 1.1111 | 0.0017 | 89.88 |
| 1 | 53559247 | rs1204521 | C | T | 2 | 0.8831 | 0.8831 | 0.9959 | 0.9959 | 0.7722 | 0 |
| 1 | 54361398 | rs1088881 | T | C | 2 | 0.8838 | 0.8838 | 0.998 | 0.998 | 0.4165 | 0 |
| 1 | 53622016 | rs1288346 | C | T | 2 | 0.8839 | 0.8839 | 1.0032 | 1.0032 | 0.399 | 0 |
| 1 | 54533644 | rs7266413 | A | G | 2 | 0.884 | 0.884 | 0.9928 | 0.9928 | 0.9228 | 0 |
| 1 | 54594629 | rs1207384 | A | G | 2 | 0.8842 | 0.8842 | 1.006 | 1.006 | 0.3817 | 0 |
| 1 | 53623664 | rs1288349 | C | T | 2 | 0.8846 | 0.8846 | 1.0032 | 1.0032 | 0.3817 | 0 |
| 1 | 53557644 | rs1208912 | C | T | 2 | 0.8852 | 0.9338 | 1.006 | 1.0043 | 0.2192 | 33.77 |
| 1 | 53884457 | rs1776421 | C | T | 2 | 0.8862 | 0.8751 | 1.0025 | 1.0029 | 0.2961 | 8.4 |
| 1 | 53572862 | rs6664572 | C | A | 2 | 0.8863 | 0.8863 | 0.9973 | 0.9973 | 0.8098 | 0 |
| 1 | 53563195 | rs3766769 | G | A | 2 | 0.8865 | 0.8174 | 1.0044 | 1.0091 | 0.215 | 34.95 |
| 1 | 53548986 | rs1015794 | C | G | 2 | 0.8871 | 0.8871 | 1.0024 | 1.0024 | 0.424 | 0 |
| 1 | 53554406 | rs1158358 | T | C | 2 | 0.8871 | 0.7825 | 1.0027 | 1.0079 | 0.162 | 48.87 |
| 1 | 54016511 | rs4927004 | G | A | 2 | 0.8876 | 0.8876 | 1.007 | 1.007 | 0.4484 | 0 |
| 1 | 53502884 | rs1710770 | C | G | 2 | 0.888 | 0.888 | 0.9952 | 0.9952 | 0.6739 | 0 |
| 1 | 53563754 | rs3563163 | C | T | 2 | 0.8881 | 0.7744 | 1.0028 | 1.0086 | 0.1514 | 51.42 |
| 1 | 53738419 | rs4129475 | C | T | 2 | 0.8891 | 0.8891 | 1.003 | 1.003 | 0.752 | 0 |
| 1 | 53628559 | rs7914920 | G | A | 2 | 0.8895 | 0.8895 | 1.0036 | 1.0036 | 0.6197 | 0 |

| | | | | | | | | | | | |
|---|---|---|---|---|---|---|---|---|---|---|---|
| 1 | 53746595 | rs2297657 | A | T | 2 | 0.8895 | 0.8895 | 1.002 | 1.002 | 0.7201 | 0 |
| 1 | 54677401 | rs3766459 | G | A | 2 | 0.8897 | 0.8897 | 1.0035 | 1.0035 | 0.5202 | 0 |
| 1 | 54674473 | rs7544763 | A | G | 2 | 0.8898 | 0.8898 | 1.0035 | 1.0035 | 0.5603 | 0 |
| 1 | 53601392 | rs1240209 | G | A | 2 | 0.89 | 0.89 | 1.0041 | 1.0041 | 0.3763 | 0 |
| 1 | 53700170 | rs6588472 | T | C | 2 | 0.89 | 0.89 | 1.002 | 1.002 | 0.4322 | 0 |
| 1 | 53542077 | rs7524881 | T | C | 2 | 0.8901 | 0.8901 | 1.0025 | 1.0025 | 0.492 | 0 |
| 1 | 54566899 | rs1120628 | T | C | 2 | 0.8904 | 0.8904 | 1.0097 | 1.0097 | 0.5554 | 0 |
| 1 | 53568144 | rs1120609 | G | A | 2 | 0.8906 | 0.8906 | 0.9961 | 0.9961 | 0.8719 | 0 |
| 1 | 53901236 | rs6703329 | C | A | 2 | 0.8912 | 0.8912 | 0.9976 | 0.9976 | 0.5178 | 0 |
| 1 | 53560171 | rs7480776 | G | A | 2 | 0.8917 | 0.9517 | 1.0056 | 1.0033 | 0.1896 | 41.89 |
| 1 | 54594263 | rs6679049 | T | C | 2 | 0.8917 | 0.8917 | 0.9946 | 0.9946 | 0.5335 | 0 |
| 1 | 53563783 | rs3521302 | C | T | 2 | 0.8922 | 0.7765 | 1.0027 | 1.0085 | 0.1502 | 51.69 |
| 1 | 54686274 | rs4130915 | C | T | 2 | 0.8922 | 0.8922 | 1.0046 | 1.0046 | 0.6293 | 0 |
| 1 | 53688076 | rs7883811 | C | T | 2 | 0.8927 | 0.8991 | 0.9862 | 0.9799 | 0.12 | 58.62 |
| 1 | 54328025 | rs6588492 | G | A | 2 | 0.8931 | 0.8931 | 0.9982 | 0.9982 | 0.3551 | 0 |
| 1 | 54588229 | rs7530710 | T | C | 2 | 0.8932 | 0.8932 | 0.9982 | 0.9982 | 0.9716 | 0 |
| 1 | 53566137 | rs2854695 | A | T | 2 | 0.8942 | 0.8942 | 0.9963 | 0.9963 | 0.7609 | 0 |
| 1 | 54672074 | rs1120630 | A | C | 2 | 0.8944 | 0.8944 | 1.0033 | 1.0033 | 0.5607 | 0 |
| 1 | 54318438 | rs7266234 | T | C | 2 | 0.8946 | 0.8946 | 0.9976 | 0.9976 | 0.6362 | 0 |
| 1 | 53611590 | rs7267088 | T | A | 2 | 0.895 | 0.895 | 1.0023 | 1.0023 | 0.7077 | 0 |
| 1 | 53761809 | rs1288498 | A | T | 2 | 0.8958 | 0.5053 | 1.0031 | 0.929 | 0.0031 | 88.55 |
| 1 | 53533756 | rs7290517 | T | C | 2 | 0.8961 | 0.8513 | 1.0038 | 1.0073 | 0.1863 | 42.75 |
| 1 | 53564255 | rs3766773 | G | A | 2 | 0.8962 | 0.8962 | 0.9963 | 0.9963 | 0.7836 | 0 |
| 1 | 54407343 | rs7451049 | T | C | 2 | 0.8962 | 0.9655 | 0.9933 | 1.0029 | 0.2131 | 35.49 |
| 1 | 53598630 | rs931158 | G | A | 2 | 0.8963 | 0.9745 | 1.0019 | 0.9993 | 0.13 | 56.37 |
| 1 | 53901655 | rs1288593 | C | T | 2 | 0.8966 | 0.8966 | 0.9977 | 0.9977 | 0.5167 | 0 |
| 1 | 53614255 | rs5944220 | T | C | 2 | 0.897 | 0.897 | 1.0022 | 1.0022 | 0.7056 | 0 |
| 1 | 54123279 | rs1405481 | A | G | 2 | 0.8984 | 0.8984 | 0.9968 | 0.9968 | 0.6821 | 0 |
| 1 | 54696743 | rs4061073 | A | G | 2 | 0.8984 | 0.8984 | 1.0019 | 1.0019 | 0.9805 | 0 |
| 1 | 53991542 | rs3013762 | G | A | 2 | 0.8986 | 0.9667 | 1.0048 | 1.002 | 0.1974 | 39.82 |
| 1 | 54300430 | rs3464082 | A | C | 2 | 0.8988 | 0.8988 | 1.0018 | 1.0018 | 0.897 | 0 |
| 1 | 54331285 | rs6661911 | A | G | 2 | 0.8989 | 0.8989 | 0.9983 | 0.9983 | 0.3417 | 0 |
| 1 | 53562802 | rs3605911 | T | C | 2 | 0.8997 | 0.7465 | 1.0022 | 1.0109 | 0.0683 | 69.91 |
| 1 | 54123270 | rs1405480 | C | T | 2 | 0.9 | 0.9 | 0.9969 | 0.9969 | 0.673 | 0 |
| 1 | 53649516 | rs6668221 | A | G | 2 | 0.9019 | 0.9019 | 1.0054 | 1.0054 | 0.756 | 0 |
| 1 | 54606222 | rs7671599 | C | T | 2 | 0.9024 | 0.9024 | 1.0045 | 1.0045 | 0.5135 | 0 |
| 1 | 53804715 | rs5751498 | C | A | 2 | 0.9027 | 0.9027 | 1.0041 | 1.0041 | 0.8538 | 0 |
| 1 | 53874457 | rs1134007 | G | A | 2 | 0.9028 | 0.8194 | 0.9978 | 0.9942 | 0.162 | 48.86 |
| 1 | 53616724 | rs1207494 | G | T | 2 | 0.9032 | 0.9032 | 1.0027 | 1.0027 | 0.3725 | 0 |
| 1 | 53752134 | rs1203592 | A | G | 2 | 0.9034 | 0.9034 | 0.9983 | 0.9983 | 0.8138 | 0 |
| 1 | 53482526 | rs1710757 | T | C | 2 | 0.9037 | 0.9037 | 0.9935 | 0.9935 | 0.6024 | 0 |
| 1 | 53553485 | rs1288399 | A | G | 2 | 0.9038 | 0.924 | 1.002 | 1.0016 | 0.3001 | 6.89 |
| 1 | 54026488 | rs1120617 | A | G | 2 | 0.9042 | 0.9042 | 0.9981 | 0.9981 | 0.498 | 0 |
| 1 | 53599574 | rs7523301 | C | T | 2 | 0.9044 | 0.9044 | 0.9831 | 0.9831 | 0.7282 | 0 |
| 1 | 54674529 | rs7547210 | T | A | 2 | 0.9048 | 0.9048 | 1.003 | 1.003 | 0.5247 | 0 |
| 1 | 53803266 | rs7289745 | C | A | 2 | 0.905 | 0.905 | 1.004 | 1.004 | 0.8848 | 0 |
| 1 | 54382686 | rs7534991 | A | G | 2 | 0.905 | 0.905 | 1.0024 | 1.0024 | 0.3774 | 0 |
| 1 | 53479892 | rs6681542 | T | C | 2 | 0.9056 | 0.9056 | 1.0046 | 1.0046 | 0.9241 | 0 |

| | | | | | | | | | | | |
|---|---|---|---|---|---|---|---|---|---|---|---|
| 1 | 53560376 | rs1300169 | C | T | 2 | 0.9057 | 0.7615 | 0.9977 | 0.9895 | 0.0873 | 65.8 |
| 1 | 53628562 | rs1288339 | C | T | 2 | 0.9061 | 0.9061 | 1.0026 | 1.0026 | 0.3686 | 0 |
| 1 | 54450998 | rs651381 | G | A | 2 | 0.9067 | 0.9067 | 1.0016 | 1.0016 | 0.8797 | 0 |
| 1 | 53630671 | rs1288334 | T | C | 2 | 0.9076 | 0.9076 | 1.0025 | 1.0025 | 0.355 | 0 |
| 1 | 53567618 | rs3766776 | A | C | 2 | 0.9079 | 0.9079 | 0.9976 | 0.9976 | 0.5627 | 0 |
| 1 | 54419730 | rs7266239 | A | G | 2 | 0.908 | 0.9695 | 0.9973 | 1.0012 | 0.1745 | 45.77 |
| 1 | 53804550 | rs5666522 | C | T | 2 | 0.9093 | 0.9093 | 1.0038 | 1.0038 | 0.8382 | 0 |
| 1 | 53627718 | rs1295454 | T | C | 2 | 0.9096 | 0.9096 | 1.0025 | 1.0025 | 0.3681 | 0 |
| 1 | 54345792 | rs6680026 | C | T | 2 | 0.91 | 0.91 | 0.9985 | 0.9985 | 0.3512 | 0 |
| 1 | 54390184 | rs1088882 | C | T | 2 | 0.9103 | 0.9103 | 1.0022 | 1.0022 | 0.3453 | 0 |
| 1 | 54359186 | rs3753419 | A | G | 2 | 0.9104 | 0.9104 | 0.9985 | 0.9985 | 0.432 | 0 |
| 1 | 54549256 | rs7891973 | A | G | 2 | 0.9108 | 0.9108 | 1.0079 | 1.0079 | 0.6506 | 0 |
| 1 | 53619879 | rs1481044 | G | A | 2 | 0.911 | 0.911 | 1.009 | 1.009 | 0.4732 | 0 |
| 1 | 53497421 | rs1207589 | G | A | 2 | 0.9113 | 0.8723 | 1.0032 | 1.0059 | 0.2111 | 36.07 |
| 1 | 53546070 | rs1272353 | A | G | 2 | 0.9113 | 0.9113 | 1.0021 | 1.0021 | 0.3205 | 0 |
| 1 | 53630531 | rs1288335 | A | G | 2 | 0.9116 | 0.966 | 1.0024 | 1.0011 | 0.2492 | 24.69 |
| 1 | 53542793 | rs899972 | G | T | 2 | 0.912 | 0.912 | 0.9969 | 0.9969 | 0.6581 | 0 |
| 1 | 54344282 | rs2272928 | C | A | 2 | 0.9121 | 0.9121 | 0.9985 | 0.9985 | 0.3301 | 0 |
| 1 | 54667039 | rs1739012 | T | C | 2 | 0.9124 | 0.9124 | 1.0037 | 1.0037 | 0.6366 | 0 |
| 1 | 53624435 | rs1288352 | T | C | 2 | 0.9125 | 0.9125 | 1.0024 | 1.0024 | 0.3616 | 0 |
| 1 | 54579276 | rs1488719 | G | T | 2 | 0.913 | 0.913 | 0.995 | 0.995 | 0.64 | 0 |
| 1 | 53755900 | rs7408683 | C | T | 2 | 0.9132 | 0.7979 | 1.0027 | 0.9892 | 0.1416 | 53.71 |
| 1 | 53559310 | rs1120609 | T | C | 2 | 0.9133 | 0.862 | 1.0021 | 0.994 | 0.089 | 65.42 |
| 1 | 53649171 | rs1209477 | A | G | 2 | 0.9134 | 0.9134 | 1.0047 | 1.0047 | 0.7375 | 0 |
| 1 | 53830719 | rs3617850 | G | A | 2 | 0.9143 | 0.9143 | 0.9985 | 0.9985 | 0.3904 | 0 |
| 1 | 53941851 | rs1203690 | C | G | 2 | 0.9143 | 0.8629 | 0.9983 | 0.9963 | 0.1822 | 43.82 |
| 1 | 54658586 | rs1205974 | T | C | 2 | 0.9143 | 0.9143 | 1.0043 | 1.0043 | 0.3978 | 0 |
| 1 | 53580839 | rs1158705 | G | A | 2 | 0.9147 | 0.9147 | 1.0043 | 1.0043 | 0.3556 | 0 |
| 1 | 54386271 | rs6696913 | T | C | 2 | 0.9153 | 0.843 | 0.9979 | 0.9929 | 0.0715 | 69.21 |
| 1 | 53745703 | rs1120613 | C | T | 2 | 0.9162 | 0.9162 | 1.0015 | 1.0015 | 0.7674 | 0 |
| 1 | 53507145 | rs1710772 | T | C | 2 | 0.9164 | 0.9164 | 0.9959 | 0.9959 | 0.6154 | 0 |
| 1 | 54024468 | rs4546869 | C | T | 2 | 0.9165 | 0.9037 | 1.0015 | 0.9964 | 0.0421 | 75.78 |
| 1 | 54586649 | rs1739121 | G | A | 2 | 0.9166 | 0.9166 | 0.9964 | 0.9964 | 0.4889 | 0 |
| 1 | 54201278 | rs797916 | G | A | 2 | 0.9169 | 0.9169 | 1.008 | 1.008 | 0.6553 | 0 |
| 1 | 54255775 | rs1387011 | C | A | 2 | 0.917 | 0.9372 | 0.9925 | 1.0075 | 0.213 | 35.53 |
| 1 | 54693400 | rs1114141 | G | A | 2 | 0.917 | 0.917 | 1.0029 | 1.0029 | 0.8014 | 0 |
| 1 | 53545995 | rs1288394 | G | A | 2 | 0.9175 | 0.9514 | 1.0103 | 1.0068 | 0.2644 | 19.71 |
| 1 | 53569492 | rs4926953 | T | C | 2 | 0.9176 | 0.8093 | 1.002 | 1.0068 | 0.1768 | 45.18 |
| 1 | 54388307 | rs1088882 | A | C | 2 | 0.9176 | 0.9176 | 1.002 | 1.002 | 0.3555 | 0 |
| 1 | 53821463 | rs6118892 | G | T | 2 | 0.9177 | 0.9177 | 0.9982 | 0.9982 | 0.5067 | 0 |
| 1 | 53913080 | rs9436924 | C | T | 2 | 0.9184 | 0.9184 | 0.9983 | 0.9983 | 0.6712 | 0 |
| 1 | 54590677 | rs3594841 | C | G | 2 | 0.9194 | 0.9194 | 1.0042 | 1.0042 | 0.4028 | 0 |
| 1 | 53611641 | rs7267088 | G | A | 2 | 0.92 | 0.92 | 1.0017 | 1.0017 | 0.6749 | 0 |
| 1 | 53495647 | rs1118493 | A | G | 2 | 0.9206 | 0.9206 | 1.0035 | 1.0035 | 0.7222 | 0 |
| 1 | 54583770 | rs5767890 | C | T | 2 | 0.9219 | 0.9312 | 0.9941 | 0.9922 | 0.1283 | 56.77 |
| 1 | 54584786 | rs1329781 | T | C | 2 | 0.9221 | 0.9727 | 1.0056 | 1.0032 | 0.097 | 63.68 |
| 1 | 53619502 | rs1481043 | A | G | 2 | 0.9228 | 0.9228 | 1.0078 | 1.0078 | 0.4717 | 0 |
| 1 | 53621695 | rs1166911 | A | G | 2 | 0.9233 | 0.9233 | 0.9923 | 0.9923 | 0.6358 | 0 |

| | | | | | | | | | | | |
|---|---|---|---|---|---|---|---|---|---|---|---|
| 1 | 54352646 | rs1078896 | A | G | 2 | 0.9236 | 0.9236 | 1.0013 | 1.0013 | 0.3352 | 0 |
| 1 | 54319977 | rs3585957 | G | T | 2 | 0.9237 | 0.9237 | 1.0019 | 1.0019 | 0.7384 | 0 |
| 1 | 53534687 | rs6697308 | C | G | 2 | 0.924 | 0.8806 | 1.0027 | 1.0053 | 0.2318 | 30.06 |
| 1 | 54612610 | rs1482078 | G | A | 2 | 0.9244 | 0.8765 | 1.0033 | 1.0065 | 0.2457 | 25.79 |
| 1 | 53740797 | rs6696068 | G | T | 2 | 0.9247 | 0.9247 | 1.0013 | 1.0013 | 0.4542 | 0 |
| 1 | 53573841 | rs3420370 | G | A | 2 | 0.9249 | 0.9249 | 0.9977 | 0.9977 | 0.5981 | 0 |
| 1 | 53820722 | rs4534330 | A | G | 2 | 0.9249 | 0.9249 | 0.9984 | 0.9984 | 0.4819 | 0 |
| 1 | 54356361 | rs1120623 | C | A | 2 | 0.9251 | 0.9251 | 0.9983 | 0.9983 | 0.7394 | 0 |
| 1 | 54604576 | rs633883 | G | A | 2 | 0.9251 | 0.9251 | 1.0103 | 1.0103 | 0.86 | 0 |
| 1 | 53826895 | rs1088878 | G | A | 2 | 0.9252 | 0.9252 | 1.0013 | 1.0013 | 0.5377 | 0 |
| 1 | 54434184 | rs7535941 | G | T | 2 | 0.9255 | 0.84 | 0.9981 | 0.993 | 0.0985 | 63.37 |
| 1 | 53654167 | rs5775194 | G | T | 2 | 0.9268 | 0.9268 | 1.0034 | 1.0034 | 0.9873 | 0 |
| 1 | 53599650 | rs1205776 | G | A | 2 | 0.9272 | 0.9797 | 1.0013 | 0.9995 | 0.1872 | 42.51 |
| 1 | 54382648 | rs7537364 | T | C | 2 | 0.9275 | 0.9275 | 1.0018 | 1.0018 | 0.3587 | 0 |
| 1 | 54359033 | rs1883454 | T | G | 2 | 0.9285 | 0.9285 | 0.9988 | 0.9988 | 0.4336 | 0 |
| 1 | 53575012 | rs1317433 | T | G | 2 | 0.9294 | 0.9294 | 1.0017 | 1.0017 | 0.441 | 0 |
| 1 | 54348138 | rs1074969 | T | C | 2 | 0.9295 | 0.9295 | 0.9988 | 0.9988 | 0.3324 | 0 |
| 1 | 54698241 | rs6706000 | T | C | 2 | 0.9301 | 0.9301 | 1.0013 | 1.0013 | 0.9393 | 0 |
| 1 | 53743677 | rs1078895 | T | C | 2 | 0.9302 | 0.9302 | 0.9988 | 0.9988 | 0.7289 | 0 |
| 1 | 53600889 | rs1213283 | C | T | 2 | 0.9303 | 0.9303 | 1.0013 | 1.0013 | 0.7762 | 0 |
| 1 | 53618037 | rs5875415 | C | A | 2 | 0.9306 | 0.9306 | 0.9985 | 0.9985 | 0.8064 | 0 |
| 1 | 53487471 | rs6675161 | C | T | 2 | 0.9307 | 0.9307 | 1.003 | 1.003 | 0.8974 | 0 |
| 1 | 54559083 | rs1208687 | A | C | 2 | 0.9309 | 0.9309 | 0.997 | 0.997 | 0.7441 | 0 |
| 1 | 53782500 | rs1288476 | A | G | 2 | 0.931 | 0.931 | 0.9987 | 0.9987 | 0.8588 | 0 |
| 1 | 53779428 | rs1288478 | C | G | 2 | 0.9315 | 0.9315 | 0.9983 | 0.9983 | 0.7226 | 0 |
| 1 | 53509793 | rs7521090 | C | T | 2 | 0.9319 | 0.9319 | 1.0021 | 1.0021 | 0.8001 | 0 |
| 1 | 54136057 | rs1256427 | T | C | 2 | 0.9319 | 0.9319 | 1.0071 | 1.0071 | 0.8677 | 0 |
| 1 | 54480398 | rs1015882 | G | C | 2 | 0.9319 | 0.9319 | 1.0012 | 1.0012 | 0.6496 | 0 |
| 1 | 53510085 | rs7533000 | A | G | 2 | 0.9321 | 0.9321 | 0.9971 | 0.9971 | 0.6113 | 0 |
| 1 | 53510119 | rs7533010 | A | T | 2 | 0.9321 | 0.9321 | 0.9971 | 0.9971 | 0.6113 | 0 |
| 1 | 53607630 | rs1240374 | T | C | 2 | 0.9326 | 0.9743 | 1.0013 | 1.0005 | 0.2627 | 20.28 |
| 1 | 53841484 | rs7289947 | T | C | 2 | 0.9326 | 0.9326 | 1.0056 | 1.0056 | 0.6481 | 0 |
| 1 | 54693290 | rs1387414 | A | T | 2 | 0.9328 | 0.9328 | 1.0022 | 1.0022 | 0.3859 | 0 |
| 1 | 53510822 | rs7529260 | G | A | 2 | 0.9332 | 0.9332 | 0.9971 | 0.9971 | 0.6131 | 0 |
| 1 | 53624338 | rs1288351 | G | A | 2 | 0.9332 | 0.9332 | 1.0018 | 1.0018 | 0.3586 | 0 |
| 1 | 53510937 | rs6588457 | A | G | 2 | 0.9333 | 0.9333 | 0.9971 | 0.9971 | 0.6148 | 0 |
| 1 | 53605310 | rs6648042 | G | C | 2 | 0.9333 | 0.9986 | 1.0012 | 1 | 0.2183 | 34.01 |
| 1 | 53533768 | rs7408241 | T | C | 2 | 0.9336 | 0.8739 | 1.0024 | 1.0063 | 0.1751 | 45.6 |
| 1 | 53904982 | rs1288599 | T | C | 2 | 0.9338 | 0.9338 | 0.9986 | 0.9986 | 0.5602 | 0 |
| 1 | 53562152 | rs3523802 | A | G | 2 | 0.9344 | 0.8316 | 1.0016 | 1.0057 | 0.1966 | 40.02 |
| 1 | 53723495 | rs915191 | A | G | 2 | 0.9346 | 0.9346 | 0.9983 | 0.9983 | 0.6728 | 0 |
| 1 | 53611972 | rs7267089 | G | A | 2 | 0.9347 | 0.9347 | 0.9986 | 0.9986 | 0.8238 | 0 |
| 1 | 53698223 | rs7656973 | T | C | 2 | 0.9351 | 0.9351 | 0.9982 | 0.9982 | 0.5676 | 0 |
| 1 | 54607546 | rs4927061 | A | T | 2 | 0.9356 | 0.9356 | 0.9989 | 0.9989 | 0.6997 | 0 |
| 1 | 54026880 | rs529674 | T | C | 2 | 0.9365 | 0.9365 | 0.9988 | 0.9988 | 0.676 | 0 |
| 1 | 54026891 | rs522441 | T | G | 2 | 0.9371 | 0.9371 | 0.9988 | 0.9988 | 0.6733 | 0 |
| 1 | 54699150 | rs1810199 | T | C | 2 | 0.9375 | 0.9375 | 0.9924 | 0.9924 | 0.7499 | 0 |
| 1 | 53774534 | rs1288487 | T | G | 2 | 0.9376 | 0.9376 | 1.0013 | 1.0013 | 0.6942 | 0 |

| | | | | | | | | | | | |
|---|---|---|---|---|---|---|---|---|---|---|---|
| 1 | 53798052 | rs7532663 | T | C | 2 | 0.9378 | 0.9378 | 1.0022 | 1.0022 | 0.6434 | 0 |
| 1 | 54407521 | rs1207854 | C | T | 2 | 0.9383 | 0.8543 | 0.9985 | 0.9932 | 0.0639 | 70.88 |
| 1 | 53520981 | rs1398574 | G | A | 2 | 0.9391 | 0.9391 | 0.9963 | 0.9963 | 0.5433 | 0 |
| 1 | 54292601 | rs1181185 | A | G | 2 | 0.9395 | 0.9395 | 1.0013 | 1.0013 | 0.8411 | 0 |
| 1 | 53628247 | rs5623274 | G | C | 2 | 0.9398 | 0.9398 | 1.0013 | 1.0013 | 0.8482 | 0 |
| 1 | 54133346 | rs3006906 | A | G | 2 | 0.9399 | 0.9399 | 0.9936 | 0.9936 | 0.3316 | 0 |
| 1 | 53592963 | rs1209755 | G | A | 2 | 0.9402 | 0.9402 | 0.9988 | 0.9988 | 0.4255 | 0 |
| 1 | 53788095 | rs7559133 | G | A | 2 | 0.9404 | 0.9404 | 0.9971 | 0.9971 | 0.5601 | 0 |
| 1 | 54347152 | rs7516269 | G | T | 2 | 0.9405 | 0.9405 | 0.999 | 0.999 | 0.3543 | 0 |
| 1 | 54628686 | rs604559 | G | A | 2 | 0.9412 | 0.9412 | 1.001 | 1.001 | 0.4907 | 0 |
| 1 | 53652786 | rs6156555 | T | C | 2 | 0.9413 | 0.9413 | 1.0027 | 1.0027 | 0.8236 | 0 |
| 1 | 53627785 | rs5604863 | T | G | 2 | 0.9416 | 0.9416 | 1.0013 | 1.0013 | 0.7879 | 0 |
| 1 | 53764072 | rs1288493 | A | G | 2 | 0.9417 | 0.9417 | 0.9987 | 0.9987 | 0.8052 | 0 |
| 1 | 54570262 | rs6699560 | T | C | 2 | 0.942 | 0.942 | 0.9963 | 0.9963 | 0.8719 | 0 |
| 1 | 53615668 | rs8016319 | A | G | 2 | 0.9425 | 0.9425 | 1.0021 | 1.0021 | 0.4488 | 0 |
| 1 | 53510038 | rs7543151 | G | A | 2 | 0.9428 | 0.9428 | 0.9975 | 0.9975 | 0.6015 | 0 |
| 1 | 53670451 | rs1205875 | G | A | 2 | 0.9436 | 0.9514 | 0.9927 | 0.9926 | 0.2338 | 29.44 |
| 1 | 54389471 | rs1206789 | C | A | 2 | 0.9441 | 0.9441 | 1.0014 | 1.0014 | 0.4167 | 0 |
| 1 | 54353730 | rs7526625 | C | A | 2 | 0.9442 | 0.9442 | 1.001 | 1.001 | 0.4829 | 0 |
| 1 | 53762189 | rs1288497 | G | A | 2 | 0.9444 | 0.4994 | 1.0016 | 0.9291 | 0.0035 | 88.28 |
| 1 | 53633492 | rs1274608 | G | A | 2 | 0.9454 | 0.9213 | 0.9983 | 0.9974 | 0.2917 | 10.04 |
| 1 | 53523963 | rs998296 | G | A | 2 | 0.9456 | 0.9456 | 1.0023 | 1.0023 | 0.6013 | 0 |
| 1 | 53643490 | rs1120612 | A | G | 2 | 0.9457 | 0.9457 | 0.9966 | 0.9966 | 0.6027 | 0 |
| 1 | 53513007 | rs1162219 | A | G | 2 | 0.9458 | 0.9458 | 1.0034 | 1.0034 | 0.4789 | 0 |
| 1 | 53568171 | rs1208701 | A | G | 2 | 0.9466 | 0.9257 | 0.9973 | 0.9942 | 0.127 | 57.07 |
| 1 | 54610389 | rs3482691 | C | G | 2 | 0.9467 | 0.887 | 1.0023 | 1.006 | 0.2365 | 28.64 |
| 1 | 54387857 | rs1088882 | T | C | 2 | 0.9468 | 0.8913 | 0.9987 | 0.9965 | 0.2036 | 38.13 |
| 1 | 53818215 | rs1162598 | T | C | 2 | 0.947 | 0.947 | 1.0025 | 1.0025 | 0.9303 | 0 |
| 1 | 53562438 | rs7248062 | C | T | 2 | 0.9476 | 0.8621 | 1.002 | 1.0069 | 0.2114 | 35.98 |
| 1 | 53744996 | rs6698933 | A | T | 2 | 0.9476 | 0.9476 | 0.9986 | 0.9986 | 0.6063 | 0 |
| 1 | 53801853 | rs5928624 | C | G | 2 | 0.9477 | 0.9477 | 1.0022 | 1.0022 | 0.8461 | 0 |
| 1 | 54023894 | rs7545676 | C | T | 2 | 0.9487 | 0.8906 | 1.0009 | 0.9959 | 0.0411 | 76.04 |
| 1 | 53630010 | rs1288336 | G | C | 2 | 0.949 | 0.9988 | 1.0014 | 1 | 0.2475 | 25.23 |
| 1 | 53476902 | rs5634760 | G | A | 2 | 0.9496 | 0.9496 | 1.0021 | 1.0021 | 0.5442 | 0 |
| 1 | 53562344 | rs1275337 | A | G | 2 | 0.9502 | 0.8376 | 1.0012 | 1.0056 | 0.1883 | 42.22 |
| 1 | 54355105 | rs1074969 | A | G | 2 | 0.9502 | 0.9502 | 0.9991 | 0.9991 | 0.4186 | 0 |
| 1 | 53814440 | rs1214115 | C | T | 2 | 0.9512 | 0.9512 | 0.999 | 0.999 | 0.6071 | 0 |
| 1 | 53561878 | rs3443110 | G | A | 2 | 0.9514 | 0.846 | 1.0012 | 1.0051 | 0.1998 | 39.17 |
| 1 | 54387577 | rs1209078 | T | C | 2 | 0.9514 | 0.9514 | 1.0012 | 1.0012 | 0.4181 | 0 |
| 1 | 53597390 | rs6688627 | G | A | 2 | 0.9516 | 0.9888 | 1.0008 | 0.9998 | 0.2344 | 29.29 |
| 1 | 53906597 | rs1295465 | C | A | 2 | 0.9519 | 0.9519 | 0.999 | 0.999 | 0.8305 | 0 |
| 1 | 53572138 | rs4926585 | C | T | 2 | 0.9522 | 0.466 | 0.9985 | 1.1009 | 0.0216 | 81.06 |
| 1 | 53571730 | rs7516241 | C | T | 2 | 0.9528 | 0.9951 | 1.0025 | 0.9996 | 0.0875 | 65.75 |
| 1 | 53562594 | rs1275374 | A | G | 2 | 0.9534 | 0.9471 | 0.9989 | 1.0016 | 0.2324 | 29.89 |
| 1 | 54370306 | rs2294512 | G | A | 2 | 0.9541 | 0.8819 | 1.0009 | 0.9962 | 0.0935 | 64.46 |
| 1 | 53756742 | rs1288504 | A | T | 2 | 0.9544 | 0.9544 | 0.999 | 0.999 | 0.5489 | 0 |
| 1 | 53744671 | rs4926591 | T | C | 2 | 0.9548 | 0.9548 | 1.0012 | 1.0012 | 0.5619 | 0 |
| 1 | 53818963 | rs7165499 | A | G | 2 | 0.9553 | 0.9553 | 0.9991 | 0.9991 | 0.4714 | 0 |

| | | | | | | | | | | | |
|---|---|---|---|---|---|---|---|---|---|---|---|
| 1 | 53593163 | rs1224025 | G | A | 2 | 0.9557 | 0.9557 | 0.9991 | 0.9991 | 0.7486 | 0 |
| 1 | 53625839 | rs1288341 | C | T | 2 | 0.9559 | 0.9559 | 1.0012 | 1.0012 | 0.3538 | 0 |
| 1 | 53562613 | rs1275388 | C | T | 2 | 0.9564 | 0.9455 | 0.9989 | 1.0017 | 0.2331 | 29.66 |
| 1 | 54614352 | rs1158549 | T | A | 2 | 0.9568 | 0.9568 | 1.0019 | 1.0019 | 0.3359 | 0 |
| 1 | 53583707 | rs1387688 | A | C | 2 | 0.9574 | 0.9917 | 1.002 | 1.0004 | 0.266 | 19.19 |
| 1 | 53628121 | rs7267089 | T | C | 2 | 0.9585 | 0.9585 | 0.9991 | 0.9991 | 0.8279 | 0 |
| 1 | 54069949 | rs2950255 | T | C | 2 | 0.9587 | 0.9587 | 1.0041 | 1.0041 | 0.5058 | 0 |
| 1 | 53802466 | rs1206674 | T | C | 2 | 0.96 | 0.96 | 1.0017 | 1.0017 | 0.9607 | 0 |
| 1 | 53902166 | rs1288594 | T | A | 2 | 0.9601 | 0.9601 | 1.0009 | 1.0009 | 0.7068 | 0 |
| 1 | 53803047 | rs1114717 | G | A | 2 | 0.9616 | 0.9616 | 1.0016 | 1.0016 | 0.9567 | 0 |
| 1 | 54381718 | rs1120624 | T | C | 2 | 0.9616 | 0.9208 | 0.9991 | 0.9977 | 0.2379 | 28.22 |
| 1 | 54352019 | rs1078896 | T | G | 2 | 0.962 | 0.962 | 1.0007 | 1.0007 | 0.3187 | 0 |
| 1 | 53757677 | rs1288500 | C | A | 2 | 0.9624 | 0.9624 | 0.9992 | 0.9992 | 0.7843 | 0 |
| 1 | 54408585 | rs1207996 | C | G | 2 | 0.9628 | 0.8665 | 0.9991 | 0.9938 | 0.0638 | 70.88 |
| 1 | 53592830 | rs1120610 | A | G | 2 | 0.9631 | 0.9631 | 0.9992 | 0.9992 | 0.424 | 0 |
| 1 | 53624489 | rs2185292 | C | G | 2 | 0.9631 | 0.9631 | 1.0008 | 1.0008 | 0.7829 | 0 |
| 1 | 53803639 | rs1208330 | C | T | 2 | 0.9631 | 0.9631 | 1.0015 | 1.0015 | 0.9541 | 0 |
| 1 | 54385047 | rs1120624 | C | A | 2 | 0.9637 | 0.9637 | 0.9991 | 0.9991 | 0.3223 | 0 |
| 1 | 53834204 | rs1162022 | G | C | 2 | 0.9638 | 0.9638 | 0.9975 | 0.9975 | 0.4526 | 0 |
| 1 | 54404522 | rs1120625 | C | T | 2 | 0.9638 | 0.9705 | 1.0009 | 1.0007 | 0.3088 | 3.45 |
| 1 | 53803856 | rs1207309 | A | G | 2 | 0.9641 | 0.9641 | 1.0015 | 1.0015 | 0.9527 | 0 |
| 1 | 54608091 | rs7771362 | T | C | 2 | 0.9642 | 0.9642 | 1.0016 | 1.0016 | 0.393 | 0 |
| 1 | 54402114 | rs1207877 | C | A | 2 | 0.965 | 0.9713 | 1.0009 | 1.0007 | 0.3092 | 3.28 |
| 1 | 53645754 | rs1337627 | T | C | 2 | 0.9666 | 0.929 | 0.9975 | 0.9866 | 0.0129 | 83.84 |
| 1 | 53754891 | rs1710820 | A | G | 2 | 0.9666 | 0.5095 | 1.0007 | 1.0413 | 0.0169 | 82.47 |
| 1 | 54410190 | rs7547247 | T | C | 2 | 0.9666 | 0.9024 | 0.9992 | 0.9967 | 0.1833 | 43.53 |
| 1 | 54163427 | rs1157397 | C | T | 2 | 0.9675 | 0.9675 | 1.0013 | 1.0013 | 0.7868 | 0 |
| 1 | 53755911 | rs1288509 | G | C | 2 | 0.9677 | 0.5367 | 1.0008 | 0.9536 | 0.0039 | 88.02 |
| 1 | 53567788 | rs1120609 | T | C | 2 | 0.9683 | 0.988 | 1.0016 | 0.9991 | 0.1659 | 47.91 |
| 1 | 53602832 | rs7165498 | G | A | 2 | 0.9685 | 0.9652 | 1.0006 | 0.9992 | 0.2167 | 34.49 |
| 1 | 54222571 | rs1183394 | A | G | 2 | 0.9685 | 0.9685 | 0.9992 | 0.9992 | 0.6053 | 0 |
| 1 | 54386643 | rs1209705 | T | C | 2 | 0.9688 | 0.9206 | 0.9992 | 0.9976 | 0.2251 | 32.03 |
| 1 | 53911812 | rs1120616 | G | C | 2 | 0.9692 | 0.9692 | 0.9994 | 0.9994 | 0.7446 | 0 |
| 1 | 54567818 | rs5884829 | C | T | 2 | 0.9692 | 0.9692 | 0.9976 | 0.9976 | 0.6941 | 0 |
| 1 | 54382524 | rs7545015 | G | A | 2 | 0.9696 | 0.9596 | 0.9992 | 0.999 | 0.3013 | 6.41 |
| 1 | 53624849 | rs5569149 | G | C | 2 | 0.9698 | 0.9698 | 0.9993 | 0.9993 | 0.9268 | 0 |
| 1 | 53548957 | rs1001513 | C | T | 2 | 0.9699 | 0.9699 | 0.9989 | 0.9989 | 0.6193 | 0 |
| 1 | 54472145 | rs7658924 | C | A | 2 | 0.9701 | 0.8418 | 0.9981 | 0.986 | 0.2032 | 38.25 |
| 1 | 53605013 | rs3503600 | C | T | 2 | 0.9702 | 0.9645 | 1.0006 | 0.9992 | 0.2103 | 36.29 |
| 1 | 53570659 | rs1120610 | G | A | 2 | 0.9712 | 0.9518 | 0.9985 | 0.9967 | 0.1833 | 43.52 |
| 1 | 53532006 | rs7939292 | T | A | 2 | 0.9714 | 0.8666 | 1.0011 | 1.0085 | 0.1184 | 58.99 |
| 1 | 54210413 | rs808859 | A | G | 2 | 0.9718 | 0.9718 | 1.003 | 1.003 | 0.7337 | 0 |
| 1 | 53597671 | rs6688914 | G | A | 2 | 0.9721 | 0.9599 | 1.0005 | 0.9991 | 0.214 | 35.23 |
| 1 | 53609303 | rs5760603 | A | G | 2 | 0.9721 | 0.9721 | 1.0006 | 1.0006 | 0.7496 | 0 |
| 1 | 54354165 | rs926456 | A | G | 2 | 0.9721 | 0.9721 | 0.9995 | 0.9995 | 0.4127 | 0 |
| 1 | 53477526 | rs7455693 | C | G | 2 | 0.9725 | 0.9725 | 1.0008 | 1.0008 | 0.5251 | 0 |
| 1 | 53569399 | rs1288420 | A | G | 2 | 0.9725 | 0.8627 | 0.9993 | 0.9954 | 0.1981 | 39.64 |
| 1 | 54254069 | rs5587273 | G | A | 2 | 0.9726 | 0.9487 | 0.997 | 1.0126 | 0.0262 | 79.78 |

| | | | | | | | | | | | |
|---|---|---|---|---|---|---|---|---|---|---|---|
| 1 | 53624083 | rs1441458 | G | A | 2 | 0.9731 | 0.9731 | 1.001 | 1.001 | 0.4574 | 0 |
| 1 | 53623911 | rs1156433 | T | C | 2 | 0.9732 | 0.9732 | 1.001 | 1.001 | 0.4574 | 0 |
| 1 | 53835125 | rs4926985 | T | C | 2 | 0.9732 | 0.9493 | 0.9995 | 0.999 | 0.2775 | 15.22 |
| 1 | 54149897 | rs3542631 | C | A | 2 | 0.9734 | 0.9734 | 0.9974 | 0.9974 | 0.6686 | 0 |
| 1 | 53596870 | rs3410087 | G | A | 2 | 0.9736 | 0.9377 | 1.0008 | 0.9972 | 0.1406 | 53.95 |
| 1 | 53782540 | rs1288475 | C | T | 2 | 0.9736 | 0.9736 | 1.0005 | 1.0005 | 0.9057 | 0 |
| 1 | 53824168 | rs1088878 | C | A | 2 | 0.9739 | 0.9739 | 0.9995 | 0.9995 | 0.4654 | 0 |
| 1 | 54199537 | rs797914 | C | T | 2 | 0.9739 | 0.9739 | 0.9975 | 0.9975 | 0.5921 | 0 |
| 1 | 53548958 | rs1120609 | G | A | 2 | 0.9755 | 0.8925 | 1.0006 | 1.0033 | 0.235 | 29.11 |
| 1 | 53549041 | rs1001511 | G | A | 2 | 0.9761 | 0.9761 | 0.9992 | 0.9992 | 0.9309 | 0 |
| 1 | 53600833 | rs7165498 | G | A | 2 | 0.9772 | 0.8545 | 0.9993 | 0.9922 | 0.0929 | 64.58 |
| 1 | 54345975 | rs6703639 | G | A | 2 | 0.9772 | 0.9772 | 0.9996 | 0.9996 | 0.3757 | 0 |
| 1 | 54346135 | rs6657508 | G | A | 2 | 0.9772 | 0.9772 | 0.9996 | 0.9996 | 0.3757 | 0 |
| 1 | 53549055 | rs1207009 | T | C | 2 | 0.9775 | 0.8984 | 1.0006 | 1.0031 | 0.2404 | 27.45 |
| 1 | 53816034 | rs1241024 | C | G | 2 | 0.9781 | 0.9781 | 0.9995 | 0.9995 | 0.4979 | 0 |
| 1 | 54385043 | rs1120624 | A | G | 2 | 0.9781 | 0.9238 | 0.9995 | 0.9976 | 0.2129 | 35.55 |
| 1 | 54196443 | rs7877053 | C | T | 2 | 0.9788 | 0.9788 | 0.9984 | 0.9984 | 0.7212 | 0 |
| 1 | 54353823 | rs7526726 | C | T | 2 | 0.9792 | 0.9792 | 0.9996 | 0.9996 | 0.408 | 0 |
| 1 | 54353450 | rs7526351 | C | T | 2 | 0.9799 | 0.9799 | 0.9997 | 0.9997 | 0.4098 | 0 |
| 1 | 53755151 | rs1203557 | T | C | 2 | 0.9806 | 0.5137 | 1.0004 | 1.0405 | 0.0179 | 82.16 |
| 1 | 53553702 | rs1288400 | A | G | 2 | 0.9807 | 0.9807 | 0.998 | 0.998 | 0.5855 | 0 |
| 1 | 53564821 | rs1288419 | T | C | 2 | 0.9813 | 0.9474 | 1.0007 | 0.9977 | 0.2392 | 27.8 |
| 1 | 53746134 | rs1288518 | G | A | 2 | 0.9815 | 0.9815 | 1.0003 | 1.0003 | 0.7226 | 0 |
| 1 | 53514385 | rs7455027 | A | G | 2 | 0.9817 | 0.9817 | 1.0016 | 1.0016 | 0.6631 | 0 |
| 1 | 54256374 | rs5759823 | C | T | 2 | 0.9819 | 0.9291 | 1.002 | 1.0178 | 0.0236 | 80.47 |
| 1 | 53908298 | rs1088879 | A | G | 2 | 0.9823 | 0.9823 | 0.9996 | 0.9996 | 0.7725 | 0 |
| 1 | 54684259 | rs3440590 | A | G | 2 | 0.9826 | 0.8244 | 0.9996 | 0.9899 | 0.016 | 82.78 |
| 1 | 53562821 | rs3412737 | G | C | 2 | 0.9829 | 0.8458 | 1.0004 | 1.0057 | 0.1694 | 47.04 |
| 1 | 54353556 | rs7548285 | G | C | 2 | 0.9833 | 0.9833 | 0.9997 | 0.9997 | 0.4059 | 0 |
| 1 | 53558680 | rs1288407 | C | A | 2 | 0.9834 | 0.9834 | 1.0018 | 1.0018 | 0.6246 | 0 |
| 1 | 53732138 | rs1710817 | C | T | 2 | 0.9834 | 0.9364 | 0.9985 | 0.9935 | 0.2789 | 14.71 |
| 1 | 53802793 | rs5640114 | T | C | 2 | 0.9834 | 0.9834 | 1.0007 | 1.0007 | 0.6104 | 0 |
| 1 | 54396725 | rs1088882 | C | T | 2 | 0.9834 | 0.9244 | 0.9996 | 0.9976 | 0.2063 | 37.39 |
| 1 | 54001468 | rs1130203 | C | A | 2 | 0.9841 | 0.9841 | 0.9975 | 0.9975 | 0.7836 | 0 |
| 1 | 54395488 | rs1120625 | G | C | 2 | 0.9841 | 0.9256 | 0.9996 | 0.9977 | 0.2073 | 37.12 |
| 1 | 53533973 | rs6089190 | A | C | 2 | 0.9843 | 0.9299 | 0.9994 | 1.0035 | 0.177 | 45.12 |
| 1 | 53917677 | rs1078895 | C | T | 2 | 0.9843 | 0.9843 | 0.9997 | 0.9997 | 0.7856 | 0 |
| 1 | 54254367 | rs5566878 | T | C | 2 | 0.9844 | 0.9304 | 1.0017 | 1.0175 | 0.0229 | 80.69 |
| 1 | 54383914 | rs5693445 | A | G | 2 | 0.9846 | 0.9846 | 1.0004 | 1.0004 | 0.4051 | 0 |
| 1 | 53591707 | rs3766795 | T | C | 2 | 0.9848 | 0.9211 | 1.0003 | 1.0022 | 0.2474 | 25.25 |
| 1 | 53620993 | rs1178987 | C | T | 2 | 0.9851 | 0.9851 | 1.0016 | 1.0016 | 0.3851 | 0 |
| 1 | 53949361 | rs1202524 | G | C | 2 | 0.9851 | 0.9851 | 0.9993 | 0.9993 | 0.4705 | 0 |
| 1 | 54246288 | rs7947251 | C | T | 2 | 0.9855 | 0.9855 | 1.0008 | 1.0008 | 0.8368 | 0 |
| 1 | 54336505 | rs1165615 | G | A | 2 | 0.9856 | 0.8641 | 1.0007 | 1.0091 | 0.1885 | 42.18 |
| 1 | 54251581 | rs7876888 | C | T | 2 | 0.986 | 0.9333 | 1.0015 | 1.0168 | 0.0233 | 80.56 |
| 1 | 53567020 | rs3570758 | C | T | 2 | 0.9864 | 0.8055 | 1.0003 | 1.0076 | 0.0937 | 64.41 |
| 1 | 53542533 | rs1077467 | G | T | 2 | 0.9866 | 0.977 | 1.0007 | 0.9983 | 0.1758 | 45.43 |
| 1 | 54382383 | rs1206531 | C | T | 2 | 0.9868 | 0.9868 | 1.0003 | 1.0003 | 0.318 | 0 |

| | | | | | | | | | | | |
|---|---|---|---|---|---|---|---|---|---|---|---|
| 1 | 54421135 | rs6693123 | A | C | 2 | 0.9868 | 0.9868 | 1.0011 | 1.0011 | 0.5673 | 0 |
| 1 | 53510126 | rs5975082 | T | C | 2 | 0.9885 | 0.9885 | 1.001 | 1.001 | 0.6487 | 0 |
| 1 | 53748665 | rs7636221 | A | G | 2 | 0.9887 | 0.9887 | 0.9997 | 0.9997 | 0.7636 | 0 |
| 1 | 54348220 | rs1078896 | C | T | 2 | 0.9887 | 0.9887 | 0.9998 | 0.9998 | 0.3812 | 0 |
| 1 | 54392302 | rs1207217 | C | G | 2 | 0.99 | 0.9332 | 0.9998 | 0.9979 | 0.2134 | 35.4 |
| 1 | 53905062 | rs5692141 | G | A | 2 | 0.9902 | 0.9902 | 1.0011 | 1.0011 | 0.4571 | 0 |
| 1 | 53799298 | rs1120614 | G | A | 2 | 0.9903 | 0.9903 | 1.0003 | 1.0003 | 0.4706 | 0 |
| 1 | 53557352 | rs7599089 | G | C | 2 | 0.9908 | 0.9704 | 1.0005 | 0.9979 | 0.1703 | 46.81 |
| 1 | 54597656 | rs7618382 | G | A | 2 | 0.9912 | 0.9912 | 1.0004 | 1.0004 | 0.6193 | 0 |
| 1 | 53904389 | rs1299818 | G | C | 2 | 0.9914 | 0.9914 | 0.9998 | 0.9998 | 0.6419 | 0 |
| 1 | 54146888 | rs1203085 | T | C | 2 | 0.9919 | 0.9919 | 0.9992 | 0.9992 | 0.6636 | 0 |
| 1 | 54350260 | rs3531451 | G | T | 2 | 0.992 | 0.985 | 1.0002 | 0.9996 | 0.2916 | 10.08 |
| 1 | 53533862 | rs7290517 | G | A | 2 | 0.9921 | 0.9187 | 1.0003 | 1.004 | 0.1888 | 42.11 |
| 1 | 53542620 | rs1077466 | G | A | 2 | 0.9922 | 0.9749 | 1.0004 | 0.9982 | 0.1887 | 42.12 |
| 1 | 53563266 | rs7267086 | C | G | 2 | 0.9923 | 0.9923 | 1.0002 | 1.0002 | 0.8924 | 0 |
| 1 | 53801240 | rs7934106 | C | A | 2 | 0.9937 | 0.9937 | 0.9997 | 0.9997 | 0.948 | 0 |
| 1 | 53857715 | rs1241003 | T | C | 2 | 0.9939 | 0.9939 | 1.0002 | 1.0002 | 0.9099 | 0 |
| 1 | 53901837 | rs1766643 | T | C | 2 | 0.9939 | 0.9939 | 0.9999 | 0.9999 | 0.7557 | 0 |
| 1 | 54399102 | rs1120625 | C | T | 2 | 0.9939 | 0.9976 | 1.0001 | 0.9999 | 0.3053 | 4.85 |
| 1 | 54404892 | rs1180381 | A | G | 2 | 0.994 | 0.9304 | 0.9999 | 0.9978 | 0.2002 | 39.07 |
| 1 | 54623066 | rs1120629 | C | A | 2 | 0.9941 | 0.9767 | 0.9999 | 1.0004 | 0.2769 | 15.41 |
| 1 | 53802054 | rs1206653 | T | C | 2 | 0.9945 | 0.9945 | 1.0003 | 1.0003 | 0.9959 | 0 |
| 1 | 53803824 | rs1206281 | G | A | 2 | 0.9945 | 0.9945 | 0.9998 | 0.9998 | 0.9852 | 0 |
| 1 | 54389200 | rs1004703 | A | G | 2 | 0.9949 | 0.8873 | 0.9999 | 0.9949 | 0.0698 | 69.59 |
| 1 | 53570642 | rs7987401 | G | A | 2 | 0.9954 | 0.9954 | 0.9998 | 0.9998 | 0.7737 | 0 |
| 1 | 53532920 | rs1710781 | C | A | 2 | 0.9955 | 0.8943 | 0.9998 | 1.0066 | 0.1246 | 57.61 |
| 1 | 53552059 | rs6701113 | G | A | 2 | 0.9956 | 0.9956 | 0.9996 | 0.9996 | 0.8139 | 0 |
| 1 | 54415698 | rs1207985 | A | G | 2 | 0.996 | 0.8945 | 1.0001 | 0.9956 | 0.1266 | 57.15 |
| 1 | 53532984 | rs7624898 | T | C | 2 | 0.9961 | 0.8841 | 1.0002 | 1.0074 | 0.1161 | 59.5 |
| 1 | 53532881 | rs7834321 | T | G | 2 | 0.9963 | 0.8968 | 0.9999 | 1.0063 | 0.1309 | 56.18 |
| 1 | 53814744 | rs1273592 | G | A | 2 | 0.9963 | 0.9963 | 0.9999 | 0.9999 | 0.7382 | 0 |
| 1 | 53598126 | rs1088876 | C | T | 2 | 0.9966 | 0.9335 | 0.9999 | 0.9985 | 0.2113 | 36.01 |
| 1 | 53598087 | rs1207666 | C | T | 2 | 0.9973 | 0.9403 | 1 | 0.9987 | 0.2152 | 34.91 |
| 1 | 54145051 | rs1202584 | G | A | 2 | 0.9973 | 0.9973 | 1.0003 | 1.0003 | 0.6596 | 0 |
| 1 | 53588942 | rs3766790 | A | G | 2 | 0.9975 | 0.9975 | 0.9997 | 0.9997 | 0.6916 | 0 |
| 1 | 53911776 | rs1504007 | A | G | 2 | 0.9977 | 0.8017 | 1.0002 | 1.0236 | 0.1367 | 54.85 |
| 1 | 54347583 | rs7553929 | T | A | 2 | 0.9982 | 0.9982 | 1 | 1 | 0.3945 | 0 |
| 1 | 53814232 | rs4926974 | G | A | 2 | 0.9986 | 0.9986 | 1 | 1 | 0.7313 | 0 |
| 1 | 53967690 | rs7290694 | G | A | 2 | 0.9987 | 0.9813 | 1.0001 | 0.9989 | 0.1626 | 48.73 |
| 1 | 53563693 | rs7422401 | C | T | 2 | 0.9988 | 0.9988 | 1 | 1 | 0.424 | 0 |
| 1 | 54361152 | rs7266237 | A | G | 2 | 0.9988 | 0.9988 | 1 | 1 | 0.7549 | 0 |
| 1 | 54694214 | rs2297026 | G | A | 2 | 0.9991 | 0.9991 | 1 | 1 | 0.9815 | 0 |
| 1 | 53618312 | rs1256599 | A | G | 2 | 0.9994 | 0.9994 | 1 | 1 | 0.6562 | 0 |
| 1 | 53545470 | rs1288390 | T | C | 2 | 0.9998 | 0.9998 | 1 | 1 | 0.4325 | 0 |
| 1 | 53858632 | rs1288584 | C | A | 2 | NA | NA | NA | NA | NA | NA |
| 1 | 53937459 | rs1288639 | C | G | 2 | NA | NA | NA | NA | NA | NA |
| 1 | 54637167 | rs625643 | T | C | 2 | NA | NA | NA | NA | NA | NA |
